# Nuclear basket protein ZC3HC1 and its yeast homolog Pml39p feature an evolutionary conserved bimodular construction essential for initial binding to NPC-anchored homologs of scaffold protein TPR

**DOI:** 10.1101/2022.09.10.507405

**Authors:** Philip Gunkel, Haruki Iino, Sandra Krull, Volker C. Cordes

## Abstract

Proteins ZC3HC1 and TPR are construction elements of the nuclear pore complex (NPC)-attached nuclear basket (NB). NB-location of ZC3HC1 depends on TPR already occurring NPC-anchored, whereas additional TPR polypeptides are appended to the NB by ZC3HC1. The current study examined the molecular properties of ZC3HC1 that enable it to bind to the NB and TPR. We report the identification and definition of a nuclear basket-interaction domain (NuBaID) of *Hs*ZC3HC1 comprising two similarly built modules, both essential for the binding to the NB’s NPC-anchored *Hs*TPR. Furthermore, we describe such a bimodular construction as evolutionarily conserved and exemplify the kinship of *Hs*ZC3HC1 by the NB- and *Dd*TPR-interacting homolog of *Dictyostelium discoideum* and by characterizing protein Pml39 as the ZC3HC1 homolog in *Saccharomyces cerevisiae*. Among several properties shared by the different species’ homologs, we unveil the integrity of the bimodular NuBaID of *Sc*Pml39p as being essential for binding to the yeast’s NBs and its TPR homologs *Sc*Mlp1p and *Sc*Mlp2p, and we further present Pml39p as enabling interlinkage of Mlp1p subpopulations. In addition to phyla-specific features, we delineate the three species’ common NuBaID as the characterizing structural entity of a one-of-a-kind protein found not in all but likely most taxa of the eukaryotic realm.

## Introduction

The nuclear basket (NB) is a delicate structure appended to the nuclear side of the nuclear pore complex (NPC), the latter the gateway between cytoplasm and nucleus in eukaryotes. The scaffold of the prototypic NB is composed of thin fibrils attached to the NPC’s nuclear ring from where they first project rectilinearly towards the nuclear interior. They then bifurcate and appear to interconnect with their neighboring fibrils laterally, with the resulting ring-like arrangement commonly called the NB’s terminal ring (TR). Possibly first described as a fish trap-like structure only sporadically perceptible during electron microscopy of ultrathin sections of monkey kidney cells (Maul, 1976), the NB was readily visible as a three-dimensional structure appended to essentially all NPCs of the nuclear envelopes (NEs) of full-grown amphibian and avian oocytes (Ris, 1989, 1991; Jarnik & Aebi, 1991; Goldberg & Allen, 1992; Goldberg *et al*, 1997). Furthermore, NBs appear also well detectable in the salivary gland cells of the midge *Chironomus tentans* (Kiseleva *et al*, 1996), with both the oocytes’ and salivary gland cells’ nuclei characterized by virtually no perinuclear chromatin and with this in turn favorable for the visualization of the NEs’ nuclear side with its NBs in top view. NBs were also detected in the protozoan *Dictyostelium* (Beck et al, 2004), and NB-like structures have been described for budding yeast (Kiseleva *et al*, 2004) and tobacco cells (Fiserova *et al*, 2009) as well, indicating that this structure might be existing in many eukaryotes and different cell types. However, while a variety of different functions have been ascribed to the NB or some of its components in one or the other cell type and species, a universal NB function that holds for all cells remains to be uncovered (e.g., Strambio-De-Castillia et al, 2010; Niepel et al, 2013; Snow & Paschal, 2014; Ashkenazy-Titelman et al, 2020; Bensidoun et al, 2021a).

Furthermore, a consensus on the NB’s complete protein inventory remains pending, even though several proteins have been described over time as components of those intranuclear fibrillar materials that occur attached to the NPCs in different species. Among these proteins is a large, coiled-coil-forming one called TPR in vertebrates (e.g., Cordes *et al*, 1997; Hase *et al*, 2001; Frosst *et al*, 2002; Krull *et al*, 2004) and its homologs in yeast, Mlp1p and Mlp2p (Strambio-de-Castillia *et al*, 1999; Kosova *et al*, 2000). Further homologs of TPR, the abbreviation standing for “Translocated Promotor Region” (Park *et al*, 1986; Mitchell & Cooper, 1992) and the Mlps (Myosin-like proteins; Kölling *et al*, 1993) have been identified in insects and plants, where they have also been named Megator and NUA, respectively (Zimowska *et al*, 1997; Kuznetsov *et al*, 2002; Xu *et al*, 2007; Jacob *et al*, 2007). Eventually, TPR homologs have been described as occurring in a wide range of species all across the eukaryotic realm (Holden *et al*, 2014). The TPR polypeptides of vertebrates and their homologs in insects and yeasts have been proposed and depicted as the backbone-forming elements of the NB’s fibrillar scaffold (Krull *et al*, 2004; Soop *et al*, 2005; Niepel *et al*, 2013; Gunkel *et al*, 2021). Finally, they were also shown to be essential for the NB’s integrity in human cells and budding yeast (Krull *et al*, 2010; Funasaka *et al*, 2012; Niepel *et al*, 2013; Duheron *et al*, 2014). In addition, various proteins have been described over time as binding partners of TPR, the Mlps, or their homologs in the other phyla, and some of them have been found co-localizing with the NPC-attached homologs of TPR and residing at the nuclear envelope (NE) in a TPR-dependent manner. However, these proteins were regarded as neither contributing to the NB’s assembly nor required for maintaining its structural integrity but instead have been considered using the NB as either an operational platform for conducting specific tasks or as a storage place at the NPC (e.g., Zhao *et al*, 2004; Scott *et al*, 2005; Palancade *et al*, 2005; Xu *et al*, 2007; Lee *et al*, 2008; Ding *et al*, 2012; Schweizer *et al*, 2013; Umlauf *et al*, 2013; Aksenova *et al*, 2020; Ouyang *et al*, 2020).

Recently, we have reported the identification of yet another NB-resident protein called ZC3HC1, which occurs positioned at the TR of the prototypic NB and represents yet another binding partner of TPR (Gunkel *et al*, 2021). As for the other NB-appended proteins, such NB residence of ZC3HC1 is TPR-dependent. However, differing from the other TPR-associated NB proteins, we found the NB-attached ZC3HC1 relevant for the NB-positioning of about half the total amount of NE-associated TPR in different cell types (Gunkel *et al*, 2021), with ZC3HC1 itself eventually found functioning as a structural NB component. Following its initial binding to the NPC-anchored TPR polypeptides, ZC3HC1 enables the subsequent recruitment of additional TPR polypeptides and their linkage to TPR already present at the NB (Gunkel & Cordes, 2022). However, it remained to be determined how such a reciprocally dependent positioning of ZC3HC1 and subpopulations of TPR at the NB can come about and which parts of ZC3HC1 enable it to engage in possibly different types of interactions with TPR.

In the current study, we have begun addressing these questions, here focusing on those molecular features and prerequisites of ZC3HC1 that enable its initial binding to the NB. We report a bimodular construction of ZC3HC1, with two similarly built modules, which, though, also exhibit distinctive features. Both modules’ integrity is essential for the protein’s binding to the NB and the TPR subpopulation already anchored to the NPC in a ZC3HC1-independent manner. Furthermore, encompassing the two modules, we present a composite sequence motif that we define as the signature of the nuclear basket interaction domain (NuBaID) of ZC3HC1. We regard such a bimodular NuBaID construction and its signature as the evolutionarily conserved characteristic feature of a unique, one-of-a-kind protein per species in many phyla of the eukaryotic realm. While the different species’ NuBaID-containing proteins often bear little primary sequence resemblance to each other beyond their shared NuBaID signature, we propose them to represent ZC3HC1 homologs nonetheless. We corroborated this conclusion experimentally by demonstrating that two other species’ NuBaID-signature-containing proteins are true homologs of *Hs*ZC3HC1, namely a NuBaID-containing protein of *Dictyostelium discoideum* and the Pml39 protein of *Saccharomyces cerevisiae*, with these proteins together with *Hs*ZC3HC1 markedly exemplifying the diversity between the NuBaID proteins of different phyla. While *Sc*Pml39p had already been found to be an Mlp1/2-binding protein (Palancade *et al*, 2005) and recently stated to be the yeast homolog of *Hs*ZC3HC1 (Gunkel *et al*, 2021; Gunkel & Cordes, 2022), we here present the experimental evidence unveiling that *Sc*Pml39p just like *Hs*ZC3HC1 possesses a bimodular NuBaID and requires integrity of its NuBaID’s two modules for NB- and Mlp-interaction. Moreover, we demonstrate that NB-association of Pml39p is a prerequisite for the positioning of additional amounts of Mlp1p at the yeast’s NBs and that Pml39p even enables interconnections between Mlp1 polypeptides at sites remote from the NB.

In addition, despite their low primary sequence identity and strikingly differing phyla-specific features, we show the different homologs’ kinship further underscored by structural similarities predicted by the artificial intelligence (AI) network-based program AlphaFold2 (Jumper *et al*, 2021; Tunyasuvunakool *et al*, 2021). While this approach also illustrates structural similarities with the target-protein-binding modules of another family of proteins, we finally discuss why ZC3HC1 and its homologs, exemplified by *Hs*ZC3HC1, *Dd*ZC3HC1 and *Sc*Pml39p, stand out as unique.

## Results

### A bimodular NB-interaction domain of ZC3HC1 is essential for its binding to TPR

Recent reports have described ZC3HC1 as an NB-resident protein (Gunkel *et al*, 2021) that interconnects distinct amounts of the NB’s scaffold protein TPR (Gunkel & Cordes, 2022). The initial goal of the at times concomitant study now presented here was to understand how these interactions can come about, at first specifying those parts of ZC3HC1 that enable its initial bonding at the NB and hence to TPR polypeptides positioned there.

For this purpose, we had studied the subcellular distribution of fluorescent protein (FP)-tagged ZC3HC1 mutants in HeLa cells. The suitability of such tags for this purpose had already been attested by experiments in which FP-tagged versions of the full-length intact protein had been ectopically expressed in different cell lines, demonstrating that ZC3HC1 tagged in such a manner was indeed binding preferentially to the NPCs (Gunkel *et al*, 2021). Moreover, complementing fluorescence-loss-in-photobleaching (FLIP) experiments had revealed that such interactions of ZC3HC1 were far longer-lasting than those between the NPC and other proteins that only transiently interact with the NPC but occur locally enriched at this structure in steady-state nonetheless (Supplemental Figure S1).

In the first set of experiments, we stepwise truncated the 502 amino acids (aa) long full-length version of human ZC3HC1, starting from its amino-terminus (NT) and carboxy-terminus (CT). Eventually, this resulted in defining two boundaries, the one close to aa 72, the other near aa 467, beyond which further truncations abolished NB binding entirely (Figure 1A and 1B2; for further ZC3HC1 deletion mutants and their capability of interacting with the NB, complemented by FLIP experiments, see Supplemental Figure S2A). A subsequent series of internal deletions revealed some additional parts we could remove without eliminating the protein’s NB binding capability (Figure 1A and 1B2; Supplemental Figure S2A). Of particular note, we could delete a large segment comprising aa 291–397 without compromising an NE association. Different prediction tools, including PONDR (Predictor of Natural Disordered Regions; e.g., Garner *et al*, 1999), had unanimously predicted most of this segment representing a natively unfolded region without intrinsic order. Furthermore, combining most of the tolerable deletions identified until then, we created a mutant, comprising aa 72–290_398–467, which still turned out capable of binding to the NE (Figure 1A and 1B2).

**Figure 1.**
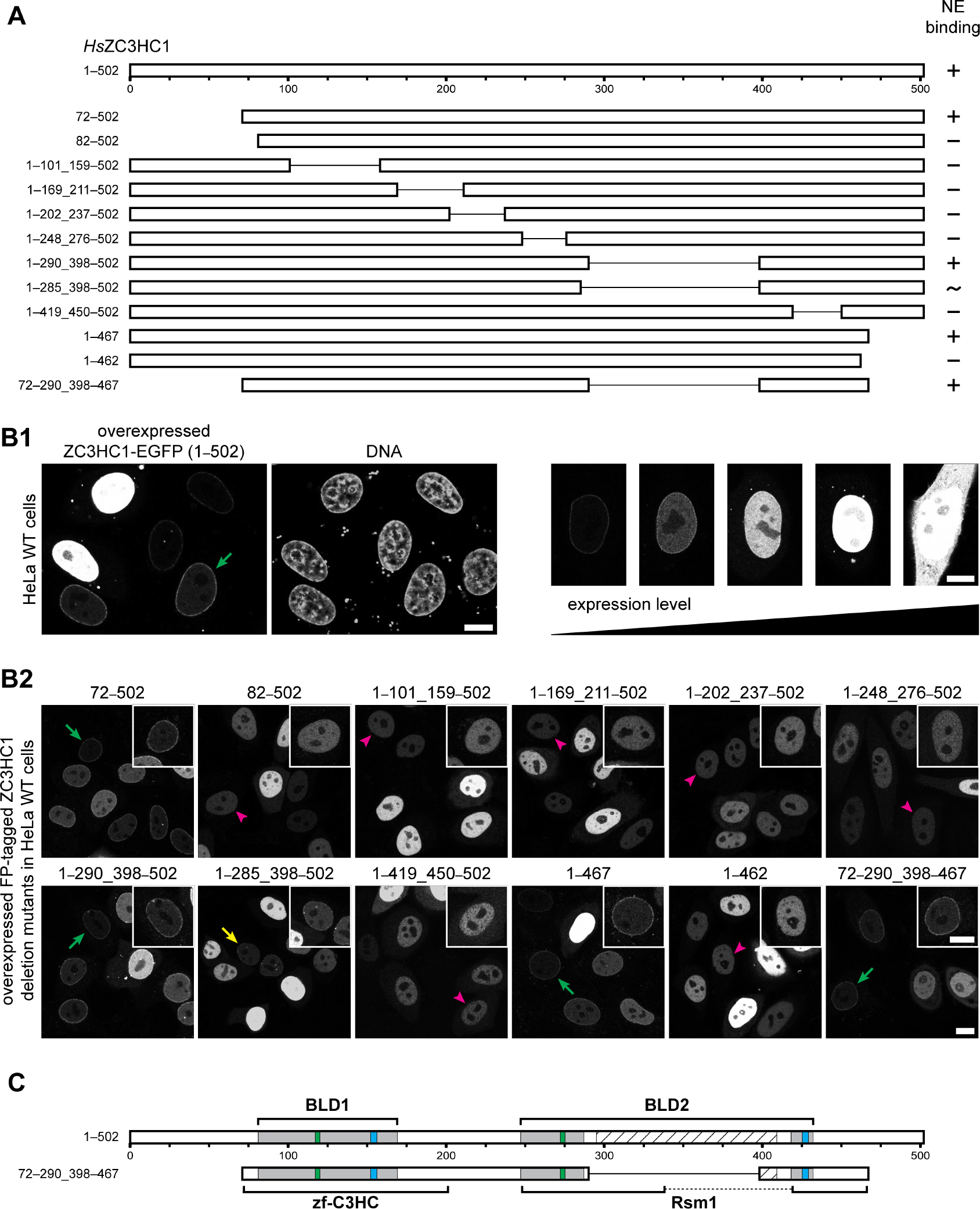
A tandem arrangement of two predicted zinc ion binding modules is essential for the NB association of ZC3HC1. **(A)** Schematic depiction, drawn to scale, of expression vector-encoded human ZC3HC1 and deletion mutants, all containing either an N-terminal EYFP-tag or a C-terminal EGFP-tag, with these tags not shown here. The capability of binding to the NE is indicated (+, binding visible; −, no binding visible; ∼, unclear or only traces of binding visible). Such rating pertained to cells in which the recombinant protein had been expressed in lower amounts, as exemplified in 1B1. **(B)** Fluorescence microscopy of HeLa P2 WT cells transiently transfected with a selection of the expression vectors referred to in 1A. Bars, 10 µm. **(B1)** Exemplifying set of cells in which NE-association of an NB-binding-competent version of *Hs*ZC3HC1, here represented by the WT protein, was only discernible at lower expression levels (one example marked by green arrow). **(B2)** Examples of cells with *Hs*ZC3HC1 mutant proteins ectopically expressed in different amounts. Some exemplary cells exhibiting a low expression level are marked, pointing at either the apparent presence of ZC3HC1 at the NE (green arrows), only trace amounts at the NE (yellow arrow), or absence from there (magenta-colored arrowheads). In addition, the marked cells are shown magnified and signal-enhanced as insets in the upper right corner for each micrograph. Additional data are presented in Supplemental Figure S2A for these and other deletion mutants in HeLa WT and Hela ZC3HC1 KO cells. **(C)** Schematic depiction of the regions encompassing the central parts of two BIR-like domains (BLDs) of ZC3HC1, relative to the simple schemes for the WT and the mutant 72–290_398– 467 shown in 1A. Boxes in green and blue, respectively, represent the positions of C-X_(2)_-C and H-X_(3)_-C sequence elements that, as part of the BLDs, are assumed to be involved in zinc ion coordination. Those regions highlighted in grey represent the BLDs’ initially estimated approximate expanse, as similarly proposed in former studies (Higashi *et al*, 2005; Kokoszynska *et al*, 2008), with the outer boundaries of the *Hs*ZC3HC1 BLD1 defined by R81 and M169, those of BLD2 by T247 and V432, and the inner boundaries of BLD2 by I287 and F418. The hatched box, corresponding to residues D295 to S409, represents an insertion that varies notably in length in the ZC3HC1 homologs of different species and that tools like PONDR predict to be largely disordered except for P322 to S329. Note that other existing segments of *Hs*ZC3HC1 also predicted unstructured by PONDR are not highlighted for reasons of simplification. Areas marked by brackets represent the expanse of regions that comprise Pfam database motifs called zf-C3HC and Rsm1, as specified in the CDD. Note that all ZC3HC1 mutants that lack any part harboring any one of the C-X_(2)_-C or H-X_(3)_-C peptide sequences were incapable of NE-binding (see 1B2).

Inspecting the remaining sequence segments of which much appeared required for NE-association, we then noted that other studies had identified local sequence similarities between the family of inhibitor of apoptosis proteins (IAPs) and ZC3HC1 (Kokoszynska *et al*, 2008), respectively between IAPs and a family of proteins, considered paralogs of the IAPs and called ILPs (IAP-like proteins). Among the latter were *Hs*ILP1 and *Sp*ILP1 (Higashi *et al*, 2005), with the one being actually identically equal to *Hs*ZC3HC1 and the other to *Schizosaccharomyces pombe* Rsm1p, a protein reported involved in RNA export (Yoon, 2004).

The IAPs had already been characterized by at least one, yet mostly several, zinc ion-binding modules called BIR (baculovirus inhibitor of apoptosis protein repeat) domains (Verhagen *et al*, 2001; Silke & Vucic, 2014; Sharma *et al*, 2017). The ILPs, in turn, had been noted to possess two BIR-like domains (BLDs), with certain sequence features resembling those of the BIR and with residues in both BLDs likely to form a zinc ion coordination sphere too.

Furthermore, the first of these two BLDs had also been noted (Kokoszynska *et al*, 2008) to correspond to a zinc finger (zf) motif listed in the Pfam database (Finn *et al*, 2006), where it was called zf-C3HC (http://pfam.xfam.org/family/PF07967). Initially based on sequences representing nine different proteins, of which three were of vertebrate origin and one corresponding to *Sp*Rsm1p, the zf-C3HC motif had already then been described as representing a domain often occurring repeated, with this motif’s initial version then actually deduced from sequence segments representing both BLDs (http://ftp.ebi.ac.uk/pub/databases/Pfam/releases/ Pfam16.0/; further below, see also Supplemental Figure S2D1). On the other hand, a segment of the second BLD of *Hs*ZC3HC1 had later been assigned (http://ftp.ebi.ac.uk/pub/databases/ Pfam/releases/Pfam24.0/; e.g., Finn et al, 2010) the then so-called Rsm1 motif (Finn *et al*, 2008; http://pfam.xfam.org/family/PF08600). The latter had initially been defined by ten non-redundant sequence segments only corresponding to the second BLD, with seven of these segments of fungal origin, again including *Sp*Rsm1 but no mammalian sequences (http://ftp.ebi.ac.uk/pub/databases/Pfam/releases/Pfam20.0/).

The IAPs, by contrast, neither possessed such a zf-C3HC nor an Rsm1 motif. Moreover, they also lacked another feature that distinguished them from the ILP proteins, namely an extensive sequence insertion that appeared to disrupt, in some of the ILP proteins like ILP1/ZC3HC1, the second BLD and the residues potentially forming its zinc finger, with nothing of the sort detectable in the BIR domains (Higashi *et al*, 2005; Kokoszynska *et al*, 2008). Remarkably, this BLD’s sequence insertion largely corresponded to the intrinsically disordered segment we had found dispensable for the appendage of ZC3HC1 to the NE.

When depicting the expanse of these different regions of ZC3HC1 and the positions of its putative zinc fingers, true to scale within a simple scheme of the wild-type (WT) protein and relative to the remaining parts of the still NB-binding-competent ZC3HC1 deletion mutant 72– 290_398–467 (Figure 1C; Supplemental Figure S2A1), it seemed evident that the integrity of both BLDs would be required for NB-binding. All the more because all of those deletion mutants that lacked any part predicted to be directly involved in zinc ion coordination had failed to bind to the NB. Nonetheless, to investigate in further detail whether and to which extent the first and second BLD and its predicted zinc ion-coordinating amino acids, next to some of its other seemingly evolutionarily conserved residues, might contribute to NE association, we created further collections of ZC3HC1 mutants. For such purpose, we initially focused on a minimal sequence signature that we had found identical in both BLDs of the vertebrate homologs of ZC3HC1. While each of the two BLDs also had its distinct, evolutionarily seemingly conserved sequence characteristic (Figure 2A1, and further below), the minimal sequence signature identical in both BLDs’ central part read C-X_(3,5)_-G-W-X_(9,15)_-C-X_(2)_-C-X_(31,152)_-H-X_(3)_-C-X-W (Figure 2A2). This signature, here numbered (1), resembled the ILP family’s minimal sequence signature already described earlier (Higashi *et al*, 2005), except for the first cysteine, characteristic for vertebrates, that preceded the signature’s G-W dipeptide. Apart from a few differences, this early signature also resembled the WebLogos (Crooks *et al*, 2004) that we built (Supplemental Figure S2D1) from those sequences that had been used for the first versions of the Pfam motifs zf-C3HC (http://ftp.ebi.ac.uk/pub/databases/Pfam/releases/ Pfam16.0/) and Rsm1 (http://ftp.ebi.ac.uk/pub/databases/Pfam/releases/Pfam20.0/).

We then altered the vertebrate signature in only one or the other BLD by introducing only a single aa substitution into the full-length and otherwise unchanged ZC3HC1 sequence at a time. In addition, we also substituted a few other single amino acids that are present only within the first BLD or are part of the loop inserted into the second BLD (Figure 2A3, Supplemental Figure S2B1). When we then ectopically expressed these mutants in HeLa cells again, we first observed that each of the solitary C-to-S mutations at positions 117, 120 and 156 in the first BLD, and 272, 275, and 429 in the second BLD completely abolished NE binding of ZC3HC1, as did H-to-A substitutions at positions 152 and 425 in the first and second BLD, respectively (Figure 2B, Supplemental Figure S2B2). In addition, NE binding was abolished by the solitary W-to-A substitutions of the flanking tryptophans at positions 107 and 158 in the first BLD as well as 256 and 431 in the second, when we expressed the corresponding mutants in cells in which the intact endogenous ZC3HC1 was present too (Figure 2B, Supplemental Figure S2B2). Beyond this selection of aa substitutions of high impact, some additional ones, at positions beyond the expanse of the minimal signature outlined above, including, for example, W458A, appeared to affect NE binding notably, yet not abolish it entirely (our unpublished data).

**Figure 2.**
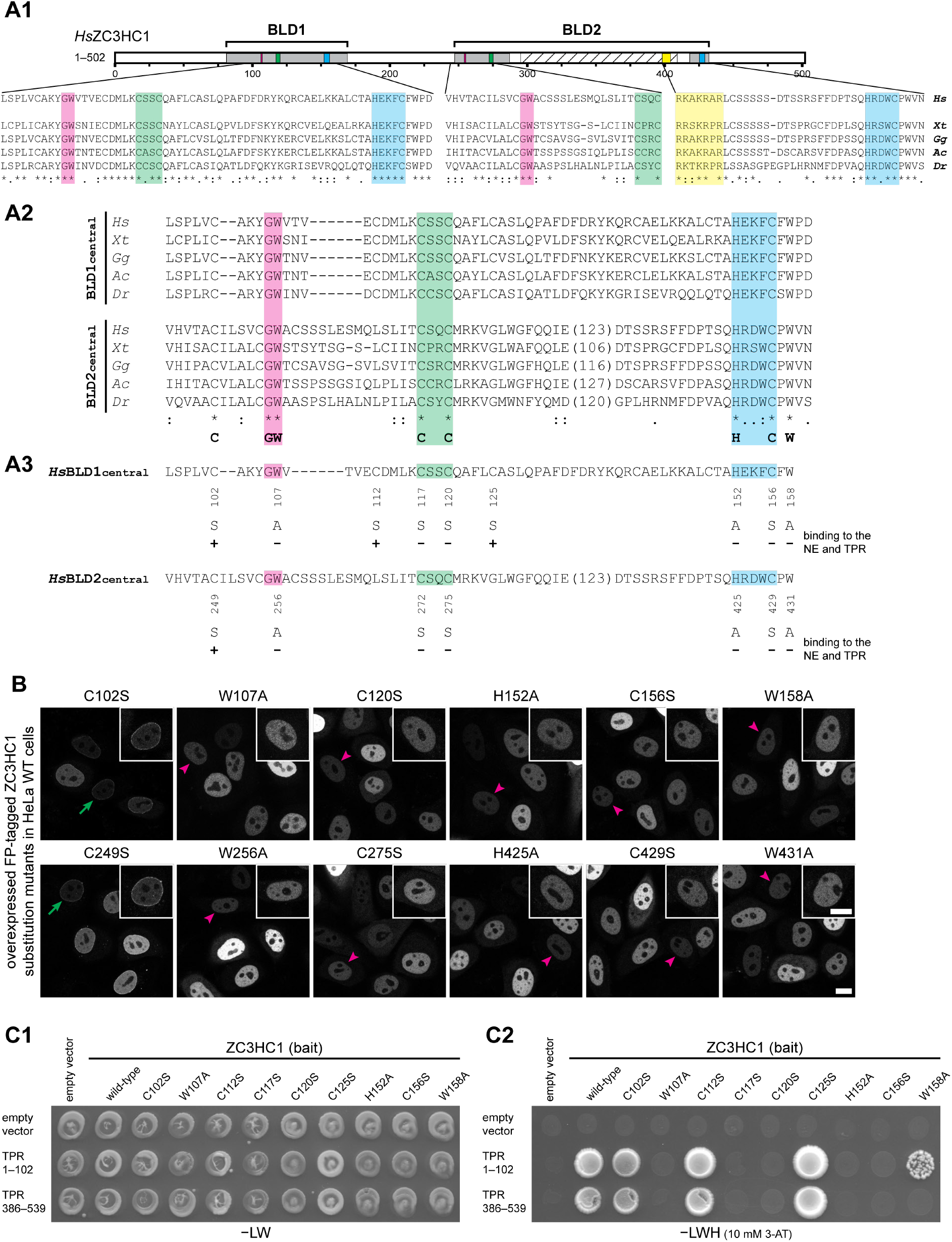
Certain amino acids of ZC3HC1 within both BLDs are essential for enabling interactions with ZC3HC1-binding domains of TPR. **(A)** Schematic depiction of the BLDs with additional sequence features, alignments of representative vertebrate ZC3HC1 homolog sequences and the summarizing depiction of the single amino acid substitution mutants of FP-tagged *Hs*ZC3HC1. **(A1)** Schematic depiction of the two BLDs of *Hs*ZC3HC1, like in Figure 1C, and alignment of the vertebrate homologs’ sequence segments corresponding to each BLD’s minimal central region. We defined the latter as comprising the BLD’s minimal sequence signature, including *i.a.*, the G-W, C-X_(2)_-C, and H-X_(3)_-C peptides, and some additional, flanking residues. Sequences aligned are from the human homolog (*Hs*), from amphibians, here represented by *Xenopus tropicalis* (*Xt*), birds (*Gallus gallus*, *Gg*), reptiles (*Anolis carolinensis*, *Ac*), and fish (*Danio rerio*, *Dr*). Areas highlighted in addition to those in Figure 1C represent the positions of evolutionarily relatively conserved G-W dipeptides (magenta) and that of the protein’s NLS (yellow), with the latter embedded within the largely unstructured insertion in BLD2. **(A2)** Alignments between the vertebrate homologs’ sequences that represent the central part of their first BLD and corresponding parts of their second BLD, yet excluding the sequences of variable length only found inserted into the latter. A minimal sequence signature identical for both BLDs of all these vertebrate homologs is provided in the bottom line. The outer boundaries of the BLD1 central region of *Hs*ZC3HC1 are here defined by L97 and D160 and those of the BLD2 by V244 and N433. The inner boundaries of BLD2, flanking the unstructured insertion, are here corresponding to E288 and D412. **(A3)** Individual amino acid substitutions within the BLD regions and their effects on the mutants’ capabilities of NE binding and interacting with TPR. Those found to be essential or non-essential for such interactions are marked accordingly. **(B)** Fluorescence microscopy of HeLa WT cells transiently transfected with a selection of expression vectors encoding for full-length *Hs*ZC3HC1 mutants, each C-terminally tagged with EGFP and harboring one of the single aa substitutions referred to in 2A3. Exemplary cells exhibiting a low expression level are marked by arrows, indicating presence at the NE (green arrows) or absence from there (magenta-colored arrowheads). In addition, the marked cells are shown magnified and signal-enhanced as insets in the upper right corner for each micrograph. Note that complementing data are presented in Supplemental Figure S2B for these and other substitution mutants (e.g., at aa 363) in HeLa WT and Hela ZC3HC1 KO cells. Bar, 10 µm. **(C)** Single aa substitution mutants of ZC3HC1 studied in Y2H experiments with two TPR segments, each of which includes one of TPR’s distinct ZC3HC1 interaction domains. **(C1)** Representative colony growth, on a selection medium lacking leucine and tryptophan (−LW), of a selection of diploid cells expressing TPR segments together with either WT ZC3HC1 or some of its mutants. **(C2)** Visualization of Y2H interactions, after replica-plating onto selection medium lacking leucine, tryptophan and histidine (−LWH), here supplemented with 10 mM 3-AT. Note that those single aa substitution mutants of ZC3HC1 that did not impair NE association (C120S, C112S or C125S) allowed for colony growth on selective medium −LHW when paired with ZC3HC1-binding domains of TPR. By contrast, no colony growth on such a selective medium was observed when single aa substitution mutants of ZC3HC1 like C156S, which had been found incapable of NE association in HeLa cells, were co-expressed with TPR’s ZC3HC1-binding domains. Note that ZC3HC1 mutant W158A, which was not notably capable of NE-association in the presence of the WT cell’s native ZC3HC1 polypeptides (2B) but readily bound to the NE in ZC3HC1 KO cells (Supplemental Figure S2B2), was found capable of an attenuated interaction with TPR, described in further detail in Supplemental Figure S2C.

Furthermore, since the BIR domains harbored other aromatic residues at those positions corresponding to *Hs*ZC3HC1 W107 and W256 (Higashi *et al*, 2005; Kokoszynska *et al*, 2008), we tested whether other aromatic aa might also be tolerable by ZC3HC1, with this revealing that W107Y, W107F, W256Y, and W256F substitutions had no notable effect on the NE association of such *Hs*ZC3HC1 mutant versions (Supplemental Figure S2C). Furthermore, solitary C-to-S substitutions of other cysteine residues of *Hs*ZC3HC1 conserved in vertebrates and some few other organisms, but again missing in others, as will be shown further below, did not notably abolish binding to the NE either (Figure 2A3, Supplemental Figure S2B). These inconspicuous substitutions included such at positions 102, 112, and 125, and another at position 249. Of these, we regarded especially the one at position 102 as of interest since it represented the cysteine that, together with C117 and C120, was part of the original Pfam zf-C3HC motif’s consensus sequence, comprising the three cysteines eponymic for the C3 within this motif’s name, in addition to the HC based on H152 and C156 (Finn *et al*, 2006; http://ftp.ebi.ac.uk/pub/databases/Pfam/releases/Pfam16.0/; Supplemental Figure S2D1).

Taking the results of our aa substitution mutations and our other findings in vertebrates into account (see also below), including, e.g., the dispensability of C102 of *Hs*ZC3HC1 and the replaceability of W for Y or F, with all this in combination with the published information available until then (Higashi *et al*, 2005; Finn *et al*, 2006; Kokoszynska *et al*, 2008; Finn *et al*, 2008, 2010), we compiled yet another minimal sequence signature. Holding for the central part of both BLDs of several potential ZC3HC1 homologs in different phyla, this signature, here numbered (2), read G-[WYF]-X_(8,72)_-C-X_(2)_-C-X_(15,524)_-H-X_(3)_-C. With two of such simple low stringency signatures arranged in tandem (Supplemental Figure S2D2), this would, in the sequel, allow for already specifically identifying only one ZC3HC1 homolog per species in various organisms, including budding yeast, in which a ZC3HC1 homolog had remained undetectable until then (see further below).

Later, to scrutinize in further detail whether any of the mutants that appeared NE binding-incompetent might possess some residual NE binding potential that we might have formerly overlooked, we also ectopically expressed these ZC3HC1 mutants in ZC3HC1 knockout (KO) cells; once these had become available (Supplemental Figure S2A2, S2B2, S2C2, and S2C3). This way of proceeding later became possible once we had disrupted the *ZC3HC1* alleles in this cell line by CRISPR/Cas9n methodology (Gunkel *et al*, 2021). The advantage of transfecting such KO cells was that the ZC3HC1 mutants did not need to compete for binding sites with the wild-type ZC3HC1. Furthermore, we scrutinized the subcellular distribution of the ectopically expressed deletion mutants and aa-substituted versions of ZC3HC1 by conducting additional FLIP experiments (Supplemental Figure S2A2, S2B2, and S2C3).

Altogether, these experiments corroborated almost all of the abovementioned results initially performed in the ZC3HC1 WT cell line. For example, they allowed for, once again, demonstrating that the likely zinc ion-coordinating residues, namely the total of three cysteines and one histidine residue per BLD, were all essential for NE association. Similarly, the tryptophan residues W107 and W256 at the N-terminal side of the first and second BLD, respectively, appeared similarly important, with again no NE-binding seen with the W256A mutant and only trace amounts of the W107A mutant at the NE, after having photobleached the protein’s soluble nuclear pool. By contrast, the mutants W158A and W431A, both appearing incapable of NE-association in the WT cells, were now clearly NE-associated in the absence of the endogenous ZC3HC1 protein (Supplemental Figure S2B2). These novel findings indicated that W158 and W431, each located at the C-terminal flank of their respective BLD, can support but are not necessarily essential for NE-binding. Later, we would even find that this result explains why a variety of residues present at the corresponding residue positions of apparent ZC3HC1 homologs in other phyla (e.g., Higashi *et al*, 2005; Kokoszynska *et al*, 2008; also see some of the other ZC3HC1 sequences listed further below) are tolerable (our unpublished data). Even more, though, our findings until then already underscored the two BLDs’ commonalities beyond mere sequence similarities, namely by having revealed that signature residues at seemingly invariant positions within both BLDs can also be functionally equivalent.

All in all, the findings till then allowed us to conclude that ZC3HC1 employs for its binding to the NE a distinct tandem arrangement of two similarly constructed binding modules, with both including residues that are likely involved in zinc ion coordination as a prerequisite.

To then scrutinize whether the NE associations that we had found depending on distinct subdomains and amino acids of ZC3HC1 would indeed reflect genuine interactions with protein TPR, we performed yeast two-hybrid (Y2H) experiments. We chose this approach because the Y2H methodology (Fields & Song, 1989) had already allowed for identifying and mapping interaction domains of TPR (e.g., Hase *et al*, 2001; Hase & Cordes, 2003), which later then also included those for ZC3HC1, as will be described in detail elsewhere (Gunkel *et al*, manuscript in preparation). Furthermore, since we here were studying Y2H interactions monitored via *HIS3* gene expression and subsequent colony growth, we additionally challenged these interactions with various concentrations of 3-amino-1,2,4-triazole (3-AT), a competitive inhibitor of the *HIS3* gene product IGP dehydratase (Brennan & Struhl, 1980; Durfee *et al*, 1993).

During these Y2H studies, we tested a collection of the single aa substitution mutants of ZC3HC1 in combination with different TPR regions, each of which included at least one of TPR’s ZC3HC1 interaction domains. While we had initially used TPR segments like, e.g., 1–175 and 172–651 for some of these experiments, all the thereby obtained results (our unpublished data) were later confirmed and corroborated with also smaller TPR segments thereof, comprising, e.g., aa 1–102 and aa 386–539 (Figure 2C; Supplemental Figure S2E1), as well as aa 1–111 and aa 275–539 (Supplemental Figure S2E2). All these segments were part of TPR’s heptad repeat-dominated, coiled coil-forming N-terminal region, called the “rod” domain (e.g., Hase *et al*, 2001). Altogether, these experiments revealed that all point mutations that did not negatively affect binding to the NB also did not prevent a robust Y2H interaction with TPR (Figure 2C, Supplemental Figure S2E). Furthermore, those single aa substitution mutants that had failed to bind to the NB in the presence of the WT protein, but had been found capable of binding in the latter’s absence, were also engaging in Y2H interactions with TPR, though tolerating only lower concentrations of 3-AT (Supplemental Figure S2E). Most strikingly, however, all single aa substitution mutants that had entirely abolished NE association in HeLa cells (Figure 2A3 and 2B), irrespective of whether the competing WT protein had been around or not (Supplemental Figure S2B), were also no longer capable of a Y2H interaction with any of the ZC3HC1-binding segments of TPR (Figure 2C, Supplemental Figure S2E).

Taken as a whole, the strict correlation between each mutant’s performance in the NB-binding and the Y2H experiments with TPR allowed us to conclude that the impaired binding of ZC3HC1 to the NB reflected an impaired interaction with TPR. Moreover, apart from having revealed that only the integrity of distinct sequence features common to both BLDs enables ZC3HC1 to bind to the NB, it was similarly evident that the BLDs are both required for establishing a functional TPR binding interface. We thus regarded the two BLDs, also called BLD1 and BLD2 in the following, as representing an experimentally ascertained functional entity that we from then on referred to as the nuclear basket-interaction domain NuBaID.

### ZC3HC1 homologs, with a NuBaID signature and capable of binding to corresponding TPR homologs, exist as NB proteins in different phyla of the eukaryotic realm

Having realized that a tandem arrangement of the alleged zinc fingers is an earmark of *Hs*ZC3HC1, allowing it to interact with TPR, we wanted to find out whether possessing a NuBaID signature might mark proteins as NB- and TPR-interaction partners also in other phyla, even if not appearing to have other immediately apparent primary sequence features in common. For such data mining, we used complementary approaches, including signature-based and primary sequence end-to-end alignment searches that made it possible to progressively comb the eukaryotic realm’s available sequence data interactively, allowing for iterative refinement of the mining process (see Supplemental Information 1). Eventually, this resulted in identifying numerous potential ZC3HC1 homologs in all eukaryotic supergroups. That is, in the Opisthokonta, Amoebozoa, and Viridiplantae (Figure 3A, see Supplemental List of Sequences for ZC3HC1 Homologs), in other divisions of the Archaeplastida, in the Excavata, in several lineages within the SAR supergroup, and in many other protist groups and genera whose affiliation was still uncertain at the time when conducting such data mining (our unpublished data). Only within some few phyla and classes, like, for example, in Porifera and Cephalochordata (Figure 3A and 3B1), a genuine ZC3HC1 homolog remained undetectable even to date (July 2022), which was also the case for most insect orders (Figure 3B2), the latter in line with ILPs having been reported not detectable in *Drosophila* (Higashi *et al*, 2005). In a class like the Insecta, the loss of recognizable ZC3HC1 homologs appears to have occurred at different time points during insect evolution and the splitting of its lineages leading to its different orders. Here, we found this exemplified by ZC3HC1 homologs not detected in the Paleoptera and most orders of the Neoptera while still present in other Neoptera orders (Figure 3B2). Furthermore, we regard it as of note that in some groups of organisms, like, for example, in the chordates’ subphylum Tunicata, the existing ZC3HC1 homologs appear to be subject to various mutations of which most, at the corresponding positions of the human homolog, would entirely abolish the latter’s ability of binding to the NB and TPR (Figure 3B1).

**Figure 3.**
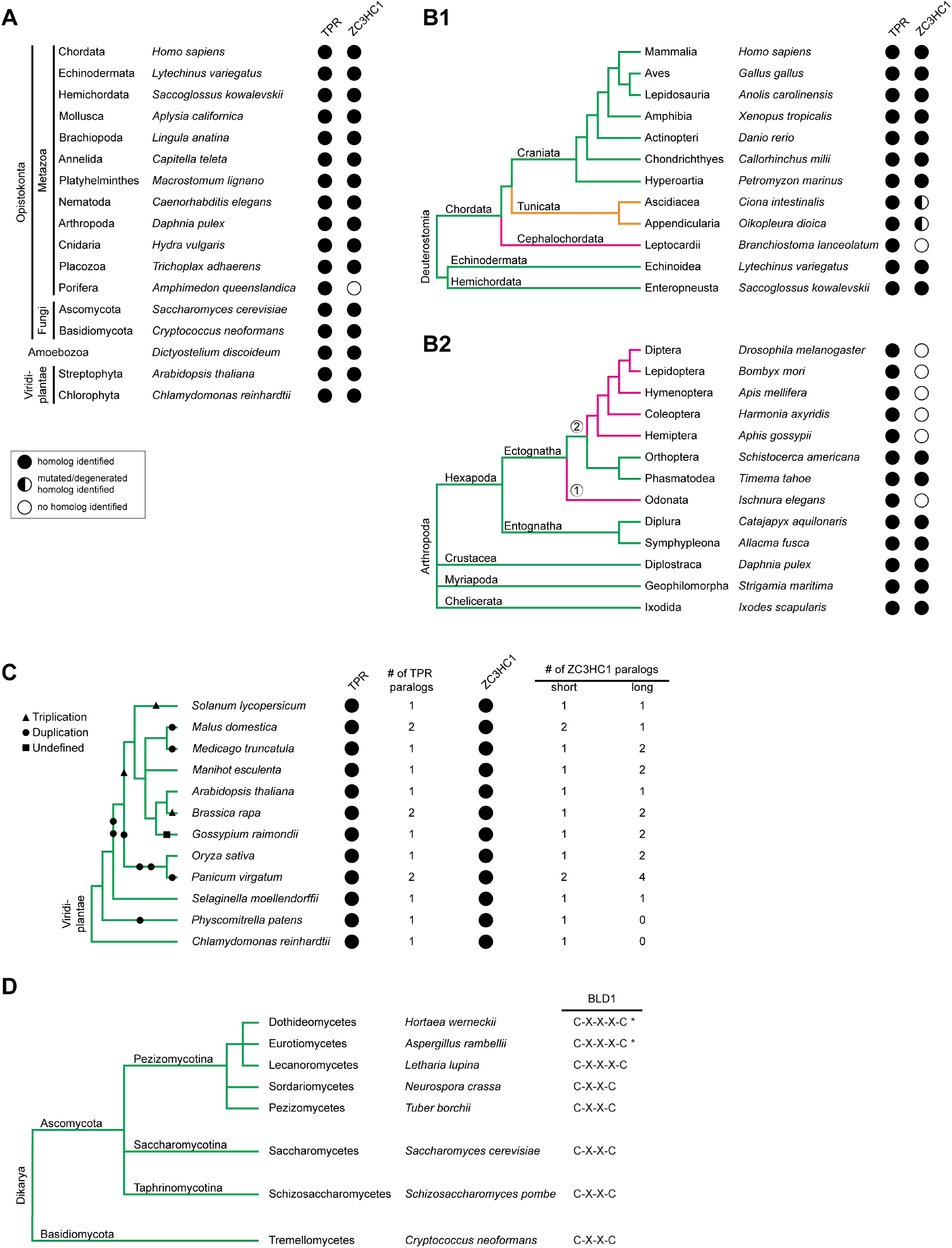
Distribution of ZC3HC1 and its homologs among eukaryotes. **(A)** Selection of phyla and divisions with representative species in which ZC3HC1 homologs were identified by searching through eukaryotic sequence databases. The presence of corresponding TPR homologs, as the potential binding partner for each species’ ZC3HC1 homolog, is depicted for comparison. The taxonomic ranks for the groups of organisms presented here mostly correspond to those used in the NCBI taxonomy database (Schoch *et al*, 2020; https://www.ncbi.nlm.nih.gov/taxonomy). Note that ZC3HC1 homologs exist in most animal phyla and the divisions of fungi, amoeba, and green plants. Among the here presented animal phyla, ZC3HC1 homologs remained undetectable merely in the sponges. **(B)** Exemplifying cladograms of the clade Deuterostomia and the phylum Arthropoda, illustrating the fate of ZC3HC1 in some subphyla. **(B1)** Cladogram of the clade Deuterostomia, listing the phyla Echinodermata, Hemichordata, and Chordata, the chordate subphyla Craniata, Tunicata, and Cephalochordata, and a selection of classes and species. Formerly presented phylogenetic trees and cladograms (e.g., Delsuc *et al*, 2006, 2018; Tassia *et al*, 2016; Zhang *et al*, 2018) guided the construction of this simplified cladogram. Note that ZC3HC1 homologs remained undetectable in the Cephalochordata. Further note that the ZC3HC1 homologs in the subphylum Tunicata harbor various mutations, differing in different classes and orders, yet all of which would prevent the human ZC3HC1 homolog from binding to the NB and TPR. **(B2)** Cladogram of the phylum Arthropoda, listing the subphyla Hexapoda, Crustacea, Myriapoda and Chelicerata, and a selection of orders and species. This cladogram was deduced from a formerly presented phylogenetic tree (Sasaki *et al*, 2013). ZC3HC1 homologs remained undetectable in the infraclass Paleoptera (①), here represented by the order Odonata, and are similarly absent in most orders of the infraclass Neoptera (②), while present in the orders Orthoptera and Phasmatodea, which in turn suggests at least two independent events that have led to the decay of the *ZC3HC1* gene, or its alteration beyond recognition, in insect evolution. **(C)** Cladogram of the kingdom Viridiplantae, with some of the orders of green plants in which genome duplication events are known to have either occurred or not, with representative species indicated. This cladogram represents an excerpt of a formerly presented phylogenetic tree (Panchy *et al*, 2016). Duplication (circles), triplication (triangles), and undefined (squares) polyploidization events are indicated. Note that several genome duplications have led to the presence of six ZC3HC1 and two TPR paralogs in the order Poales, here represented by the switchgrass *Panicum virgatum.* By contrast, in other orders that have experienced genome duplication, the second *ZC3HC1* gene appears to have been inactivated again, for example, in the spreading earth moss *Physcomitrella patens*. **(D)** Cladogram of the subkingdom Dikarya, with its two divisions Basidiomycota and Ascomycota, listing for the latter division the three subdivisions Pezizomycotina, Saccharomycotina, and Taphrinomycotina, and some of their classes, plus representative species. This cladogram represents an excerpt of a formerly presented one (Spatafora *et al*, 2017). Note that while most fungi possess a ZC3HC1 homolog with a C-X_(2)_-C tetrapeptide as part of their BLD1 zinc finger, some classes of the Pezizomycotina instead feature a C-X_(3)_-C. Note, though, that while the occurrence of the C-X_(3)_-C pentapeptide in the classes Eurotiomycetes and Dothideomycetes is far more dominant, a few of their orders nonetheless only feature sequences with a C-X_(2)_-C, with these classes here therefore additionally marked by an asterisk.

Also remarkable, in species with a non-duplicated genome, we found only one gene coding for a protein with a NuBaID signature, together with a few complementing features that we eventually defined as characterizing a prototypic ZC3HC1 homolog (see further below). Only in some groups of organisms in which one or more rounds of whole-genome duplications appear to have occurred (e.g., Sinha *et al*, 2017; Li *et al*, 2018; Qiao *et al*, 2019) could one identify two or more of such NuBaID-encoding genes per species. The latter was the case, listing only some examples, in plants (Figure 3C), in some fungi, here exemplified by *Hortaea werneckii* (see Supplemental List of Sequences for ZC3HC1 homologs) as a member of the class Dothideomycetes, and in some hexapods, like in springtails, here represented by *Allacma fusca* (Figure 3B2; Supplemental List of Sequences). In such organisms, the similarities between the respective proteins’ sequences were evident also beyond their NuBaID signatures (our unpublished data), again in line with early findings of two closely related IAPs in *Arabidopsis thaliana* (Higashi *et al*, 2005), here now referred to as ZC3HC1 homologs.

While we would currently describe a multitude of putative ZC3HC1 homologs so far detected within the eukaryotic realm by a NuBaID signature variant that reads G-[WYF]-X_(6,24)_-C-X_(2)_-C-X_(17,82)_-H-X_(3)_-C-X-[WY]-X_(48,232)_-G-[WYF]-X_(8,140)_-C-X_(2)_-C-X_(14,994)_-H-X_(3)_-C-X-[WY], with this signature here numbered (3) for simplified classification, we found the likely ZC3HC1 homologs of other species reflecting an even wider range of diversity. Among these were, for example, some fungal homologs that appeared to have come up during evolution with different spacing between the first two cysteines of the first BLD’s zinc finger signature, reading C-X_(3)_-C instead of C-X_(2)_-C, and which as such would not have been detectable with a NuBaID signature that would not consider such variance. However, after having created additional *Hs*ZC3HC1 mutants with such C-X_(3)_-C spacing, we found these well capable of binding to the NE of human cells, even in the presence of the wild-type version of ZC3HC1 (Figure 3D; Supplemental Figure S3). Furthermore, we also noted conspicuous residue diversity between alleged homologs at positions corresponding to the tryptophan residues W158 and W431 of *Hs*ZC3HC1, which are not essential for NE-association in a ZC3HC1 KO cell (see above).

Such diversity was then illustrated by another lower-stringency version of the NuBaID signature, here again simplistically numbered (4), in which we expanded the spacings also beyond those actually detected. This signature, which we used as yet another one of several versions for repeatedly data mining over time the SWISS-PROT and TrEMBL databases (Bairoch & Apweiler, 1997) with the ScanProsite tool (de Castro *et al*, 2006) read G-[WYF]-X_(5,25)_-C-X_(2,3)_-C-X_(10,100)_-H-X_(3)_-C-X-[WYFML]-X_(40,250)_-G-[WYF]-X_(5,150)_-C-X_(2)_-C-X_(10,1500)_-H-X_(3)_-C-X-[WYFRCV]. Once again, we had found some residues of this signature tolerable in the context of the human protein when we had them replace one of the human homolog’s tryptophans (Supplemental Figure S2C and our unpublished data).

However, while the more detailed description of the data mining’s findings and outcome went beyond the scope of the current study, we chose two of the proteins we had newly identified as potential ZC3HC1 homologs for already here addressing two questions. Namely, first, whether the possession of a NuBaID signature would also mark a different phyla’s protein as one that would be positioned next to its species’ NPC and interact with its TPR homolog, irrespective of how little sequence similarity such a putative ZC3HC1 homolog might share with *Hs*ZC3HC1. And second, whether residues we had found essential for allowing *Hs*ZC3HC1 to bind to *Hs*TPR might also be similarly essential for a distant, NuBaID-containing relative and its binding to a corresponding TPR homolog.

Even though some other species’ putative ZC3HC1 homologs appeared even more exotic when compared to *Hs*ZC3HC1, we chose the following two putative homologs for being more closely investigated. The one was DDB0349234 from *D. discoideum*, and the other, in particular, the non-essential protein Pml39 from *S. cerevisiae*, which had already been found to be a binding partner of the TPR homologs in yeast, namely Mlp1p and Mlp2p (Palancade *et al*, 2005). However, not sharing any immediately evident sequence similarity with any human protein, it was not surprising that *Sc*Pml39p had not been correlated with *Hs*ZC3HC1 so far. Accordingly, sequence mining for ILP/ZC3HC1 proteins had reportedly not allowed for detecting a homolog in *S. cerevisiae* either (Higashi *et al*, 2005). Likewise, with most of the different search tools we used for sequence database mining, one could neither detect *Hs*ZC3HC1 starting from *Sc*Pml39p nor vice versa (see Supplemental Information 2).

While the amoebic protein had a zf-C3HC motif assigned to it (http://ftp.ebi.ac.uk/pub/ databases/Pfam/releases/Pfam18.0/; e.g., Finn et al, 2008) when we conducted such searches, this was not so for *Sc*Pml39p, with the latter only listed as a zf-C3HC motif-possessing protein later (http://ftp.ebi.ac.uk/pub/databases/Pfam/releases/Pfam30.0/). Even so, the zf-C3HC motif has not yet (July 2022) been assigned to *Sc*Pml39 in some databases, like, for example, NCBI’s Conserved Domain Database (CDD; https://www.ncbi.nlm.nih.gov/Structure/cdd/cdd.shtml; and further below). Moreover, neither the amoebic protein nor *Sc*Pml39p had an Rsm1 motif assigned to it back then, which remains to be the case to date (http://ftp.ebi.ac.uk/pub/databases/ Pfam/releases/Pfam35.0/; https://pfam.xfam.org/family/PF08600; and further below).

By contrast, already when having used the tandem arrangement (Supplemental Figure S2D2) of the abovementioned minimalist signature (2) for database searches via ScanProsite, we had found *Sc*Pml39 to be the only protein in the budding yeast complying with this motif, and this then also held for all further derivatives of this signature, including (3) and (4) mentioned above. While the amoebic protein had not been identifiable with the tandem arrangement of signature (2), it too was then readily detectable with (3) and (4).

The sequence diversity between *Hs*ZC3HC1, *Sc*Pml39p, and DDB0349234, which we will refer to as *Dd*ZC3HC1, manifested itself in various ways, including size, pI, aa composition and a low degree of overall sequence similarity. While the human homolog is a 502 aa long protein of 55.3 kDa with a pI of 5.44, the 386 aa long *Sc*Pml39p of 39.2 kDa has a pI of 9.13. The *Dd*ZC3HC1 again (accession number ON368701), of 72.5 kDa and 635 aa long, with a pI of 8.38 (see also Supplemental Figure S4B) and dominated by clusters of asparagine residues, is a protein with an aa composition strikingly different from both *Sc*Pml39p and *Hs*ZC3HC1.

Aside from a few additional amino acids whose contributions to NB binding were not a topic of the current study, the three proteins’ only common primary sequence feature appeared to be, once again, describable by a NuBaID signature variant. In its simple version, here numbered (5), it would read G-W-X_(9,14)_-C-X_(2)_-C-X_(31,34)_-H-X_(3)_-C-X-W-X_(77,96)_-G-[WY]-X_(10,89)_-C-X_(2)_-C-X_(16,183)_-H-X_(3)_-C-X-[WY]. This signature was then, of course, again sufficient for the pattern-based specific detection of all three proteins in their respective species via ScanProsite. Especially eye-catching, the different parts of the NuBaID signature appeared separated from each other by sequence segments that strikingly differ in length between the different species’ proteins (Figure 4A and 4B; also addressed further below). Moreover, in these and numerous other species, we noted that those segments embedded between the different parts of the NuBaID signature often did not share any notable sequence similarity, sometimes even not among closely related species. The large insertions, in particular, were predicted to mainly have in common only that they are likely to be intrinsically disordered loops (see also Oates *et al*, 2013, and further below).

**Figure 4.**
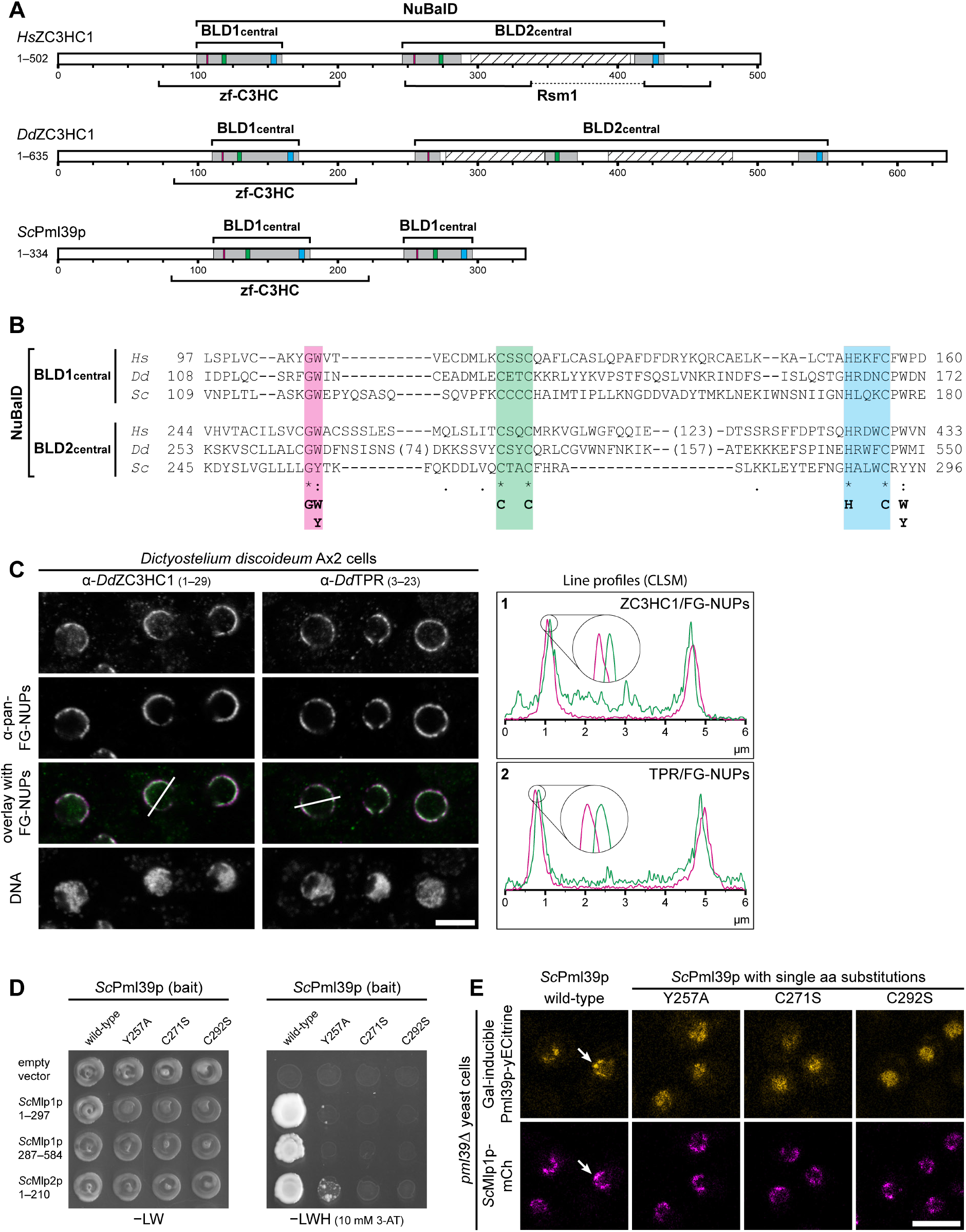
NB- and TPR/Mlp-interacting ZC3HC1 homologs with an evolutionary conserved NuBaID signature exist in *Saccharomyces cerevisiae* and *Dictyostelium discoideum*. **(A)** Schematic depiction of *D. discoideum* ZC3HC1 *and S. cerevisiae* Pml39p, next to *Hs*ZC3HC1 for comparison. The areas highlighted in the central regions of their two BLDs, together representing the bimodular NuBaID, correspond to those already described in Figures 1C and 2A1, with the boxes in magenta now depicting the position of an evolutionarily relatively conserved dipeptide that here can either read G-W or G-Y, with the latter dipeptide part of the BLD2 of Pml39p. Since the known NLS of *Hs*ZC3HC1 and the unknown, only conjecturable NLS of the two other homologs appear to be differently positioned relative to each other within regions not regarded as part of the BLDs, neither of them are depicted here. In the case of *Hs*ZC3HC1, the boundaries for the here shown minimal central regions of both BLDs are identical to those specified in Figure 2A2. The outer boundaries of the *Dictyostelium* homolog’s central BLD1 and BLD2 regions are here defined by I108 and D172 and by K253 and I550, respectively. The inner boundaries flanking the two apparent insertions within the *Dd*ZC3HC1 BLD2 are here presented as corresponding to S273 and D348 and to K371 and A529, respectively. Parts of these *Dd*ZC3HC1 insertions that the PONDR algorithm predicts to comprise mostly unstructured residues, except for V418 to N433, range from N277 to K347 and N393 to S482 (hatched boxes). For *Sc*Pml39p, the outer boundaries of its BLD1 central region are here defined by V109 and E180, and those for BLD2 by K245 and N296. Bracket-marked areas represent the regions to which the Pfam database has assigned a zf-C3HC or Rsm1 motif to date, with no Rsm1 motif though attributed to *Dd*ZC3HC1 and *Sc*Pml39 so far. **(B)** Sequence alignments of the BLDs’ central regions, according to those presented in Figure 2A2. A minimal sequence signature identical for both BLDs of all three proteins is provided in the bottom line. **(C)** Double-labeling IFM of *D. discoideum* Ax2 cells with pan-FG-NUPs antibodies and antibodies for either *Dd*ZC3HC1 or *Dd*TPR. Micrographs of the DNA-stained cells are shown as a reference. Confocal laser scanning microscopy, with the focal plane approximately at the equator of most nuclei, revealed specific labeling of the slime mold’s NE. Furthermore, sections of nuclei marked by lines 1 and 2 in the overlay micrographs were analyzed by ImageJ software, with line profiles plotted. Note that the 4× enlarged line profile sections from both sides of the corresponding nuclei reveal the offset location of *Dd*ZC3HC1 and *Dd*TPR, relative to the NPCs’ immunolabeled FG-repeat nucleoporins, towards the nuclear interior. Bar, 5 µm. **(D)** Representative Y2H data obtained with expression vectors for segments of Mlp1p and Mlp2p, in combination with vectors coding for the intact version of Pml39p or a selection of Pml39p mutants with single aa substitutions of the NuBaID signature. Y2H interactions were visualized, as described in Figure 2C, by replica-plating the expression vector-transformed cells grown on a –LW and –LWH selection medium; the latter supplemented with 10 mM 3-AT (complementing data are presented in Supplemental Figure S5B). Note that while a robust Y2H interaction occurred between the intact Pml39p and distinct parts of the Mlp proteins, no colony growth on the selective medium –LHW occurred when single aa substitution mutants of Pml39p, like Y257A, C271S, and C292S, were co-expressed with the Mlp proteins’ Pml39p-binding domains. **(E)** Live-cell imaging of *pml39*Δ yeast cells endogenously expressing mCherry-tagged Mlp1p, which had then been transformed transiently with expression vectors that allowed for the Gal-inducible ectopic expression of yECitrine-tagged versions of Pml39p. The latter comprised the intact wild-type protein and a selection of Pml39p mutants, each harboring a single aa substitution of Pml39p’s NuBaID signature. Confocal laser scanning microscopy, with the focal plane approximately at the nuclei’s equator, revealed that the newly synthesized wild-type Pml39p primarily accumulated at the NE (arrow). By contrast, the Pml39p mutants, no longer capable of binding to the NE, were found distributed throughout the nuclear interior instead. Asterisks mark a few cells in which Pml39p now and then was found not having been expressed. Bar, 5 µm.

As one of the next steps, we turned towards *Dd*ZC3HC1 to assess whether a protein possessing a prototypic NuBaID signature in a phylum beyond the opisthokonts might also occur positioned next to the NPC and its TPR homolog. Given the sequence peculiarities of *Dd*ZC3HC1, which also held for its large loop-like insertions, we considered this alleged homolog well suited for addressing whether the NuBaID signature might represent the evolutionarily conserved common hallmark of a distinct class of NB- and TPR-interacting proteins. Furthermore, *D. discoideum* had already been shown to harbor NBs (Beck *et al*, 2004), and this species thus met the requirement of holding ready, in principle, a potential binding site for *Dd*ZC3HC1 at the NE. In addition, we had already noted that *D. discoideum* possesses a TPR homolog, namely DDB0308586, with features also characteristic for, e.g., metazoan TPR homologs (Supplemental Figure S4A1 and S4A2; Kuznetsov *et al*, 2002), and a sequence-deduced Mr of 235 kDa (accession number ON368702; Supplemental Figure S4A3). Furthermore, antibodies that we had raised against *Dd*TPR had already allowed us to locate the protein at the *Dictyostelium* NPC’s nuclear side (see further below), with this finding now also in line with recent reporting of *Dd*TPR, when tagged with mNeonGreen, to be located at the NE (Mitic *et al*, 2022). Moreover, initial Y2H interaction studies had revealed that a *Dd*ZC3HC1 deletion mutant, from which we had removed the two largest poly-asparagine-dominated loops (Supplemental Figure S4B), was well capable of robustly binding to parts of *Dd*TPR’s N-terminal domain (Supplemental Figure S4C).

Then, to address where *Dd*ZC3HC1 locates in *Dictyostelium*, we investigated this by immunofluorescence microscopy (IFM). The antibodies raised against *Dd*ZC3HC1 specifically labeled a single protein with an electrophoretic mobility corresponding to about 80 kDa in total *D. discoideum* cell extracts, in line with the *Dd*ZC3HC1 protein’s sequence-deduced molecular mass. In addition, some of the immunoblotting-(IB) and IFM-compatible antibodies that we had raised against *Dd*TPR allowed for similarly specific labeling of a protein of about 250 kDa (Supplemental Figure S4D). On the other hand, several *D. discoideum* proteins with molecular masses exceeding 100 kDa could be labeled with “pan-FG-NUPs” antibodies (Supplemental Figure S4D) that we had generated for being reactive with various species’ NPC proteins (nucleoporins) via the binding to their common FG-repeat domains.

In IFM then, the *Dd*ZC3HC1 and *Dd*TPR antibodies specifically labeled the slime mould’s NE, revealing a clear association with the NPC (Figure 4C). Since we had not managed to generate pairs of antibodies from different species that would have allowed for conventional double-labeling of *Dd*TPR and *Dd*ZC3HC1 directly within the same specimen, we performed the alternatively possible double-labeling with the pan-FG-NUPs antibodies. Applying the same IFM protocol and using the same microscope settings for inspecting all specimens side by side, the positions of *Dd*ZC3HC1 and *Dd*TPR could thus be visualized relative to those of the FG-repeat nucleoporins, which revealed an offset location of *Dd*ZC3HC1 and *Dd*TPR towards the nuclear interior (Figure 4C).

In parallel to having shown that *Dd*ZC3HC1, as the only NuBaID-signature-possessing protein in this species of the phylum Amoebozoa, occurs positioned just like its binding partner *Dd*TPR adjacent to the slime mould’s NPCs, we concomitantly had been focusing on the budding yeast protein Pml39 and its relationship with the Mlp proteins. In this context, we had been particularly eager to determine whether the integrity of the NuBaID signature of Pml39p too might be required for its binding to the Mlp proteins, as the yeast’s TPR homologs.

To this end, we had first performed Y2H experiments, having created for such purpose bait and prey constructs that were coding for (i) Mlp protein segments, among which were ones also similar to those formerly found interacting with *Sc*Pml39p (Palancade *et al*, 2005), for (ii) wild-type Pml39p, and for (iii) a collection of Pml39p mutants with single aa substitutions. Apart from confirming that Pml39p could interact with distinct parts of the Mlp proteins (Supplemental Figure S5A), these Y2H experiments then additionally revealed that Pml39p mutations, at aa positions equivalent to those of *Hs*ZC3HC1 essential for TPR binding, abolished or markedly impaired also Pml39’s ability to bind to Mlp1p and Mlp2p (Figure 4D; Supplemental Figure S5B). By contrast, other single aa substitutions of the yeast’s minimal NuBaID signature again, like Y257W, which transformed the second BLD’s dipeptide G-Y to G-W, thus identical then to the corresponding G-W of *Hs*ZC3HC1, did not notably impair binding to any of the Mlp1p and Mlp2p segments capable of Y2H-interacting with wild-type Pml39p (our unpublished data).

Next, to investigate whether, in particular, those Pml39p mutations that had prevented Y2H interactions with the Mlp proteins would also impair binding to the NB *in vivo*, we generated yeast strains stably expressing all Mlp1p as mCherry-tagged polypeptides in a *pml39*Δ strain. We then used the resulting *PML39* deletion strain for the galactose-inducible expression of yECitrine-tagged intact and mutant versions of Pml39p in the absence of any potentially competing endogenous Pml39p. With this experimental set-up later equivalent to the one in which we had transfected human ZC3HC1 KO cells with expression vectors coding for different versions of ZC3HC1 (Supplemental Figure S2B2), we inspected the transformed yeast cells by live-cell imaging. First, and in line with former findings (Palancade *et al*, 2005), we saw the yECitrine-tagged intact version of Pml39p and the mCherry-tagged Mlp1 polypeptides co-localizing at the NE. By contrast, we found all of the Pml39p mutants with those single aa substitutions that had attenuated or abolished Y2H-binding to the Mlp proteins (Supplemental Figure S5B) also notably impaired in or not capable of NE-binding at all and distributed throughout the nuclear interior instead (Figure 4E, Supplemental Figure S5C). This outcome was thus the same as for the corresponding aa substitutions introduced into *Hs*ZC3HC1, which had abolished interaction with *Hs*TPR and the NB in human cells.

All in all, the findings made during our excursion into the phyla Ascomycota and Amoebozoa prompted us to conclude that a protein’s possession of an intact, prototypic NuBaID signature stands for a high probability that such a protein is a genuine ZC3HC1 homolog capable of binding its species’ TPR/Mlp1 polypeptides. Even though our results did not allow us to conclude that every protein with such a signature will eventually turn out to be an NB-located TPR binding partner, as discussed further below, they already indicated, as attested in three widely different species, that a ZC3HC1:TPR ensemble at the NB is in use widely across the eukaryotic realm.

However, at this point, one question still demanded an answer: whether some ZC3HC1 homologs, apart from sharing a NuBaID signature and the ability to bind to their respective TPR homologs, might also share with *Hs*ZC3HC1 a functional property, like its capability of interconnecting pools of TPR polypeptides.

### *Sc*Pml39p, as the budding yeast homolog of *Hs*ZC3HC1, is required for NE-positioning of subpopulations of *Sc*Mlp1p and enables their interlinkage in nuclear foci

With (i) Pml39p known to be a protein only located at the NE because of its binding to the Mlp proteins, in particular to Mlp1p (Palancade *et al*, 2005; Supplemental Figure S6A1), with (ii) ZC3HC1 being a protein whose NB-association depends on its binding to TPR (Gunkel *et al*, 2021), and with (iii) both ZC3HC1 and Pml39p sharing a NuBaID sequence signature whose integrity was essential for binding to each species’ NBs and TPR homologs, the next inevitable question was apparent. On the one hand, the deletion of *PML39* had been reported as not affecting the NE localization of Mlp1p or Mlp2p (Palancade *et al*, 2005), and on the other, we knew that ZC3HC1 was required for the positioning of subpopulations of TPR polypeptides at the NB (Gunkel *et al*, 2021). Even though it was not unimaginable that the ZC3HC1 homologs in humans and yeasts might differ in terms of some of their functional properties, given their poor overall degree of sequence similarity, we nonetheless examined to what extent Pml39p might differ from ZC3HC1’s role in interconnecting TPR subpopulations.

First, we investigated the NE positioning of Mlp1p and Mlp2p in the presence and absence of Pml39p. We thereby hardly noticed any unambiguous effect regarding the Mlp2 polypeptides’ NE association when comparing a *PML39* wild-type (*wt*) and a *pml39*Δ version of a strain in which we had tagged the *MLP2* gene with yEGFP (Supplemental Figure S6A2), with this observation in line with the formerly reported findings. Even after repeatedly inspecting populations of Mlp2p-yEGFP-expressing *PML39wt* and *pml39*Δ cells in parallel, having also tested different growth conditions in this context, we only sporadically noted at most some minor diminishment of the *pml39*Δ cells’ NE-associated overall signal intensities for Mlp2p-yEGFP. More commonly, when we had randomly acquired live-cell images with the same microscope settings from populations of *PML39wt* and *pml39*Δ cells, each loaded onto the imaging slides in duplicate, and had then coded these images and compiled mixed collections of them, we arrived at the following result: When we attempted to unambiguously sort these coded images of Mlp2p-yEGFP-expressing cells back into groups that, after decoding, would have represented only either *PML39wt* or *pml39*Δ cells, this turned out not to be possible. Similarly, having yEGFP-tagged other components of the yeast’s NPC and NB, we also found that the NE-positioning of some of the other NB-associated proteins, like Mad1p, Sac3p, and Ulp1p, was not notably or only very moderately affected in the *pml39*Δ cells as compared to the corresponding wild-type (Supplemental Figure S6B).

By contrast, a strikingly different situation manifested itself when comparing corresponding yeast strains in which we had tagged the *MLP1* gene with yEGFP. In the *pml39*Δ cell populations, compared to those of the *PML39wt* cells grown in parallel, the NE-attached overall amounts of Mlp1p appeared notably diminished (Figure 5A, Supplemental Figure S6A3). Such differences were, in fact, evident enough to recurrently allow for unambiguously distinguishing *pml39*Δ from WT cells, and we could sort mixed compilations of coded live-cell micrographs of such Mlp1p-yEGFP-expressing *PML39wt* and *pml39*Δ cells faultlessly back into groups representing only the one and other strain. Furthermore noteworthy, in many of the cells within a *pml39*Δ population, a certain amount of yEGFP-tagged Mlp1p appeared diffusely distributed throughout the nuclear interior, a feature generally not noted within the *PML39wt* cells (Figure 5A). Moreover, we made essentially the same observations when comparing *PML39wt* and *pml39*Δ strains in which we had tagged the *MLP1* gene with mCherry (our unpublished data; but see also Supplemental Figure S6B).

**Figure 5.**
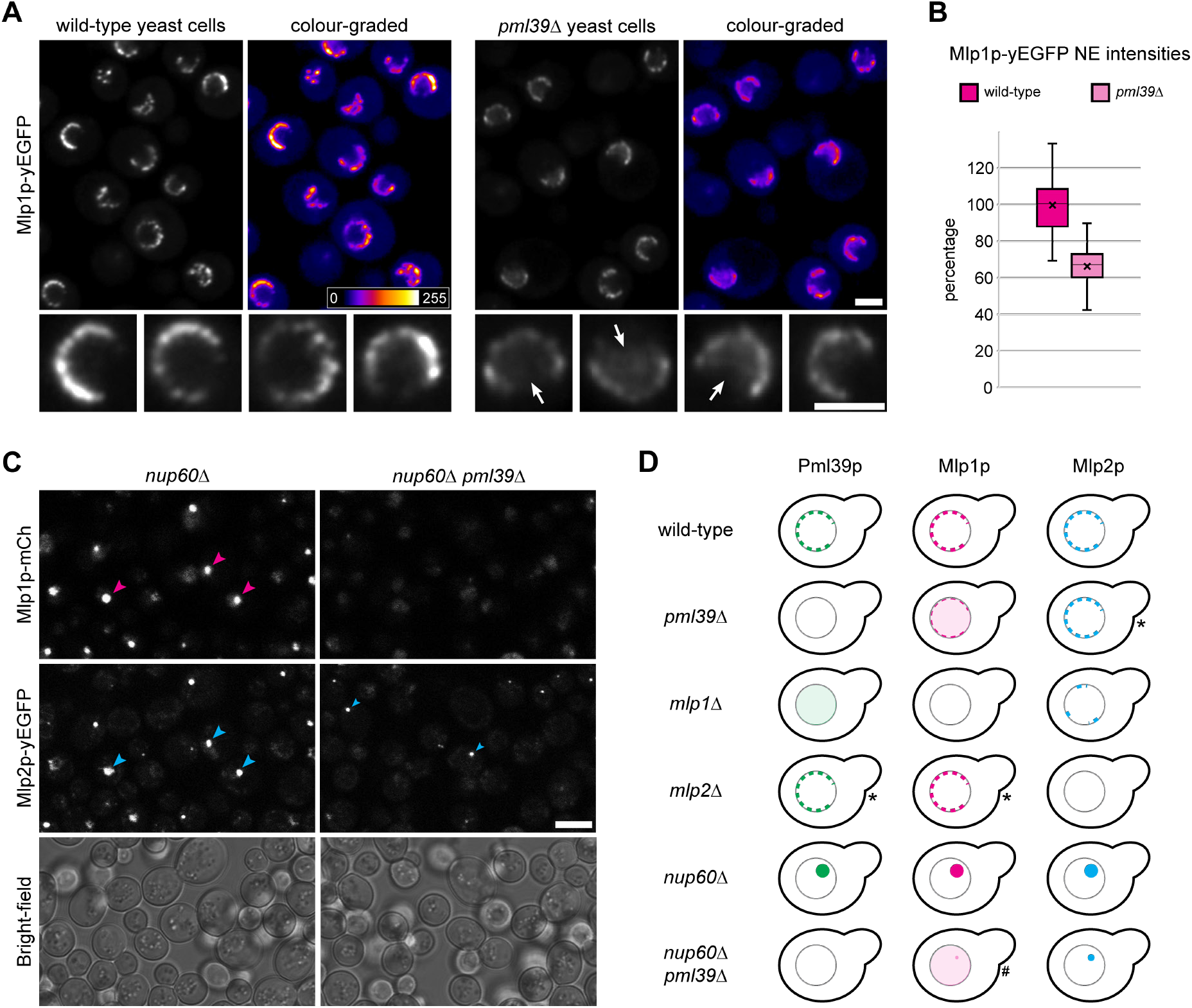
Absence of Pml39p results in reduced amounts of NE-associated Mlp1p in *pml39*Δ cells and essentially prevents in *nup60*Δ *pml39*Δ cells the accumulation of Mlp1p in nuclear clusters. **(A)** Live-cell fluorescence microscopy of *PML39wt* and *pml39Δ* yeast cells endogenously expressing all Mlp1p as yEGFP-tagged polypeptides. The two strains had been grown as asynchronous populations, loaded into separate wells next to each other, with these specimens then analyzed in parallel and images acquired with identical microscope settings. Next to a color look-up table, representative overview images are also presented as color-graded. Some of the nuclei in the overview images are also shown at higher magnification. Note that the yEGFP signal intensities at the NEs of the *pml39Δ* cells appeared overall notably reduced compared to the *PML39wt* cells. Further note that a certain amount of Mlp1p-yEGFP was often found distributed throughout the *pml39Δ* cells’ nuclear interior instead, with arrows pointing at examples of such a nucleoplasmic pool of Mlp1p. Bars, 2 µm. **(B)** Quantification of signal yields of yEGFP tagged to Mlp1p at the NEs of *PML39wt* and *pml39Δ* cells. Populations of *PML39wt* and *pml39Δ* cells had been grown under identical conditions in parallel to each other, harvested in the exponential growth phase, and then imaged as live cells using the same microscope settings. ImageJ software was used for determining the signal intensities of NE segments from essentially all NEs that could be seen in an equatorial focus plane in several randomly chosen overview micrographs for each of the two strains’ populations. In order to compare only truly NE-associated fluorescence levels, GFP signals from cytoplasmic and nucleoplasmic areas were subtracted. The data presented here represent the mean results of two separate experiments (n = 50 nuclei per dataset) in which populations of *PML39wt* and *pml39Δ* cells had been grown next to each other, with the two pairs of *PML39wt* and *pml39*Δ cell populations harvested on different days and then evaluated independently from the other. The two experiments’ individual results are presented in Supplemental Figure S6C1. Box plots display the relative signal intensity values, with the arithmetic means marked by x, with the ones for the *PML39wt* cells set to 100%, and the standard deviations provided (for an alternative presentation, in which the *pml39Δ* cells’ mean value is set to 100%, see Supplemental Figure S6C2). While the signal intensities between individual nuclei differed notably, with such variation somewhat more pronounced in the *PML39wt* than in the *pml39*Δ cells, note that the mean Mlp1p-yEGFP signal yields for the *pml39Δ* cells’ NEs were in general found to be reduced notably. **(C)** Live-cell imaging of *nup60*Δ cells and *nup60*Δ *pml39*Δ cells endogenously expressing mCherry-tagged Mlp1 and yEGFP-tagged Mlp2 polypeptides. Bright-field micrographs are shown as a reference. Note that in the Nup60p-deficient cells, both Mlp1p and Mlp2p were no longer attached to the NPCs. Instead, they often occurred focally accumulated, together with Pml39p (Supplemental Figure S6E), within prominent nuclear clusters (some marked by magenta-and blue-colored arrowheads). By contrast, in *nup60*Δ *pml39*Δ cells, Mlp1p was hardly or no longer detectable as part of these nuclear clusters, whilst Mlp2p could still be present in such foci (two marked by a small blue arrowhead), which then, though, typically were of smaller size. Bar, 5 µm. **(D)** Schematic depiction of the subcellular distribution of FP-tagged Mlp and Pml39 polypeptides in some of the yeast KO strains investigated in the current study. In contrast to the majority of strains, for which the observations were rated as being unequivocal, the outcome for a few strains, here marked by asterisks, appeared to vary moderately between replicated rounds of inspection. Such ambiguity held for some minor reduction in the amounts of (i) NE-associated Mlp2p in the *pml39*Δ cells, (ii) NE-associated Pml39p in the *mlp2*Δ cells, and (iii) NE-associated Mlp1p in the *mlp2*Δ cells, with such reductions noted only in some replicates but not others, and with such variations appearing to correlate to some extent with cell culture growth phases. By contrast, the *nup60*Δ *pml39Δ* strain is marked by a hash because we technically could not unambiguously exclude the possibility that some residual minor amounts of Mlp1p, apart from those Mlp1 polypeptides distributed throughout the nucleoplasm, might still be associated with this strain’s small-sized Mlp2p foci. Finally, note that additional data are summarized in Supplemental Figure S6F).

Altogether, such a reduction in the NE-associated amounts of Mlp1p, accompanied by a nucleoplasmic pool of Mlp1p due to Pml39p deficiency, resembled the effects we had by then noted in human cells, in which knockdown of ZC3HC1 by RNAi had caused a reduction in the NE-associated amounts of TPR, accompanied by a nucleoplasmic pool of TPR. Later, such findings also turned out in line with very similar results in ZC3HC1 KO cells (Gunkel *et al*, 2021) and eventually upon degron-mediated ZC3HC1 degradation too (Gunkel & Cordes, 2022), with the latter findings now also resembling the outcome of a study in which degron-mediated degradation of Pml39p is shown affecting the NE-association of Mlp1p (Bensidoun et al, 2021b).

To approximate the extent to which the *pml39*Δ cells’ NE-associated amounts of Mlp1p were diminished, we determined the signal yields for Mlp1p-yEGFP at the WT and *pml39*Δ cells’ NEs. In doing so, we found the NE-associated signal intensities between individual *PML39wt* cells varying notably. Such cell-to-cell differences were more pronounced than the lower degree of variation between TPR’s NE-associated relative amounts in human cells, both in synchronized and asynchronous cell populations, with and without ZC3HC1 (Gunkel *et al*, 2021), when having conducted such quantifications in the same manner as for the yeast cells. Nonetheless, when we determined the mean Mlp1p-yEGFP signal intensity of all NEs seen in equatorial focus in randomly acquired live-cell images, we found up to about one-third of the cells’ ordinarily NE-associated total amounts of Mlp1p no longer located there in the absence of Pml39p (Figure 5B; Supplemental Figure S6C).

Then, to gain more insight into how Pml39p might allow for the interconnection of Mlp1 polypeptides, we conducted additional experiments, once again similar to those we had carried out or were still performing at that time in human cells. In the latter, knockdown of NPC protein NUP153 by RNAi had long been known to cause TPR to accumulate in cytoplasmic and nuclear foci (e.g., Hase & Cordes, 2003). Later though, we had noted that such TPR-containing foci were mostly no longer detectable when both NUP153 and ZC3HC1 together had been knocked down by RNAi, suggesting early on that ZC3HC1 might be required for such foci’s formation (our unpublished data). Eventually, we found that the TPR foci also did not arise when knocking down NUP153 in ZC3HC1 KO cells once the latter had become available (Gunkel *et al*, 2021; our unpublished data).

Having wanted to know whether an absence of *Sc*Pml39p would have a similar impact on Mlp foci, we performed experiments in the budding yeast that corresponded to those we conducted in human cells. With protein *Sc*Nup60 considered homologous to NUP153, concerning both proteins’ role in recruiting TPR/Mlp1 polypeptides (e.g., Hase & Cordes, 2003), we asked how the prominent nuclear clusters of then NPC-detached Mlp1p and Mlp2p, which arise in the yeast cell’s nucleus in the absence of Nup60p (e.g., Feuerbach *et al*, 2002), might be affected when Pml39p was additionally absent. To address this question, we generated and studied further yeast strains.

In *nup60*Δ cells, we found conspicuous nuclear clusters of no longer NPC-attached Mlp1p and Mlp2p (Figure 5C, Supplemental Figure S6D and S6E1), confirming former findings (e.g., Feuerbach *et al*, 2002). Furthermore, we found Pml39p co-localizing with the Mlp polypeptides in such foci (Supplemental Figure S6E1), which was similarly in line with what had been observed earlier (Palancade *et al*, 2005). The novel result then, which we regarded as of particular interest, was the seeming absence of Mlp1p foci in the absence of Pml39p, with such foci hardly or no longer detectable within the *nup60*Δ *pml39*Δ cells (Figure 5C). Instead, we found Mlp1p then often appearing distributed diffusely throughout these cells’ nuclei instead, which thus argued for Pml39p being required for keeping the Mlp1 polypeptides place-bound even at sites other than the NPC (Figure 5C and 5D); with this result reciprocally confirmed by also inspecting separately generated *nup60*Δ *pml39*Δ strains (our unpublished data). In addition, we also considered it of interest that the focal accumulation of Mlp2p appeared less affected by Pml39p deficiency (Figure 5C and 5D), indicating that interactions between Pml39p and Mlp2p differ from the likely more complex arrangements between Pml39p and Mlp1p. Along this line, we regarded it as of further note that the absence of Mlp2p did not prevent the formation of conspicuous foci in *nup60*Δ *mlp2*Δ cells, with Mlp1p and Pml39p still co-localizing in such foci, while the notably smaller-sized and less often observed Mlp2p foci in *nup60*Δ *mlp1*Δ cells only appeared to attract minor amounts of Pml39p (Supplemental Figure S6E and S6F).

However, at least concerning the Mlp1p foci, present in *nup60Δ* cells while absent in *nup60Δ pml39Δ* cells, we rated these foci as equivalent to the TPR foci in the NUP153-deficient but ZC3HC1-positive human cells while absent in the NUP153- and ZC3HC1-deficient cells. Moreover, since we later could prove via degron-mediated ZC3HC1 degradation that the TPR foci are indeed held together by ZC3HC1 (Gunkel & Cordes, 2022), we eventually also considered it legitimate to interpret our yeast data accordingly, namely by also regarding Pml39p as a protein with a direct role as a linker element between Mlp1 polypeptides even at sites remote from the NB.

Apart from having unveiled many features common to both ZC3HC1 and Pml39p, we need to mention, though, that we had also tested whether Pml39p and ZC3HC1 were exchangeable with regard to an interaction with the respective other species’ TPR homologs. However, we did not find Pml39p capable of binding to the NB and *Hs*TPR in human cells, neither in the presence of ZC3HC1 nor later in ZC3HC1 KO cells. Furthermore, we also did not find Pml39p capable of a Y2H interaction with the ZC3HC1-binding regions of *Hs*TPR, just as no Y2H interactions were detectable between *Hs*ZC3HC1 and the Pml39-binding parts of Mlp1p and Mlp2p (our unpublished data). However, we regarded these negative findings as likely explainable by Mlp- and TPR-binding interfaces that probably share only a limited degree of sequence similarity. This explanation appeared even more plausible once it became evident that the proteins’ common residues of the minimal NuBaID signature are not likely to engage in intermolecular interactions and that they are thus unlikely residues of the Mlp- and TPR-binding interfaces (see further below).

Therefore, even without the ultimate proof that exchangeability of homologs would have provided, we felt confident having proven kinship between *Hs*ZC3HC1 and *Sc*Pml39p. While our findings did not allow us to tell whether all species’ ZC3HC1 homologs are capable of interconnecting pools of TPR/Mlp1 polypeptides, as discussed further below, we regarded it as justified to conclude that at least *Hs*ZC3HC1 and *Sc*Pml39p are genuine homologs that share at least one similar task within their respective cells. This conclusion, though, raised the next question. Since these proteins share properties like NB binding and interacting with their TPR homologs yet appear so strikingly different at the primary sequence level, apart from their NuBaID signature, which structural features do they have in common then?

### ZC3HC1 structure predictions illustrate the NuBaIDs’ evolutionarily conserved construction and allow for newly defining each BLD’s boundaries

At this point of our research, we regarded X-ray crystallographic analyses of the ZC3HC1 structures as the next natural step, which for the human homolog, though, turned out more challenging than anticipated. Therefore, until crystallographic data becomes available, we turned towards inspecting the ZC3HC1 structures that some of the recent neural network-based deep-learning programs allowed predicting computationally, with us eventually using DeepMind’s deep-learning program AlphaFold2 (Jumper *et al*, 2021; Tunyasuvunakool *et al*, 2021) and the ColabFold platform (Mirdita *et al*, 2021). Beforehand, though, we had scrutinized how the composition and variation of the information packages that represent the input materials used for such predictions would affect any subsequently predicted BLD structures (Supplemental Information 3, Supplemental Figures S7 and S10A). The outcome of these trials dissipated some initial concerns regarding certain aspects of a BLD structure prediction and the comparability of the predicted ZC3HC1 structures. From then on, we considered the predictions for the BLDs’ structured parts, as presented in the following, conclusive within the range of accuracy and precision that we regarded as sufficient in the context of the current study.

The predictions provided by such AI contradicted some other models of what the zinc ion-coordination topology of ZC3HC1 might look like (Supplemental Figure S8), thereby supporting a model already formerly proposed (Higashi *et al*, 2005). In addition, the current predictions now illustrated the striking degree of structural similarity between the different homologs’ BLD1 domains, and they similarly uncovered the close resemblance between the different species’ BLD2 domains, as will be outlined further below (Figure 6A–C). In addition, they revealed some structural similarities, which sequence alignments had not indicated, also beyond those regions initially defined as the BLDs, as will be specified in the following too (Figure 6C and 6D). On the other hand, these predictions also visualized some striking differences between the three homologs, with the loop-like insertions within the human and amoebic ZC3HC1 and their absence from *Sc*Pml39p being especially conspicuous ones (Supplemental Figure S9A), among others. However, when blanking out such loop-like insertions, a compact arrangement of the two BLDs as two adjoining modules became apparent for all three species (Supplemental Figure S9B1). Closer inspection revealed an evolutionary conserved BLD1:BLD2 binding interface, underscoring that the two BLDs are not only a functional ensemble but also a compact structural entity (Supplemental Figure S9B2 and S9B3). With the BLDs being the current study’s main topic concerning ZC3HC1, we addressed their predicted structures in more detail, focusing on the BLD’s central parts first (Figure 6A and 6B). Guided by the initial delineation of these parts that had stemmed from primary sequence alignments and had been the basis for Figure 4A, we introduced only some minor adjustments to these boundaries based on an initial superimposition and alignment of the three homologs’ AlphaFold2-predicted structures. Next, to only expose the structural elements of the BLDs’ central parts, we blanked out the major loop-like insertions within the BLD2 of the human and amoebic homolog and then superimposed the different homologs’ BLD modules onto each other once again. Altogether, this approach allowed for illustrating the structural similarity between the central parts regarded as contributing to the zinc ion coordination spheres. Such similarities were evident when comparing the three homologs’ corresponding BLDs (Figure 6A) as well as each homolog’s two BLDs among themselves (Figure 6B).

**Figure 6.**
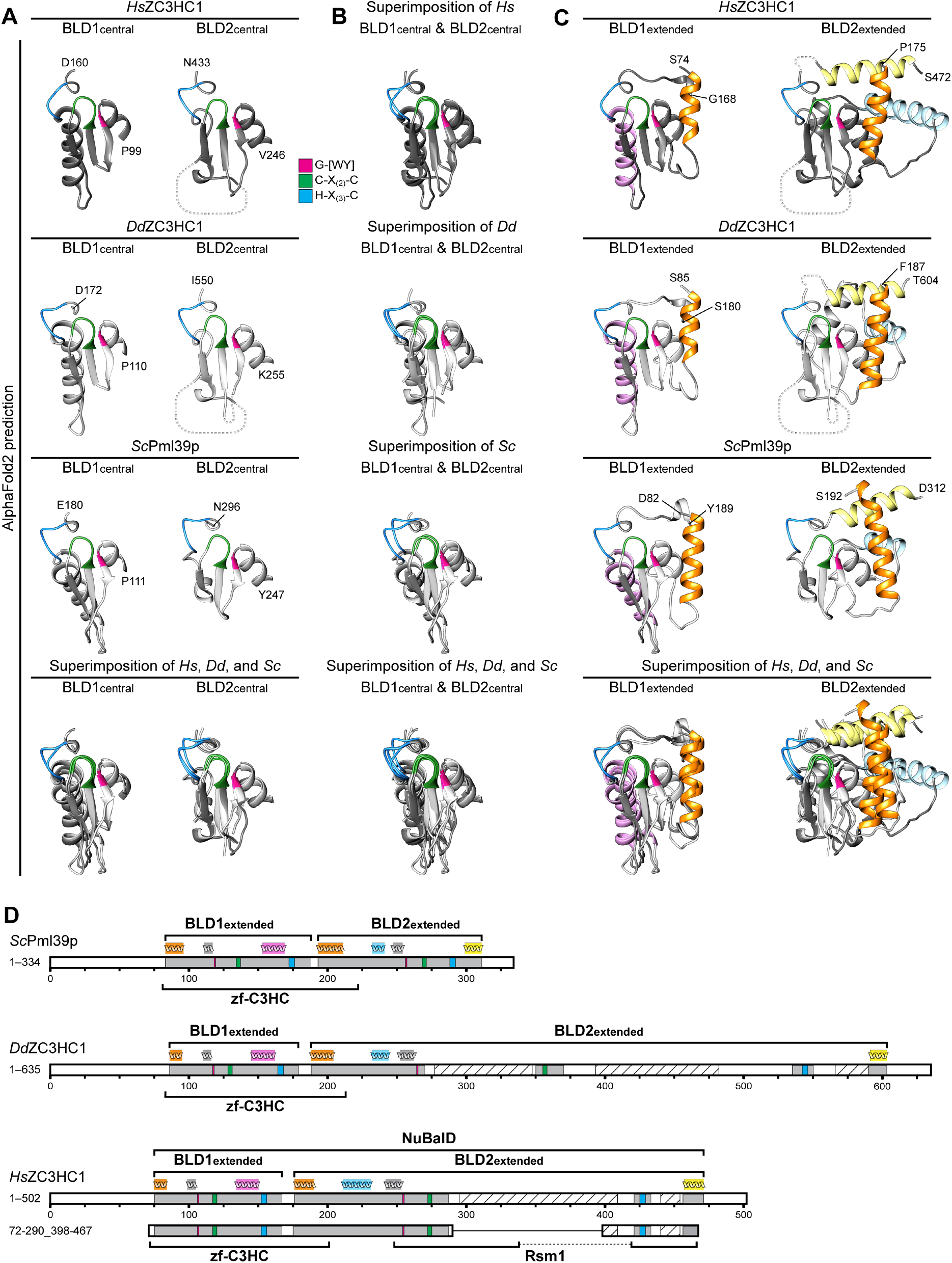
Tertiary structure predictions by AlphaFold2 underscore the striking similarities between the BLD modules of *Hs*ZC3HC1, *Dd*ZC3HC1, and *Sc*Pml39p and allow for a redefinition of their boundaries. **(A)** Structures predicted for the central regions of the BLD1 and BLD2 modules of *Hs*ZC3HC1, *Dd*ZC3HC1 and *Sc*Pml39p. The outer boundaries of the here shown central part of BLD1 are P99 and D160 for *Hs*ZC3HC1, P110 and D172 for *Dd*ZC3HC1, and P111 and E180 for *Sc*Pml39p. The outer boundaries of the central part of BLD2 here correspond to V246 and N433 of *Hs*ZC3HC1, K255 and I550 of *Dd*ZC3HC1, and Y247 and N296 of *Sc*Pml39p. Having blanked out from the human homolog’s BLD2 the major loop-like insertion, and similarly the two major insertions from the amoebic BLD2, the inner BLD2 boundaries of *HsZ*C3HC1 shown here correspond to I287 and F418, and those of *Dd*ZC3HC1 to I270 and K350, and I370 and E535, respectively. The loop-like insertions are schematically depicted as dashed lines not to scale. The source for all *Hs*ZC3HC1 and *Sc*Pml39p structures presented here and in 6B and 6C was the AlphaFold database. The structure for the sequence-corrected version of *Dd*ZC3HC1 (accession number ON368701) was determined with the source code of AlphaFold2. Superimposition of all BLD1 structures onto each other, and similarly for all BLD2 structures, was achieved with Chimera’s MatchMaker tool. The NuBaID signature’s sequence elements C-X_(2)_-C and H-X_(3)_-C, assumed to be involved in zinc ion coordination, and the G-[WY] dipeptide are colored like in Figure 4A. Note the similarities between the different homologs’ central BLD1 structures and those between their BLD2 structures. **(B)** Superimposition of the central parts of BLD1 and BLD2 onto each other. Aside from an evolutionarily conserved α-helix seemingly specific for BLD1, the structural similarity between the other central parts of both BLDs, regarded as contributing to zinc ion coordination, appears evident. **(C)** Structures predicted for those segments of *Hs*ZC3HC1, *Dd*ZC3HC1 and *Sc*Pml39p that we now regard as their essentially entire BLD1 and BLD2 modules, according to their newly defined extent. With one additional residue shown appended to each BLD’s presented boundaries, for facilitating their recognition, the BLD1 of *Hs*ZC3HC1, comprising K75–F167, is presented as aa S74–G168. BLD2, comprising A176–S471, is here shown as aa P175–S472, yet with the BLD2’s major loop-like insertion, and now also a smaller second one between I434 and E455, edited out like in 6A. Accordingly, the BLD1 of *Dd*ZC3HC1 is presented as aa S85– S180, instead of N86–F179, and its BLD2 as aa F187–T604, instead of Q188–S603. Again, the two major loops within the amoebic BLD2 were blanked out, and additionally a smaller one between aa V551 and I589. The BLD1 of *Sc*Pml39p is presented correspondingly as aa D82– Y189, instead of L83–E188, and its BLD2 as aa S192–D312, instead of S193–E311. Note that the BLD1-specific α-helix is colored in light pink, whereas α-helices seemingly specific for the BLD2 module of all three homologs are colored in light yellow and light blue. By contrast, the α-helix common to the N-terminal boundary of both BLD1 and BLD2 is highlighted in bright orange. As an aside, note that all segments shown here primarily comprise residues for which AlphaFold2 assigned, with only a few exceptions, a high per-residue confidence score of at least 70, yet mostly exceeding a score of 90. **(D)** Schematic depiction of *Hs*ZC3HC1, *Dd*ZC3HC1 and *Sc*Pml39p like in Figure 4A, but with the BLD modules newly defined. Unlike in the preceding Figures, the current schemes depict an additional minor insertion within the BLD2 of *Hs*ZC3HC1 (aa 434–455) and *Dd*ZC3HC1 (aa 551–589), close to each BLD2’s C-terminal boundary, with those parts exceeding PONDR score values of 0.5 (*Hs*ZC3HC1 aa 440–454, *Dd*ZC3HC1 aa 566–590) again shown hatched. In contrast, other regions beyond the BLDs’ outer boundaries, existing in all three homologs and predicted unstructured too by PONDR and AlphaFold2, are not highlighted for reasons of simplification. Schematic indications of α-helices above each homolog’s scheme and colored in orange, pink, blue and yellow represent the relative positions of those α-helices correspondingly colored in 6C. Note that for reasons outlined in Figure 4A, the relative position of each protein’s NLS is not depicted. Further note that according to the BLDs’ novel delineation, the Pfam database’s zf-C3HC motif would now comprise sequences not only encompassing the entire BLD1 but also part of the BLD2, while the Rsm1 motif, so far assigned only to *Hs*ZC3HC1 but not *Dd*ZC3HC1 and *Sc*Pml39p, would now only correspond to parts of the BLD2 and its loop-like insertions. Finally, note that the minimal NE binding-competent *Hs*ZC3HC1 mutant 72–290_398–467, here schematically depicted for comparison too, comprises, except for the four residues 468–471, the newly defined BLD regions in their entirety.

The similarity in the central regions of their BLDs manifested itself by the same anti-parallel arrangement of several β-sheets and an identical tetrahedral arrangement of each BLD’s three cysteines and one histidine, forming the likely zinc ion-coordination sphere. Moreover, each BLD featured the same positioning, relative to its CCHC arrangement, of the G-W/G-Y dipeptide as part of the G-[WY]-X_(9,89)_-C-X_(2)_-C peptide and of the aromatic residue at its H-X_(3)_-C-X-[WY] peptide’s C-terminal side (Figure 6A and 6B). In particular, the tryptophans’ indole and the tyrosines’ phenol rings were predicted to be similarly positioned relative to the histidines’ imidazole ring and the center of the zinc ion coordination spheres (Supplemental Figure S9C).

These structure predictions now also attested that those residues we had defined as the NuBaID’s minimal sequence signature are not likely those that will engage in direct intermolecular interactions with TPR as part of a yet-to-be-defined TPR binding interface. Instead, the NuBaID signature residues, and several of the few other ones evolutionarily conserved across different phyla, contribute to establishing or maintaining the BLDs’ central structures (see also further below). These residues apparently represent not only those directly involved in zinc ion coordination but also such that engage in other intra-BLD interactions required for establishing the BLD’s core structure and a stable zinc ion coordination sphere.

Furthermore, the structural features of the intact BLDs’ central parts were predicted to be similar to those characterizing the BIR domains of the IAPs (Supplemental Figure S10A–C), including, for example, their anti-parallel arrangement of several β-sheets flanked by an α-helix (Supplemental Figure S10C1), with this latter α-helix here corresponding to those shown in grey among the other colored a-helices of the BLDs shown further below (Figure 6C and 6D). Of further note, such an arrangement of the BLDs’ central parts was similar to the formerly proposed BLD1 structure of *Hs*ILP1/ZC3HC1 and its structural similarity with the BIR domain of human survivin/BIRC5 (Higashi *et al*, 2005).

Moreover, beyond the similarities of their central, zinc ion coordination spheres, we further noted that the BIR domains and the BLD1 of the vertebrate ZC3HC1 homologs also have yet other features in common, like, e.g., a conspicuous groove exposed on the domains’ surface. In the case of *Hs*ZC3HC1, this groove is formed by residues located between L128 and L144 (Supplemental Figure S10D) and resembles, to some extent, the so-called IBM (IAP binding motif) groove of the type II BIR domains (e.g., Cossu *et al*, 2019; later, see also Supplemental Discussion 1). By contrast, such a groove was not evident in the BLD2 of *Hs*ZC3HC1; there, a few protruding residues, exposed on the BLD’s surface and evolutionarily relatively conserved, were predicted to be positioned instead (Supplemental Figure S10D, and further below).

However, despite the structural similarities between the BLDs’ and BIR domains’ central parts and additional features common to the BIR and one or the other BLD, all of the BIR domains nonetheless turned out structurally more similar among themselves than the ZC3HC1 homologs’ BLD1 and BLD2 domains compared to each other. While sharing several structural features, the BLDs were easily distinguishable nonetheless by additional α-helices inserted into the one or other BLD at different positions.

Regarding the central parts of BLD1 and BLD2, such structural difference manifested itself primarily by a rather extended, evolutionarily conserved α-helix as part of the BLD1 module (Figure 6A; light pink-colored α-helix in Figure 6C, ranging from F134 to T150 in *Hs*ZC3HC1). Such an α-helix is absent from BLD2, and an equivalent α-helix is not part of the BIR domains either (Supplemental Figure S10E). Moreover, predicted structures flanking the two BLDs’ central parts also included α-helices seemingly specific for the BLD2 vicinity and absent in the BLD1. One of these BLD2-specific α-helices (light blue-colored in Figure 6C and Supplemental Figure S10E2, ranging from E211 to E231 in *Hs*ZC3HC1) was predicted to occur positioned outside of the initially defined BLD2 region (Kokoszynska *et al*, 2008, and Figure 1). This α-helix is present in many ZC3HC1 homologs while absent, e.g., in some but not other families of the nematodes (our unpublished data). The other BLD2-specific α-helix, evolutionarily widely conserved and also located beyond the initial BLD2 boundaries (light yellow-colored α-helix in Figure 6C and Supplemental Figure S10E2), ranges from G457 to S471 in *Hs*ZC3HC1. However, even these two α-helices, the one in its entirety and the other with almost all of its residues, were part of the minimalist *Hs*ZC3HC1 deletion mutant we had found still capable of NE-association (Figure 1). Of further note, at least the light blue-colored BLD2-specific α-helix was found absent from not only the BLD1 but also the BIR domains (Supplemental Figure S10E). Altogether, these predictions indicated that both BLD1 and BLD2 include features that distinguish them structurally from the BIR domains.

At this point, there was yet another conspicuous α-helix to be addressed. Initially, this one, present in all three homologs (orange-colored α-helices in Figure 6C), appeared specific for the BLD1, before we only here now assign a corresponding α-helix also to BLD2. In the case of the *Hs*ZC3HC1 BLD1, this α-helix was predicted to start at K75, meaning that all (Kokoszynska *et al*, 2008) or some of this domain’s residues (Higashi *et al*, 2005; Figure 1) were already part of the initially defined BLD1 region. Of particular note, several residues of this BLD1 α-helix appeared to contribute to the establishment of the above-mentioned BLD1:BLD2 interface (Supplemental Figure S9B), while several of its other residues appeared to engage in intra-BLD contacts, as addressed further below. Moreover, having found that a mutant version of *Hs*ZC3HC1 starting at S72 was well capable of binding to the NE, while deletion of aa 1–81 had abolished such binding (Figure 1), we could now conclude that integrity of this particular α-helix, ranging from at least aa K75 to T84, was essential for a functional bimodular NuBaID of *Hs*ZC3HC1 capable of NB-binding. Thus, when newly defining the boundaries of BLD1, this α-helix would need to be included.

So far, however, the corresponding α-helix associated with BLD2 had not been considered part of this BLD. Instead, the N-terminal BLD2 boundary had been regarded as located at H226 or even further away from BLD1. On the other hand, the α-helix we were referring to as the orange-colored one of BLD2 comprises aa A176 to C190. Since we had found deletion mutants lacking either aa 170–210, aa 170–188, or aa 170–178 to all be incapable of NE-association (Figure 1, Supplemental Figure S2A), one could thus consider the integrity of also this BLD2 α-helix as likely essential for NB-binding of *Hs*ZC3HC1.

While this α-helix of BLD2, unlike the orange-colored one of BLD1, did not appear to contribute directly to the BLD1:BLD2 interface (Supplemental Figure S9B2), both of these α-helices had one particular feature in common that we regarded as marking them as equivalent nonetheless. This property was their possession of an evolutionarily conserved arginine of apparently similar function. In the human BLD2, we found this arginine positioned at R185, while the corresponding one of BLD1 locates at R81 (Supplemental Figure S10E2), with each appearing to be involved in stabilizing the position of its α-helix relative to the BLD’s more central parts. Moreover, each one was predicted to contribute to additionally stabilizing the position of one of the aromatic residues we had studied earlier, namely W158 and W431, and such intra-BLD residue contacts, corresponding to R81:W158 and R185:W431, turned out to be characteristic for *Dd*ZC3HC1 and *Sc*Pml39p too (Supplemental Figure S10F and S10G).

Consequently, guided by the holistic consideration of the outcome of our ZC3HC1 deletion experiments (Figure 1; Supplemental Figure S2A), by AlphaFold2’s structure predictions, and Chimera’s structure-based alignments of the different homologs’ sequences, we now regarded the BLD1 of *Hs*ZC3HC1 as at least comprising aa K75 to F167. Its BLD2, correspondingly, would then start at A176 and reach up to S471, even though the very last residues of this BLD’s C-terminal α-helix had been found not essential for NE association (Figure 1). However, dispensability of a few residues of this particular α-helix of BLD2 appeared in line with the corresponding α-helix of *Sc*Pml39p, naturally much shorter than in *Hs*ZC3HC1 (Figure 6C), being sufficient for the yeast homolog’s binding to the NB.

The schematic depiction of the newly defined BLDs’ regions, now also accordingly for *Dd*ZC3HC1 and *Sc*Pml39p, illustrated how close the BLD1 and BLD2 modules of all three homologs are actually positioned next to each other also with regard to the homologs’ primary sequences (Figure 6D). Furthermore, the redefinition of the BLDs’ boundaries meant that both BLDs actually differ in size, ranging from 93–107 aa for BLD1 and 119–144 aa for BLD2 for the three here presented homologs, when not counting the loop-like sequence insertions that are part of the BLD2 of many organisms.

Regarding the newly defined entirety of each of the two BLDs, it is now apparent that each exhibits its unique characteristics, clearly distinguishing one from the other, and both BLDs from the related BIR domains, while nonetheless all sharing also essentially identical structural elements. Of further note, it is now also evident that the Pfam database’s zf-C3HC motif encompassed residues that corresponded to the entire BLD1 and part of BLD2, while the Rsm1 motif, which so far had only been assigned to *Hs*ZC3HC1 but not to *Dd*ZC3HC1 or *Sc*Pml39p, referred to some parts of BLD2 while missing others (Figure 6D). Among the residues now newly assignable to BLD2, and thus, in principle, also to an Rsm1 motif, was the R185 of *Hs*ZC3HC1 BLD1. As mentioned above, we had found this residue corresponding to R81, the latter also present in *Dd*ZC3HC1 and *Sc*Pml39p and part of the Pfam zf-C3HC motif.

Finally, with AlphaFold2’s predictions having allowed for more accurately delineating the ZC3HC1 homologs’ NuBaID and its BLD modules, it is now even more apparent that these boundaries are close to those of the minimal NE-binding-competent mutant of *Hs*ZC3HC1 (Figure 1). The structure predicted for this mutant, comprising 289 aa, reveals a compact polypeptide mainly composed of the two BLDs with only a few residual unstructured parts (Supplemental Figure S11A), again illustrating that it is the two BLDs’ largely integrity (Figure 6D) that is essential and sufficient for the initial binding of ZC3HC1 to the NB.

## Discussion

In the current study, we identified and characterized the protein domain of *Hs*ZC3HC1 that allows for its initial interaction with the NB and the NPC-anchored TPR polypeptides. Based on a combination of experimental data, the definition of a novel NuBaID signature, sequence database mining, and AI-based structure predictions, we demonstrate that the NuBaID, composed of two adjoining modules, is an evolutionarily conserved functional and structural entity, which also functions as the NB- and Mlp1/2-binding domain of the *Sc*Pml39 protein that we here present as the budding yeast’s ZC3HC1 homolog. In the following, we discuss some of the conjectures and conclusions one can draw from the current study’s findings.

### Introducing the NuBaID of ZC3HC1 as the evolutionary conserved bimodular TPR interaction domain of a unique protein present in numerous eukaryotes

The NuBaID that allows ZC3HC1 to bind to TPR at the NPC includes two predicted zinc ion coordination units that we found both essential for such binding. Together they comprise six cysteines and two histidines as part of an evolutionarily conserved signature, with the minimal consensus for each of the two zinc fingers reading C-X_(2)_-C-X_(>14)_-H-X_(3)_-C in vertebrates and the majority of species across the eukaryotic realm that possess a ZC3HC1 homolog. In addition, several classes of fungi have come up during evolution with different spacing between the first two cysteines of the first BLD’s zinc finger signature, reading C-X_(3)_-C instead of C-X_(2)_-C, with such spacing here found also permitting NB-association of *Hs*ZC3HC1.

Our finding of a modular built NuBaID is actually in line with the abovementioned study (Higashi *et al*, 2005), in which data mining for homologs of *Arabidopsis* IAPs had resulted in identifying several proteins that were then called ILPs, including a human ILP that we now know to be ZC3HC1, and described as containing two zinc fingers of the C2HC-type. Equivalent to the CCHC arrangement of the two zinc fingers of the here outlined NuBaID, such former designations should nonetheless neither be confused with the likewise called C2HC-type of zinc fingers, with its minimal consensus C-X_(4)_-C-X_(12)_-H-X_(5)_-C (Kim & Hudson, 1992), nor with the so-called CCHC-type of zinc finger, also known as the zinc knuckle, with its minimal consensus C-X_(2)_-C-X_(4)_-H-X_(4)_-C (Green & Berg, 1989). We further wish to point out that the actual similarities and differences between the BLDs’ zinc fingers and the zinc finger of the human IAPs’ BIR domain, with its minimal consensus C-X_(2)_-C-X_(16)_-H-X_(6)_-C, are addressed in detail in the Supplemental Discussion 1.

The first half of the NuBaID signature, applying to the first of the predicted two zinc fingers, resembles, to some extent, the Pfam database’s zinc finger motif called zf-C3HC (Finn *et al*, 2006; http://pfam.xfam.org/family/zf-C3HC). Indicating a total of four specific cysteines (see also Supplemental Figure S2D1), we can imagine that this motif’s name had been eponymic for the gene’s name *ZC3HC1* (zinc finger C3HC-type protein 1; https://www.genenames.org/data/gene-symbol-report/#!/hgnc_id/HGNC:29913). The NuBaID signature, however, does not include this fourth cysteine as we found it dispensable for NE-binding and TPR-interaction. Moreover, numerous proteins across the eukaryotic realm that we consider ZC3HC1 homologs do not possess such a fourth cysteine (also see Supplemental Figure S10H).

Furthermore, when we database-mined protein sequences from all across the eukaryotic realm for such that likely represent ZC3HC1 homologs, it became evident that the second half of the NuBaID signature, applying to the predicted other zinc finger, was often but not always predicted to overlap with sequence segments that included a Pfam signature called the Rsm1 motif, or parts thereof. This motif’s name stems from the fission yeast protein Rsm1p (Yoon, 2004) that we, like others (Higashi *et al*, 2005), regard being the homolog of *Hs*ZC3HC1 in *Schizosaccharomyces pombe,* with *Sp*Rsm1 too containing a prototypic NuBaID signature, and now also a structure (https://alphafold.ebi.ac.uk/entry/O94506), as predicted by AlphaFold2, whose BLDs resemble those of *Sc*Pml39p.

Our finding that not all prototypic NuBaID proteins are predicted to possess an Rsm1 motif might, in parts, be explainable by the highly variable spacing between the C-X_(2)_-C tetrapeptide and the H-X_(3)_-C pentapeptide of the NuBaID’s second zinc finger. These insertions can include loop-like connecting sequences that are sometimes so extraordinarily long that the Rsm1 consensus might not always tolerate them. Furthermore, in some homologs, very long sequence insertions also exist between the G-W dipeptide and the C-X_(2)_-C tetrapeptide of the BLD2, e.g., in *D. discoideum*, *A. thaliana*, and *C. reinharditii* (see also Supplemental List of Sequences), and these insertions too might sometimes prevent the assignment of an Rsm1 motif to these ZC3HC1 homologs. On the other hand, it is noteworthy that the Rsm1 motif is also not detected in some of the other genuine NuBaID proteins in which the spacer sequence is very short, like in the case of *Sc*Pml39p, where it consists of only 16 amino acids. In again other cases, though, the reason for an Rsm1 motif not being detectable would simply be that the corresponding ZC3HC1 homolog has been mutated in the course of evolution and lacks a BLD2 domain (e.g., Figure 3B1 and Supplemental List of Sequences), as later will also be discussed further below.

While we regard a detailed study of the phylogenetic history of ZC3HC1, with its presence and fate in different organisms, as a topic beyond the scope of the current work, we nonetheless summarize and discuss some of these findings here briefly: Clearly, whenever a sequence was predicted to possess a zf-C3HC signature together with an Rsm1 motif, or even when only one of the two was part of an incomplete sequence in hands, we eventually were able to class the corresponding protein as a ZC3HC1 homolog, or at least as a fragment thereof, next to some also naturally occurring mutated versions of ZC3HC1. Such assignment was based on the protein’s possession of a prototypic NuBaID signature or at least unquestionable parts thereof, the latter then usually affirmed by a few additional residues, which too are characteristic for this type of protein even though some are evolutionarily less conserved and mostly specific for either only BLD1 or BLD2 (e.g., Supplemental Figure S2D1 and S10H). Moreover, apart from some rare ambiguous exceptions, we found a species’ ZC3HC1 homolog to consistently co-occur with a genuine TPR/Mlp homolog, indicating that such a co-existence might be mandatory for ZC3HC1 to persist within a taxon. On the other hand, such co-existence does not appear needed for TPR, which also exists in organisms without an evident ZC3HC1 homolog.

Furthermore, apart from the minimal NuBaID signature and a few other residues, sequence conservation of the ZC3HC1 homologs’ other parts appeared, in general, relatively poor. Hence, this also suggested that these proteins’ common properties would be defined by and depend on the NuBaID. Especially noteworthy, species with a non-duplicated genome were found possessing at most only one NuBaID-containing protein, indicating early on that the NuBaID in these species stands for a particular task to be carried out by a one-of-a-kind protein only.

However, since the NuBaID signature does not appear to define the TPR-binding interface, as discussed further below, and since the residues contributing to this interface still need to be determined in more detail, it remains to be clarified whether the two or more ZC3HC1 paralogs of those species in which genome duplications have occurred, all represent TPR-binding proteins. The latter question holds, for example, for the *Arabidopsis* ILP proteins At1g17210 and At1g48950 (e.g., Higashi *et al*, 2005), and it holds even more so for the three or more ZC3HC1 homologs found in a few species, as in plant genera that have undergone several rounds of genome duplication. The resistance of at least two such plant ZC3HC1 paralogs and their NuBaID signatures against the pressure of evolutionary elimination could mean that one of the two has acquired yet another BLD-involving function, the latter then possibly plant-specific, as also considered for other pairs of paralogs that have persisted in plants (e.g., Blanc & Wolfe, 2004; Veitia, 2005). In this context, we regard it as of note that one of the *Arabidopsis* paralogs of ZC3HC1, At1g48950, was isolated in a genetic screen for proteins involved in the regulation of DNA demethylation pathways, with the gene’s inactivation leading to hypermethylation phenotypes and to naming the protein MEM1 (methylation elevated mutant 1; Lu *et al*, 2020). Further investigations then revealed that MEM1 is also involved in preventing genomic DNA damage (Wang *et al*, 2022). We now consider it of particular interest whether MEM1, which we note to be representing the 594 aa-long and thus shorter one of the two ZC3HC1 paralogs in *Arabidopsis*, will turn out located at the plant’s NBs, in order to there fulfill its function. Moreover, we currently wonder (i) whether the longer paralog of 958 aa, with its conspicuously long loop-like insertion (Supplemental Figure S12A; Supplemental List of Sequences), might exhibit similar or other properties, (ii) where it will turn out located, and (iii) whether both paralogs interact with the *Arabidopsis* homolog of TPR or with different proteins. So far, though, AlphaFold2’s predictions of the BLDs of the two *Arabidopsis* paralogs have not revealed pronounced differences that would hint at one of them unambiguously no longer being able to bind to TPR. Both paralogs’ BLD1 and BLD2 domains appear similarly constructed and equipped with the BLD-characteristic α-helices that are also part of the corresponding BLDs of *Hs*ZC3HC1, *Dd*ZC3HC1 and *Sc*Pml39p (our unpublished data). Nonetheless, without full knowledge of all prerequisites required for a functional TPR binding interface, it remains a matter of speculation whether both bind to the NB or not.

In the case of *Hs*ZC3HC1, though, its NuBaID currently appears monogamous for only one stably to be bound protein, namely *Hs*TPR, also since other proteins formerly proposed as regular ZC3HC1 binding partners, like components of the SCF complex, CCNB1, and FANCD2 (e.g., Bassermann *et al*, 2005; Kreutmair *et al*, 2020), have been ruled out (Gunkel *et al*, 2021) or assessed as unlikely (Supplemental Figure S13). However, we do not exclude the possibility that some ZC3HC1 paralogs, like in plants, have established binding interfaces for stably interacting partners other than TPR that are yet unknown.

Therefore, as long as we cannot be sure that the current study’s NuBaID signatures will turn out to mark every ZC3HC1 paralog with such a signature as a nuclear basket-interacting protein, one could argue that one should conceive another, more universally applicable naming for this signature, without already assigning a function to it. Thus, we can imagine proposing various alternative names, for example, the zf-C2HC-tandem, the BLD-bimodule, the dual-BLD, or simply the zf-(C2HC)2 motif. Apart from that, we wish to propose to those with corresponding expertise to consider merging the NuBaID, the zf-C3HC, and the Rsm1 signatures into a single, all-characteristics-encompassing and then again database-deposited novel signature, since present knowledge now allows to argue that the current individual signatures all represent the same one-of-a-kind type of protein.

### To Have or Have Not: The absence of a ZC3HC1 homolog in some species and a trade-off underlying its presence in others?

Both the presence of ZC3HC1 in many organisms across the eukaryotic realm and its absence in others raises many questions. Some of these will be addressed in the following.

On the one hand, the ZC3HC1 homolog of diverse organisms was found to be a non-essential protein, like in budding and fission yeast (Yoon, 2004; Palancade *et al*, 2005), in the nematode *C. elegans* (Rual *et al*, 2004; Sönnichsen *et al*, 2005), and in mice (e.g., Illert *et al*, 2012; Aherrahrou *et al*, 2021; our unpublished data). Moreover, the inactivation of the human *ZC3HC1* gene by CRISPR/Cas9n technology did not notably affect cell growth and normal cell cycle progression of tumor and non-tumor cell lines (e.g., Hart *et al*, 2015; Gunkel *et al*, 2021). In mice, however, some of the phenotypes and limitations that come along with no ZC3HC1 being around would probably prevent them from surviving for long in a competitive natural environment outside their cage. While such KO mice often appear lively and in a metabolically seemingly advantageous condition, they generally are far more slender and fine-boned than their wild-type kinship, often further accompanied by one or the other kind of a skeletal abnormality (our unpublished data; www.mousephenotype.org/data/genes/MGI:1916023; www.informatics.jax.org/marker/MGI:1916023). On the other hand, though, the disadvantage of no longer possessing a ZC3HC1 homolog appears less evident for other organisms, and those seemingly naturally devoid of a ZC3HC1 homolog appear to come along fine with its absence anyhow. On the one hand, this raises the question of which general, species-spanning advantage ZC3HC1 provides in the numerous organisms where some selection pressure ensures that this protein persists. On the other, one could wonder which property of ZC3HC1 might have been waived in those organisms that no longer possess this protein, having turned dispensable for them at some point, or perhaps even disadvantageous. Furthermore, these questions inevitably come along with another one: *TPR* in some organisms, including mammals and insects, is an essential gene, meaning that those TPR polypeptides appended to the mammalian NPC independently of ZC3HC1 are indispensable, which we had to realize also being the case in cultured human cells, in which it turned out impossible to generate TPR KO cell lines by CRISPR/Cas9n technology (our unpublished results). However, the ZC3HC1-appended TPR polypeptides apparently are not essential since currently known ZC3HC1 KO organisms are viable and since one could presume that absence of ZC3HC1 caused by evolution could also come along with the inability to append additional amounts of TPR to the NB. Thus, one also needs to ask why such TPR subpopulations would be dispensable.

It is evident, though, that such ZC3HC1-deficient organisms can still possess an NB, attested by its presence in insects. In these, the NBs have been studied in the relatively large nuclei of the salivary gland cells of the midge *Chironomus tentans* (Kiseleva *et al*, 1996, 1998), which belongs like *Drosophila* to the order Diptera, possesses a TPR homolog (e.g., Soop *et al*, 2005) just as likely all insects do, but lacks an identifiable ZC3HC1 homolog, just as it is the case for most insect orders. Nonetheless, the midge’s NB structure appears similar to the NB commonly regarded as prototypic in the ZC3HC1-containing vertebrate oocyte. Even though the reported length and diameter of the insect NB’s fibrils (Kiseleva *et al*, 1996) would have them be shorter and thicker than those in vertebrates, there described as more extended and notably thinner (e.g., Ris, 1991, 1997; Jarnik & Aebi, 1991; Gunkel *et al*, 2021), these differences likely reflect, at least to a large part, the outcome of different sample preparation procedures, with some including the specimens’ coating with relatively thick layers of heavy metals.

However, if one assumes that no conspicuously different structures are hidden beneath such coats of metal that would distinguish the insects’ from the vertebrates’ NB fibrils, this would raise the next inevitable question; namely, where those TPR polypeptides are positioned that are appended to the vertebrate NB by ZC3HC1, with the same question also applying for yeast and its Pml39p-dependent Mlp1p subpopulation. In theory, a thinkable answer would sketch a scenario in which there actually are no significant differences between the insects’ and the vertebrates’ NBs, simply because insects would still possess a ZC3HC1 homolog whose NuBaID signature merely diverged from the current consensus beyond recognizability during insect evolution, while still acting as a TPR-interlinking structural NB component nonetheless. One could then speculate whether this might have come along with, for example, other combinations of cysteine and histidine residues or with residues like aspartate and glutamate as zinc ion coordinating ligands instead of the one or other of the NuBaID’s cysteines and histidines (Laitaoja *et al*, 2013). Along a similar line, one could alternatively also conceive that the function of ZC3HC1 at the NB has been taken over by another yet unknown insect protein capable of binding to the NB and of NB-appending TPR polypeptides in a manner analogous to ZC3HC1. However, while we deem such scenarios not yet proven ruled out, we regard the other signs pointing at a ZC3HC1 homolog having been lost without substitution in various organisms, including insects, as more compelling. We felt this assumption also underscored after having searched AlphaFold’s database with the recently available protein structure search tool Foldseek (van Kempen *et al*, 2022). The latter allowed us to seek protein structures of *Drosophila melanogaster* that might come into question as structural homologs of *Hs*ZC3HC1, *Dd*ZC3HC1, or *Sc*Pml39p. However, other than known BIR proteins, these searches, for now, did not reveal *Drosophila* structures that we would instantaneously regard as likely candidates. By contrast, when searching the database-deposited human, amoebic, and budding yeast structures with either the *Hs*ZC3HC1, *Dd*ZC3HC1, or *Sc*Pml39p structure as the only query, the two other species’ ZC3HC1 structures were in each case readily identifiable as the best matches (Supplemental Figure S14).

However, concerning the NB’s structure, this situation would momentarily leave us with neither a homolog nor an analog of ZC3HC1 in insects, and consequently, no additional TPR amounts appended to the insects’ NB, with such a scenario confronted by micrographs of similarly looking NBs in both insects and vertebrates. This, in turn, leaves us with the task of providing an answer elsewhere (Gunkel *et al*, manuscript in preparation) as to how at least the ZC3HC1-dependent TPR polypeptides in vertebrates are arranged at the NB.

Next, again presuming that various eukaryotes once possessed a functional ZC3HC1 homolog, yet from some point on no longer, this brings us back to asking how this might have come about and for what reason. While several scenarios are considered in Supplemental Discussion 2, we outline one of them also here. In such a scenario, we can imagine that the presence of a ZC3HC1 homolog might have turned disadvantageous for some species in certain situations and environments, with evolutionary forces then expediting its elimination. Such a notion of a trade-off between advantages and disadvantages finds support in several systematic studies in *S. cerevisiae*. In these, homozygous *pml39*Δ cells were found well viable in a range of growth conditions that are considered approximations of typical environments experienced by wild, domesticated and laboratory yeast strains, with *PML39* deficiency in these conditions not or only minimally affecting the KO cells’ competitive fitness as compared to the WT strains (e.g., Breslow *et al*, 2008; Qian *et al*, 2012). However, when exposed to a plethora of chemical, physical or nutritional stress conditions, the *pml39*Δ cells not only turned out hypersensitive to some of these treatments, whereon they exhibited a notably decreased competitive fitness, but they also clearly outclassed the WT strains in other conditions (Brown *et al*, 2006; Hillenmeyer *et al*, 2008). While a detailed reflection on the wealth of information on Pml39p in these systematic studies goes beyond the scale of the current discussion, we wish to point out some findings. For example, while *pml39*Δ cells were found to be more sensitive to some kinds of acute stress and the triggering of distinct signal transduction pathways, they were more tolerating than the wild-type when it came to nutritional deficiencies or other types of stress (Brown *et al*, 2006; Hillenmeyer *et al*, 2008). Altogether, these screening data prompt the conclusion that the existence of Pml39p in budding yeast reflects a balancing act between pros and cons.

We also find it remarkable that many of the especially prominent phenotypes observed with the *pml39*Δ strains were similarly pronounced in *mlp1*Δ strains, while they were notably different from those of the homozygous or heterozygous deletion strains of some of the other known NB-associated proteins like Mad1p, Sac3p, Ulp1p, and, in particular, also Mlp2p (Hillenmeyer *et al*, 2008). We consider this finding in line with the notion of a special structural-functional relationship between Pml39p and Mlp1p. On the other hand, though, a few other phenotypes appear to be specific, at first sight, for Pml39p, which could indicate some additional, standalone functions of this protein. However, we can also imagine that such findings might point to those Mlp1 polypeptides that occur NB-attached via Pml39p, with us anticipating that some of their functions are distinct from those Mlp1 polypeptides anchored to the NPC independently of Pml39p. Finally, in the context of Pml39p’s function, we regard it as noteworthy that some of the disadvantages caused by homozygous deletion of Pml39p are also observed, though less pronounced, in the heterozygous Pml39p mutant (Hillenmeyer *et al*, 2008), pointing at haploinsufficiency and the need for sufficient copy numbers of Pml39p to fulfil its function properly.

In the context of trade-offs between advantages and disadvantages in different life stages, we also regard it as noteworthy that Pml39p belongs to a minor group of less than 4% of all protein-coding yeast genes whose deletion notably extends the yeast’s replicative lifespan (McCormick *et al*, 2015). This finding, in turn, raises the question of whether this might be due to the enhanced tolerance toward oxidative stress in the absence of Pml39p (Brown *et al*, 2006; Hillenmeyer *et al*, 2008).

Furthermore, indications that the absence of a ZC3HC1 homolog might also have some moderate life span-prolonging effect in mammals come from studies in ZC3HC1 KO mice (Aherrahrou *et al*, 2021). Such effect, in turn, appears in line with elderly ZC3HC1 KO mice outperforming, concerning some physiological parameters, their wild-type siblings with whom they shared a cage for year-long periods (our unpublished data). Such observations of ZC3HC1 KO mice possibly being more long-living, despite those deficits they had to live with since being juveniles, and then barely exhibiting additional deficits at a higher age, in contrast to their ZC3HC1-possessing siblings, underscores the notion that lacking a ZC3HC1 homolog does not only need to come along with irrelevance in some species and disadvantages in others but can also have its benefits. Such advantages would then manifest themselves at a later life stage. Conversely, regarding the WT mouse, ZC3HC1 would thus manifest most of its benefits in young adults but start revealing its disadvantages at an older age, with such a change in the weighting of properties conforming to the original definition for antagonistic pleiotropy (Williams, 1957).

Provided that the existence of a unique but non-essential *ZC3HC1*/*PML39* gene in some organisms and its absence in others would thus indeed reflect a dynamic balance between advantages in certain situations and life stages and disadvantages in others, *ZC3HC1* would also conform to the current, broader definition of a gene whose possession comes along with antagonistic pleiotropy and thus multiple opposing effects on fitness (e.g., Kirkwood, 2002; Gavrilov & Gavrilova, 2002; Elena & Lenski, 2003; Mitchell-Olds *et al*, 2007; Montano & Long, 2011; Qian *et al*, 2012; Nikolin *et al*, 2012; Anderson *et al*, 2011, 2013; Austad & Hoffman, 2018). If so, the absence of ZC3HC1 in some organisms would no longer be interpretable as the fate of a gene that has turned out useless and no longer required at some point. Instead, it then would reflect a gene whose protein has turned into being more detrimental than advantageous, eventually exhibiting an unbearable property that demanded to be selected against by evolutionary forces to achieve its elimination.

In conclusion, while future research might quite naturally focus on elucidating the benefits that ZC3HC1, and the TPR it attaches to the NB, will need to convey to those that have a ZC3HC1 homolog not disposed of by evolution, we feel that we should keep in mind that there might also be a dark side to possessing this protein and the ZC3HC1-dependent TPR polypeptides at the NB. As outlined, we can imagine that this protein’s advantages and disadvantages, if existing as such, might not only manifest themselves in different environmental conditions; if ZC3HC1 were indeed the product of a trade-off gene, we could envision its presence causing antagonistic pleiotropic effects also in the young and old. Future work would then need to find out how these effects can be explained by ZC3HC1 and TPR being structural proteins and how one can correlate them with their functions at the NB.

Furthermore, we propose ZC3HC1 as a general model for studying different aspects of both adaptive and neutral protein evolution (e.g., Akashi *et al*, 2012; Galtier, 2016; Albalat & Cañestro, 2016) by gaining insight into which types of evolutionary causes have affected the different ZC3HC1 homologs’ evolutionary history. Being a protein (i) probably non-essential in most organisms, (ii) nonetheless existing as a functional version in many species while appearing mutated or absent in others, (iii) exhibiting some features strikingly different among species, like a sprawling BLD2-inserted loop in some and its complete lack in others, while at the same time (iv) featuring an evolutionarily conserved centerpiece, ZC3HC1 combines all the ingredients that compose a challenging riddle posed by evolution, with such riddle now demanding to be solved.

### Computational predictions reveal commonalities and some differences between the BLDs’ structures, prompting some conclusions and further considerations

The deep-learning program AlphaFold2 (Jumper *et al*, 2021; Tunyasuvunakool *et al*, 2021; Varadi *et al*, 2022) and the general accessibility of its structure database have allowed for gaining a first insight into the likely structures of the different phyla’s ZC3HC1 homologs. AlphaFold2 predicted that the homologs share strikingly similar structural arrangements, despite these proteins’ low overall sequence identity. Some of these findings also allowed us to redefine the BLDs’ expanse, thereby unfolding that the C-terminal part of the zf-C3HC motif actually corresponds to the N-terminal part of the adjacent BLD2.

We regarded the BLDs’ predicted structural similarities as all the more remarkable when we also realized that the homologs’ low degree of sequence identity still manifested itself after having aligned pairwise only those sequences of the different homologs that represented their redefined pairs of BLDs (Supplemental Figure S11B). Altogether, we considered these findings to add to a plausible explanation (see also Supplemental Information 2) for the former reporting that a ZC3HC1 homolog had not been detectable in *S. cerevisiae* when data mining primary sequences (Higashi *et al*, 2005).

In addition to the information gained from our experimental work, the AlphaFold2-predicted structural similarity of the different species’ BLDs allowed for further underscoring our notion that an intact ZC3HC1 homolog possesses two similarly built zinc ion coordination modules that together constitute the NuBaID as an evolutionarily conserved entity. Moreover, such insight into the BLDs’ likely structure also corroborated our assumption that the current version of the NuBaID’s minimal sequence signature only represented residues involved in establishing the BLDs’ central structures and not the binding interface for the different species’ TPR homologs. Therefore, one can further conclude that the integrity of both BLDs’ core regions is also a prerequisite for forming a fully functional TPR binding interface, the latter including residues also positioned elsewhere. In other words, aa substitutions within the NuBaID’s minimal sequence signature that prevent the formation of the domains’ core structures would, in turn, prevent the establishment of a properly functioning total of TPR attachment points.

Noteworthy, too, in this context, only some of the few other ZC3HC1 residues that appear evolutionarily relatively well conserved between the eukaryotes’ different phyla appear to be surface-exposed (Supplemental Figure S10D and S10H), while most seem to be projecting towards or embedded within the BLDs, where they appear to contribute to intra-BLD arrangements, according to the structure predictions (e.g., Supplemental Figure S10H). One could thus further imagine that the TPR binding interfaces of the different species’ ZC3HC1 homologs might eventually turn out not to share more than a few residues evolutionarily conserved among distant eukaryotic phyla.

Furthermore, we consider it particularly noteworthy that each BLD’s core region with its zinc ion coordination sphere is surrounded by conspicuous α-helices, defining the BLD’s outer perimeter. We can imagine that the outwardly exposed parts of some of these α-helices are suitably positioned for and eventually will turn out to be contributing to a TPR binding interface, and we can picture a scenario in which some of these α-helices align with distinct segments of the coiled-coil homodimers of TPR. We can further imagine that such interactions require the different ZC3HC1 α-helices to be in a distinct constellation relative to each other, with such positioning also appearing evolutionary conserved and at least in some cases defined by interactions with the BLD’s central parts. In fact, assuming that those aa substitutions that bear on the zinc ion coordination sphere will not abolish the α-helices as such while abolishing all interactions with TPR nonetheless, one could interpret the effects caused by these mutations by regarding the latter as no longer permitting the establishment of a central BLD core, with this then, in turn, no longer allowing for keeping the BLD’s outward positioned α-helices correctly in place. The currently available AI, though, did not yet allow for assessing the effects of distinct single aa substitutions on the BLDs’ overall structure, in line with AlphaFold2 having been mentioned to have neither been trained nor expected to capture the effect of protein-destabilizing single aa mutations (https://www.ebi.ac.uk/about/news/perspectives/alphafold-potential-impacts/; see also Supplemental Figure S7D). However, we can imagine that future algorithms will eventually also be able to correctly assess the effects caused by single aa substitutions on the protein’s actual folding processes, in addition to the current computing, whose training was guided by intact mature proteins (see also Supplemental Information 3).

Apart from the BLDs’ α-helices, other structural features specific to one or the other BLD stir up curiosity. Among such single-BLD-specific features, we noted, for example, a conspicuous surface groove of the BLD1 in vertebrates and some other organisms (Supplemental Figure S10D; also see Supplemental Discussion 1). By contrast, this groove appears absent or altered in the BLD2, which we noted instead exhibiting, at the BLD1-groove-corresponding position, the abovementioned small cluster of evolutionarily relatively well conserved surface-exposed residues. Next to the BLDs’ α-helices, the functions of also these and other single BLD-specific features will need to be a topic of research that will eventually have to clarify which parts of the two BLDs engage in direct interactions with TPR as part of the ZC3HC1 protein’s TPR-binding interface.

It also remains to be determined whether the commonality shared by all the natural ZC3HC1 homologs with an intact NuBaID might be restricted to their binding to the NB and the respective organism’s TPR homolog or whether all of them are also capable of recruiting yet further TPR polypeptides. We do not regard it as unlikely that some ZC3HC1 homologs, even with an intact NuBaID and possibly well capable of binding to their species’ NBs, might nonetheless lack the potential to recruit additional TPR polypeptides. In other words, while we regard an intact NuBaID that allows for an intact TPR binding interface as sufficient for NB association, we can imagine that this on its own is generally, or even categorically, not sufficient for the subsequent recruitment of TPR polypeptides. Instead, we regard it as likely that such recruitment capability requires additional sequence segments of ZC3HC1 that might not be part of all homologs (our unpublished data). This assumption, in turn, raises the question of why evolution would have allowed some species to possess a ZC3HC1 homolog with a prototypic NuBaID capable of NB-binding but then be unfit to recruit more TPR.

Future work will thus need to scrutinize whether the presumed properties of some ZC3HC1 homologs indeed match our prediction of them not being capable of recruiting TPR and whether the binding of only ZC3HC1 itself to the NPC-anchored TPR polypeptides might already be functionally relevant on its own, like in a scenario already outlined earlier (Gunkel *et al*, 2021). Back then, we had pondered whether ZC3HC1 might also act as an adjustable insulator of TPR segments, with such function possibly even uncoupled from the recruitment of additional TPR polypeptides. Such a simple setting would nonetheless allow for additional, species-specific functions of ZC3HC1, for example, by the one or other type of BLD2-inserted loops present in the one but not other species.

### Intrinsically disordered regions as a characteristic feature of many but not all ZC3HC1 homologs

Having mapped *Hs*ZC3HC1 for those parts required for its binding to the NB, we found some segments dispensable, with the most prominent one likely to be largely unstructured, comprising more than 100 aa embedded within the NuBaID’s second BLD. Moreover, our subsequent search for homologs of ZC3HC1 in other eukaryotes then led to the identification of numerous homologs that also harbored large insertions corresponding to the major one of *Hs*ZC3HC1. The largest ones of these unstructured loops comprised even more than 500 amino acids, with those in the ZC3HC1 homologs of the stick insects of the superorder Orthopterida possibly exceeding even these sizes by far, which raises further questions as to their presence in these insects (Supplemental Figure S12A) while ZC3HC1 homologs apparently no longer exist in other Neoptera insects. Furthermore, and by similarly striking contrast, no such additional sequence insertion was present within the BLD2 of other genuine homologs of ZC3HC1, as exemplified by *Sc*Pml39p.

Except for a few conserved residues, which are part of Pfam’s Rsm1 motif (e.g., Supplemental Figure S10H) and located at the transitions between the central part of the BLD2 and the loop-like insertion, these insertions do not share any obvious sequence similarity between the ZC3HC1 homologs of different phyla. So far, we cannot tell which physiological relevance these large loops might have in those species in which they have been favored and expanded by evolution.

While large loop-like insertions do not occur within the first BLD of whichever species’ homolog, suggesting that the BLD1, when located at the NB, faces a binding interface or local environment that generally does not allow for extensive, space-filling appendices, the possibility of having loops BLD2-inserted suggests that they there have the opportunity to occupy free spaces. Particularly in the case of the very long loops, we enjoy wildly speculating that these might even project away from the NB scaffold and perhaps contribute to some highly flexible barriers, with the latter hedging in the perimeter of the NB’s terminal ring and thereby contributing to the demarcation of some distinct cargo pathways, possibly even in a species-specific manner. These loops might then also be targets of cellular signaling processes, resulting in their post-translational modification, thereby allowing for adjusting their performance at the NB’s perimeter.

In fact, in the case of *Hs*ZC3HC1, its BLD2-embedded loop harbors especially many sites that have been found distinctly phosphorylated also in interphase (Supplemental Figure S12B) after a range of different stimuli (e.g., Christensen *et al*, 2010; Moritz *et al*, 2010; Weintz *et al*, 2010; Yu *et al*, 2011; Robitaille *et al*, 2013; Sos *et al*, 2014; Sharma *et al*, 2014; Hu *et al*, 2015, our unpublished data; see also further below). One could now further imagine, for example, that such a more or less phosphorylated loop might then repel specific molecules in a more or less efficient manner. Alternatively, we can also conceive of such ZC3HC1 loops as playing a role as spacer elements, defining distances between the NB and neighboring chromatin. Furthermore, while dispensable for the initial binding of ZC3HC1 to the NB, we can also imagine a scenario in which such loops might play a role as lariats in the regulatable formation of higher-order cylindrical arrangements between TPR and ZC3HC1 of the type discussed recently (Gunkel *et al*, 2021; Gunkel & Cordes, 2022).

Future work aiming at understanding the role of the ZC3HC1 loop might eventually also provide insight into why a distinct aa polymorphism of *Hs*ZC3HC1 has been associated with phenotypes connected to atherosclerosis of coronary arteries, ischemic stroke and distinct blood pressure profiles. Such pathological conditions have been assigned to an arginine at the aa position 363, while a histidine at the same position instead was regarded as a non-effect or protective residue (e.g., Schunkert *et al*, 2011; Deloukas *et al*, 2013; Jones *et al*, 2016; Wirtwein *et al*, 2016; Linseman *et al*, 2017; Jafaripour *et al*, 2019), which thus locates centrally within the large, unstructured loop of *Hs*ZC3HC1. In some studies, such phenotypes have been considered to reflect some dysfunction of ZC3HC1 in its formerly proposed role (Bassermann *et al*, 2005) as an SCF component ensuring CCNB1 degradation (Jones *et al*, 2016; Linseman *et al*, 2017; see also López-Mejías *et al*, 2013; Kunnas & Nikkari, 2015). However, since our recent data have shown that ZC3HC1 is neither an SCF component nor involved in some direct regulation of cellular CCNB1 amounts (Gunkel *et al*, 2021), we can now rule out such a formerly presumed causal relationship.

Momentarily though, we cannot tell how this R to H exchange, or the opposite one, neither of which notably affects the initial binding of ZC3HC1 to the NB, and which locates within a region not sequence-conserved amongst vertebrates, would provoke the different effects assigned to this polymorphism. One could speculate, for example, that the two residues at position 363 might, as also considered elsewhere (Jones *et al*, 2016; Linseman *et al*, 2017; Wang *et al*, 2019), affect phosphorylation of nearby serine or threonine residues differently, perhaps by changing kinase binding preferences (e.g., Ren *et al*, 2010). The phenotypes assigned to this polymorphism might then be related to differences in phosphorylation affecting the yet unknown function of this disordered loop.

### Pml39p as the budding yeast’s homolog of ZC3HC1

Whatever the loop might eventually turn out to be good for, it is evident that the budding yeast’s protein Pml39 does not possess such a loop. Furthermore, we found Pml39p not capable of stably binding to *Hs*TPR, and we neither found *Hs*ZC3HC1 capable of binding to the Mlp proteins. Nonetheless, as already pointed out earlier (Gunkel *et al*, 2021; Gunkel & Cordes, 2022), we regard Pml39p as the one and only homolog of *Hs*ZC3HC1 in *S. cerevisiae*.

We base this conclusion not merely on our finding that the non-essential protein Pml39 has a prototypic NuBaID signature but also on our other experimental results. Some of them confirmed former data that had shown Pml39p locating at the yeast NE (Huh *et al*, 2003) via binding to the Mlp proteins (Palancade *et al*, 2005), just like the non-essential protein ZC3HC1 binds to the NBs of vertebrates via its binding to TPR (Gunkel *et al*, 2021; Gunkel & Cordes, 2022). Furthermore, Pml39p had been found to bind to parts of the Mlp proteins, including regions at the NT (Palancade *et al*, 2005), which correspond to those of TPR that we found binding ZC3HC1 (Figure 2C, and Gunkel *et al*, manuscript in preparation). In addition, we have validated Pml39p as the yeast’s ZC3HC1 homolog by showing that both NB-association and binding to Mlp proteins require the integrity of the Pml39 protein’s NuBaID. Even single aa substitutions of its NuBaID signature already prevented Pml39p from interacting with the Mlps, just like it happens to be so for ZC3HC1 and its binding to TPR.

Furthermore, our finding of the notably diminished NE-attached amounts of Mlp1p in *pml39Δ* cells, in line with recent observations of a similar kind (Bensidoun et al, 2021b), indicates that specific arrangements at the nuclear periphery between subpopulations of Mlp polypeptides depend on Pml39p. Again, such a finding was similar to ZC3HC1 being a protein required to keep TPR subpopulations tethered to the NBs in human cells (Gunkel *et al*, 2021). Moreover, our current study’s finding of Pml39p being a prerequisite for the focal accumulation of Mlp1 polypeptides in *nup60Δ* cells was equivalent to ZC3HC1 having been shown responsible for keeping TPR polypeptides accumulated in NPC-remote foci in NUP153-deficient cells (Gunkel & Cordes, 2022). Apart from these findings, an additional observation hinting at perhaps another commonality between *Hs*ZC3HC1 and *Sc*Pml39 is mentioned in Supplemental Information 4. Finally, with AlphaFold2 predicting striking similarity between the NuBaID’s construction in *Hs*ZC3HC1 and *Sc*Pml39p, we regarded the proteins’ kinship at long last also displayed at the structural level.

ZC3HC1 not having been identified earlier as a Pml39p homolog might have had several reasons. One of them, already mentioned further above, will have been the poor sequence similarity between *Sc*Pml39p and *Hs*ZC3HC1, which prevented finding the other species’ homolog through standard primary sequence alignment searches. The other reason why Pml39p, identified in a synthetic lethal screen with a *nup133*Δ mutant (Palancade *et al*, 2005), appears not to have been generally considered a bona fide NB protein (Köhler & Hurt, 2007, 2010; Grossman et al, 2012; Niepel et al, 2013; Floch et al, 2014; Ptak et al, 2014; Obado et al, 2016; Lin & Hoelz, 2019; Fernandez-Martinez & Rout, 2021; Dultz et al, 2022) might be similar to the reason which for long prevented detecting *Hs*ZC3HC1 as an NB protein: While ZC3HC1 is a protein stably bound to the NB under physiological conditions, it is rapidly detached when exposed to the non-physiological conditions of standard cell fractionation protocols (Gunkel *et al*, 2021; Gunkel & Cordes, 2022). Furthermore, once the genuine *in vivo* interactions between native TPR and ZC3HC1 polypeptides have been disrupted in such a way, notable amounts of these parted proteins do not appear inclined to readily re-associate again *in vitro*, even when having re-instated conditions that more closely again resemble those within cells. Since common yeast and mammalian cell fractionation protocols share non-physiological similarities, we can now imagine that sensitivity towards such conditions also applies to the interactions between the Pml39 and Mlp polypeptides (for further comments along this line, see Supplemental Information 5).

Given the abovementioned commonalities between *Hs*ZC3HC1 and *Sc*Pml39p, the question inevitably arising is whether a function formerly assigned to Pml39p might also apply to ZC3HC1: Pml39p had been described as a protein acting as an upstream effector of the Mlp proteins in the retention of improper mRNPs, with Pml39p regarded as specifically involved in the nuclear retention of incompletely spliced mRNAs (Palancade *et al*, 2005; see also Bonnet *et al*, 2015). In such a model, Pml39p was depicted as a protein using the NB and Mlp proteins merely as an operational platform for interacting directly, in principle also autonomously, with distinct mRNA-binding proteins and potentially also components of the splicing machinery. Such interactions, in turn, were to prevent intron-containing pre-mRNAs from exiting the nucleus (Palancade *et al*, 2005).

Concerning *Hs*ZC3HC1, though, none of our currently available data suggests some autonomous function for ZC3HC1 that the protein would effectively execute when uncoupled from TPR. On the other hand, we cannot tell for sure so far whether there might be, apart from specific kinases, some proteins that only transiently, yet ordinarily nonetheless, interact with ZC3HC1 once it occurs positioned at the NB. However, having investigated whether ZC3HC1 or TPR appended to the NB via ZC3HC1 might be involved in some general surveillance of intron-containing transcripts in HeLa cells, we eventually could conclude that this was not the case. In these cells, we neither found ZC3HC1 nor the ZC3HC1-appended TPR polypeptides playing a universal role in the monitoring of introns and their splice sites in mRNAs (Iino, 2017, and our unpublished data), with these findings in conflict with a former report (Rajanala & Nandicoori, 2012) but in line with TPR not promoting nuclear retention of 5’splice site-containing transcripts in U-2 OS cells either (Lee *et al*, 2020).

However, since such human cells differ from yeasts by a generally far larger nucleus, an abundance of introns, and open mitosis, the bulk of correct splicing will need to be quality-controlled already early on in a human cell, and thus mostly deep within the nuclear interior to avoid work-overload at the NPC and unfulfilled splicing tasks at the onset of mitosis. Therefore, we do not regard our data from human cells as necessarily in conflict with those that argue for one or another type of mRNA quality control directly occurring at the budding yeast’s NPCs and NBs, particularly for subsets of yeast genes transcribed at the nuclear periphery.

We can, though, also imagine interpreting some of the formerly observed phenotypes in an alternative manner, thereby assigning this protein only an indirect contribution. Instead of conceiving Pml39p as directly interacting with different RNA-binding proteins, we consider it possible that some of the phenotypes observed upon the absence and overexpression of Pml39p might have reflected a consequence of subpopulations of Mlp1 polypeptides then having occurred mislocalized. Just like mislocalized subpopulations of TPR happen to be the case when ZC3HC1 has been eliminated or highly overexpressed (Gunkel *et al*, 2021; Gunkel & Cordes, 2022). However, one should also keep in mind that neither the nucleoplasmic Mlp1p amounts in the pml39Δ cells (Figure 5A) nor the large nucleoplasmic pools of soluble TPR in different ZC3HC1 KO cell lines (Gunkel & Cordes, 2022) appear to interfere with normal cell cycle progression, which argues against some TPR/Mlp1p-interacting proteins being mislocalized or sequestered within these cells’ nucleoplasm in pivotal amounts.

To gain further insight into the events occurring at the NB, we now regard it as necessary to also unveil the TPR and Mlp proteins’ molecular contributions to their relationships with ZC3HC1 and Pml39p. We consider such knowledge a prerequisite for any more profound understanding of the supramolecular arrangements at the NB. We furthermore regard it as easily conceivable that only these arrangements at the NB will allow for the one or other type of molecular interactions that can only occur at an intact NB and not with the individual NB components on their own, like when these occur elsewhere within the nucleoplasm.

Now aware that ensembles of TPR and ZC3HC1 homologous to those in vertebrates also exist in other model organisms, we feel confident that each of these species will contribute to providing answers to the plethora of questions still twining around the NB so that it can eventually unfold its mysteries.

## Materials and Methods

### Antibodies, expression vectors, cell lines, and yeast strains

Mouse monoclonal antibody 203-37 against *Hs*TPR (Cordes *et al*, 1997), whose epitope was mapped to a region comprising aa 1462–1500 (Hase *et al*, 2001; Gunkel *et al*, 2021), has been described earlier. Similarly, guinea pig peptide antibodies against *Hs*TPR aa 2063–2084 (Cordes *et al*, 1997) and *Hs*ZC3HC1 aa 307–355 (Gunkel *et al*, 2021) have been described. The same pool of pan-FG-NUPs rabbit antibodies already used earlier (Göttfert *et al*, 2013) represented antibodies obtained after immunization with the FG-repeat domain of the *Xenopus* oocyte NB-associated protein *Xl*GANP. From these anti-*Xl*GANP sera, we had isolated by sequential affinity-chromatography, using a series of overlapping FG domain peptides and recombinant proteins, as illustrated for other proteins (Gunkel *et al*, 2021), antibody subpopulations either specific for xlGANP or cross-reactive with numerous FG-NUPs. The most broadly cross-reactive ones of these pan-FG antibodies targeted all of a comprehensive collection of bacterially expressed and purified FG-repeat domains of *Xenopus* and mammalian FG-NUPs. Novel peptide antibodies against synthetic peptides (Peptide Specialty Laboratories, Heidelberg, Germany), coupled via a C-terminal (i) or N-terminal (ii) cysteine to keyhole limpet hemocyanin, were raised in guinea pigs, followed by peptide affinity purifications using standard procedures. These peptides included such corresponding to aa 3–23 of *Dd*TPR (accession number ON368702) and aa 1–29 of *Dd*ZC3HC1 (ON368701). All secondary antibodies were from Jackson ImmunoResearch (Cambridgeshire, United Kingdom), as listed earlier (Gunkel *et al*, 2021). All expression vectors coding for human and yeast polypeptides are listed in Supplemental Table 1. The human HeLa cell line P2 and its ZC3HC1 KO progeny cell line, obtained after CRISPR/Cas9n-editing, and their growth conditions, have been described earlier (Gunkel *et al*, 2021). The axenic *D. discoideum* strains Ax2, Ax3, and Ax4 (e.g., Bloomfield *et al*, 2008) were kindly provided by Martin Kollmar and Katarina Gunkel. The budding yeast strains used in this study, isogenic to S288c, are listed in Supplemental Table 2, among which are also yeast strains with gene deletions obtained from Dharmacon (Lafayette, CO, USA; Winzeler *et al*, 1999).

### Transfections and transformations for live-cell imaging, immunofluorescence microscopy, and immunoblotting

Culturing of HeLa cells of line P2 and transfection with expression vectors encoding FP-tagged polypeptides, using PolyJet (SignaGen Laboratories, Rockville, MD, USA), followed by cell inspection not later than 24 h post-transfection, was performed as described (Gunkel *et al*, 2021). HeLa cell transfections with small interfering RNAs (Supplemental Figure S13), using HiPerFect (QIAGEN, Hilden, Germany), followed by cell harvest at three days post-transfection, and subsequent IFM, were performed as described, as were immunoblottings of cell extracts (Gunkel *et al*, 2021). The lithium acetate method (Gietz & Schiestl, 2007) was used to transform Mlp1p-mCherry-expressing *pml39*Δ yeast cells with vectors coding for yECitrine-tagged WT and mutant versions of Pml39p. For the galactose-induced expression of the yECitrine-tagged polypeptides, the transformed yeast cells were first grown in a non-inducing medium with raffinose and then transferred into a galactose-containing medium; followed by live cell imaging about 30 min later.

### Live-cell imaging and immunofluorescence microscopy

Live-cell imaging and IFM of cultured HeLa cells of line P2 were performed as described (Gunkel *et al*, 2021; Gunkel & Cordes, 2022), using a 63x Lambda objective and a Leica TCS SP5 or SP8 confocal laser-scanning microscope (Leica Microsystems, Wetzlar, Germany) for image acquisition. FLIP of transiently transfected HeLa ZC3HC1 KO cells or HeLa WT cells ectopically expressing FP-tagged polypeptides was conducted with the Leica TCS SP5. Selected areas were subjected to pulses of full laser power for 2 min, usually followed by the acquisition of post-bleach images. Additional brightness-enhancement of a subset of images (Figures 1B2 and 2B, and Supplemental Figure S2) carried out after the image acquisition was as described (Gunkel & Cordes, 2022). For live-cell imaging of yeast, cells from a culture in the logarithmic growth phase were transferred into Ibidi µ-slides with 18 wells (ibidi µ-Slide 18 Well ibiTreat; ibidi GmbH, Gräfelfing, Germany). Once the cells had settled, they were imaged at RT with a 63x Lambda objective at the Leica SP5 confocal microscope or with a 63x NA1.4 objective at the ZEISS LSM880 with Fast Airyscan detector microscope (Carl Zeiss, Oberkochen, Germany). Fiji/ImageJ (versions 1.50i–1.51t, National Institutes of Health, USA; Schneider *et al*, 2012) was used for quantifying the NE signal intensities on the yeast cell micrographs obtained by live-cell imaging, applying the quantification procedure described earlier (Gunkel *et al*, 2021). Suspended *Dictyostelium* Ax2 cells were grown in HL5 medium with glucose (HLG01, Formedium, Hunstanton, United Kingdom) for 2–3 days to reach confluency in a culture dish, from which they were then detached by gentle pipetting, collected in a 15 ml tube, and centrifuged at 300 *g* for 3 min. The sedimented cells were washed in 10 ml PBS with 10 mM of freshly added MgCl_2_. Centrifugation and resuspension of the sedimented cells in PBS with MgCl_2_ were repeated, followed by pipetting 400 µl of cell suspension per well onto 12 mm #1.5H coverslips (Gerhard Menzel B.V. & Co.KG, Braunschweig, Germany) already placed in a 24-well plate. The plate was centrifuged in a swing-out rotor at 300 *g* for 3 min to allow the cells’ homogenous settling onto the coverslips. The PBS solution was then carefully aspirated and immediately replaced with a fixation solution containing 1% formaldehyde and 0.54% methanol in PBS (formalin solution diluted 1:37) and 0.1% TX-100. After incubation for 15 min, the fixed and permeabilized samples were quenched with 50 mM NH_4_Cl in PBS for 5 min, blocked in 1% BSA in PBS for 30 min, and then immunolabeled as described for cultured human cells earlier (Gunkel *et al*, 2021). Fiji/ImageJ was used for generating line profiles across immunolabeled *Dictyostelium* nuclei.

### Cloning of Dictyostelium discoideum cDNAs

For the isolation of total RNA, *D. discoideum* Ax4 cells were (i) homogenized in TRIzol reagent (Invitrogen, Carlsbad, CA, USA), followed by (ii) RNA purification and on-column-digestion of genomic DNA using the Direct-zol RNA Miniprep kit (Zymo Research, Freiburg, Germany), and (iii) subsequent cDNA synthesis using oligo(dT) primers and final RNA digestion with the Superscript III First-Strand Synthesis System (Invitrogen), thereby performing all steps according to the manufacturers’ instructions. Primers used for PCR, subcloning and sequencing of *Dd*TPR and DdZC3HC1 sequences are listed in Supplemental Table 3. cDNA sequences were deposited in the Genbank database (https://www.ncbi.nlm.nih.gov/genbank/), the latter providing accession numbers ON368701 (*Dd*ZC3HC1) and ON368702 (*Dd*TPR).

### Editing of yeast genes

Genetic manipulations were performed in yeast strains that were isogenic to *S. cerevisiae* S288c (for the list of strains, see Supplemental Table 2). Tagging of yeast genes with the yeast codon-optimized ORFs for yEGFP or mCherry, resulting in C-terminally tagged proteins, was by chromosomal integration of PCR amplified cassettes, as described earlier (Janke *et al*, 2004). Null mutants were generated by replacing whole genes with integration cassettes. In brief, the integration cassette, including a tag’s ORF or solely a stop codon followed by a selection marker under the control of the *Ashbya gossypii* TEF-promoter, was amplified with primers of up to about 100 nt in length, including 55–78 nt long overhangs complementary to the 5’ and 3’ region of the later to be targeted genomic location. The lithium acetate method (Gietz & Schiestl, 2007) was used to transform the WT or auxotrophic yeast cells with the purified PCR product, followed by colony selection via antibiotic resistance or on drop-out plates. Isolated strains were further analyzed by live-cell imaging, when purposeful, and by PCR, using genomic DNA, isolated as described (Harju *et al*, 2004).

### Yeast two-hybrid experiments

Haploid *S. cerevisiae* MATa strain CG-1945 or AH109 and MATα strain Y187 (Harper *et al*, 1993; Feilotter *et al*, 1994; James *et al*, 1996) were used for single transformations (Dohmen *et al*, 1991) with bait or prey constructs, respectively (all based on original vectors from Clontech Laboratories; for further plasmid details, see Supplemental Table 1). After selection on either −Trp (prey) or −Leu (bait) plates, single colonies were picked and grown as liquid culture. Mating was first done in solution in a 96-well plate with subsequent plating onto adenine-containing YPD (YPDA) agar plates. After two days of growth at 25°C, the cells were replica-plated onto agar plates with a selective medium (−Trp/−Leu; −LW) for the diploid cells to monitor the success of mating. After three days of growth at 25°C, cells were again replica-plated, now onto −Leu/−Trp/−His (−LWH) drop-out plates, without 3-amino-1,2,4-triazole (3-AT) and with increasing concentrations of 3-AT. These cells were further cultured at 25°C for up to 10 days, allowing the growth of those cells expressing protein pairs capable of interacting with each other. In addition, each bait construct was tested for self-activation by mating the corresponding MATa strain with a MATα strain transformed with the empty prey vector.

### Sequence database mining, protein structure predictions and molecular graphics tools

Details regarding the sequence database mining approaches are provided in Supplemental Information 1 and 2. In brief, the ScanProsite tool (de Castro *et al*, 2006; https://prosite.expasy.org/scanprosite) was commonly used for scanning the Swiss-Prot and TrEMBL protein sequence databases (Bairoch & Apweiler, 1997; https://www.uniprot.org/help/uniprotkb_sections), while the original BLAST tools (Altschul *et al*, 1990) were used for the mining of NCBI’s nucleotide and protein sequence databases. In addition, BLASTP was also used for reverse BLAST approaches to identify false positive sequences within whole genome sequencing (WGS) datasets, e.g., contaminating DNAs from a species’ food sources or fungal or other evident contaminations. Furthermore, searches were conducted with tools using position-specific score matrices (PSSMs) like position-specific iterated (PSI)-BLAST (Altschul *et al*, 1997), via pattern hit-initiated (PHI)-BLAST (Zhang *et al*, 1998), and via domain enhanced lookup time accelerated (DELTA)-BLAST (Boratyn *et al*, 2012). In addition, other profile-based tools like pHMMER (Finn *et al*, 2011; Potter *et al*, 2018), next to others making use of HMMs; see Supplemental Information 3), were used for other profile-based approaches. WebLogos (Crooks *et al*, 2004) were generated with an online tool (https://weblogo.berkeley.edu/), using for this the original Pfam MSAs from the Pfam-A full datasets of different releases (http://ftp.ebi.ac.uk/pub/databases/Pfam/releases/), retrieved from Pfam database’s FTP server (http://ftp.ebi.ac.uk/pub/databases/Pfam/). For generating HMM logos as vector graphics, Pfam seed sequences retrievable from the Pfam website (https://pfam.xfam.org/) were processed with the Skylign tool (Wheeler *et al*, 2014; http://skylign.org/). Predicted structures presented in the current study were either retrieved as PDB files from the structure database of AlphaFold (https://www.alphafold.ebi.ac.uk; Jumper et al, 2021; Tunyasuvunakool et al, 2021) or predicted using the platform ColabFold (AlphaFold2 using MMseqs2; https://colab.research.google.com/github/sokrypton/ColabFold/blob/main/Alpha Fold2.ipynb; Mirdita et al, 2021), using the latter also for structures not yet present in the AlphaFold database. While ColabFold also makes use of the source code of AlphaFold2, it uses a different tool for generating the input MSAs (multiple sequence alignments), a feature that we, in turn, exploited for assessing possible limitations in the BLD structure predictions (Supplemental Information 3). Most of the structures presented in the current study, thus stemming either from the AlphaFold database or obtained after a ColabFold-prediction process, as specified in the Figures, had then been processed further in the UCSF Chimera system (e.g., Pettersen *et al*, 2004; https://www.rbvi.ucsf.edu/chimera) for coloring, structural alignments, superimpositions, and *in silico* truncations, and then exported from Chimera as Figures. In particular, structural alignments, which in turn generated a new sequence alignment, were performed with the Match-Align tool of the Chimera system. For the superimposition of the different homologs’ BLD modules onto each other, we used Chimera’s superimposition tools, including MatchMaker. Chimera’s structural analysis tool was used to compute and illustrate potential contacts of designated atoms of an aa side chain with neighboring aa residues (distance ≤ 4 Å). The UCSF Chimera tools enabling molecular graphics and analyses had been developed, with support from NIH P41-GM103311, by the Resource for Biocomputing, Visualization, and Informatics at the University of California, San Francisco. Fast structural comparisons of protein structure datasets were performed on the Foldseek web server (https://search.foldseek.com/search; van Kempen et al, 2022), using the PDB files from the AlphaFold database for the human, amoebic and budding yeast ZC3HC1 homologs (Uniprot identifiers Q86WB0, Q54PS8, and Q03760) and the *Drosophila melanogaster* Diap2 protein (Q24307) as query structures. Foldseek searches for structures within the AlphaFold/Proteome v2 database were conducted in the 3Di/AA mode, using the taxonomic filter for the respective other species.

## Abbreviations

aa: amino acid
AI: artificial intelligence
BIR: baculovirus inhibitor of apoptosis protein repeat
BLD: BIR-like domain
FLIP: fluorescence loss in photobleaching
FP: fluorescent protein
IAP: inhibitor of apoptosis protein
ILP: IAP-like protein
IB: immunoblotting
IFM: immunofluorescence microscopy
KO: knockout
NB: nuclear basket
NE: nuclear envelope
NPC: nuclear pore complex
NuBaID: nuclear basket interaction domain
NUP: nucleoporin
TR: terminal ring
Y2H: yeast two hybrid
zf: zinc finger

## Acknowledgements

We gratefully acknowledge Frank Schwarz and Kerstin Mohr at the Core Facility for Genomics and Proteomics at the German Cancer Research Center (DKFZ) for contributions to Y2H experiments, and Gabriele Hawlitscheck at the Department for Cellular Logistics at the Max Planck Institute for Multidisciplinary Sciences for contributions to cloning. Furthermore, we thank Dirk Görlich for generous support, and Eberhard Bodenschatz, Steffen Frey, Dirk Görlich, Katharina Gunkel, Martin Kollmar, Arturo Vera Rodriguez, and Frank Schwarz for kindly providing research materials. In addition, we appreciate Thomas Güttler, Michael Ridders and Arturo Vera Rodriguez for technical advice.

## Author contributions

Conceptualization, V.C.C. and P.G.; data curation, P.G. and V.C.C.; formal analysis, P.G., V.C.C., H.I., and S.K.; investigation, P.G., V.C.C, H.I., and S.K.; project administration, V.C.C. and P.G.; supervision, V.C.C.; validation, P.G., V.C.C., H.I., and S.K.; visualization, P.G., V.C.C., and H.I.; writing—original draft preparation, V.C.C. and P.G.; writing—review and editing, V.C.C., P.G., and H.I. All authors have read and agreed to the submitted version of the manuscript.

