## Supplemental material for "Nuclear basket protein ZC3HC1 and its yeast homolog Pml39p feature an evolutionary conserved bimodular construction essential for initial binding to NPC-anchored homologs of scaffold protein TPR"

Running title: The bimodular homologs ZC3HC1 and Pml39p

#### Supplemental Information, Figures, Discussion, Tables, Sequences and References

#### Supplemental Information

##### Supplemental Information 1. Motif-based scanning of sequence databases for ZC3HC1 homologs.

Our searching for potential ZC3HC1 homologs also in other phyla and clades added to former studies relating to this issue (e.g., Higashi *et al*, 2005; Kokoszynska *et al*, 2008). In particular, we early on wanted to know whether possessing only one type of NuBaID signature-encoding gene per species, which we then knew for sure was the case in vertebrates, might also be common to species beyond the chordates. Or whether some organisms might make wider use of the NuBaID signature, and the type of construction it represents, by featuring it as part of very different proteins. While we used a wider range of search tools (see Material and Methods and Supplemental Information 2) in the later course of recurrently searching the sequence databases over the years, we made use of only two tools at the very beginning of our data mining for ZC3HC1 homologs across the eukaryotic realm. On the one hand, this was the omnipresent basic local alignment search tool BLAST (Altschul *et al*, 1990), which we used for the complementary mining of NCBI's nucleotide and protein sequence databases, thereby using BLASTP also for regularly conducting reverse BLAST searches to identify sequences falsely assigned to a given species (see Material and Methods). On the other hand, we used, initially most commonly, the ScanProsite tool (de Castro *et al*, 2006; https://prosite.expasy.org/scanprosite) for scanning the reviewed, manually annotated Swiss-Prot and the unreviewed, computationally annotated TrEMBL protein databases.

One of the early signature motifs we used for such a purpose was the minimalist BLD sequence signature G-W-$X_{(9,15)}$-C-$X_{(2)}$-C-$X_{(31,152)}$-H-$X_{(3)}$-C-X-W that we had assembled after having compared the vertebrate ZC3HC1 homologs and having found some of their cysteine residues dispensable (Figure 2A2). We arranged two of these signatures in tandem, thus then representing the two BLDs of *Hs*ZC3HC1 with their corequisite zinc fingers, once we had found the integrity of both of them essential for the protein's interaction with TPR. It turned out immediately evident, though, that this signature did not allow for detecting some possible homologs in other phyla, like, for example, the fission yeast protein Rsm1p (Yoon, 2004), whose sequence similarity with *Hs*ILP1/ZC3HC1 had already been noted earlier (Higashi *et al*, 2005; Finn *et al*, 2006; http://pfam.xfam.org/family/PF07967).

Therefore, knowing that the spacing between the second BLD's two pairs of suspected zinc-coordinating residues, i.e., between the C-$X_{(2)}$-C and H-$X_{(3)}$-C sequences, could be highly variable (Higashi *et al*, 2005; Kokoszynska *et al*, 2008), and also aware by then that the large insertion within the second BLD of *Hs*ZC3HC1 was dispensable for NE association, we made use of this knowledge for creating several additional versions of this novel motif assemblage. We thereby also took the conclusions into account that we had been able to draw from our single aa substitution experiments, namely that some positions of the signature for the BLDs of *Hs*ZC3HC1 did not tolerate certain substitutions, while others like W107Y, W107F, W256Y, and W256F had no notable effect on the NE association of the respective mutant versions of *Hs*ZC3HC1. Furthermore, having then attentively considered also the published information available until then (Higashi *et al*, 2005; Kokoszynska *et al*, 2008), one of our newly created second-generation signatures read G-[WYF]-$X_{(8,72)}$-C-$X_{(2)}$-C-$X_{(15,524)}$-H-$X_{(3)}$-C. With the latter signature arranged in tandem (Supplemental Figure S2D2) and with the linker's length between the two copies initially deduced from published information (Higashi *et al*, 2005; Kokoszynska *et al*, 2008), we could then readily identify not only *Sp*Rsm1p but also *Sc*Pml39p as a putative homolog in budding yeast, in which a ZC3HC1 homolog had remained undetectable until then.

With these and next-generation signatures notably differing in complexity in the course of our study, we nonetheless collectively designated all full-length versions of them as simplifying signatures of the NuBaID, of which we later rated some as representative of a prototypic ZC3HC1 homolog in most phyla. For assembling the later versions, we also used information then gained from experiments conducted in ZC3HC1 KO cells once the latter had become available. This approach allowed for identifying several residues whose substitution for other amino acids was tolerable, i.e., still allowed for NE-binding of the human homolog as long as such mutants did not need to outcompete a more binding-competent WT version of ZC3HC1 within the same cell. Among several residues, this held, e.g., for substitutions of W158 and W431 of *Hs*ZC3HC1 (e.g., Supplemental Figure S2B2). This latter information also turned out to be of value when later interpreting the NuBaID signatures of other groups of organisms, in which we found a whole range of residues to occur at positions corresponding to W158 and W431 of *Hs*ZC3HC1. Later, studying other facets of the ZC3HC1 protein's structure and function, we found some of these residues, too, allowing for NB and TPR association, as will be presented in another context elsewhere (our unpublished data).

All of these motif versions were used to scan the Swiss-Prot and TrEMBL protein databases via ScanProsite. Later, we then also complemented this approach and sometimes replaced it by searching NCBI-based sequences with the pattern-hit initiated BLAST (PHI-BLAST) program (Zhang *et al*, 1998). The latter uses as input not only a signature to search for pattern-conforming subject sequences but also a query sequence, in our case first only the one for *Hs*ZC3HC1 and later also the ones of clearly identified ZC3HC1 homologs, to subsequently construct local alignments next to the pattern's residues, between the query and the identified sequences. This hybrid strategy of PHI-BLAST allowed for sorting out more easily than with ScanProsite those sequences whose possession of relaxed NuBaID signatures of very low sequence stringency was regarded as random, namely when no additional traces of sequence similarity in the signature's vicinity indicated kinship.

The motif-based searches were constantly complemented by primary sequence alignment searches within the freely accessible nucleotide and protein sequence databases, using for this purpose the BLAST tools and, as query sequences, a selection of those from the steadily increasing collection of putative ZC3HC1 homologs that we were identifying in different taxa. We used such latter sequences for also screening expressed sequence tag (EST) and whole-genome shotgun (WGS) databases via TBLASTN. Thereby, we occasionally also ventured to re-interpret genomic information, particularly by newly predicting and assembling exon sequences and by re-defining open reading frame (ORF) boundaries in those cases in which we felt sure that computational ORF predictions and automatic annotations had not deciphered the corresponding gene correctly. In addition, to further validate or supplement sequences already deposited in the databases, we isolated mRNAs for cDNA synthesis and sequencing from some organisms of interest. Beyond that, when TBLASTN-searching the nucleotide sequence databases of protist phyla, we considered that some protists exhibit exceptions to the standard nuclear genetic code in eukaryotes (https://www.ncbi.nlm.nih.gov/Taxonomy/Utils/wprintgc.cgi).

Furthermore, the abovementioned approaches were later complemented by checking the identified sequences for additional signature elements conforming to either complete or partial versions of the Pfam motifs zf-C3HC and Rsm1. The latter was done even though the Rsm1 motif, in particular, often did not allow for identifying proteins we had been able to define by then as prototypic NuBaID-containing ones like, for example, *Sc*Pml39p and the *D. discoideum* ZC3HC1 homolog DDB0349234 presented in the current study. Both of these ZC3HC1 homologs still have not been assigned an Rsm1 motif, as defined by Pfam, to date (July 2022). Nonetheless, we further inspected those Pfam database-deposited sequences and species listed there as possessing a zf-C3HC or an Rsm1 motif to search for potential candidates that might have remained undetected by the other abovementioned local sequence alignment searches and the pattern-based ones using a NuBaID signature.

Altogether, the combination of complementary approaches allowed us to progressively comb through the eukaryotic realm in a reiterative and interactive manner. Such database mining eventually resulted in identifying numerous potential ZC3HC1 homologs in all eukaryotic supergroups, namely in the Opisthokonta, Amoebozoa, Excavata, Archaeplastida, and several lineages within the SAR supergroup. In addition, we could identify likely homologs in many other protist groups and genera whose affiliation was still uncertain (e.g., Adl *et al*, 2012; Pawlowski, 2013; Burki, 2014) at times when we intermittently conducted rounds of such signature-based data mining for ZC3HC1 homologs also in lower eukaryotes.

However, while we had also realized by then that certain organisms appear to lack a functional ZC3HC1 homolog, we were also aware that other species possessed a ZC3HC1 protein that had neither been detectable by the pattern-based nor the primary sequence alignment searches conducted till then. Subsequently, we defined further variants of the NuBaID signature, which, for example, also tolerated different spacing between the first two cysteines of the first BLD's zinc finger signature, then reading C-$X_{(3)}$-C instead of C-$X_{(2)}$-C (see also Supplemental Figure S3). Finally, for a re-scanning of the database-positioned sequences of those species for which we had neither been able to detect a ZC3HC1 homolog with any NuBaID signature version nor with the zf-C3HC or Rsm1 motifs of the Pfam database, we eventually also assembled low stringency NuBaID signatures that incorporated characteristic sequence features of the BIR domains, described further below (see also Supplemental Figure S10F). While these still did not allow for detecting a likely ZC3HC1 homolog in some species, like, for example, in *Drosophila* (our unpublished data), we momentarily cannot exclude for sure that there might also exist ZC3HC1-homologous proteins evolutionarily altered beyond recognition, or analogous proteins of equivalent function, at the NBs of such species.

**Supplemental Information 2. Low overall sequence similarity and a lack of shared, database-deposited sequence motifs as an explanation for a kinship so far gone unrecognized between distinct ZC3HC1 homologs.**

Unlike when scanning fungal sequences via ScanProsite with the NuBaID signatures, we could not detect *Sc*Pml39p when conducting local alignment searches via standard protein-protein BLASTP (Altschul *et al*, 1990) when starting with *Hs*ZC3HC1 as the query sequence, and neither was this possible vice versa. Furthermore, finding the other species' homolog was also not possible with tools using position-specific score matrices (PSSMs), like position-specific iterated (PSI)-BLAST (Altschul *et al*, 1997), when we had been searching the genus *Saccharomyces* with *Hs*ZC3HC1 as the query, and neither was this possible, again, vice versa when searching for the vertebrate homologs with *Sc*Pml39p. Similarly, the homology search tool MMseqs2 (Steinegger & Söding, 2017), as used for ColabFold-based protein structure predictions (Mirdita *et al*, 2021, 2022), did not detect the human or yeast homolog with the respective other homolog's sequence either. Furthermore, even when using as the input either the human or the yeast sequence together with one of the abovementioned NuBaID signatures for then conducting searches via pattern hit-initiated (PHI)-BLAST (Zhang *et al*, 1998; https://blast.ncbi.nlm.nih.gov/Blast.cgi?PAGE=Proteins), the other homolog was not detected. In addition, other profile-based approaches, including tools like JackHMMER (Johnson *et al*, 2010) or pHMMER (Finn *et al*, 2011; Potter *et al*, 2018; https://www.ebi.ac.uk/Tools/hmmer/) that make use of Hidden Markov Models (HMM) built from multiple sequence alignments (MSA), did not allow for the identification of *Sc*Pml39p when searching the genus *Saccharomyces* with default settings and *Hs*ZC3HC1 as the query. Again, neither were the mammalian homologs identified using *Sc*Pml39p for searches via the HMMER tools. Furthermore, one did also not detect the human homolog when using the tool domain enhanced lookup time accelerated (DELTA)-BLAST (Boratyn *et al*, 2012) for a search starting with *Sc*Pml39p. However, DELTA-BLAST allowed for detecting *Sc*Pml39p when starting the search with *Hs*ZC3HC1.

These latter results can be briefly explained as follows. DELTA-BLAST makes use of the signatures and HMMs present in the Conserved Domain Database (CDD; https://www.ncbi.nlm.nih.gov/Structure/cdd/cdd.shtml) for constructing an MSA for those proteins to which such motifs have been attributed. Such MSAs are then the prerequisite for computing PSSMs eventually used for searching the sequence databases. In other words, DELTA-BLAST searches the CDD with a query sequence and then uses the domains the query gets aligned with to create a PSSM. This approach thus differs from other search tools commonly used for detecting distantly related homologs, with some among the latter using, for example, an MSA for constructing an HMM then used for database searching or, as another example, with some creating a PSSM based on an MSA that derives from a regular BLASTP search.

Regarding *Hs*ZC3HC1, the Pfam zf-C3HC and Rsm1 motifs were assigned to it long ago, with this information then also deposited in the CDD. The latter thus allows for its alignment with sequences possessing the same motifs, for deriving a PSSM from such an MSA, and for then using the latter for sequence database searches. However, concerning *Sc*Pml39p, the Pfam database has not attributed an Rsm1 motif to it to date (July 2022; http://ftp.ebi.ac.uk/pub/databases/Pfam/releases/Pfam35.0/). Moreover, while a zf-C3HC motif has been assigned to *Sc*Pml39p recently (http://ftp.ebi.ac.uk/pub/databases/Pfam/releases/Pfam30.0/), this information has not yet been incorporated into the CDD. The latter, in turn, means that there is no CDD-deposited profile for *Sc*Pml39p that would allow its alignment with its homologs possessing such Pfam motifs, which means that with *Sc*Pml39p as the query sequence, DELTA-BLAST will not be able to use a PSSM for its sequent search. In other words, since DELTA-BLAST *"owes its generally very good performance regarding search sensitivity and quality of alignment to the information available in the CDD"* (Boratyn *et al*, 2012), a situation in which it is not possible to attribute any of this information to a query sequence of interest, results in DELTA-BLAST conducting merely a BLASTP search with this sequence. The latter thus happens to be the case for *Sc*Pml39p, for which we already knew that a BLASTP search does not allow for detecting the human homolog.

**Supplemental Information 3. Assessment of the input datasets' respective contributions to user-initiated BLD structure predictions by AlphaFold2.**

After having inspected the predicted structures of *Hs*ZC3HC1 and *Sc*Pml39p available in the AlphaFold database for the first time and also having compared the human homolog's BLDs with both the crystal and AlphaFold2-predicted structures of the BIR domains (see, e.g., Supplemental Figure S10), we had considered it justified to look at these domains' predicted similarities with some caution, both despite and because of the human and yeast ZC3HC1 homologs' overall sequence dissimilarities on the one hand and the BLDs' and BIR domains' profile HMM similarities on the other.

In brief, we were aware that AlphaFold2 uses the primary amino acid sequence, i.e., the query sequence, for first searching both protein sequence and protein structure databases, then converts these search results into distinct input datasets, and then uses the latter for its further computations (Jumper *et al*, 2021). On the one hand, the searching of the sequence databases would result in the construction of an MSA composed of sequences from evolutionarily related proteins, with this process involving tools like Jackhmmer (Johnson *et al*, 2010; Eddy, 2011; https://www.ebi.ac.uk/Tools/hmmer/search/jackhmmer) and HHblits (Remmert *et al*, 2011; https://toolkit.tuebingen.mpg.de/tools/hhpred). On the other hand, AlphaFold2 would search the Protein Data Bank (PDB)-deposited crystal structures for structures it regards as potentially similar to the one the query sequence would adopt. In fact, the second of AlphaFold2's input datasets are constructions that it calls the "pair representations", with these the outcome of having aligned PDB structure templates and query sequence for computing some initial representations of the query's structure (Jumper *et al*, 2021). In the subsequent computation steps, AlphaFold2 would then refine the MSA and the pair interactions, thereby exchanging information between the MSA and the structure templates iteratively. Finally, AlphaFold2 would use the exhaustively refined MSA and pair representation to construct a threedimensional structure model (Jumper *et al*, 2021; see also https://www.blopig.com/blog/ 2021/ 07/alphafold-2-is-here-whats-behind-the-structure-prediction-miracle/).

Having then noted that for constructing the abovementioned pair representations, a ZC3HC1 query sequence would be assigned to the PDB-deposited BIR domain crystal structures as templates for AlphaFold2's computations for a BLD structure, we had wondered to which extent an alignment of a BLD with a given BIR structure, channeling the prediction into one direction, might introduce a discussible level of bias into the prediction. In fact, since AlphaFold2 was known to have been trained to produce a prediction that would be the one "*most likely to appear as part of a PDB structure*" (Jumper *et al*, 2021), we wondered how informative it would be when a structure predicted for a query sequence would look very similar to already available crystal structures considered related. In other words, with the profile HMMs of query sequences being used for database searches via HMM comparisons (Jumper *et al*, 2021), using tools like, e.g., HHSearch and HHblits (Söding, 2005; Remmert *et al*, 2011; Zimmermann *et al*, 2018) and Jackhmmer (Johnson *et al*, 2010), we wondered how such a correlation of similar but not identical HMMs with one type of crystal structure would affect the outcome of a structure prediction. In particular, since HHSearch was apparently identifying Pfam's BIR profile with a *Hs*ZC3HC1 query sequence via the HMM of its zf-C3HC motif, this would result in assigning ZC3HC1 homologs with such a profile HMM to the numerous BIR crystal structures already deposited in the PDB.

Beyond that, we had noticed that AlphaFold2 had been mentioned to take ligands and ions into account when these *"are predictable from the sequence alone"*, with AlphaFold2 *"likely to produce a structure that respects those constraints implicitly"* (Jumper *et al*, 2021), yet then found this not appear to be so for the BLDs' likely zinc ion coordination spheres (see further below, and Supplemental Figure S7D). Such latter findings, though, were in line with statements elsewhere, according to which AlphaFold2 does not make predictions about any non-protein components that might be part of a protein of interest (https://www.embl.org/news/ science/alphafold-potential-impacts/). The latter, in turn, meant that AlphaFold2 would also not take into account any potential role that the ZC3HC1 homologs' zinc ions could execute in the protein's folding process. Since it is known, though, that zinc ions can play a crucial role in the proper folding of zinc proteins, in addition to stabilizing a resulting fold (Maret & Li, 2009; Gomes & Wittung-Stafshede, 2010; Padjasek *et al*, 2020), and with this also imaginable for ZC3HC1, we wondered whether non-consideration of the zinc ions' contributions by AlphaFold2 would demand caution when interpreting the predicted structures.

Further along this line, we had realized in an initial series of trial predictions that AlphaFold2 does not appear to predict the structural impact that substitutions of NuBaID's signature residues should have, which both held for mutations naturally occurring *in vivo* and others experimentally created in the current study. On the one hand, the AlphaFold2-predicted structures of such BLD single aa substitution mutants appeared essentially indistinguishable from the wild-type protein's predicted structure. On the other hand, we had found such mutations abolishing the protein's ability to bind specifically to the NB and TPR. Moreover, additional experimental evidence suggested that at least some of the aa-substituted versions were no longer correctly folded *in vivo*. In contrast to the wild-type protein, such mutants were often more rapidly degraded and more readily sedimented by centrifugation, next to non-specifically interacting with other proteins (e.g., Gunkel & Cordes, 2022, and our unpublished data), indicating that such mutations had indeed disrupted the BLDs' natural structure. At some point then, we found it mentioned that *"AlphaFold has not been trained or validated for predicting the effect of mutations"* and was "*not expected to capture the effect of point mutations that destabilise a protein"* (https://www.embl.org/news/science/alphafold-potential-impacts/). It was also inferable why this might be so since AlphaFold2 had been trained to correlate sequence information with only the end products of protein folding processes (Jumper *et al*, 2021) and with such end products having been primarily the proteins' wild-type versions or parts thereof. Nonetheless, such realization meant that we had to be cautious also in this context and to avoid pitfalls when interpreting the predicted structures of, for example, distinct ZC3HC1 homologs that represent some particular, naturally existing mutant versions.

Given such preliminary insight, yet lacking the expertise to comment on the program's algorithms and codes, while at the same time wanting to better assess the range of AlphaFold2's opportunities and restraints in order to define how far we could go in interpreting predicted BLD structures, we conducted, in a semi-systematic manner, some additional series of simple trial predictions. With some, we wanted to assess the degree of interdependency between the MSA-based and the template-based part of AlphaFold2's prediction process and how the PDB-deposited structures would contribute to a user-initiated structure prediction. In other words, we wanted to know how the exclusion of PDB information, i.e., not permitting the program to use the PDB-deposited BIR domain crystal structures as reference templates, might affect the predicted BLD structures' appearance (e.g., Supplemental Figure S7A and S7B). Furthermore, since we also intended to use the ColabFold platform (https://colab.research.google.com/github/sokrypton/ColabFold/blob/main/AlphaFold2.ipynb) for conducting such trial predictions, to benefit from its accelerated predictions made possible by combining the fast homology search of the MMseqs2 program with AlphaFold2 (Mirdita *et al*, 2021), instead of conducting all structure predictions directly via AlphaFold2, we considered it necessary to compare the AlphaFold2 predictions with those obtained via the ColabFold-accelerated approach (Supplemental Figure S7A and S7B).

Then, with other trials, we tested the outcome of using MSAs only composed of sequences lacking Pfam Rsm1 motifs while representing ZC3HC1 homologs nonetheless, according to our criteria. In other words, we wanted to know how far sequences lacking such a profile HMM, which might also not have been part of the MSAs used for the PDB template alignments during AlphaFold2's initial training sessions, would allow for user-initiated BLD structure predictions resembling those that one obtains with the programs' default settings (e.g., Supplemental Figure S7C). In again other words, with approaches of this kind, we aimed to assess how the MSA's composition and how selectively changing it would affect the outcome of a prediction process.

Finally, we wanted to learn more about AlphaFold2's inability to predict the effects caused by mutations. So far, we knew that the sequence of an individual aa substitution mutant, used as the query, would lead to an MSA primarily composed of wild-type sequences. Since this wild-type-sequence-dominated MSA would then be the input dataset guiding the subsequent structure prediction process, we wondered whether it might be the dominance of wild-type sequences, "assimilating" the mutant sequence, that would prevent appreciating a single mutation's structural consequences. Now, we wanted to know whether the outcome would be different if such an MSA were composed exclusively of sequences similarly mutated. In other words, using the approach via ColabFold (Mirdita *et al*, 2021), we wanted to feed AlphaFold2's "evoformer" (Jumper *et al*, 2021) with an MSA that we would force into being composed merely of mutant sequences all harboring the same mutations (Supplemental Figure S7D). With this approach, we aimed to assess whether or not AlphaFold2's neural network training might have already educated it so that even compilations of mutations within an MSA might no longer be recognizable as such.

In brief, the outcome of these trial experiments can be summarized as follows. First, using ColabFold's default settings, we noted that the BLD and BIR domain structures predicted via ColabFold were essentially indistinguishable from those derived directly from AlphaFold2's database of predicted structures (e.g., Supplemental Figures S7A, S7B, and S10A2).

Second, we noted that user-initiated predictions of BLD and BIR domain structures, when not accessing PDB template information, resulted, nonetheless, in structures that too were essentially indistinguishable from those derived directly from AlphaFold2's database, as long as sufficiently informative MSAs were provided (e.g., Supplemental Figures S7A, S7B, and

S10A2). In other words, once AlphaFold2's initial neural network training had been completed, crystal structures like, e.g., those of the BIR domains were apparently dispensable as templates for a then user-initiated prediction of structures like, e.g., those of the BLDs. For such template-free modeling, we sometimes used the AlphaFold-Colab notebook (https://colab.research.google.com/github/deepmind/alphafold/blob/main/notebooks/AlphaFold.ipynb), the latter a simplified version of AlphaFold2 omitting in its predefined settings an alignment with PDB templates. However, for systematically conducting trials of this kind, we used ColabFold instead. For example, having noted that those MSAs compiled by ColabFold's default settings, e.g., for *Hs*ZC3HC1 and *Sc*Pml39p, would be linked, e.g., via the HHsearch tool, to the PDB-deposited metazoan BIR structures, we made use of ColabFold's options for barring or including such PDB information. Nonetheless, the template-free approach also led to predicting essentially the same BLD structures as with AlphaFold2's default settings (e.g., Supplemental Figure S7A and S7B).

Third, we found that a BLD2 structure prediction that had started with an MSA only composed of sequences without a pre-existing Rsm1 profile, while representing ZC3HC1 homologs nonetheless, could result in structures closely resembling those obtained with the template-based and template-barring settings of ColabFold and the default settings of AlphaFold2. For example, even without any template information and no profile HMM appertaining to the input sequences representing the homologs' BLD2, the latter at least so at the time when we initially conducted such trials before, e.g., the HHpred web server updates in 2022 (Zimmermann *et al*, 2018; https://toolkit.tuebingen.mpg.de/tools/hhpred), the predicted structure of the *Sc*Pml39p BLD2 still closely resembled the initial predictions based on MSAs composed of sequences with Rsm1 motifs already assigned to them (Supplemental Figure S7C).

Fourth, we found that AlphaFold2 did not predict the effect of amino acid substitution mutations even when several of such mutations had been introduced into all sequences of an MSA then to be the input dataset for a structure prediction. For example, having introduced several mutations into all of the MSA's sequences used for predicting the BLD structures of *Sc*Pml39p, even a high sensitivity HMM comparison tool like HHpred, again at the time prior to its 2022 web server updates, could no longer correlate an HMM deduced from such an MSA with any BLD-reminiscent structure. Nonetheless, the structure predicted for such a manifoldly mutated BLD2 was still essentially the same as obtained with ColabFold's default settings (Supplemental Figure S7D).

As a whole, these trial experiments provided some guiding principles for us as to how far we could go in carefully interpreting the predicted structures presented in the current study, with these trials' findings having outlined the confines within which we considered it justified to deduce conclusions cautiously from the data provided by AlphaFold2. Aware that the predictions, even for the BLDs' core parts, are likely not perfect, not expecting them to provide already the definite positions of the BLDs' structural elements, we regard further efforts that aim at still gaining BLD crystal structure information as reasonable. On the other hand, though, we regard the information provided by these predictions as already highly valuable within the range of resolution and presumed accuracy we considered sufficient for the current study's objectives. One of the latter was to provide a first overall impression of the ZC3HC1 homologs' molecular features, including the approximate positions of their BLDs' different structural elements relative to each other. Furthermore, such predictions provided hints as to where to expect the ZC3HC1 homologs' potential binding sites for their respective TPR homologs, allowing us to conjecture which additional structural parts of ZC3HC1 might be promising for further molecular manipulations. Finally, by illustrating the similarity and equivalence of their structural elements, these predictions markedly underscored the kinship of *Hs*ZC3HC1 and *Sc*Pml39p.

**Supplemental Information 4. A relationship between *Sc*Pml39p/*Hs*ZC3HC1 and *Sc*Nup157p/*Hs*NUP155?**

Pondering on similarities between *Hs*ZC3HC1 and *Sc*Pml39p, we consider it justified to bring a former Y2H screen's reported outcome to mind again. This screen of a yeast genomic library identified Nup157p as another potential binding partner of Pml39p (Palancade *et al*, 2005). With *Sc*Nup157p being the homolog of *Hs*NUP155 (Aitchison *et al*, 1995), this result might perhaps represent another one pointing towards kinship between Pml39p and ZC3HC1, even though our Y2H-screening of a human cDNA library with ZC3HC1 (Gunkel *et al*, manuscript in preparation) had not provided a NUP155 cDNA. We had also not found NUP155 notably co-detached with ZC3HC1 from NEs upon NB disassembly in *Xenopus* oocytes (Gunkel *et al*, 2021), and only minor amounts of NUP155, or none at all, were released concomitantly to the degradation of NBs in human cell lines expressing degron-tagged NB components (Gunkel & Cordes, 2022; our unpublished data). Remarkably, however, when immunoprecipitating ectopically expressed *Hs*ZC3HC1, we found peptides of NUP155 as the only representative of the NPC proper among the co-sedimented materials (our unpublished data).

Our subsequent attempts to illuminate this potential link between ZC3HC1 and NUP155 *in vivo* did not provide much further insight, with both proteins' subcellular positioning not appearing directly affected in the absence of the respective other. This latter finding, though, was again in line with *NUP157* deletion not having affected Pml39p localization either (Palancade *et al*, 2005). Nonetheless, we continued wondering whether the observed interactions between *Sc*Pml39p and *Sc*Nup157p and between *Hs*ZC3HC1 and *Hs*NUP155 might reflect some genuine relationship common to both species or just coincidentally some same type of unspecific interaction.

**Supplemental Information 5. Instability of the Mlp-dependent association of *Sc*Pml39p with NPCs under common cell fractionation conditions?**

The TPR-dependent association of ZC3HC1, both in humans and amphibians, is sensitive to low temperatures and commonly depends on the presence of bivalent cations, among other conditions, as described earlier (Gunkel *et al*, 2021). By contrast, some of the standard cell fractionation protocols, using ice-cold solutions without adequate amounts of bivalent cations, can cause all or substantial amounts of ZC3HC1 to be detached from the NE, together with the ZC3HC1-dependent population of TPR polypeptides (Gunkel *et al*, 2021). With some of these conditions also employed at specific steps of certain yeast fractionation procedures, we can imagine that sensitivity towards non-physiological conditions might also apply to the interactions between the Pml39 and Mlp polypeptides, causing Pml39p detachment from its Mlp binding partners.

Concerning Pml39p, a notion of a similar kind had already been expressed earlier when authors stated that the Mlp-dependent association of Pml39p with NPCs *"might thus be either transient or unstable under biochemical purification conditions"* (Palancade *et al*, 2005). Along this line, except for two studies (Niepel *et al*, 2013; Bensidoun *et al*, 2021), affinity purifications of Mlp1p or Mlp2p in other investigations appear not to have come along with identifying Pml39p in notable amounts, despite co-isolating other NPC proteins and even nucleocytoplasmic transport factors and mRNA-associated proteins (e.g., Niepel *et al*, 2005; Bretes *et al*, 2014; Saroufim *et al*, 2015; Kim *et al*, 2018). Moreover, just like ZC3HC1 had no longer been detectable among the proteins of purified rat NPCs (Cronshaw *et al*, 2002), reproducibly a consequence of its sensitivity to the conditions mentioned above (Gunkel *et al*, 2021), Pml39p had not been identifiable during the pathbreaking mass spectrometry-based proteomics of purified yeast NPCs either (Rout *et al*, 2000), with the corresponding *YML107C*/*PML39* ORF sequence possibly available at the time (Bowman *et al*, 1997). However, just as discussed for ZC3HC1 in detail (Gunkel *et al*, 2021), we see no incompatibility in Pml39p being an NB component whose interaction stability with the Mlp proteins can vary under certain non-physiological conditions.

#### Supplemental Figures

**Supplemental Figure S1.** Photobleaching experiments with fluorescent protein-tagged proteins ectopically expressed in HeLa WT cells.

**Supplemental Figure S2.** Characterization of ZC3HC1 deletion and single aa-substituted mutants, ectopically expressed in HeLa WT and ZC3HC1 KO cells, by fluorescence microscopy and FLIP, complemented by additional Y2H experiments.

**Supplemental Figure S3.** Functional tolerance of *Hs*ZC3HC1 for a C-$X_{(3)}$-C spacing within its first BLD, as commonly present in some fungal ZC3HC1 homologs.

**Supplemental Figure S4.** Characterizing the *Dictyostelium* homologs of TPR and ZC3HC1.

**Supplemental Figure S5.** Experiments complementing the characterization of *Sc*Pml39p and its NuBaID.

**Supplemental Figure S6.** Complementing studies on the contribution of *Sc*Pml39p in keeping subpopulations of NE-associated Mlp1p polypeptides positioned at the NB and within Mlp1p-containing nuclear foci.

**Supplemental Figure S7.** Assessment of the contributions of sequence database search-derived MSAs and PDB database-deposited structures as templates for BLD structure predictions.

**Supplemental Figure S8.** Former considerations regarding the zinc ion-coordination topology of ZC3HC1.

**Supplemental Figure S9.** The tertiary structures of *Hs*ZC3HC1, *Dd*ZC3HC1, and *Sc*Pml39p *in toto*, as predicted by AlphaFold2, and closer looks at a BLD1:BLD2 interface and at distinct aromatic amino acids flanking the zinc ion coordination spheres.

**Supplemental Figure S10.** Comparing the predicted structural characteristics and sequence features of the BLDs of ZC3HC1 with the BIR domains' structures and sequences.

**Supplemental Figure S11.** *In vivo* and *in silico* deletion mutants of ZC3HC1 homologs and their tertiary structures predicted by AlphaFold2 via ColabFold.

**Supplemental Figure S12.** ZC3HC1 deficiency in human cell lines not affecting the cellular amounts and subcellular distribution of FANCD2, in line with no evident robust interaction between ZC3HC1 and FANCD2 at the NE or in cell extracts.

**Supplemental Figure S13.** Size variations of loop-like sequence insertions within exemplary ZC3HC1 homologs, and the BLD2 loop of *Hs*ZC3HC1 as a prime target for phosphorylation.

**Supplemental Figure S14.** Searching AlphaFold2's protein structure datasets via Foldseek, using ZC3HC1 and BIR protein structures as queries.

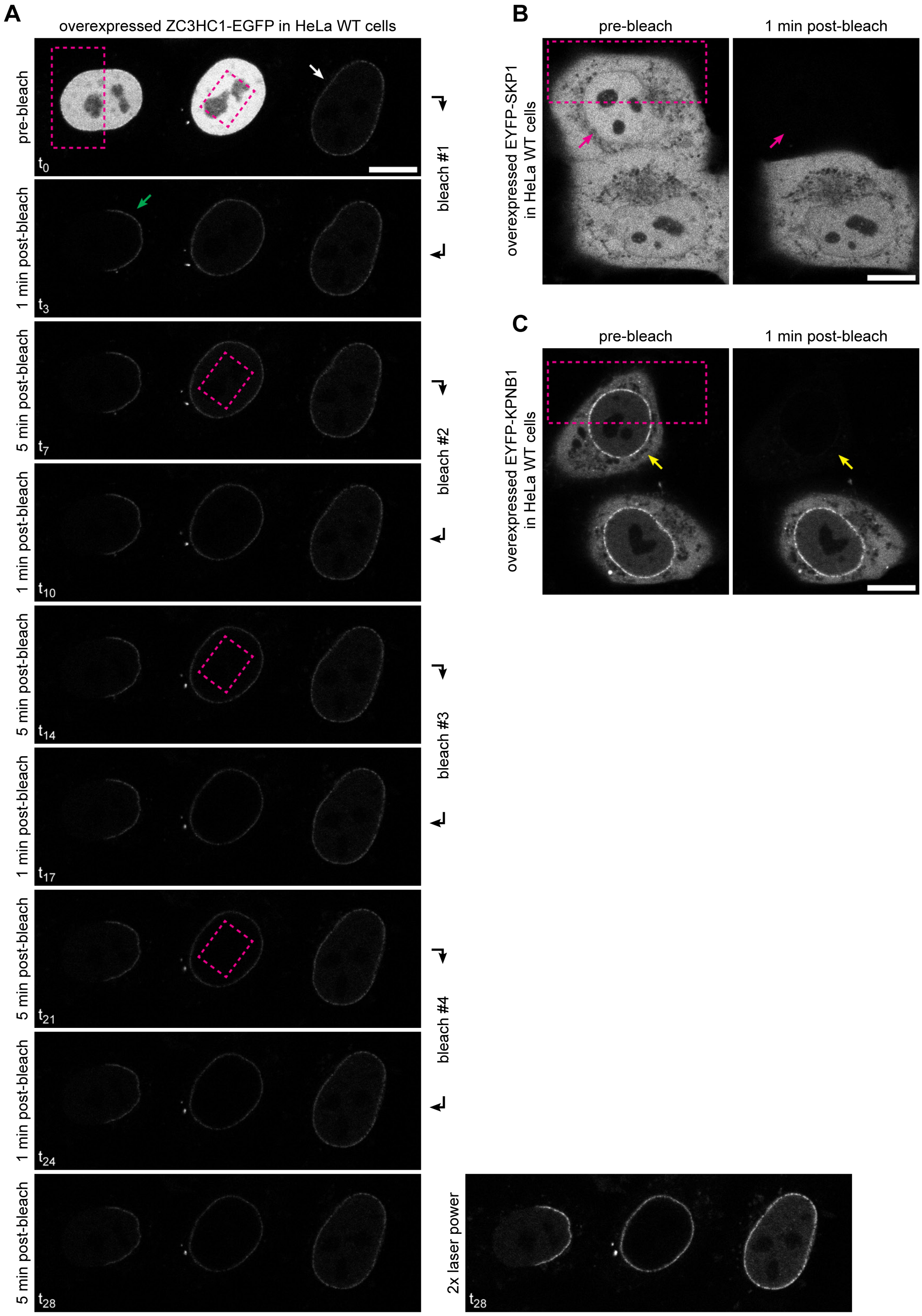

**Supplemental Figure S1. Photobleaching experiments with fluorescent protein-tagged proteins ectopically expressed in HeLa WT cells.**

Upon constitutive ectopic expression of either an N-terminally EYFP-tagged or a C-terminally EGFP-tagged version of the WT ZC3HC1 protein, we had found such recombinant proteins, when synthesized only in low amounts, primarily located at the NE, which suggested early on that the NE was a preferred binding site for these tagged versions of ZC3HC1. However, upon further overexpression, we had seen the tagged ZC3HC1 polypeptides distributed throughout the nuclear interior, then resembling the formerly described subcellular localization of ZC3HC1 (Ouyang *et al*, 2003). Such increased levels of ZC3HC1 often made it no longer possible to detect any preferred location at either the NE or any other site within the nucleus, and later we found this issue particularly applying to various mutant versions of ZC3HC1. Live-cell photobleaching of such nuclear pools of tagged ZC3HC1 polypeptides was thus mainly performed to more reliably assess whether specific ZC3HC1 mutants might be capable of binding to the NE. Later on, once ZC3HC1 KO cells had become available, such fluorescence-loss-in-photobleaching (FLIP) experiments were also conducted with the KO cells, with the latter then too ectopically expressing versions of ZC3HC1 tagged with fluorescent proteins (FP), as will be shown further below (see Supplemental Figure S2).

On the other hand, not yet aware at this point that one would prove ZC3HC1 to be a structural element of the NB years later (Gunkel & Cordes, 2022), the purpose of the initial photobleaching experiments shown here, performed with the EGFP-tagged WT version of ZC3HC1, was to allow for an early assessment as to how durably the ectopically expressed ZC3HC1 polypeptides might be positioned at the NE. Such experiments were thus to provide a first impression regarding the degree of exchange between the NE-located and nuclear pools of ZC3HC1 before addressing this issue at a later time point also in more detail by fluorescence recovery after photobleaching (FRAP) experiments, which will be presented as part of another study (our unpublished data).

Moreover, the initial photobleaching experiments shown here were to be conducted for comparison also with cells expressing other FP-tagged proteins, including the nuclear import factor importin β/KPNB1, which shuttles between the nucleus and cytoplasm (Görlich & Kutay, 1999) and SKP1, being a formerly reported direct binding partner of ZC3HC1 (e.g., Bassermann *et al*, 2005a, 2005b, 2007; Klitzing *et al*, 2011), with this latter notion, though, later refuted (Gunkel *et al*, 2021).

**(A)** Live-cell fluorescence micrograph of HeLa WT cells transiently transfected with constitutive expression vectors coding for ZC3HC1-EGFP, showing NE-staining in cells expressing only lower amounts of ZC3HC1-EGFP (white arrow) and others in which seemingly vast amounts were located throughout the nucleus. Photobleaching experiments were performed to assess the extent to which such intranuclear pools of ZC3HC1 might be mobile and to which extent there might be an exchange between this nuclear pool and the ZC3HC1 polypeptides at the NE. The two marked rectangular areas in the pre-bleach image were subjected to pulses of full laser power for 2 min, followed by the acquisition of post-bleach images 1 min and 5 min after the bleaching. The cell shown in the center was then subjected to an immediate next round of bleaching for 2 min, again followed by image acquisition and the repetition of this procedure until four rounds of bleaching and image acquisitions had been conducted. For comparison, the cell expressing only low amounts of ZC3HC1-EGFP in the same field of imaging (white arrow) was left unbleached, also in order to assess the degree of signal reduction due to bleaching during image acquisition. Note that the surplus of ZC3HC1 polypeptides deeper within the nuclear interior of the centrally located cell apparently had no immobile natural binding partner that would allow for a longer-lasting interaction. In fact, repeated bleaching of the nuclear interior of this ZC3HC1-overexpressing cell resulted in quantitative elimination of the nuclear EGFP fluorescence, pointing at a highly mobile pool of nuclear ZC3HC1 polypeptides. On the other hand, the NE of this bleached middle cell concomitantly emerged as the only structure that was still fluorescent even after having bleached its nuclear interior repeatedly. This result indicated a steady and lasting *in vivo* interaction between the NE and a certain amount of ZC3HC1. This conclusion was underscored further by the result relating to the left cell's bleached NE fragment (green arrow), which had remained faded over the monitored period of about 32 min, without any remarkable recovery of EGFP fluorescence. This result of such a first type of preliminary FRAP experiment still held when the corresponding image had been acquired with doubled laser power (last image on the right side). As an aside, note that we obtained essentially identical results also after having performed such photobleaching experiments with HeLa cells expressing EYFP-ZC3HC1 (data not shown). Bar, 10 µm.

**(B**, **C)** Live-cell fluorescence micrographs of HeLa WT cells that had been transiently transfected with constitutive expression vectors coding for EYFP-SKP1 and EYFP-KPNB1 and then used for representative photobleaching experiments. In both S1B and S1C, each bleached area (large rectangles) had been subjected once to pulses of full laser power for 2 min. By contrast, the reference cells, one each in S1B and S1C, had been left unbleached to show that the degree of bleaching due to image acquisition prior to and after the actual bleaching procedure was negligible. Especially when compared to the NE-associated pool of ZC3HC1-EGFP in S1A, it was evident that all subcellular populations of these other proteins were far more mobile, yet to a variable extent.

**(B)** In the case of the EYFP-SKP1-expressing cell, the single round of bleaching of part of its nucleus and cytoplasmic compartment led to a complete loss of this cell's EYFP fluorescence. Note, in particular, that also no traces of signal were seen at the non-bleached NE (magenta-colored arrows), with this suggesting early on that this particular protein does not engage in any lasting interaction with any of the NE's components. As an aside, also note that we obtained essentially identical results after performing such photobleaching experiments with HeLa cells expressing SKP1-EGFP (data not shown). Bar, 10 µm.

**(C)** In the case of the EYFP-KPNB1-expressing cells, and in contrast to those expressing EYFP-SKP1, the overexpressed protein was not only distributed throughout the nucleus and cytoplasm but also enriched at the NE. However, this did not come as a surprise as such enrichment was known to reflect the naturally occurring subcellular distribution of endogenous KPNB1 (Görlich *et al*, 1995). Nonetheless, just like in the case for EYFP-SKP1, a single round of only bleaching part of the nucleus and the cytoplasmic compartment was sufficient to obliterate all fluorescence from both compartments, in line with KPNB1 being primarily a highly mobile nuclear transport receptor. Furthermore, and in striking contrast to the findings for ZC3HC1-EGFP in S1A, the EYFP-KPNB1 signal at the unbleached part of the NE too (yellow-colored arrows) was nearly abolished, confirming that KPNB1 polypeptides mainly engage in only transient physical interactions with components of the NPC. Bar, 10 µm.

## A1

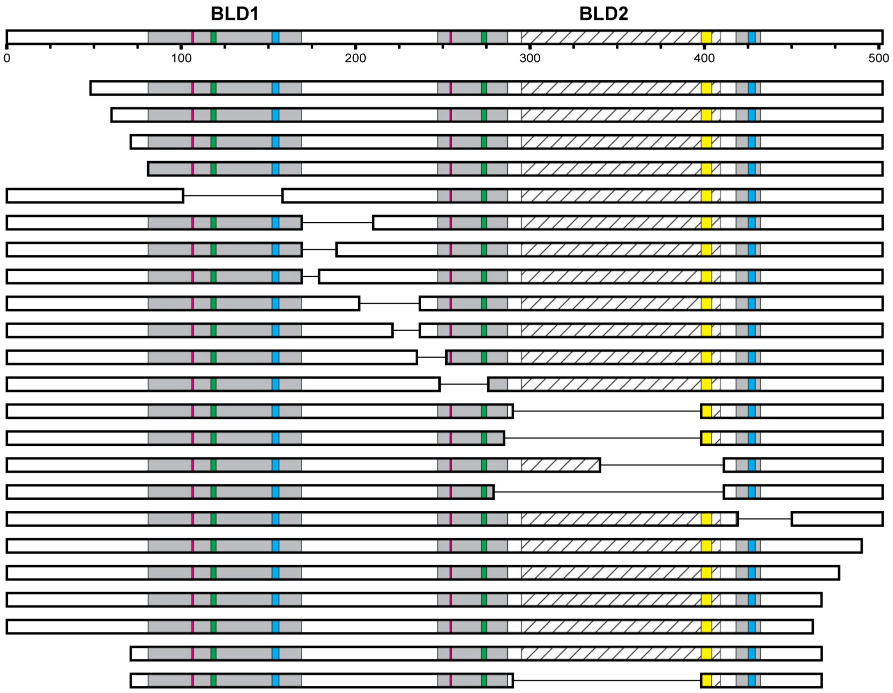

| Construct | binding to the NE: low exp. WT | low exp. KO | FLIP WT | FLIP KO |
|---|---|---|---|---|
| 1–502 | + | + | + | + |
| 49–502 | + | + | + | + |
| 61–502 | + | + | + | + |
| 72–502 | + | + | + | + |
| 82–502 | − | − | − | − |
| 1–101_159–502 | − | − | − | − |
| 1–169_211–502 | − | − | − | − |
| 1–169_189–502 | − | − | − | − |
| 1–169_179–502 | − | − | − | − |
| 1–202_237–502 | − | − | − | − |
| 1–221_237–502 | + | + | + | + |
| 1–235_252–502 | − | − | − | − |
| 1–248_276–502 | − | − | − | − |
| 1–290_398–502 | + | + | + | + |
| 1–285_398–502 | ~ | + | + | + |
| 1–340_412–502 | + | + | + | + |
| 1–279_412–502 | − | − | + | + |
| 1–419_450–502 | − | − | − | − |
| 1–490 | + | + | + | + |
| 1–477 | + | + | + | + |
| 1–467 | + | + | + | + |
| 1–462 | − | − | − | − |
| 72–467 | + | + | + | + |
| 72–290_398–467 | + | + | + | + |

**S2 (1/8)**

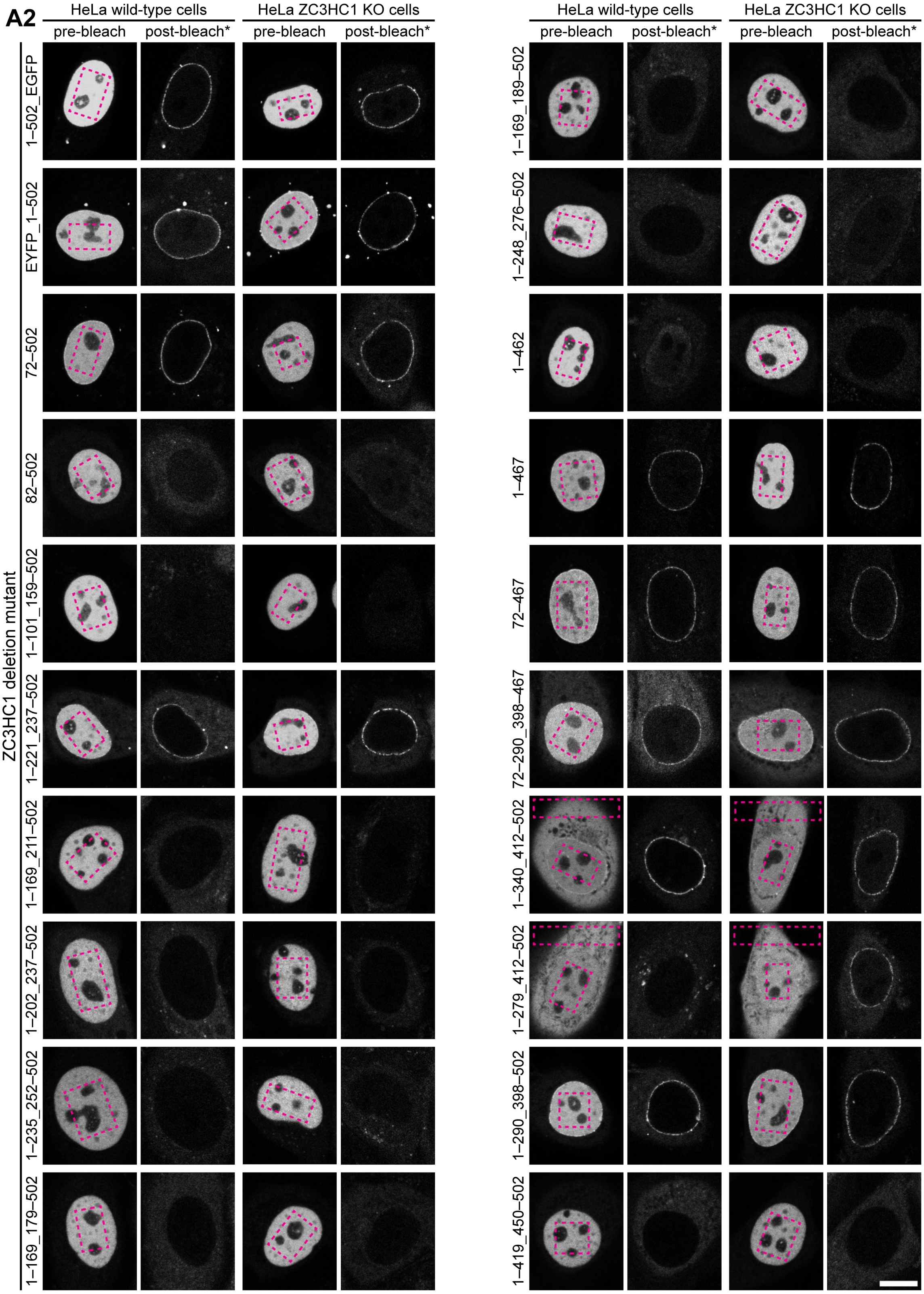
A2
HeLa wild-type cells
HeLa ZC3HC1 KO cells
pre-bleach
post-bleach*
ZC3HC1 deletion mutant
1–502_EGFP
EYFP_1–502
72–502
82–502
1–101_159–502
1–221_237–502
1–169_211–502
1–202_237–502
1–235_252–502
1–169_179–502
1–169_189–502
1–248_276–502
1–462
1–467
72–467
72–290_398–467
1–340_412–502
1–279_412–502
1–290_398–502
1–419_450–502

**S2 (2/8)**

**B1**

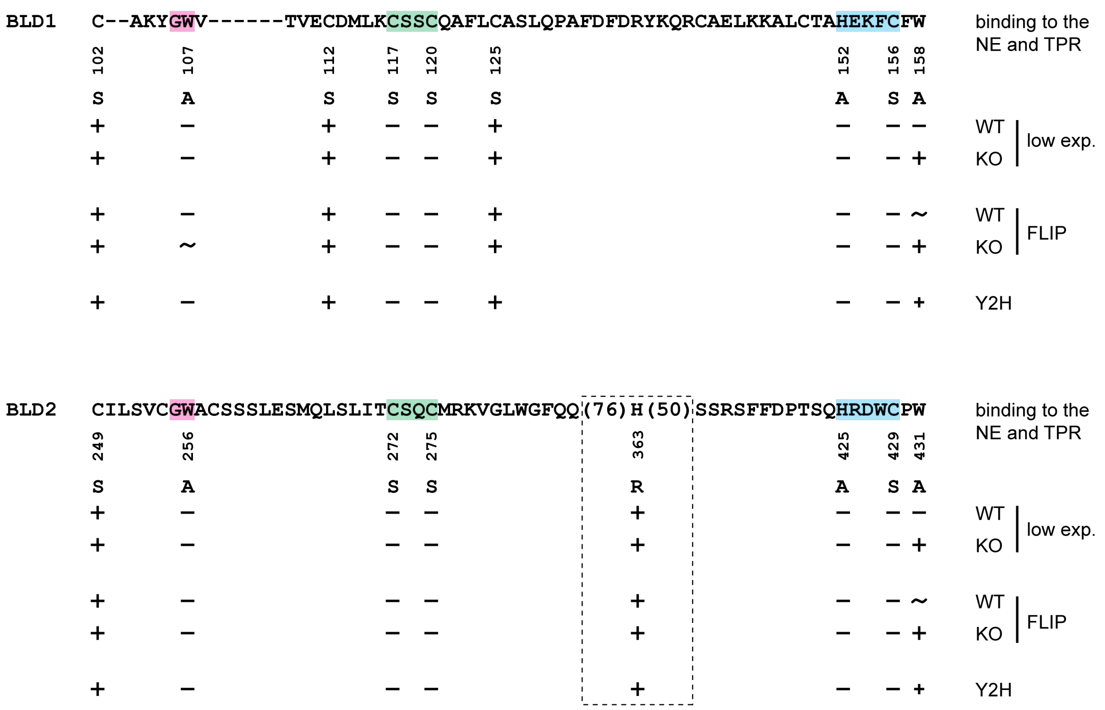
BLD1
C--AKYGWV------TVECDMLKCSSCQAFLCASLQPAFDFDRYKQRCAELKKALCTAHEKFCFW
binding to the NE and TPR
102 107 112 117 120 125 152 156 158
S A S S S S A S A
WT
KO
low exp.
FLIP
Y2H
BLD2
CILSVCGWACSSSLESMQLSLITCSQCMRKVGLWGFQQ(76)H(50)SSRSFFDPTSQHRDWCPW
binding to the NE and TPR
249 256 272 275 363 425 429 431
S A S S R A S A
WT
KO
low exp.
FLIP
Y2H

S2 (3/8)

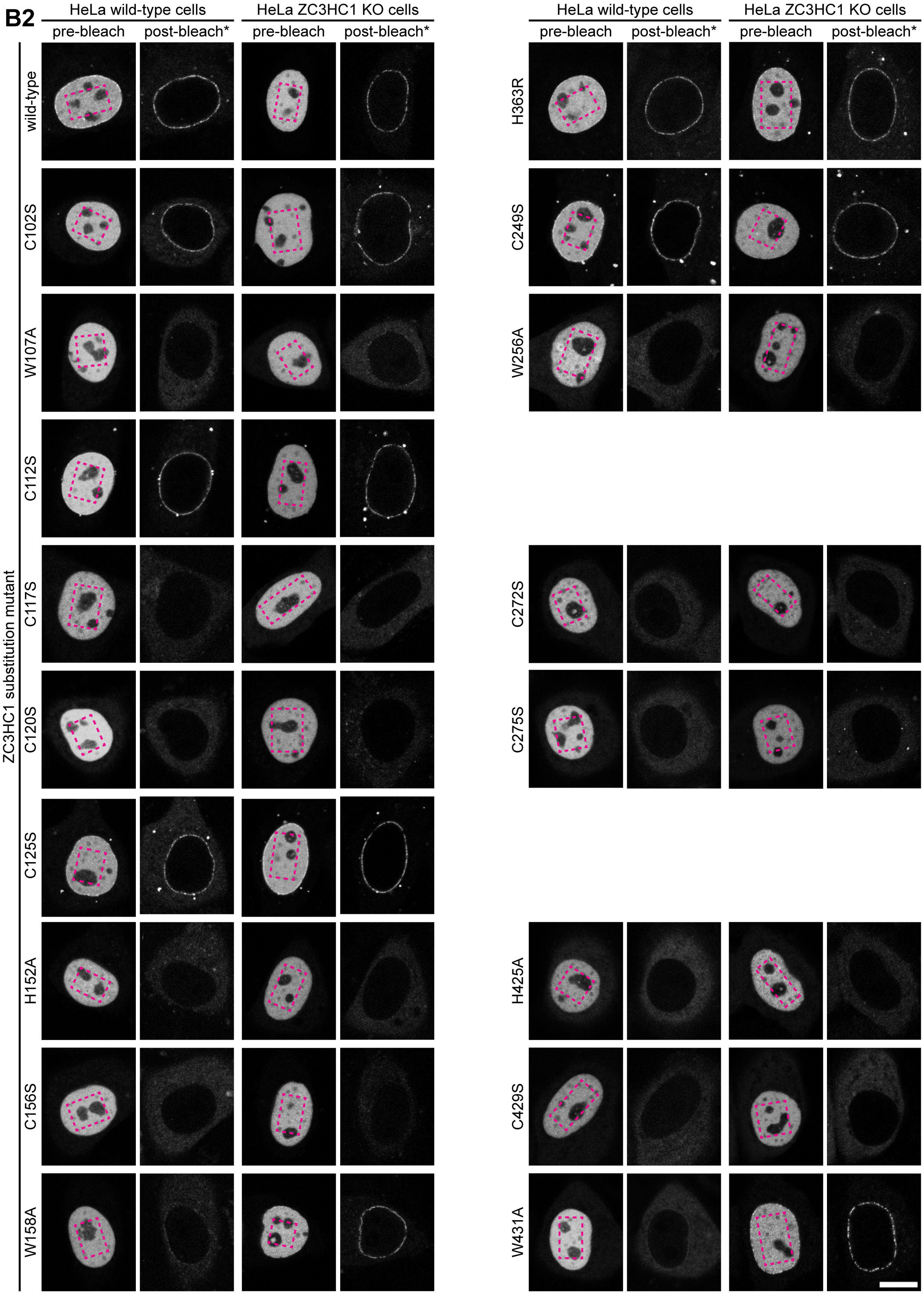
B2
HeLa wild-type cells
HeLa ZC3HC1 KO cells
pre-bleach
post-bleach*
ZC3HC1 substitution mutant
wild-type
C102S
W107A
C112S
C117S
C120S
C125S
H152A
C156S
W158A
H363R
C249S
W256A
C272S
C275S
H425A
C429S
W431A

S2 (4/8)

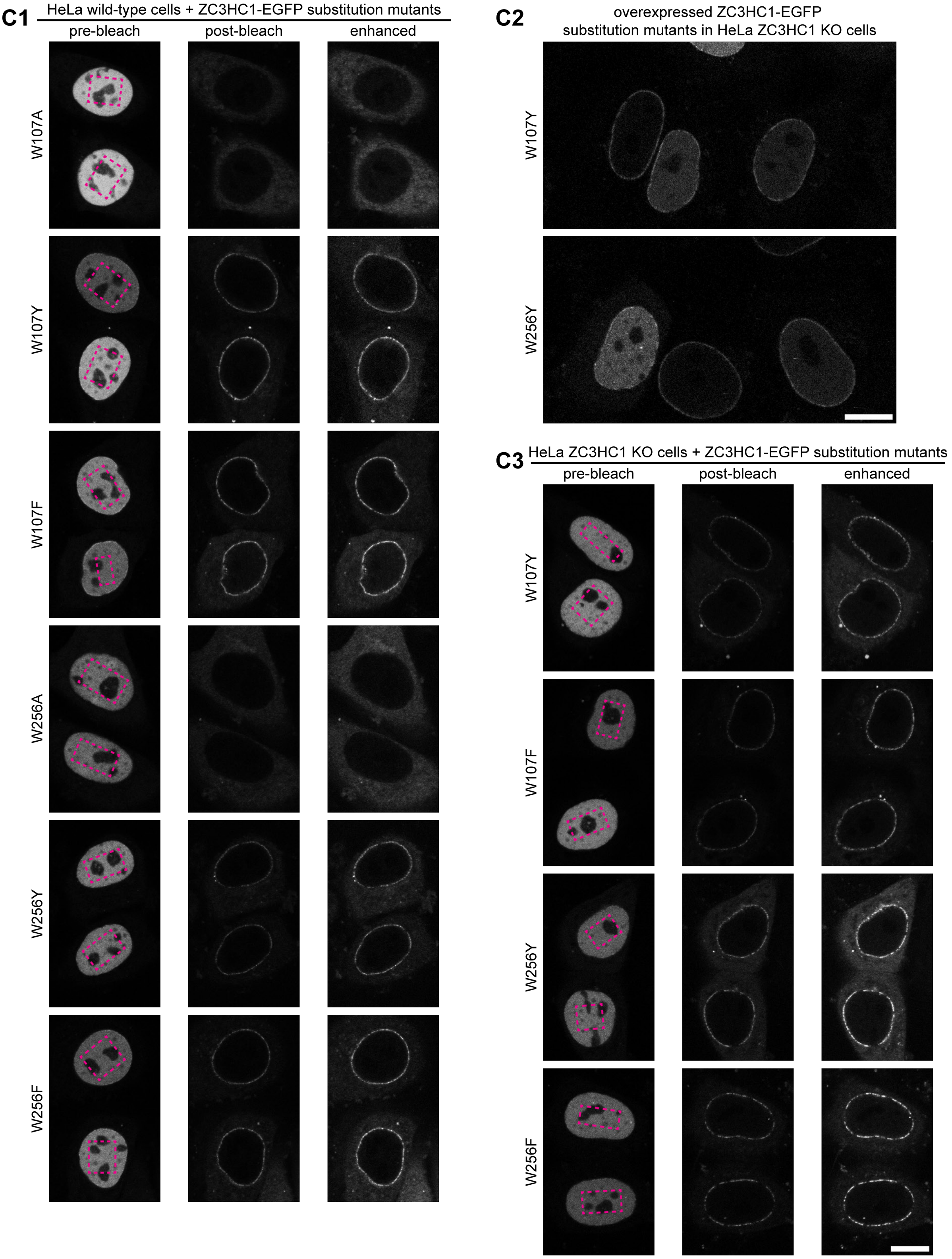
C1
HeLa wild-type cells + ZC3HC1-EGFP substitution mutants
pre-bleach
post-bleach
enhanced
W107A
W107Y
W107F
W256A
W256Y
W256F
C2
overexpressed ZC3HC1-EGFP
substitution mutants in HeLa ZC3HC1 KO cells
W107Y
W256Y
C3
HeLa ZC3HC1 KO cells + ZC3HC1-EGFP substitution mutants
pre-bleach
post-bleach
enhanced
W107Y
W107F
W256Y
W256F

**S2 (5/8)**

**D1**

zf-C3HC / PF07967 (Pfam 16 release)

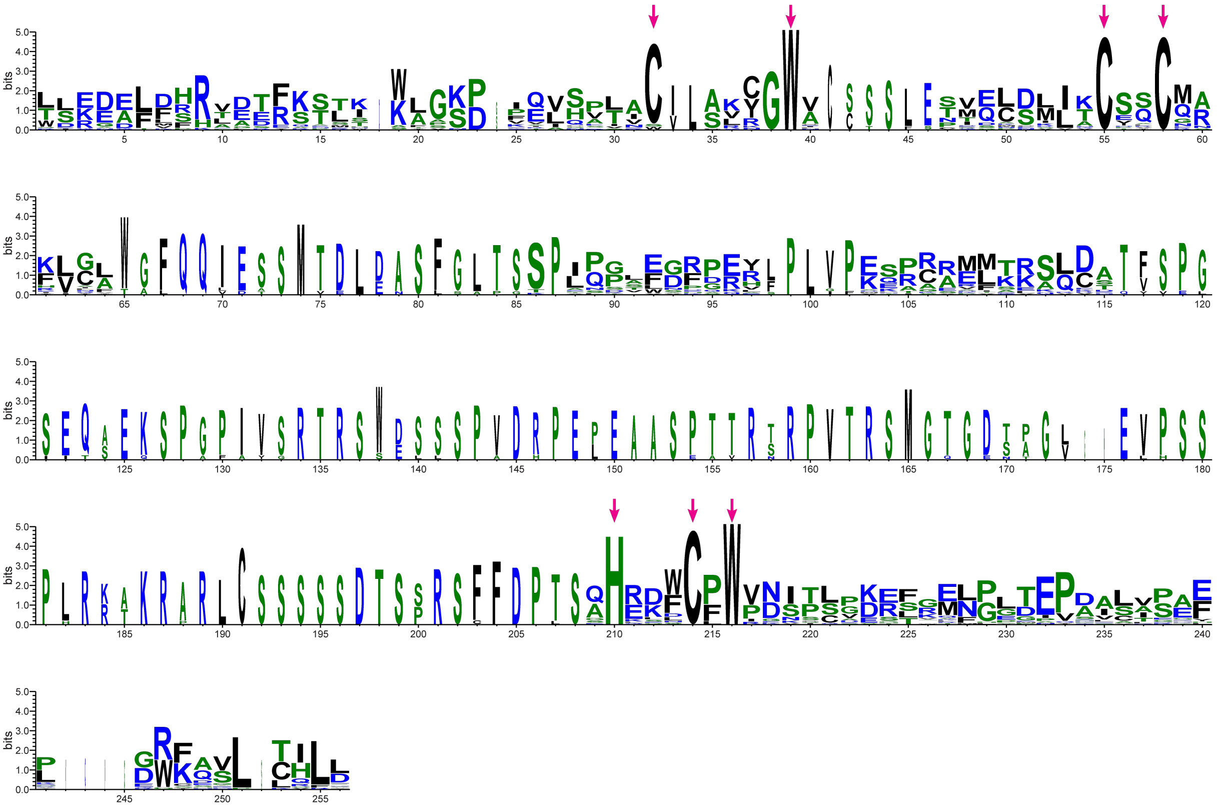

Rsm1 / PF08600 (Pfam 20 release)

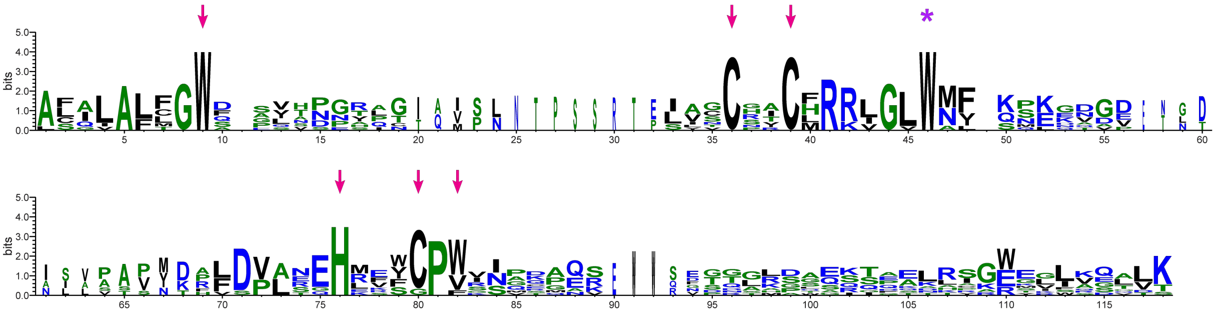

**D2**

$$G\text{-}[WYF]\text{-}X_{(8,72)}\text{-}C\text{-}X_{(2)}\text{-}C\text{-}X_{(15,524)}\text{-}H\text{-}X_{(3)}\text{-}C\text{-}X_{(62,117)}\text{-}G\text{-}[WYF]\text{-}X_{(8,72)}\text{-}C\text{-}X_{(2)}\text{-}C\text{-}X_{(15,524)}\text{-}H\text{-}X_{(3)}\text{-}C$$

**S2 (6/8)**

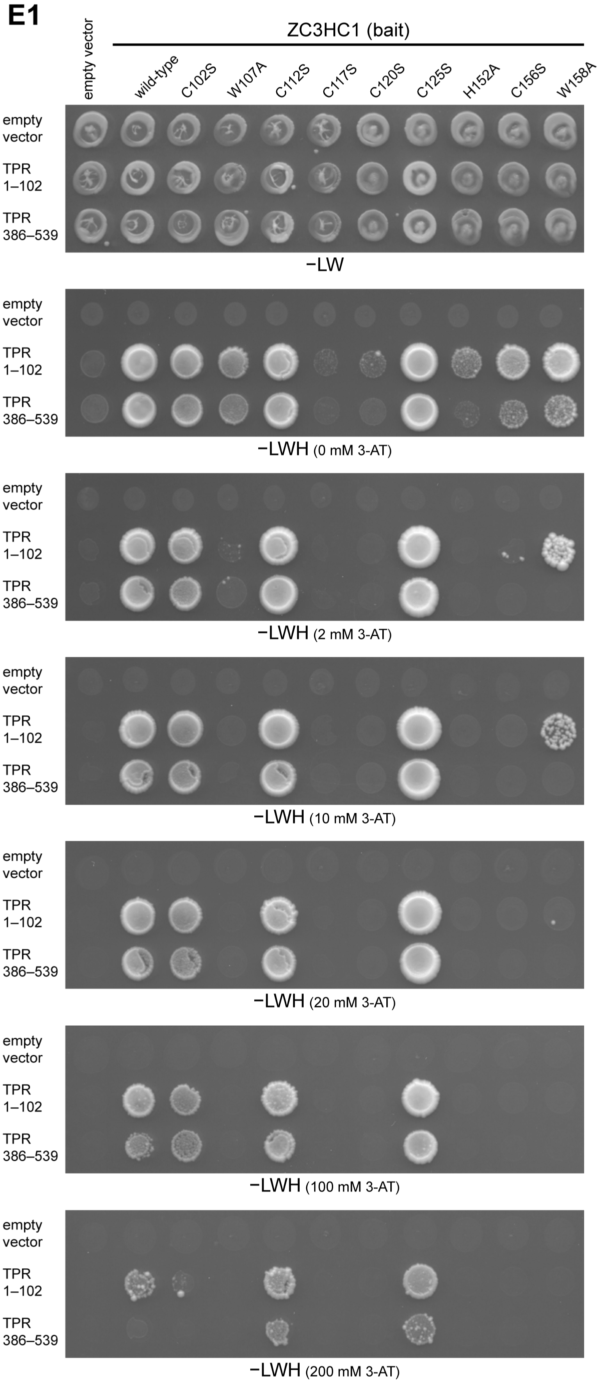
E1
empty vector
ZC3HC1 (bait)
wild-type
C102S
W107A
C112S
C117S
C120S
C125S
H152A
C156S
W158A
empty vector
TPR 1–102
TPR 386–539
−LW
−LWH (0 mM 3-AT)
−LWH (2 mM 3-AT)
−LWH (10 mM 3-AT)
−LWH (20 mM 3-AT)
−LWH (100 mM 3-AT)
−LWH (200 mM 3-AT)

S2 (7/8)

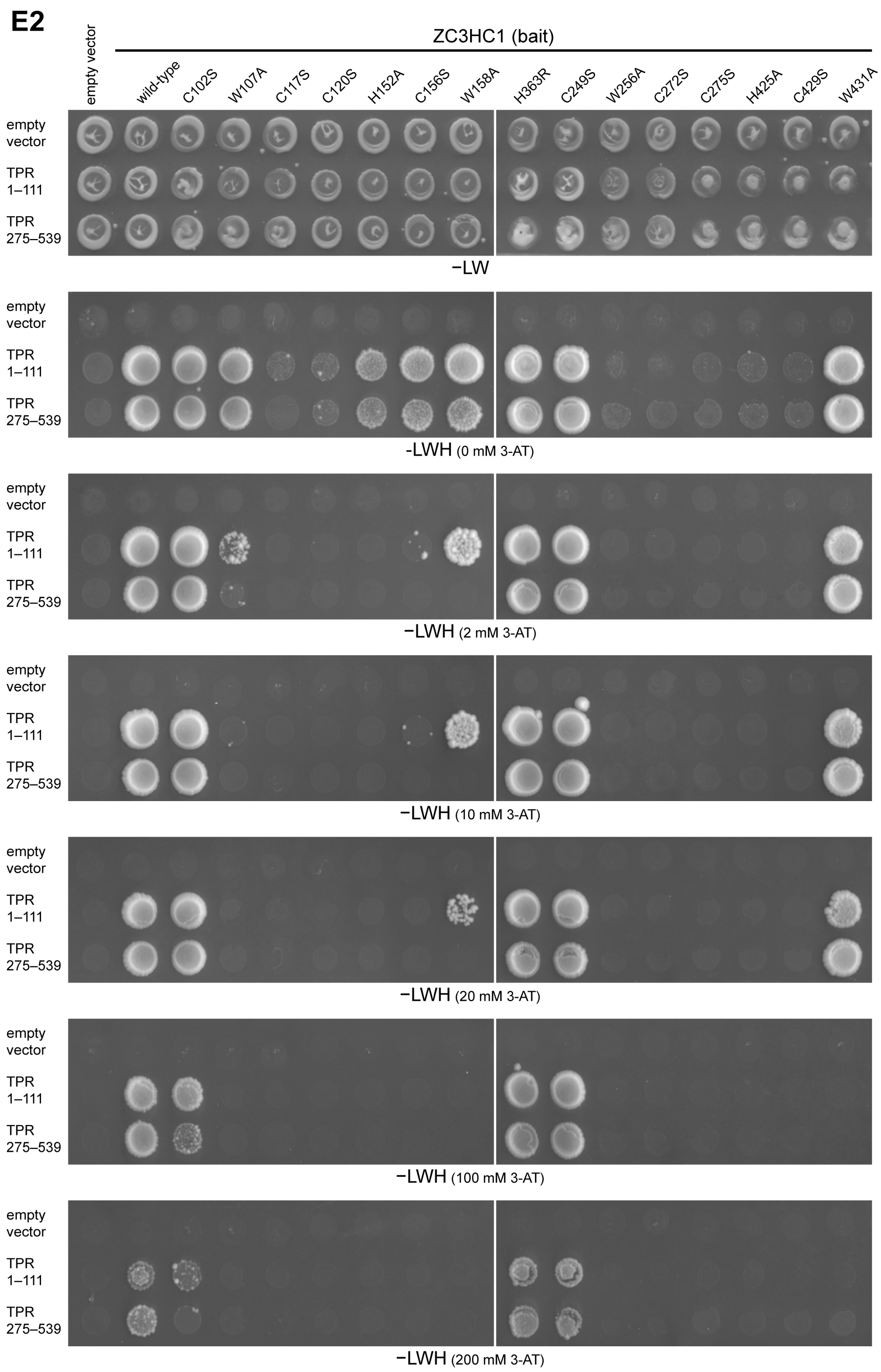

**Supplemental Figure S2. Characterization of ZC3HC1 deletion and single aa-substituted mutants, ectopically expressed in HeLa WT and ZC3HC1 KO cells, by fluorescence microscopy and FLIP, complemented by additional Y2H experiments.**

**(A, B)** Photobleaching experiments with FP-tagged deletion and single aa substitution mutants of ZC3HC1.

**(A1)** Schematic depiction of expression vector-encoded deletion mutants of ZC3HC1, all containing either an N-terminal EYFP-tag or a C-terminal EGFP-tag, with these tags not depicted though (for details, see Supplemental Table 1). Schemes are designed corresponding to the main Figure 1C, with the expanse of the central parts of the by then explained BLDs in grey, with each BLD's putative zinc ion coordinating sequences highlighted as green and blue boxes, respectively. Areas additionally highlighted represent the positions of evolutionarily conserved G-W dipeptides (magenta-colored boxes) and the protein's NLS (yellow box). The capability of binding to the NE as observed at low expression levels in HeLa P2 WT cells and HeLa ZC3HC1 KO cells is indicated in the first and second of the four columns at the right margin. Note that once such KO cells had become available later in our study (Gunkel *et al*, 2021), we conducted the experiments with them similarly to those initially done with only the HeLa WT cells. Until then, we had not yet been able to exclude that some of the ectopically expressed ZC3HC1 mutants that seemingly were NE-binding-incompetent in HeLa WT cells might merely be unable to successfully compete against the transfected cells' endogenous ZC3HC1 for NB binding sites. All the experiments with the KO cells were thereby correspondingly also carried out once again with the WT cells in parallel to allow for direct data comparability, with these thus repeated WT cell experiments confirming the findings obtained initially with the WT cells only.

Note further that the rating in the first two columns did not relate to those cells in which very high expression levels had resulted in the proteins' distribution throughout the nuclear interior and then even in notable occurrence in the cytoplasm despite possessing a functional NLS. Furthermore, the third and fourth columns indicate whether NE-localization was still or only notable after having photobleached the nuclear interior of WT and KO cells expressing the recombinant proteins in high amounts after having been transfected with the listed expression vectors. Note that this approach revealed that one of the deletion mutants, ZC3HC1 1–279_412–502, which had not been seen at the WT cell's NE, can bind to the NE of the ZC3HC1 KO cell, also despite lacking the protein's NLS, the latter comprising at least aa 398–404 (described as 396–402 in Ouyang *et al*, 2003). Even though the loss of this NLS largely impairs nuclear import of ZC3HC1, we found small amounts of this mutant protein eventually entering the nucleus nonetheless, which also held for ZC3HC1 1–340_412–502 and which was likely due to NPCs not being perfect permeability barriers (Güttler & Görlich, 2011; Kırlı *et al*, 2015). While we already found it noteworthy at this point that mutants like 1–279_412–502 and 1–340_412–502 were capable of NE-association, we had not yet been able though to exclude another scenario in which minor amounts of NLS-deficient ZC3HC1 polypeptides would have been co-imported with TPR, as a result of some sporadic binding to TPR already in the cytoplasm. However, a later study (Gunkel & Cordes, 2022) eventually provided unambiguous evidence proving that NB association in the absence of the NLS reflected the intact ZC3HC1 protein's capability of binding to the NB also without engaging in a cytoplasmic interaction with TPR.

Further note that among those ZC3HC1 mutants not listed in the dataset presented in Figure 1 are also such that we had created for further addressing the question as to whether distinct sequence segments of region aa 159–254, located between those parts of the minimal core sequence signature of the first and second BLD found essential for NB-binding, might be required too. This revealed that even further minor deletions within this area, including the aa segments 170–210, 170–188, 170–178, 203–236, and 236–251, abolished its positioning at the NE, even though these mutants' C-$X_{(2)}$-C and H-$X_{(3)}$-C peptide sequences were intact. These findings indicated that other distinct sequence features or a certain length of this inter-domain region are also needed to allow for an NB-binding interface. We found merely the deletion of one of these short segments (222–236) tolerable.

**(A2)** Live-cell imaging and photobleaching of HeLa WT and HeLa ZC3HC1 KO cells transiently transfected with the expression vectors encoding the deletion mutants referred to in S2A1. Since it had sometimes turned out difficult to judge whether some of the ectopically expressed mutant versions of FP-tagged ZC3HC1 had actually engaged in some interaction with the NE, next to being uniformly distributed throughout the nuclear interior, we bleached this latter pool of polypeptides in order to this then perhaps allow for visualizing higher numbers of the FP-tagged polypeptides at the NE, which otherwise might have remained undetectable due to signal intensities at the NE not exceeding those concomitantly accumulating within the nuclear interior.

Marked rectangular areas in the pre-bleach images of representative cells were subjected to full laser power, followed by the acquisition of a post-bleach image, with the laser settings then again as for the pre-bleach image. Signal brightness of the here shown post-bleach images was enhanced electronically using the Multiply command in the Math Submenu of the ImageJ/Fiji software (version 2.0.0-rc-64/1.51t, National Institutes of Health, USA), with such enhancement here indicated by an asterisk. Each pixel value was thereby multiplied by the same multiplication factor, allowing the signal intensity relationships between the electronically brightness-enhanced images to remain essentially the same as between the corresponding raw images beyond their background zero values. The same procedure was also applied to obtain the signal-enhanced images presented in S2B2. Note that after near-quantitative elimination of the nuclear fluorescence, the NEs of many cells ectopically expressing either one or the other mutant version of ZC3HC1 were also not detectable. In contrast, in other cells expressing yet other mutants, the NE emerged as the only structure that was still fluorescent, pointing at an *in vivo* interaction between the NE and the respective ZC3HC1 mutant. As an aside, note that in those cells ectopically expressing the NLS-deficient ZC3HC1 mutants, the cytoplasmic compartment had also been bleached to render potential signals at the NE better visible. Bar, 10 µm.

**(B1)** Compilation of the single aa substitutions of *Hs*ZC3HC1 presented or referred to in the current study context. Indicated are the mutations' effects (i) on the protein's capability of binding to the NB, as studied in HeLa WT and ZC3HC1 KO cells with and without photobleaching, as done for the deletion mutants presented in S2A. In experiments without photobleaching, NE-association was rated in cells with a low level of the mutant protein's constitutive expression, while in the FLIP experiments, such cells were bleached in which expression levels were regarded as high. Indicated, furthermore, are the mutations' effects (ii) on the protein's capability of interacting with TPR in Y2H experiments (see further below). Thus, for example, we found ZC3HC1 with either the single aa substitution W158A or W431A to occur at the NE of a ZC3HC1 KO cell, while both did not appear capable of NE-association when the native ZC3HC1 was present. Furthermore, also note that the dashed rectangle accentuates one particular aa and its substitution, namely R363H, that is not part of one of the actual BLD sequence stretches depicted in grey in S2A1 but whose performance with regard to NB- and TPR-binding we studied nonetheless. Back then, R363H had been recently reported to represent a naturally occurring single aa polymorphism of ZC3HC1 in humans, with pathophysiological phenotypes connected to coronary artery diseases (CAD) assigned to the presence of an arginine at aa position 363 while a histidine at the same position instead was regarded as a non-effect residue (Schunkert *et al*, 2011). Positioned within the large, apparently unstructured loop inserted into the second BLD of ZC3HC1 that we had found dispensable for NB association, we nonetheless inspected whether one of the two amino acids at 363 might notably affect the protein's binding to the NB and TPR. However, studying both variants upon their ectopic expression in HeLa cells in parallel, we found both the R363 and the H363 version of full-length ZC3HC1 seemingly similarly well capable of binding to the NE. Furthermore, both variants were found equally well capable of interacting with both of TPR's ZC3HC1 binding domains in Y2H experiments (see also further below).

**(B2)** Fluorescence microscopy and photobleaching of HeLa WT and HeLa ZC3HC1 KO cells transiently transfected with the expression vectors encoding the aa substitution mutants referred to in S2B1. Experiments were performed, and data were presented, like in S2A2. Bar, 10 µm.

**(C)** Live-cell imaging and FLIP of HeLa WT (S2C1) and ZC3HC1 KO cells (S2C2 and S2C3) transiently transfected with the expression vectors coding for further ZC3HC1 mutants, with either the single aa substitution W017Y, W107F, W256Y, or W256F. In those experiments initially conducted in the WT cells, FLIP of the aa substitutions W017A and W256A were conducted in parallel for comparison. Note that the W017Y, W107F, W256Y, and W256F substitutions already allowed for NE association in the WT cells, revealing that these aa substitution mutants could successfully compete with the endogenous ZC3HC1 for NB binding sites, in contrast to W107A and W256A. As an aside, note that the here presented experiments in the WT and ZC3HC1 KO cells had been conducted at different timepoints, in contrast to those presented in S2B2 in which the initial experiments in WT cells later had been repeated, next to the then available KO cells studied in parallel, to identify substitution mutants that were only capable of NE binding in the absence of ZC3HC1. Bars, 10 µm.

**(D)** Illustration of the sequence signatures of the initial versions of the Pfam zf-C3HC and Rsm1 motifs, next to an early low-stringency signature of the BLD-tandem motif. The latter and other versions of the BLD-tandem signature were later collectively referred to as signatures of the nuclear basket-interaction domain (NuBaID; see further below).

**(D1)** WebLogos (Crooks *et al*, 2004) that we had generated with an online tool (https://weblogo.berkeley.edu/) and the original Pfam MSAs for the zf-C3HC and Rsm1 motifs. The latter had first been described in the Pfam 16.0 and 20.0 releases, respectively, with the underlying MSAs deposited in the corresponding Pfam-A full datasets (http://ftp.ebi.ac.uk/pub/databases/Pfam/releases/Pfam16.0/; http://ftp.ebi.ac.uk/pub/databases/Pfam/releases/Pfam20.0/; e.g., Finn *et al*, 2006, 2008). For the original version of the Rsm1 motif, the MSA comprised 11 sequence segments from 10 species, of which seven were of fungal origin, including *Sp*Rsm1p, but initially no mammalian sequences. Furthermore, the sequence segments for the Rsm1 motif corresponded only to the second BLD, with one of these sequences being redundant. On the other hand, for the original zf-C3HC motif, the MSA included a total of 32 sequence segments from seven species, including three vertebrates and *Schizosaccharomyces pombe*, the latter again represented by *Sp*Rsm1p. Furthermore, 18 sequence segments corresponded to the first BLD and 14 to the second BLD, with 9 and 11, respectively, being redundant. Of further note, the zf-C3HC motif had already in the Pfam release 16.0 (released November 2004) been described as representing a domain *"often repeated, with the second domain usually containing a large insert (approximately 90 residues) after the first three cysteine residues"* (http://ftp.ebi.ac.uk/pub/databases/Pfam/releases/Pfam16.0/). Such realization might explain why 14 of the 32 sequence segments initially used for the motif definition represented a BLD2. In other words, for the initial version of the zf-C3HC motif, sequences corresponding to both BLD1 and BLD2 had been used.

In addition to the WebLogos, also note that some residues are marked by magenta-colored arrows, with these residues representing a selection of those for which aa substitutions are presented in the current study. Other aa substitutions, e.g., a tryptophan marked by an asterisk in purple as part of the Rsm1 motif, will be presented in another context elsewhere. Further note that even minimal signature versions deducible from the zf-C3HC motif based on these sequences, like C-$X_{(3,6)}$-G-W-$X_{(9,15)}$-C-$X_{(2)}$-C-$X_{(31,149)}$-H-$X_{(3)}$-C-X-W, would not have allowed for identifying the Pml39 protein of budding yeast, while the ZC3HC1 homolog in *Dictyostelium discoideum* would have been detectable. Regarding the Rsm1 motif, a minimal signature like G-W-$X_{(10,25)}$-C-$X_{(2)}$-C-X-R-$X_{(4)}$-W-$X_{(15,29)}$-H-$X_{(3)}$-C-P-W would have neither allowed for detecting the budding yeast nor the *D. discoideum* homolog.

**(D2)** The tandem arrangement of the minimal BLD signature referred to in the main text as signature (2), having taken into account, i.a., the dispensability of C102 of *Hs*ZC3HC1 (Supplemental Figure S2B) and the replaceability of W for Y or F (e.g., Supplemental Figure S2C). Furthermore, this BLD-tandem signature had considered the Pfam Database's information and the other published data available until then (Higashi *et al*, 2005; Kokoszynska *et al*, 2008), having also deduced from the latter the here chosen spacing between the two identical BLD signatures. Note that while the resulting BLD-tandem signature, G-[WYF]-$X_{(8,72)}$-C-$X_{(2)}$-C-$X_{(15,524)}$-H-$X_{(3)}$-C-$X_{(62,117)}$-G-[WYF]-$X_{(8,72)}$-C-$X_{(2)}$-C-$X_{(15,524)}$-H-$X_{(3)}$-C, allowed for identifying the budding yeast's Pml39 protein via ScanProsite, it did not detect the *D. discoideum* homolog.

**(E)** Single aa substitution mutants of ZC3HC1 that were studied in Y2H experiments, in combination with two different sets of two ZC3HC1 interaction domain segments of TPR (S2E1 and S2E2). Here, ZC3HC1 was expressed as the Y2H bait, representing a fusion protein including the N-terminally appended GAL4 DNA-binding domain (GAL-BD). Correspondingly, the empty vector represented the one only expressing the GAL4 DNA-activation domain (GAL4-AD), while the TPR segments, as the Y2H preys, represented GAL4-AD-TPR fusion polypeptides. Both data sets include the presentation of representative colony growth on the selection medium lacking leucine and tryptophan (−LW), and the actual Y2H interactions, after replica-plating onto the selection medium lacking leucine, tryptophan, and histidine (−LWH), supplemented with different concentrations of 3-AT. Some of the data shown here, included for comparison, are also presented in Figure 2C. Note that those single aa substitution mutants of ZC3HC1 that did not impair NE association (in S2B1, those marked with NE+) allowed for colony growth on selective medium minus LHW when paired with ZC3HC1-binding domains of TPR. By contrast, no colony growth on −LHW medium was observed when single aa substitution mutants of ZC3HC1, which have been found incapable of NE association in HeLa cells (NE− in S2B1), had been co-expressed with TPR's ZC3HC1-binding domains. Of additional note, we found ZC3HC1 mutant W158A capable of a seemingly more robust interaction with those segments of TPR that harbored a ZC3HC1 binding domain located near TPR's NT, here represented by TPR 1–102 and TPR 1–111, while already at a concentration of 2 mM 3-AT, no interaction was observed between ZC3HC1 W158A and either TPR 275–539 or TPR 386–539, both harboring another ZC3HC1 binding site of TPR. However, the analysis of this finding in further detail was not considered a topic of the current study.

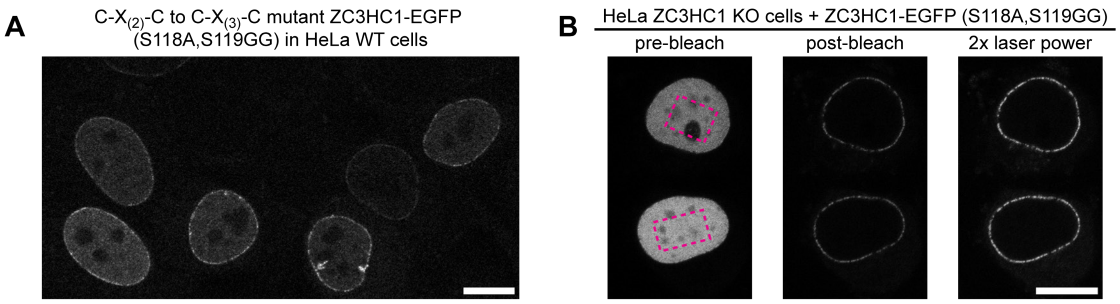

**Supplemental Figure S3. Functional tolerance of *Hs*ZC3HC1 for a C-X$_{(3)}$-C spacing within its first BLD, as commonly present in some fungal ZC3HC1 homologs.**

Since the ZC3HC1 homologs of some fungi, for example, of the genus *Aspergillus*, exhibited a different spacing between the first two cysteines of the first BLD's zinc finger signature, reading C-X$_{(3)}$-C, this had raised the question of whether such a particular C-X$_{(3)}$-C constellation might still allow for NB-association. This, in turn, had prompted us to create yet additional *Hs*ZC3HC1 mutants. Among these was one in which we exchanged the human homolog's original C-S-S-C sequence between aa 117–120 for the correspondingly positioned sequence C-A-G-G-C, which is present, for example, in *Aspergillus rambellii*.

(**A**) Fluorescence microscopy of HeLa P2 WT cells transiently transfected with an expression vector coding for the mutant version of ZC3HC1 in which we had replaced the first BLD's sequence C-S-S-C by C-A-G-G-C. Note that we found this *Hs*ZC3HC1 mutant, even at low expression levels, well capable of binding to the NE of human cells, with this being so, remarkably enough, even in the presence of the endogenous wild-type version of ZC3HC1.

(**B**) Photobleaching of HeLa ZC3HC1 KO cells transiently transfected with the expression vector encoding the C-X$_{(3)}$-C aa substitution mutant referred to in S3A. Experiments were performed like in Supplemental Figure S2A2, except that the here shown signal-enhanced post-bleach image did not represent the outcome of an electronic signal enhancement but an additional image acquired with doubled laser power after the first post-bleach image had been taken, like also in S1A. Note that the mutant remained stably bound to the NE after having eliminated the fluorescence of the nuclear pool. Bars, 10 µm.

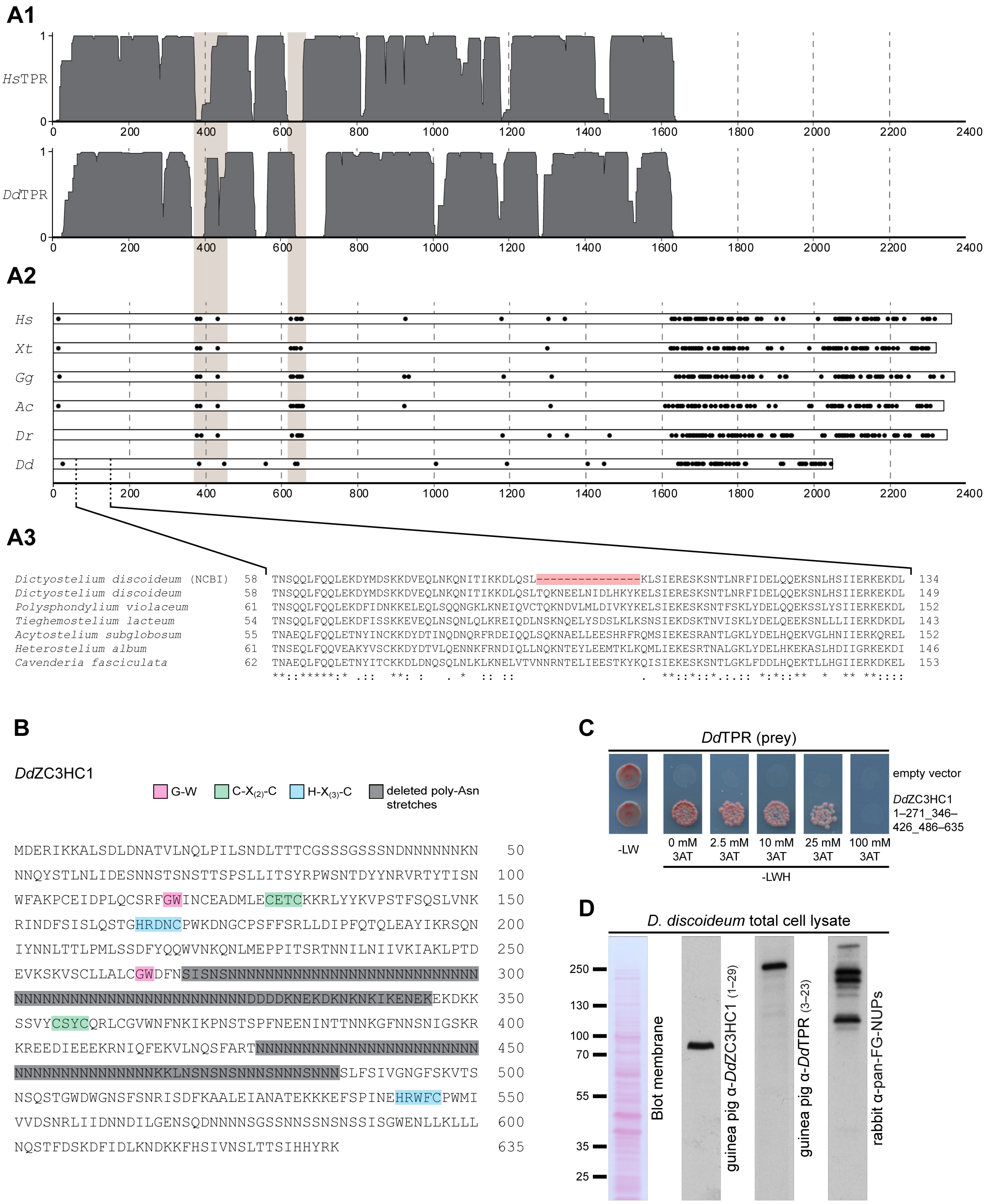

**Supplemental Figure S4. Characterizing the *Dictyostelium* homologs of TPR and ZC3HC1.**

**(A)** Sequence features of *Dictyostelium discoideum* TPR compared to TPR homologs from other species.

**(A1)** The coiled-coil (CC)-forming regions of *Dd*TPR compared to those of *Hs*TPR, as here predicted by the PCOILS algorithm (Zimmermann *et al*, 2018; https://toolkit.tuebingen.mpg.de/

#/tools/pcoils) based on windows of 28 residues. The CC-forming parts are shown as areas in dark grey. The two vertical rectangles in light brown delineate the area between about aa 400–600 of *Hs*TPR formerly found containing those sequence elements required for TPR's binding to the NE. These elements are flanked by short, hinge-forming sequence stretches involving evolutionary-conserved prolines or other coiled-coil-disrupting features at these sites (Hase *et al*, 2001; Kuznetsov *et al*, 2002; Krull *et al*, 2004; Gunkel *et al*, 2021). The 2052 aa-long *Dd*TPR sequence (NCBI accession number ON368702) used for these predictions is the result of RNA isolation from *D. discoideum* Ax4 cells and subsequent cDNA synthesis and sequencing.

**(A2)** Schemes of vertebrate TPR homologs, comprising the one from humans (*H*s, accession number NP_003283.2), *Xenopus tropicalis* (*Xt*, XP_002933814.3), *Gallus gallus* (*Gg*, XP_004943365.4), *Anolis carolinensis* (*Ac*, XP_016850554.1) and *Danio rerio* (*Dr*, NP_001025294.1), next to the one for *Dd*TPR (ON368702). The black dots within each of the horizontal rectangles representing the different TPR homologs illustrate the positions of proline residues, of which, as expected, only a few are present within the CC-forming parts. In particular, though, proline residues flanking the NPC/NB-binding elements of *Hs*TPR, and evolutionarily conserved in other vertebrates, also exist at similar positions in the CC domain of *Dd*TPR. By contrast, proline residues are abundant in the TPR homologs' carboxyterminal domains (see also Kuznetsov *et al*, 2002), with this domain shown to be largely unstructured (Hase *et al*, 2001).

**(A3)** Alignment of an aa sequence segment from the N-terminal domain of different amoebic species, including two sequences for *D. discoideum* (accession numbers XP_636884.1 and ON368702, first and second line, respectively), with the first one derived from an annotated genomic sequence (NC_007091), lacking residues evolutionarily conserved in the slime molds. The other aligned sequences stem from hypothetical proteins representing evident TPR homologs of other species of the class Dictyostelia, including *Polysphondylium violaceum* (KAF2071913.1), *Tieghemostelium lacteum* (KYR02800.1), *Acytostelium subglobosum* LB1 (XP_012748738.1), *Cavenderia fasciculata* (XP_004357092.1) and *Heterostelium album* PN500 (XP_020436491.1), with the latter sequence only representing a segment of this species' TPR homolog.

**(B)** Sequence of the ZC3HC1 homolog of *D. discoideum*. Residues highlighted in magenta, green, and blue represent the positions of the two BLDs' G-W, C-$X_{(2)}$-C, and H-$X_{(3)}$-C sequence elements. Those regions highlighted in grey represent the residues that we removed, by cloning, from some of the *Dd*ZC3HC1 sequences used for expression experiments in budding yeast cells (e.g., in S4C) and in mammalian cells (data not shown). The 635 aa-long sequence (NCBI accession number ON368701) results from our RNA isolation from *D. discoideum* Ax4 cells and subsequent cDNA synthesis and sequencing. The protein's C-terminal residues differ from the NCBI-deposited sequence for a 647 aa-long *Dd*ZC3HC1 from Ax4 cells (XP_638576.1), the latter derived from an annotated genomic sequence (NC_007090).

**(C)** Exemplary Y2H data obtained with expression vectors coding for a GAL4-BD version of *Dd*ZC3HC1 as the bait polypeptide, here with its two BLDs both intact while lacking those asparagine-dominated sequence segments highlighted in grey in S4B. Correspondingly, the empty vector represented the one only expressing the GAL4-BD, while the *Dd*TPR segment used here as the Y2H prey was thus fused to GAL4-AD.

**(D)** Immunoblotting (IB) of the total of proteins from *D. discoideum* cells of line Ax2, with those IFM-compatible antibodies for *Dd*TPR and *Dd*ZC3HC1 that were also used for the micrographs presented in Figure 4C. IB with pan-FG-NUPs antibodies, cross-reactive with the FG repeat domains of numerous nucleoporins, are shown for comparison. Target regions of *Dd*ZC3HC1 and *Dd*TPR are given in parentheses. Immunolabeling was performed on the representative Ponceau S-stained membrane shown here and on replicates of the identical kind. As an aside, note that generating some of the required antibodies for *Dd*ZC3HC1 turned into an unexpectedly challenging enterprise, with each antibody having to be versatile for both IB and IFM while at the same time not allowing it to be cross-reactive with unrelated proteins. In particular, basic requirements that had to be met by all selected antibodies also included being suitable for at least one same IFM protocol from among the various published, including notably different ones established for studying different target proteins in *Dictyostelium*. After systematically testing such published IFM protocols and others that we had newly conceived for *Dictyostelium*, we realized that several of here not shown IB-compatible antibodies against *Dd*TPR and *Dd*ZC3HC1 could not be used for double-labeling IFM because some performed only in the one while others only in the other IFM protocol. However, despite such unanticipated difficulties concerning IFM in *Dictyostelium*, we obtained at least some affinity-purified antibodies that allowed for the proteins' comparison under identical IFM conditions.

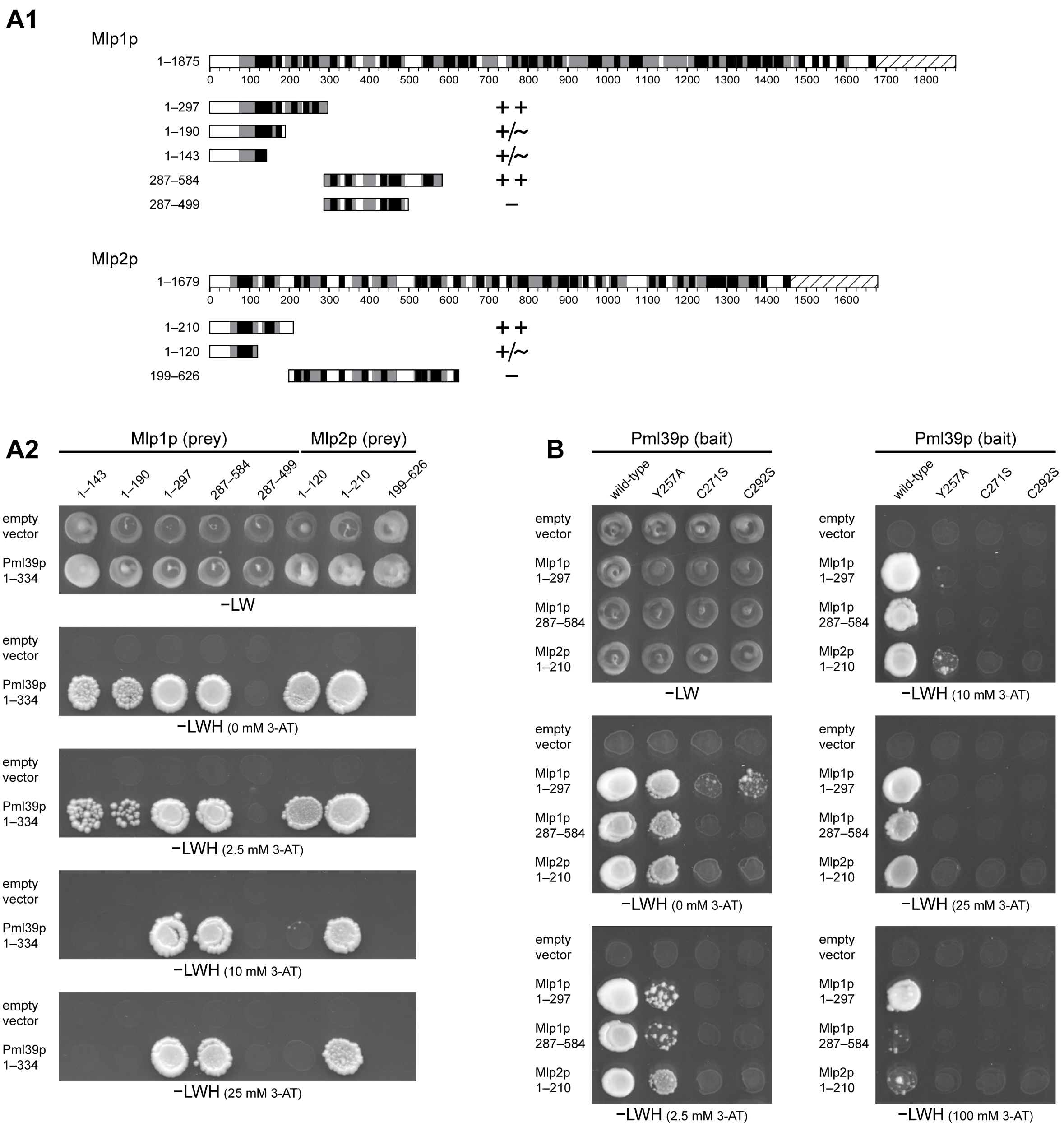
A1
Mlp1p
1–1875
0 100 200 300 400 500 600 700 800 900 1000 1100 1200 1300 1400 1500 1600 1700 1800
1–297 ++
1–190 +/~
1–143 +/~
287–584 ++
287–499 −
Mlp2p
1–1679
0 100 200 300 400 500 600 700 800 900 1000 1100 1200 1300 1400 1500 1600
1–210 ++
1–120 +/~
199–626 −
A2
Mlp1p (prey)
Mlp2p (prey)
1–143 1–190 1–297 287–584 287–499 1–120 1–210 199–626
empty vector
Pml39p 1–334
−LW
−LWH (0 mM 3-AT)
−LWH (2.5 mM 3-AT)
−LWH (10 mM 3-AT)
−LWH (25 mM 3-AT)
B
Pml39p (bait)
wild-type Y257A C271S C292S
empty vector
Mlp1p 1–297
Mlp1p 287–584
Mlp2p 1–210
−LW
−LWH (10 mM 3-AT)
−LWH (0 mM 3-AT)
−LWH (25 mM 3-AT)
−LWH (2.5 mM 3-AT)
−LWH (100 mM 3-AT)

**S5 (1/2)**

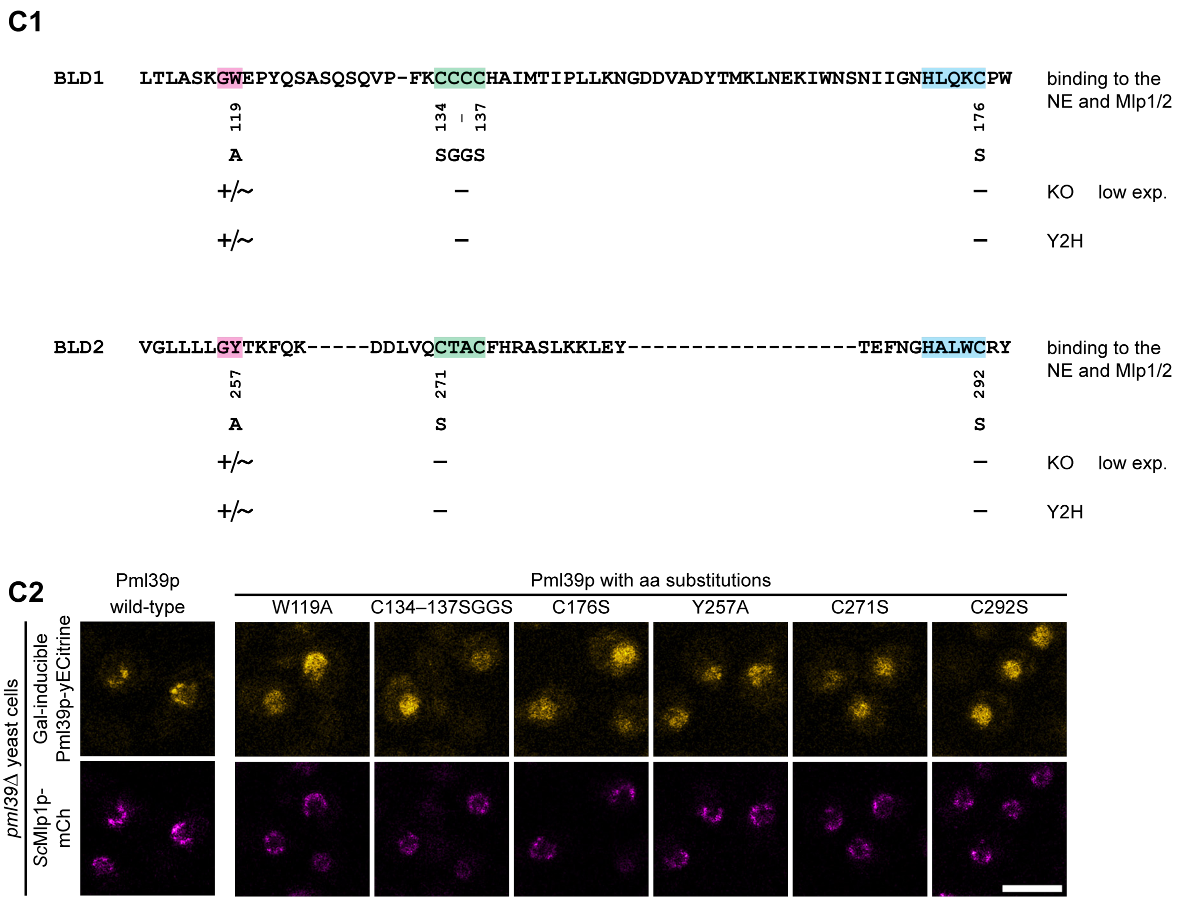

**Supplemental Figure S5. Experiments complementing the characterization of *Sc*Pml39p and its NuBaID.**

**(A)** Representative Y2H data obtained with expression vectors coding for *Sc*Mlp1p and *Sc*Mlp2p segments, fused to GAL4-AD (prey), and for the intact, wild-type version of *Sc*Pml39p, fused to GAL4-BD (bait).

**(A1)** Schemes of full-length Mlp1p and Mlp2p drawn to scale, next to segments of Mlp1p and Mlp2p encoded by Y2H expression vectors. The hatched rectangles represent the Mlp proteins' C-terminal domain. The boxes in grey and black represent those parts of each polypeptide for which the probability of forming a coiled-coil is here predicted to be at least 80% by the PCOILS algorithm (Zimmermann *et al*, 2018) within windows of 14 aa (black) and 28 aa (plus grey parts), respectively. Each segment's capability of a Y2H interaction with Pml39p is indicated (++, robust colony growth even in elevated concentrations of 3-AT; +/~, notably attenuated colony growth; −, no colony growth).

**(A2)** Representative colony growth on selection medium lacking leucine and tryptophan (−LW), revealing successful mating between the yeasts harboring the indicated GAL4-AD and GAL4-BD expression vectors, and visualization of Y2H interactions, after having replicaplated the diploid cells onto selection medium lacking leucine, tryptophan and histidine (−LWH), either without or with different concentrations of 3-AT as indicated.

**(B)** Representative Y2H results obtained with prey vectors coding for the three Mlp1p and Mlp2p segments that had allowed for Pml39p interaction in the presence of 25 mM 3-AT (S5A2) and bait vectors encoding the wild-type version of Pml39p, next to additional Pml39p mutants with single aa substitutions of the NuBaID signature. Some of the data shown here, included for comparison, are also presented in Figure 4D. Note that each of these single aa substitutions abolished the interaction with all three Mlp segments, yet at different concentrations of 3-AT. Note that the here shown Y257A mutant allowed for some attenuated Y2H interactions with the Mlps at low 3-AT concentrations, in line with some residual co-localization with Mlp1 at the NE upon ectopic expression of this mutant for live-cell imaging (see S5C2). As an aside, further note that an additional mutant, in which we introduced the single aa substitution C134S, was also capable of an attenuated Y2H interaction with both Mlp1p and Mlp2p, indicating that the Pml39 protein's three cysteines from C135 to C137 could compensate for the loss of C134 to some extent (data not shown).

**(C)** Summary and presentation of additional live-cell images of *pml39*Δ yeast cells with ectopically expressed versions of yECitrine-tagged Pml39p.

**(C1)** Sequence segments of Pml39p, representing the corresponding parts of its first and second BLD, together with a compilation of the Pml39p aa substitutions presented in the current study, next to the summary of those data shown in main Figure 4E and here in S5C2. Indicated are the mutations' effects on the protein's capability of binding to the NE, as studied in *pml39*Δ cells, upon the induced ectopic expression of the yECitrine-tagged versions of Pml39p. NE-association was rated in cells in which the mutant proteins' expression levels were regarded as still low or only moderate. In addition, provided for comparison are the summarizing results of those corresponding Y2H data available, including those presented in Figures 4D and S5B and others not shown as images, reflecting each mutant protein's capability of interacting with the Mlp proteins.

**(C2)** Live-cell imaging of yeast *pml39*Δ cells with endogenously expressed mCherry-tagged Mlp1 polypeptides and additional, galactose-induced ectopically expressed yECitrine-tagged Pml39p mutant versions, harboring either a single aa substitution of the NuBaID signature or the quadruple aa substitution C134–137SGGS. These additional mutants are here shown next to those micrographs already presented in the main Figure 4E, comprising the ones for the ectopically expressed WT version of the FP-tagged Pml39p and the single aa substitution mutants Y257A, C271S, and C292S. Note that while the wild-type Pml39p primarily accumulated at the NE, all Pml39p mutants but W119A and Y257A were found no longer capable of binding to the NE and instead distributed throughout the nuclear interior. Concerning the W119A and Y257A mutants, we detected some residual co-localization with Mlp1 at the NE, in addition to these mutants' distribution throughout the nuclear interior. This latter finding resembled the attenuated NE association of the corresponding *Hs*ZC3HC1 mutant W107A upon its ectopic expression in HeLa ZC3HC1 KO cells (Supplemental Figure S2B). Furthermore, the residual NE localization of the W119A and Y257A mutants was also in line with these mutants being capable of attenuated Y2H interactions with the Mlps at low 3-AT concentrations (S5B and S5C1). Bar, 5 µm.

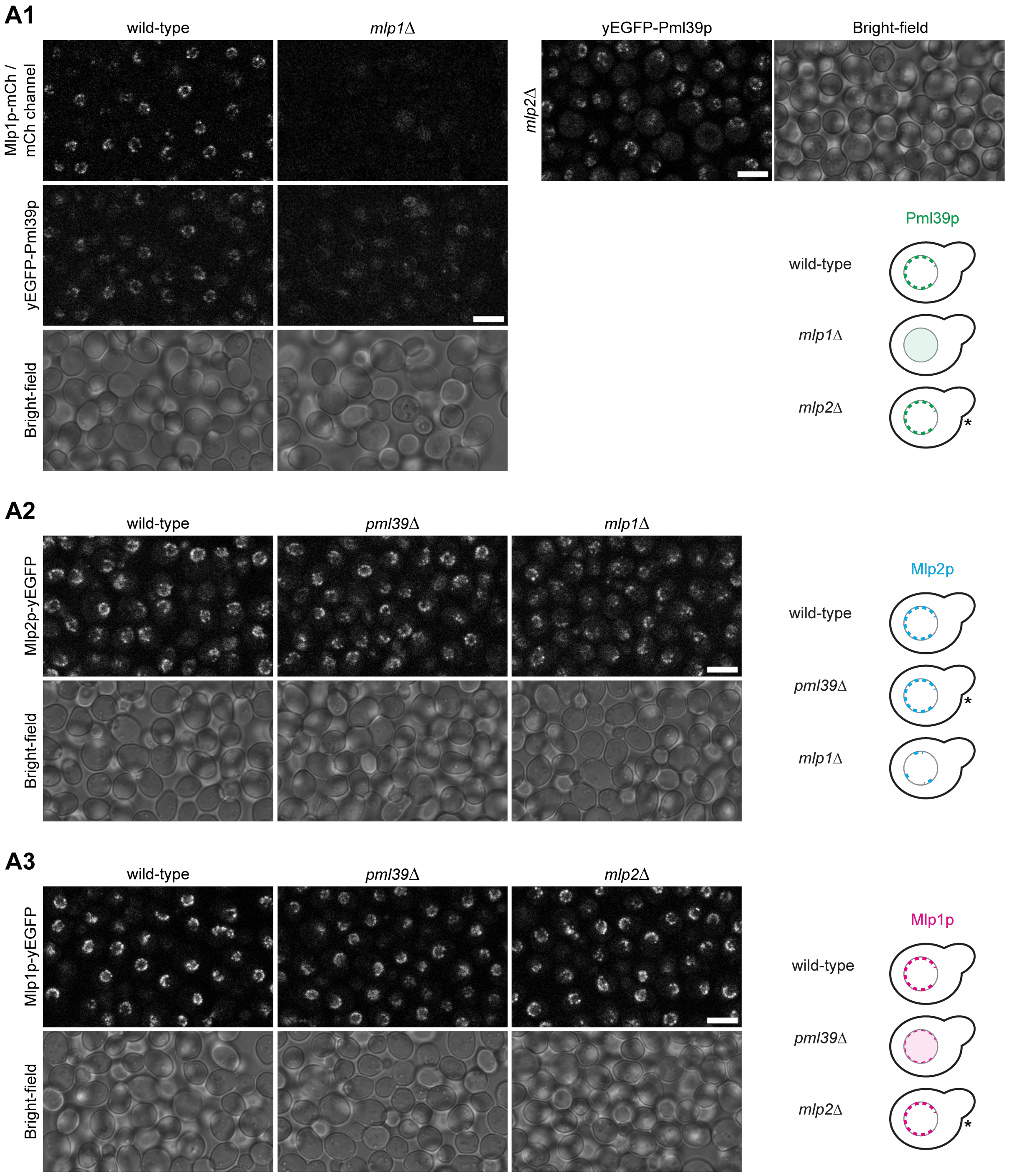
A1
wild-type
mlp1Δ
Mlp1p-mCh / mCh channel
yEGFP-Pml39p
Bright-field
yEGFP-Pml39p
Bright-field
mlp2Δ
Pml39p
wild-type
mlp1Δ
mlp2Δ
*
A2
wild-type
pml39Δ
mlp1Δ
Mlp2p-yEGFP
Bright-field
Mlp2p
wild-type
pml39Δ
mlp1Δ
*
A3
wild-type
pml39Δ
mlp2Δ
Mlp1p-yEGFP
Bright-field
Mlp1p
wild-type
pml39Δ
mlp2Δ
*

**S6 (1/3)**

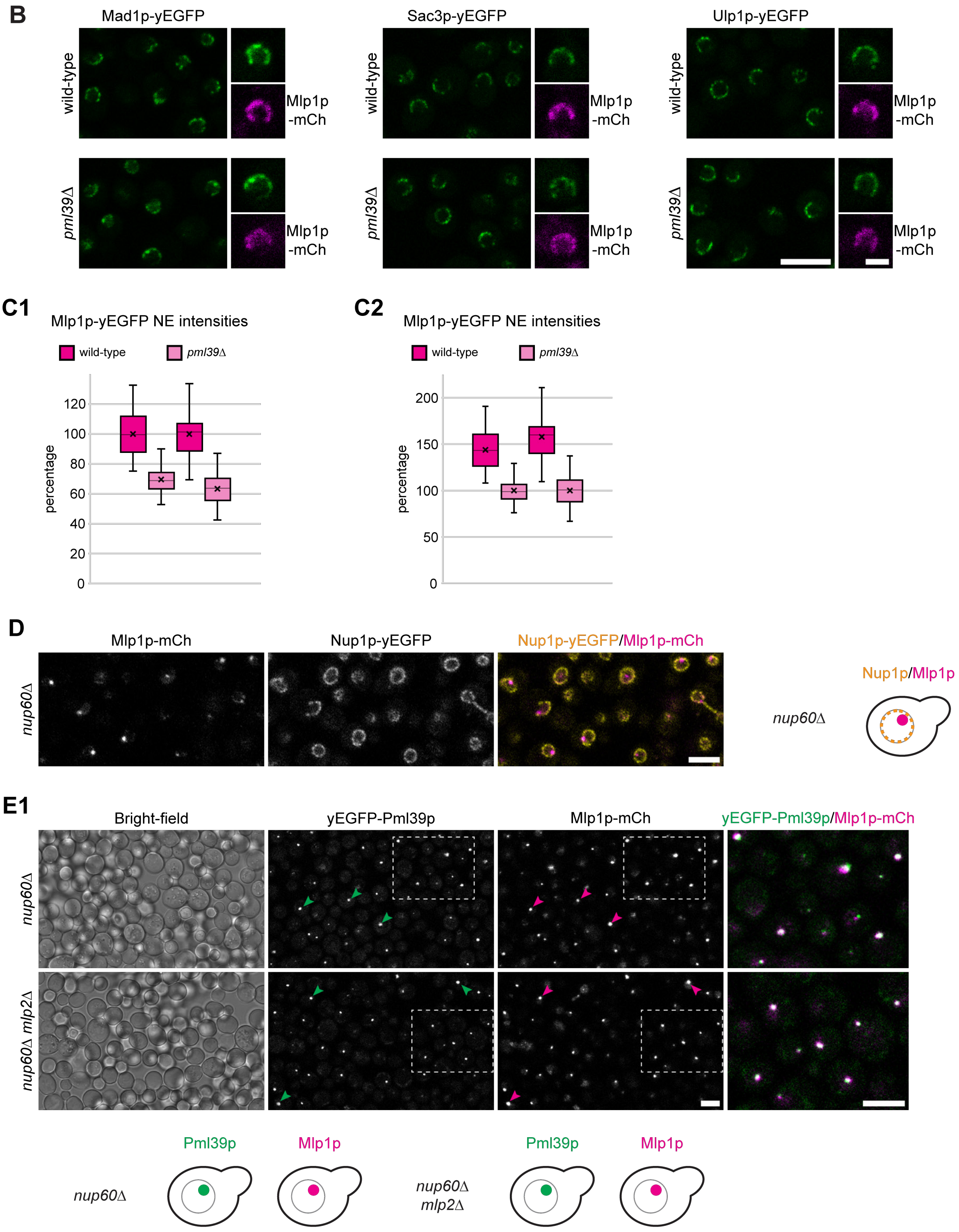
B
Mad1p-yEGFP
Sac3p-yEGFP
Ulp1p-yEGFP
wild-type
pml39Δ
Mlp1p -mCh
C1
Mlp1p-yEGFP NE intensities
wild-type
pml39Δ
percentage
120
100
80
60
40
20
0
C2
Mlp1p-yEGFP NE intensities
wild-type
pml39Δ
percentage
200
150
100
50
0
D
Mlp1p-mCh
Nup1p-yEGFP
Nup1p-yEGFP/Mlp1p-mCh
nup60Δ
Nup1p/Mlp1p
nup60Δ
E1
Bright-field
yEGFP-Pml39p
Mlp1p-mCh
yEGFP-Pml39p/Mlp1p-mCh
nup60Δ
nup60Δ mlp2Δ
Pml39p
Mlp1p
nup60Δ
Pml39p
Mlp1p
nup60Δ mlp2Δ

S6 (2/3)

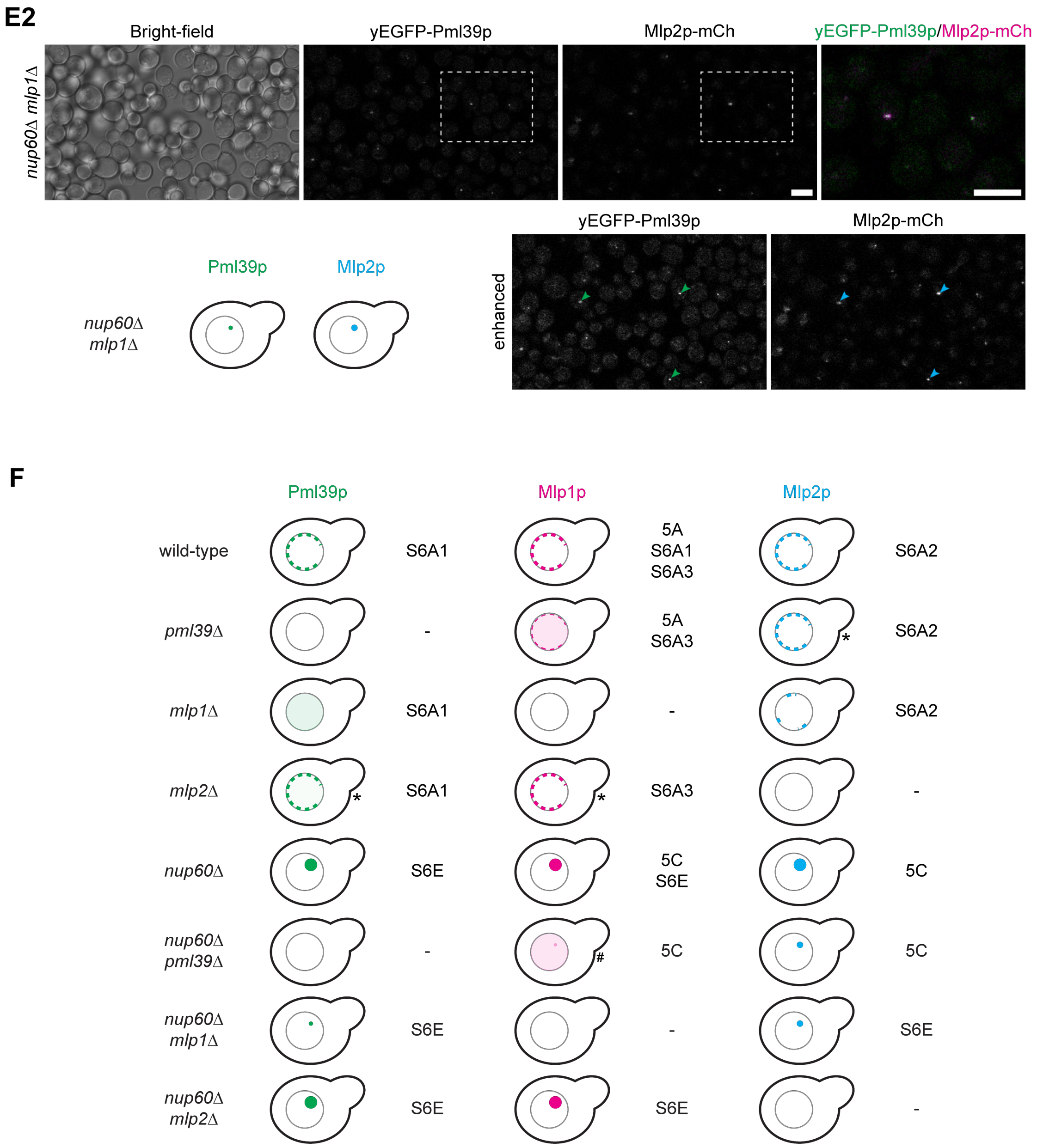

**Supplemental Figure S6. Complementing studies on the contribution of *Sc*Pml39p in keeping subpopulations of NE-associated Mlp1p polypeptides positioned at the NB and within Mlp1p-containing nuclear foci.**

**(A)** Subcellular localization of Mlp1p, Mlp2p, and Pml39p in wild-type and *mlp1*Δ, *mlp2*Δ, and *pml39*Δ cells, with a selection of live-cell images here presenting such cells, endogenously expressing either all Pml39p, Mlp2p, or Mlp1p as yEGFP-tagged polypeptides (S6A1–3) or Mlp1p tagged with mCherrry (S6A1). Bright-field micrographs are shown as a reference. Note that images of *pml39*Δ cells expressing Mlp1p-yEGFP presented in S6A3 and equivalent micrographs in Figure 5A represent different datasets. Further note that images from the different strains presented in S6A2 and S6A3 were acquired on the same day in parallel to each other, using identical microscope settings to allow for adequate comparability of data. The subcellular distribution of the yEGFP-tagged Mlp and Pml39 polypeptides in the wild-type and KO strains is again schematically depicted on the right. The asterisks mark those combinations of KO strains and endogenously expressed FP-tagged proteins for which the results appeared to vary moderately between replicated rounds of inspection. Such ambiguity held for some minor reduction in the amounts of (i) the NE-associated Pml39p in the *mlp2*Δ cells (S6A1), (ii) the NE-associated Mlp2p in the *pml39*Δ cells (S6A2), and (iii) the NE-associated Mlp1p in the *mlp2*Δ cells (S6A3), with us noting such reductions only in some replicates but not others, and with such variations appearing to correlate to some extent with the growth phases of the cell cultures. Further note that not all but most of the results presented in S6A1–3 confirm previously reported findings (Palancade *et al*, 2005).

**(A1)** Live-cell fluorescence microscopy of wild-type and *mlp1*Δ cells, with both strains expressing all Pml39p tagged with yEGFP and Mlp1p tagged with mCherry. In addition, an *mlp2*Δ strain expressing yEGFP-tagged Pml39p is shown. Note that Pml39p and Mlp1p co-localize at the wild-type cells' NEs. Further note that Pml39p was essentially absent from the NEs of the *mlp1*Δ cells and instead primarily distributed diffusely throughout the nuclear interior, next to only some very rarely seen small-sized foci. By contrast, the absence of Mlp2p in *mlp2*Δ cells appeared to come along with only minor reductions in the NE-associated amounts of Pml39p. Bars, 5µm.

**(A2)** Live-cell fluorescence microscopy of wild-type, *pml39*Δ, and *mlp1*Δ cells, with all strains expressing all Mlp2p tagged with yEGFP. Note that while the NE-associated amounts of Mlp2p were notably reduced in *mlp1*Δ cells, the positioning of Mlp2p at the NEs of *pml39*Δ cells appeared barely affected. Bar, 5µm.

**(A3)** Live-cell fluorescence microscopy of wild-type, *pml39*Δ, and *mlp2*Δ cells, with all strains expressing all Mlp1p tagged with yEGFP. Note that the positioning of Mlp1p at the NEs of the *mlp2*Δ cells appeared barely affected. By contrast, in many of the cells within the *pml39*Δ population, a subpopulation of the yEGFP-tagged Mlp1 polypeptides appeared distributed diffusely throughout the nuclear interior, a feature generally not noted within the *PML39wt* cells. Moreover, the amounts of the NE-associated Mlp1p-yEGFP appeared notably reduced. These results, which we regarded as unequivocal, were in line with essentially the same observations we made when inspecting Mlp1p-yEGFP-expressing *PML39wt* and *pml39*Δ cells at higher resolution in a separate experiment (Figure 5A). Bar, 5µm.

**(B)** Live-cell fluorescence microscopy for investigating the cellular localization within *pml39*Δ cells of other proteins known to occur NB-associated in wild-type cells. To study these proteins' fate in the absence of Pml39p, we had created yet further *PML39wt* and *pml39*Δ strains, all endogenously expressing mCherry-tagged Mlp1p and, in addition, either all Mad1p, Sac3p, or Ulp1p as yEGFP-tagged polypeptides. Note that we found the positioning of these other NB-associated proteins at the NE not notably or only very moderately affected in the *pml39*Δ cells compared to their localization at the wild-type cells' NEs. Bars, 5 µm (overviews) and 2 µm (insets), respectively.

**(C)** Quantification of signal yields of yEGFP tagged to Mlp1p at the NEs of *PML39wt* and *pml39*Δ cells. The quantification procedure is described in the legend of Figure 5B. The data presented here represent the individual results of the two separate experiments (n = 50 nuclei per dataset), whose means are presented in Figure 5B. The box plots display the relative signal intensity values, with the arithmetic means marked by x and the standard deviations (SD) provided. In S6C1, those arithmetic means set to 100% relate to the *PML39wt* cells, while in S6C2, representing the same measured values, they relate to the *pml39*Δ cells. While both ways of presentation illustrate that the mean Mlp1p-yEGFP signal yields were found notably reduced at the *pml39*Δ cells' NEs, we regard the presentation in S6C2 as perhaps better reflecting the copy number relationships between the different Mlp1p subpopulations at the NE. In fact, we can imagine that those Mlp1 polypeptides anchored to the NPC independently of Pml39p represent a somewhat more homogeneous population in terms of copy numbers than those additionally appended in the presence of Pml39p.

Along the same line, we find it informative that we generally found the signal yields' mean SD values smaller for the population of *pml39*Δ cells than for the wild-type cells. Again, we regard this finding, too, as in accord with a model in which the Mlp1p polypeptides that occur NPC-anchored independently of Pml39p would reflect an Mlp1p subpopulation with a more defined copy number per NPC. In contrast, the copy numbers of those Mlp1 polypeptides that can be additionally appended to the NB in a Pml39p-dependent manner could, to some extent, be more variable. In other words, we consider those NBs that remain NPC-appended in the absence of Pml39p to be more similar, regarding their Mlp1p copy numbers, both between cells and between individual NBs within the same cell, than in the presence of Pml39p.

**(D)** Recapitulating the formation of Mlp-containing nuclear foci in *nup60*Δ cells (e.g., Feuerbach *et al*, 2002) as the prerequisite for later investigating how Pml39p contributes to such foci's occurrence. Here, the formation of such Mlp1p-containing foci, usually a single one per nucleus, is exemplified by the live-cell images of *nup60*Δ cells endogenously expressing mCherry-tagged Mlp1p, next to the NPC protein Nup1p tagged with yEGFP, the latter shown as a reference protein that remains located at the NE. In the two-color overlay image, note that

Mlp1p within such *nup60*Δ cells appeared only detectable within the nuclear foci, with the foci's subcellular location evident as they were surrounded by NPCs labeled with Nup1p-yEGFP. Bar, 5 µm.

**(E)** Assessing the contribution of Mlp1p and Mlp2p to the formation of the Mlp-containing foci in *nup60*Δ cells.

**(E1)** Live-cell fluorescence microscopy of *nup60*Δ and *nup60*Δ *mlp2*Δ strains endogenously expressing all Pml39p as yEGFP-tagged polypeptides and all Mlp1p as tagged with mCherry. In the overview images, demonstrating the commonness of the Pml39p- and Mlp1p-containing foci within both strains' populations, some of these foci are marked by green- and magenta-colored arrowheads. Furthermore, the images' marked rectangular areas are shown as two-color overlays at higher magnification on the right, revealing quasi co-localization of Pml39p and Mlp1p in these foci known to be located within these cells' nuclei. Note that signal displacement often observable between the two color channels was caused by the non-fixed cells' residual mobility during live-cell imaging, even though the cells had been allowed to settle to the wells' bottom. In particular, though, note that the absence of Mlp2p did not prevent the formation of such foci. Further note that the overview images presented here in S6E1 and the following in S6E2 were acquired from specimens all inspected side-by-side, using the same microscope settings, to allow for direct data comparability. Bars, 5 µm.

**(E2)** Live-cell fluorescence microscopy of a *nup60*Δ *mlp1*Δ strain endogenously expressing all Pml39p as yEGFP-tagged polypeptides and all Mlp2p as tagged with mCherry. The overview images are also shown electronically signal-enhanced, with a multiplication factor of two. Note that hardly any foci, except for a few sporadic ones, were detectable, some of which are marked by green and blue arrowheads, revealing that Pml39p and Mlp2p together are not well capable of forming large foci when Mlp1p is absent. Bars, 5 µm.

**(F)** Summarizing schematic depiction of the subcellular distribution of the FP-tagged Mlp and Pml39 polypeptides in some of the different yeast strains investigated in the current study (for a list of strains, see Supplemental Table S2). In contrast to the majority of strains, for which we rated the observations as unequivocal, the outcome for a few strains, here marked by asterisks, appeared to vary moderately between replicated rounds of inspection, as mentioned in the legends of Figure 5D and Supplemental Figure S6A. Figure numbers of micrographs representing the different experiments are provided on the right side of each yeast cell's scheme. Based on the results *in toto*, we regarded it justified to conclude that Pml39p can keep subpopulations of Mlp1p positioned at the NB and even at sites remote from the NB, pointing at Pml39p functioning as a protein connecting Mlp1p polypeptides.

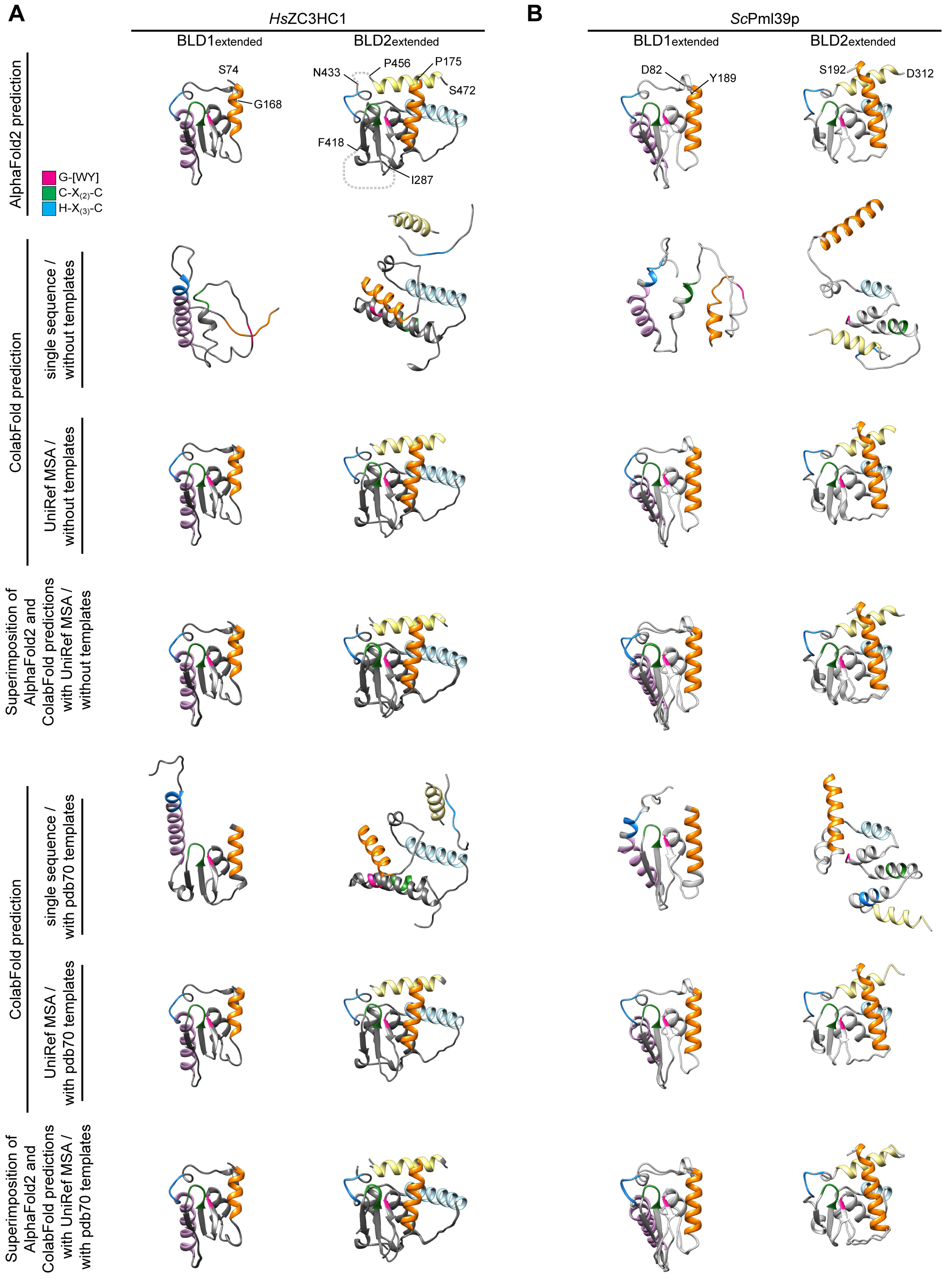
A
HsZC3HC1
BLD1extended
BLD2extended
B
ScPml39p
BLD1extended
BLD2extended
AlphaFold2 prediction
S74
G168
N433
P456
P175
S472
F418
I287
D82
Y189
S192
D312
G-[WY]
C-X(2)-C
H-X(3)-C
ColabFold prediction
single sequence / without templates
UniRef MSA / without templates
Superimposition of AlphaFold2 and ColabFold predictions with UniRef MSA / without templates
ColabFold prediction
single sequence / with pdb70 templates
UniRef MSA / with pdb70 templates
Superimposition of AlphaFold2 and ColabFold predictions with UniRef MSA / with pdb70 templates

**S7 (1/2)**

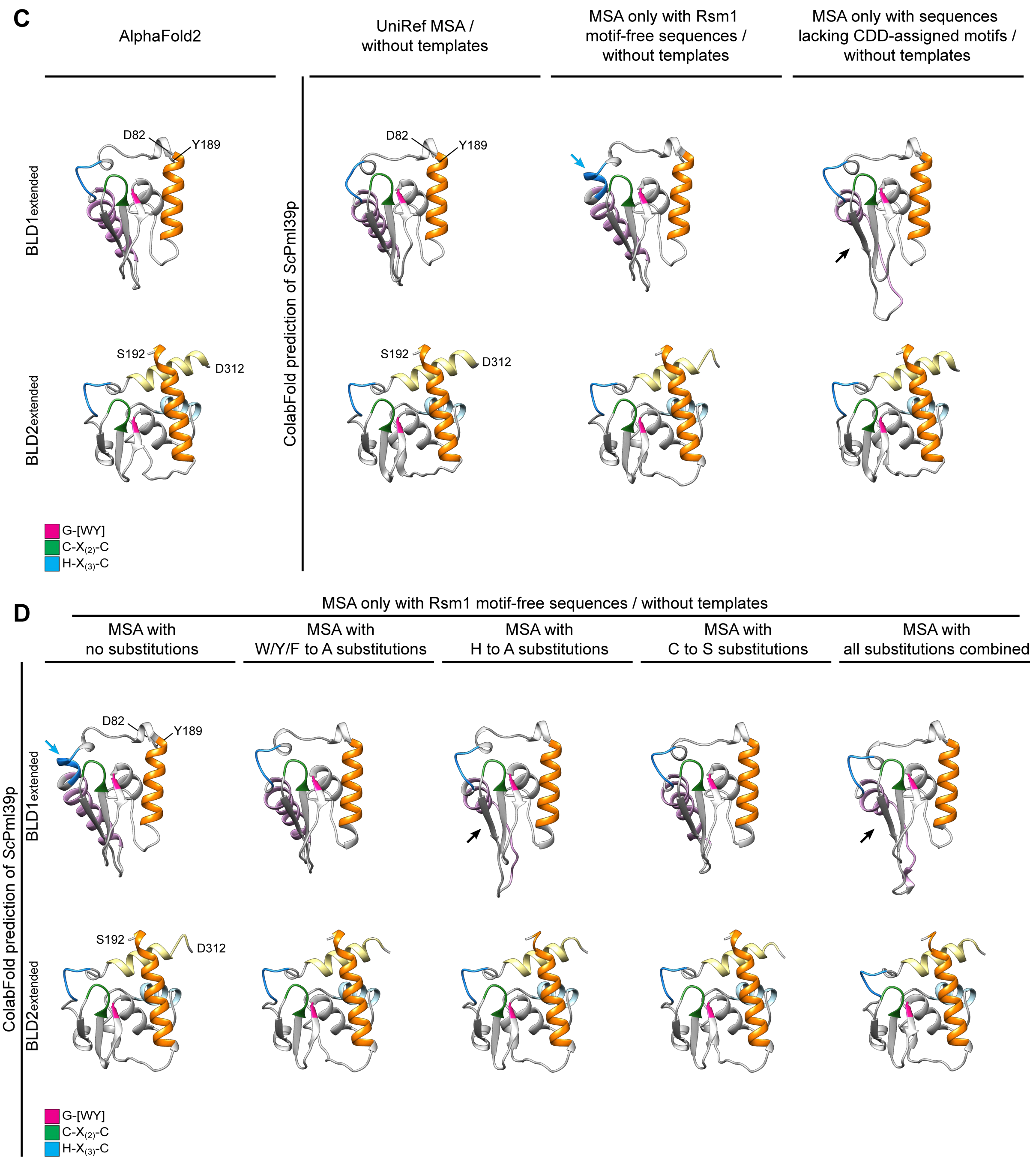

**Supplemental Figure S7. Assessment of the contributions of sequence database search-derived MSAs and PDB database-deposited structures as templates for BLD structure predictions.**

Since AlphaFold2 uses, as one of the input datasets for predicting a query protein's structure, also PDB-deposited crystal structures as templates in its computations (Jumper *et al*, 2021), we had wondered whether such templates might bias the predictions of the BLDs' structures (Supplemental Information 3). Furthermore, we wanted to assess to which extent distinct sequences that were part of the MSAs, the latter representing the other input dataset for such structure predictions, might influence the outcome of the predictions by introducing some bias. In particular, we were interested to find out whether structure predictions starting from an MSA composed of protein sequences with a Pfam Rsm1 motif already assigned to them would differ from predictions based on an MSA solely composed of sequences without such a pre-existing profile HMM while nonetheless representing ZC3HC1 homologs according to our criteria, which included possession of a NuBaID signature (Supplemental Information 3). Since an Rsm1 motif has not been attributed to *Sc*Pml39p so far, and as this happens to be the case for numerous other proteins too that we consider true ZC3HC1 homologs, including such in other species of the family Saccharomycetaceae, we wondered which structures would be predictable starting from such a "Pfam Rsm1 motif-free" MSA. However, we also noted that tools like HHsearch and HHpred (Söding, 2005; Steinegger *et al*, 2019; https://toolkit.tuebingen.mpg.de/tools/hhpred) would assign the PDB structures and profile HMM of metazoan BIR domains to such a subset of Pfam motif-free yeast sequences nonetheless. Therefore, we also attempted to mutate the input sequences by *in-silico*-exchanging a small number of residues so that the BIR domains' HMM would no longer be correlatable with the mutated yeast sequences, with this latter approach overlapping with our additional attempt to gain insight also into how AlphaFold2 handles an MSA only composed of mutated sequences (see further below). With such approaches, we aimed to impede the program's access to HMM and PDB template information by which it might correlate *Sc*Pml39p with *Hs*ZC3HC1 or the BIR domains during a user-initiated prediction process, unlike in AlphaFold2's training procedures during which such HMM and crystal structure information had been used (Jumper *et al*, 2021).

In addition, we addressed some of our concerns about the structure predictions of mutant versions of ZC3HC1 (i) that naturally occur *in vivo*, those (ii) that we had constructed and cloned in the course of the current study, and yet additional ones (iii) that we had designed *in silico* in order to test certain aspects of the program's performance. In Supplemental Information 3, the rationale for conducting such trial structure predictions is described in further detail, while exemplary data sets in the following illustrate the outcome of such trials.

**(A)** Structures predicted for the essentially entire BLD1 and BLD2 modules of *Hs*ZC3HC1, as defined as the "extended" version in Figure 6C. Some of the ColabFold-predicted structures were then superimposed onto those from AlphaFold2. The MMseqs2-derived UniRef MSA used for some of these predictions comprised 799 sequences from various organisms. Note that predictions based on only the protein's own sequence did not allow for any prediction closely resembling the initial prediction by AlphaFold2 or via ColabFold's default settings (UniRef MSA and without template). Further note, in particular, that including PDB template information did not notably affect the outcome of the ColabFold predictions starting from the UniRef MSA, with such template-considering predictions closely resembling those achieved with the programs' default, "template-free" settings. The additional information provided by the PDB templates only improved the outcome to some extent, though merely for the BLD1, when using only the *Hs*ZC3HC1 sequence as the sequence input.

**(B)** Structures predicted, as in S7A, for the essentially entire BLD1 and BLD2 modules of *Sc*Pml39p. Some of the ColabFold-predicted structures were again superimposed onto those from AlphaFold2. The UniRef MSA used for some of the here-presented predictions comprised a total of 27 fungal sequences, including two with an Rsm1 motif attributed to them. A zf-C3HC motif had been assigned to 26 of them, as in Pfam release 35.0, while in the CDD, this was the case for only 13 of them. Note that predictions based on the *Sc*Pml39p sequence alone, i.e., without employing an MSA, notably differed from the initial one provided by AlphaFold2. Again, however, just like for *Hs*ZC3HC1, note that the exclusion of PDB template information did not notably affect the outcome of those predictions that only made use of information provided by the UniRef MSA.

**(C)** ColabFold-predicted structures of the essentially entire BLD1 and BLD2 modules of *Sc*Pml39p, all obtained without accessing the PDB's template information, next to those predicted with AlphaFold2s' default settings. The UniRef MSA used by ColabFold for the structures in the second column, as already presented in S7B, comprised the total of again 27 sequences, among which was also a sequence that would represent a BLD1-mutated version of a Pml39p homologue (XP_037145256.1) and two short segments only comprising part of the respective homolog's BLD1 sequence. Removing these three sequences together with the remaining one with an Rsm1 motif assigned to it resulted in an MSA comprising 23 Rsm1-motif-free sequences, representing ZC3HC1 homologs with an intact BLD1. The BLD1 and BLD2 structures of *Sc*Pml39p computed on the basis of this MSA are presented in the third column. The forth column then represents the structures predicted after having additionally removed all those sequences from the MSA to which a zf-C3HC motif had been assigned in the CDD. Aware that the remaining sequences still had a zf-C3HC attributed to them by Pfam, we nonetheless regarded also this structure prediction as noteworthy, for reasons outlined below.

We already regard, though, the BLD2 structure presented in the third column as of particular note. Even without any template information and no Rsm1 profile HMM appertaining to the input sequences, this ColabFold-predicted BLD2 still closely resembled the initial one based on the UniRef MSA, with this again essentially identical to those from the AlphaFold2 structure database. However, one also needs to know that a program like HHpred (Zimmermann *et al*,

2018; https://toolkit.tuebingen.mpg.de/tools/hhpred), even though it, back then, did not detect ZC3HC1 homologs of other phyla when using the Rsm1-motif-free yeast sequences, would still correlate the MSA composed of these sequences with the PDB-deposited structure and HMM profile of BIR domains. In other words, some profile comparison tools using such Rsm1-motif-free yeast sequences would still detect similarity with the BIR domains' profile even when the closer kinship of these yeast proteins remained undetectable. In this context, we also regard it as noteworthy that the removal of the Rsm1 motif-containing sequences resulted in a BLD1 prediction, which then, in some part, more closely resembled a BIR domain: In fact, the pink-colored α-helix of the BLD1 was then predicted C-terminally extended (blue arrow), with the histidine of the first BLD's H-$X_{(3)}$-C pentapeptide then part of this α-helix, just like the histidine of the BIR domains' H-$X_{(6)}$-C octapeptide is part of an α-helix similarly positioned relative to the domain's centrally positioned β-sheets.

Furthermore, we regard it as noteworthy that a total of only 13 remaining sequences had turned out sufficient to predict a structure whose appearance was still very similar to the one predicted when starting from the UniRef MSA, with about twice as many sequences. Merely the prediction of the pink-colored α-helix of BLD1, then no longer predicted as such in its entirety, differed notably (here marked by a black arrow). Initially, we had been uncertain whether such a small number of very similar sequences, all from the same family Saccharomycetaceae, would provide a bandwidth of non-conserved residues high enough to accentuate those evolutionarily conserved residues that were the structurally most relevant ones. In other words, we initially were doubtful whether such a small dataset would suffice because Alphafold2's prediction accuracy had been mentioned to decrease notably when an MSA was composed of less than 30 sequences (Jumper *et al*, 2021).

**(D)** ColabFold-predicted structures of wild-type and mutant versions of *Sc*Pml39p, all obtained without accessing the PDB's template information. The MSA for predicting the wild-type structure on the left, as already presented in S7C, again comprised the total of 23 yeast sequences that we regarded as representing functionally intact ZC3HC1 homologs, including *Sc*Pml39p, while all of them lacked an Rsm1 signature as defined by the Pfam database. For predicting the structure of various mutant versions of *Sc*Pml39p, we exchanged distinct residues both within the *Sc*Pml39p input sequence and at the corresponding positions of the other 22 sequences for either alanine or serine, with these MSAs and their modified sequences to be then used for the mutant *Sc*Pml39p's structure prediction. The residues exchanged corresponded, in the case of *Sc*Pml39p, to H172A and H288A of the two BLDs' H-$X_{(3)}$-C pentapeptides (H to A), and to W119A and Y257A of the G-W and G-Y dipeptides, with one of the other yeast sequences having featured a G-F at the corresponding position instead, which was changed accordingly (W/Y/F to A). Furthermore, for another mutant *in silico*, all cysteine residues of the two BLDs' C-$X_{(2)}$-C tetrapeptides and H-$X_{(3)}$-C pentapeptides involved in zinc coordination were exchanged for serine, resulting in C134S, C137S, C176S, C268S, C272S, and C292S (C to S). Finally, all these mutations were combined (W/Y/F to A, C to S, and H to A). It needs to be remarked that upon introducing the W/Y/F to A mutations or the H to A mutations, the high sensitivity tool HHpred would still correlate these MSAs with BIR domain structures. However, this was no longer the case for the MSA representing the C to S mutations when initially using HHpred, i.e., before the HHpred web server updates in 2022, and the same applied to the one whose sequences combined all the substitutions. Furthermore, while other tools like HHblits (Remmert *et al*, 2011) and HMMER3 (Eddy, 2011), used by AlphaFold2 too (Jumper *et al*, 2021), would still allow for identifying a few fungal wild-type ZC3HC1 homologs when using some of the mutant sequences as query, no homologs were immediately identifiable beyond the Mycota kingdom during first-round searches. The latter also held when conducting such searches with the other search tools mentioned in the current study.

The findings outlined in S7D conveyed to us the following impressions. First, with the *Sc*Pml39p sequence part of a subtracted MSA, which at first appeared no longer alignable with BIR domain structures when we had initially conducted such computations, and with such an MSA neither then nor currently allowing for direct detection of ZC3HC1 homologs beyond the fungi, we initially considered it reassuringly remarkable that the *Sc*Pml39p BLD structures predicted on these terms were still strikingly similar to those of *Hs*ZC3HC1. Second, we were surprised, though, that after having introduced aa substitutions in all 23 sequences that would have abolished the establishment of a zinc ion coordination sphere, the AlphaFold2/ColabFold-predicted structures for these mutants appeared only very moderately affected. We regarded this observation as particularly noteworthy since we already knew that each individual substitution mutation, already on its own, abolished or severely attenuated *Sc*Pml39p binding to the NB (this study), with at least some of them also known to prevent proper folding of *Hs*ZC3HC1 (Gunkel & Cordes, 2022, and our unpublished data). Even though we were aware that AlphaFold2 had been mentioned not to be trained for predicting the effect of substitution mutations within a query sequence, we were nonetheless amazed that even manifoldly mutating all the sequences of an MSA would hardly affect the outcome of a prediction process. We considered it enigmatic that even in the case of those BLD1 and BLD2 mutants that each included five aa substitutions, the program still predicted the three centrally positioned conspicuous β-sheets and those positions usually occupied by residues involved in establishing the zinc ion coordination sphere to remain essentially unchanged. With these central arrangements predicted unaffected, the only finding no longer a surprise to us was that the positioning of the BLDs' α-helices relative to these central structures was then also predicted to have remained unchanged, with this in contrast, too, to what we would expect based on our experimental data. Therefore, in light of such trial predictions of the kind exemplified here and the insight they had allowed us to gain, we considered it justified to exert some caution when interpreting some of the structure predictions by AlphaFold2; to avoid misinterpretations and not draw conclusions from computed structures that this prediction tool had not been trained for providing.

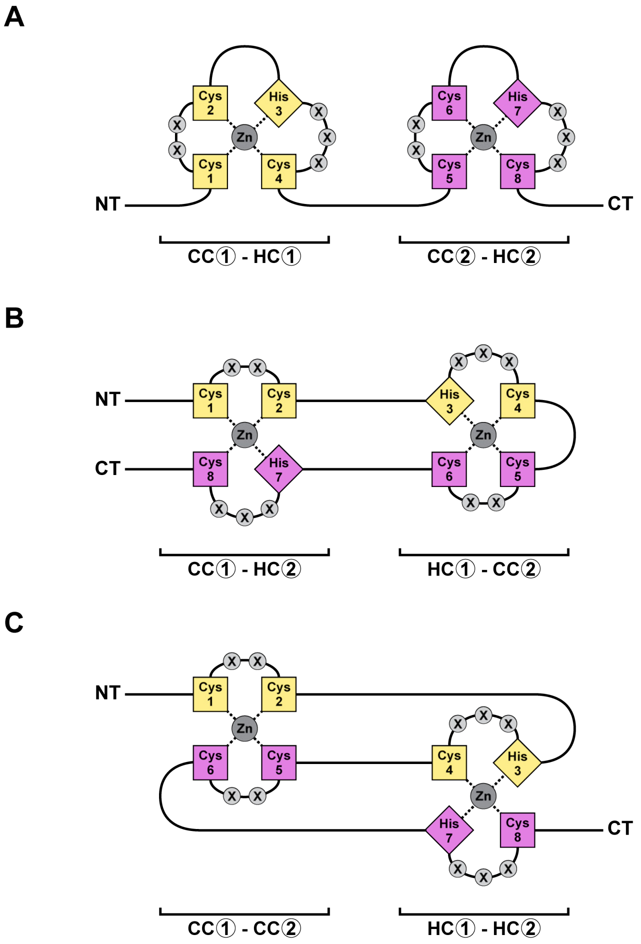

**Supplemental Figure S8. Former considerations regarding the zinc ion-coordination topology of ZC3HC1.**

Speculating in the early stages of our research on ZC3HC1 how the protein's two pairs each of C-$X_{(2)}$-C tetrapeptides and H-$X_{(3)}$-C pentapeptides are arranged relative to each other, each in order to coordinate a zinc ion likely in a tetrahedral manner, we had initially considered different constellations hypothetically conceivable. Even though the NuBaID signature early on suggested an arrangement of two zinc finger modules one after the other, also paraphrased as CC①-HC① and CC②-HC② in the following and schematically depicted as such in S8A, and even though this appeared in line with the type of zinc ion coordination sphere already proposed based on the BLD's similarity with the BIR domains (Higashi *et al*, 2005), we at times had not yet regarded it as justified to exclude other scenarios categorically. Such mind games also took place in light of the variety of arrangements within different proteins that by then were known to allow for binding zinc ions in a tetrahedral geometry.

For example, configurations with pairs of zinc ion-coordinating residues belonging to different parts of the primary sequence separated from each other by other pairs of residues involved in coordinating yet another zinc ion had been described for some zinc fingers of the RING/FYVE/PHD-types (e.g.,Capili *et al*, 2001; Legge *et al*, 2004; Houben *et al*, 2005; Gamsjaeger et al., 2007; Kandias *et al*, 2009; Deshaies & Joazeiro, 2009; Wei & Sun, 2010) and for some other types of zinc fingers too (e.g., He *et al*, 2007; Massiah *et al*, 2007). These proteins were characterized by the especially pronounced winding of their aa chains to achieve an interleaved arrangement of the zinc ion-coordinating residues, also referred to as a cross-braced configuration.

Furthermore, while the coordination of a zinc ion by two cysteines and two histidines was known to commonly occur in a non-interleaved manner in numerous zinc finger proteins harboring a C2H2 zinc finger domain (e.g., Brayer & Segal, 2008), zinc ion coordination involving two histidines had also been found occurring within cross-braced arrangements, for example in some B-box zinc finger proteins (e.g., Massiah *et al*, 2006; He *et al*, 2007).

If such latter coordination topology had applied for ZC3HC1, the *Hs*ZC3HC1 protein's two pairs of C-$X_{(2)}$-C tetrapeptides and H-$X_{(3)}$-C pentapeptides would have had to allow for an interleaved CC①-CC② and HC①-HC② configuration, as schematically depicted in S8C. Furthermore, we also did not want to discard yet another scenario too early in which the protein would adopt a clothespin-like CC①-HC② and CC②-HC① configuration, as outlined in S8B.

In principle, our finding that eliminating a single one of the BLDs' zinc-coordinating residues had been sufficient for abolishing TPR-binding would have been reconcilable with all three scenarios. Thus, without crystallographic data for ZC3HC1 at hand, we had initially conceived it inappropriate to take the CC①-HC①_CC②-HC② configuration for granted and already rule out the other scenarios, even though sequence similarities between the BLDs and the related BIR domains, for which crystal structures were already available (e.g., Supplemental Figure S10), were pointing attractively towards the CC①-HC①_CC②-HC② version.

Eventually, then, with AlphaFold2 predicting such a configuration too (Figure 6, Supplemental Figure S9), not only for *Hs*ZC3HC1 but also for other homologs with which the human one barely shared any aa sequence identity, like *Sc*Pml39p (see in this context also Supplemental Information 3 and Supplemental Figure S7), we regarded it as reasonable to treat the CC①-HC①_CC②-HC② configuration as the one genuinely existing in all probability. The finding of an evolutionarily conserved BLD1:BLD2 binding interface (Supplemental Figure S9B) contributed to this conclusion, with the latter, in turn, based on the following considerations: On the one hand, regarding the similarity of the BLDs' predicted central structures with the BIR domains' crystal structures, we had initially considered it appropriate to interpret such similarities with caution, as reasoned further above (e.g., Supplemental

Information 3). On the other hand, though, the BIR domains' AlphaFold2-predicted arrangements relative to each other, for example, in the four human proteins that contain more than one BIR domain (BIRC1 to BIRC4), appeared dissimilar from the BLD1:BLD2 interface of the three here presented ZC3HC1 homologs. This, then again, meant that the three homologs' BLD1:BLD2 interface similarities were less likely to have been influenced by some structural similarity with the BIR domains and thus by an initially considered "discussible level of bias" affecting the BLD structure predictions (Supplemental Information 3).

Nonetheless, even though all current evidence argues for the NuBaID's CC①-HC①_CC②-HC② configuration, we still consider it adequate to illustrate the other initially imagined constellations for comparison in the following. The tetrahedrally coordinated zinc ions are thereby presented as dark grey spheres in these schematic depictions. The zinc ion-coordinating residues of the first and second BLD are depicted as squares and rhombuses, representing cysteine and histidine residues, with those corresponding to the first BLD colored in yellow and those of the second in purple. In addition, the numbers of 1–8 displayed within these quadrilaterals correspond to the total of eight zinc ion-coordinating residues of the two BLDs and reflect their order relative to each other along the protein's linear sequence of amino acids. **(A)** Depiction of the two BLDs of ZC3HC1 as two separate zinc-binding modules consecutively arranged one after the other in tandem. The coordinating residues of the first and second BLD conform with the order CC①-HC①_CC②-HC② and the arithmetic sequence 1,2,3,4,5,6,7,8. Note that this arrangement is in line with AlphaFold2's structure predictions for ZC3HC1 homologs like *Hs*ZC3HC1 (https://alphafold.ebi.ac.uk/entry/Q86WB0) and *Sc*Pml39p (https://alphafold.ebi.ac.uk/entry/Q03760). As such, they closely resemble the consecutive arrangements of zinc fingers that are part of the BIR domain-containing proteins, the latter compared with the BLDs' structures in Supplemental Figure S10, with the ZC3HC1 homologs' BLDs though exhibiting an evolutionarily conserved BLD1:BLD2 interface (Supplemental Figure S9B). By contrast, in the single BIR domain-containing human proteins BIRC5 to BIRC8, an equivalent BIR:BIR interface is self-evidently absent, while in the human BIRC1 to BIRC4 (https://alphafold.ebi.ac.uk/entry/Q13075; https://alphafold.ebi.ac.uk/entry/Q13490; https://alphafold.ebi.ac.uk/entry/Q13489; https://alphafold.ebi.ac.uk/entry/P98170), with three BIR domains each, their arrangements relative to each other appear notably different (see also further comments in the legend to Supplemental Figure S9B).

**(B)** Depiction of a zipper-like arrangement that could have arisen when the second half of the ZC3HC1 protein, harboring the second BLD, would have folded back onto itself. Such constellation could then have allowed positioning each of the one BLD's two pairs of zinc ion-coordinating residues face-to-face with each of the other BLD's two pairs, resulting in a CC①-HC②_CC②-HC① arrangement and the number order 1,2,7,8,3,4,5,6. With this possibly having positioned the protein's N- and C-terminal parts next to each other, such a constellation might have created the impression also of a clothespin-like arrangement.

**(C)** Depiction of an interleaved, cross-braced type of arrangement, which would have required marked winding of the aa chain to achieve such a conformation. In contrast to the scenarios in S8A and S8B, this model of a CC①-CC②_HC①-HC② arrangement, equivalent to a number order of 1,2,5,6,3,4,7,8, would have demanded that the first zinc ion would have been coordinated only by cysteine residues while both histidine residues would have been involved in the coordination of the second zinc ion.

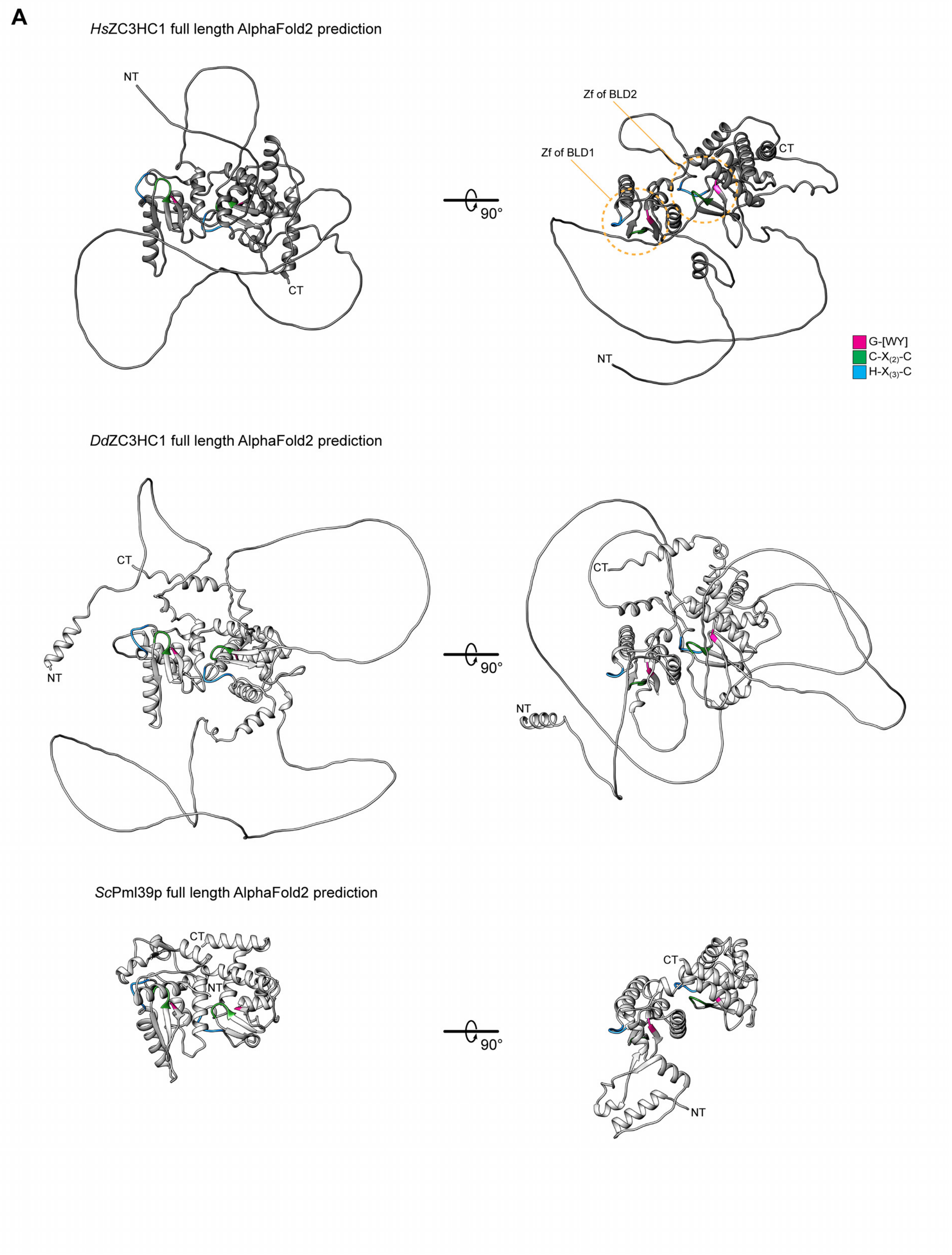
A
HsZC3HC1 full length AlphaFold2 prediction
NT
CT
90°
Zf of BLD2
Zf of BLD1
CT
NT
G-[WY]
C-X(2)-C
H-X(3)-C
DdZC3HC1 full length AlphaFold2 prediction
CT
NT
90°
CT
NT
ScPml39p full length AlphaFold2 prediction
CT
NT
90°
CT
NT

**S9 (1/3)**

**B1**

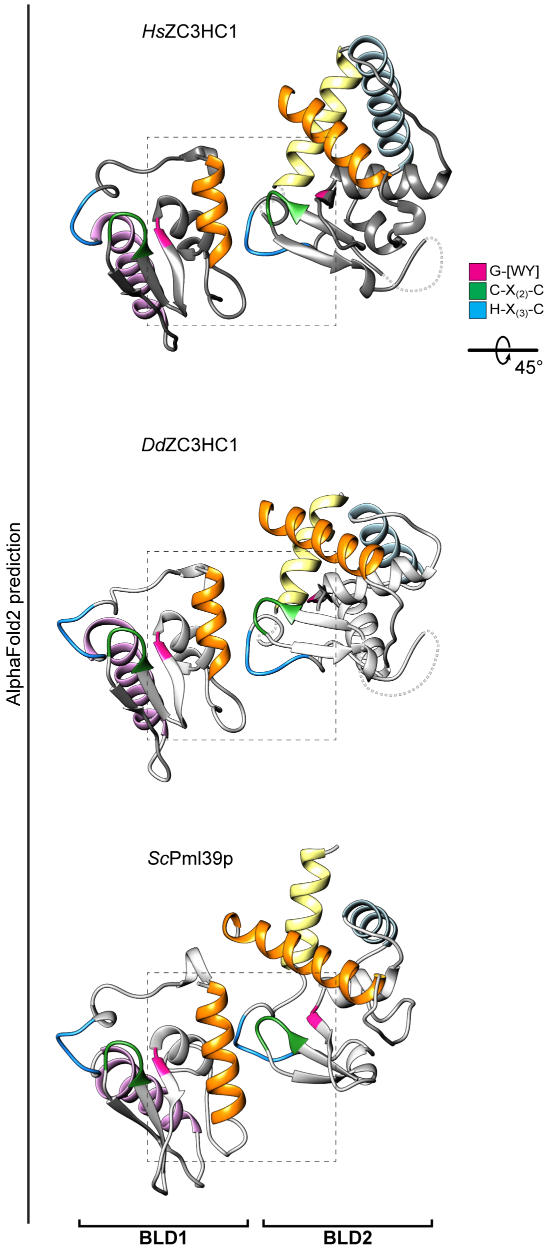

**B2**

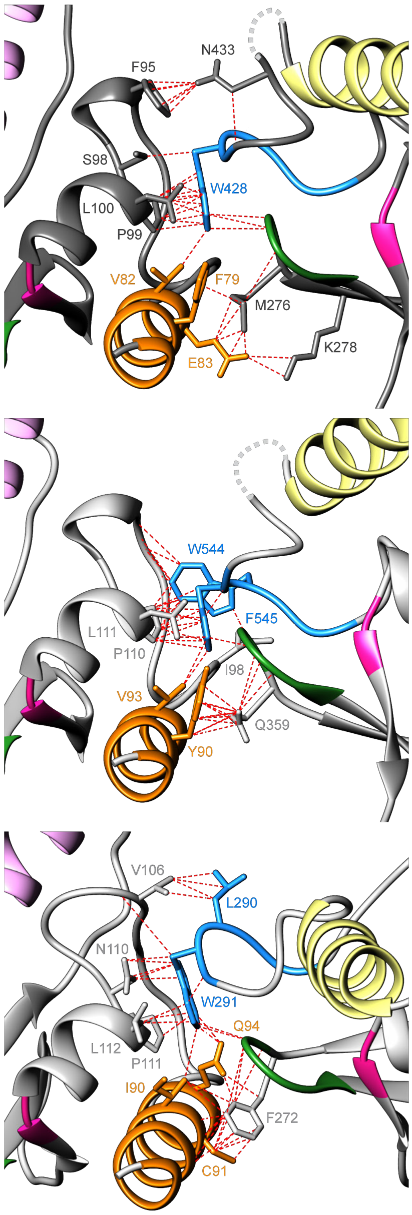

| BLD1 | / BLD2 |
|---|---|
| E83 | - M276 |
| E83 | - K278 |
| F79 | - K278 |
| V82 | - W428 |
| F95 | - N433 |
| S98 | - W428 |
| P99 | - W428 |
| L100 | - W428 |

| BLD1 | / BLD2 |
|---|---|
| Y90 | - Q359 |
| V93 | - F545 |
| I98 | - W544 |
| P110 | - F545 |
| L111 | - F545 |

| BLD1 | / BLD2 |
|---|---|
| I90 | - F272 |
| I90 | - W291 |
| C91 | - F272 |
| Q94 | - F272 |
| Q94 | - W291 |
| V106 | - L290 |
| N110 | - W291 |
| P111 | - W291 |
| L112 | - W291 |

**B3**

```
HsZC3HC1_BLD1  74  SKEAFFSRVETFSSLKWAGKPFELSPLVCAKYGWVTVE------CDMLKCSSCQAFLCASLQPAFDFDRYKQRCAELK--KA-LCTAHEKFCFWPDS 161
DdZC3HC1_BLD1  85  SNTDYYNRVRTYTISNWFAKPCEIDPLQCSRFGWINCE------ADMLECETCKKRLYYKVPSTFSQSLVNKRINDFS--ISLQSTGHRDNCPWKDN 173
ScPml39p_BLD1  82  DLRALLKRICSIQNYTR(7)VRWVNPLTLASKGWEPYQSASQSQVPF-KCCCCHAIMTIPLLKNGDDVADYTMKLNEKIWNSNIIGNHLQKCPWREN 181

HsZC3HC1_BLD2 175  PAILVSEFLDRFQSLCHLDL(52)TACILSVCGWACSSSLESMQLSLITCSQCMRKVGLWGFQQIE--(123)-DTSSRSFFDPTSQHRDWCPWVNI 434
DdZC3HC1_BLD2 187  FQTQLEAYIKRSQNIYNNLT(49)VSCLLALCGWDFNSI(79)-KSSVYCSYCQRLCGVWNFNKIK--(157)-ATEKKKEFSPINEHRWFCPWMIV 551
ScPml39p_BLD2 192  SSQNLIREIERIHTEIDRIV(36)SLVGLLLLGYTKF-----QKDDLVQCTACFHRA--------------------SLKKLEYTEFNGHALWCRYYNK 297
                                                   *:               *  *                                  *   *:
                                                   GW               C  C                                  H   CW
                                                    Y                                                         Y
```

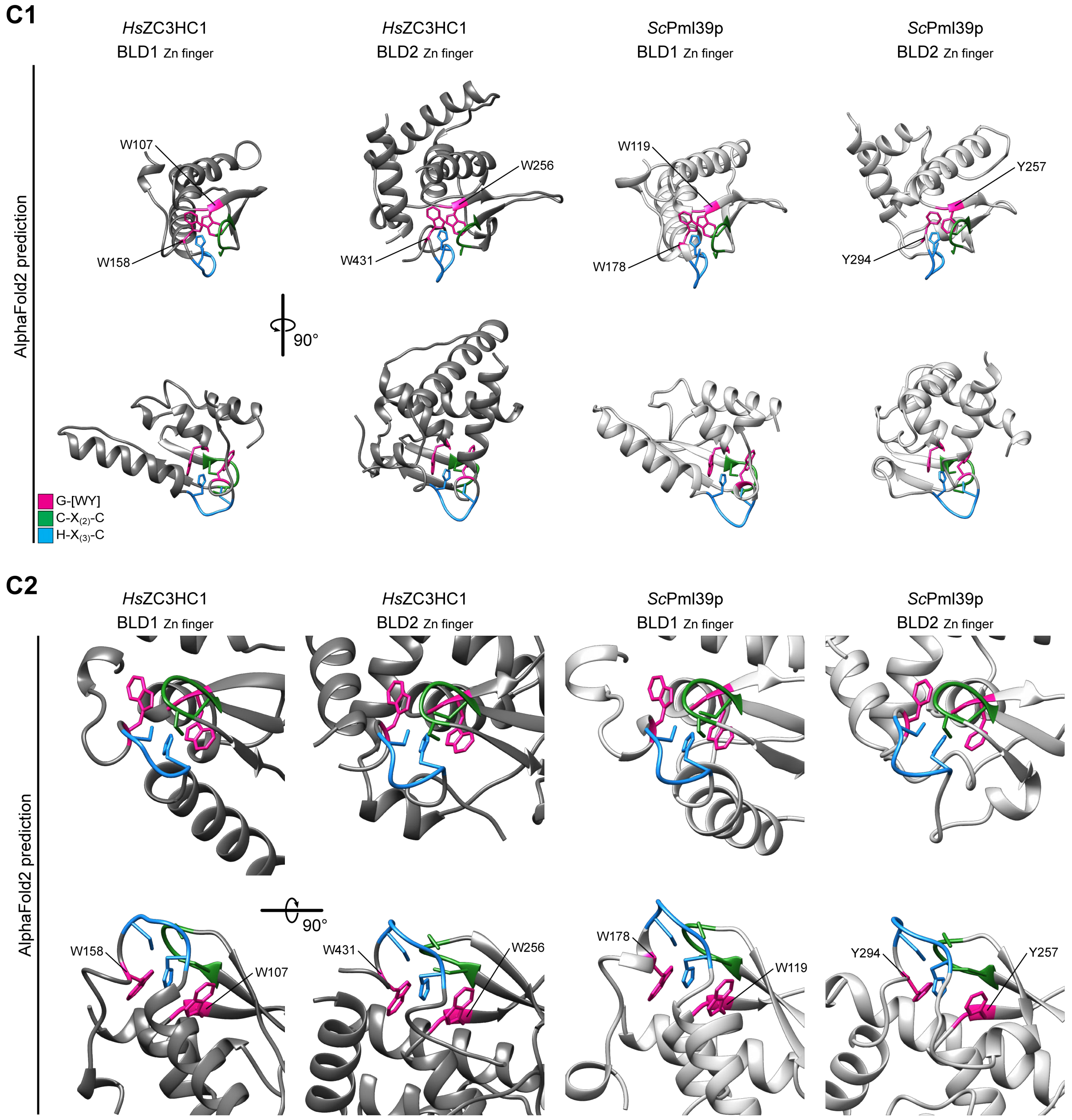

**Supplemental Figure S9. The tertiary structures of *Hs*ZC3HC1, *Dd*ZC3HC1, and *Sc*Pml39p *in toto*, as predicted by AlphaFold2, and closer looks at a BLD1:BLD2 interface and at distinct aromatic amino acids flanking the zinc ion coordination spheres.**

To allow for a comparison of the, at first sight, notably different appearance of the human, amoebic and budding yeast homolog of ZC3HC1, AlphaFold2's predictions for these proteins in their full length are here presented in S9A. In addition, these predictions unveiled an evolutionarily conserved BLD1:BLD2 binding interface with distinct inter-BLD contacts, as shown in S9B, and conspicuous intra-BLD arrangements, like, for example, those of the evolutionarily conserved aromatic acids W107, W158, W256, and W431 of *Hs*ZC3HC1 and

W119, W178, Y257, and Y294 of *Sc*Pml39p. The latter residues' positions relative to the zinc ion coordination spheres are outlined in S9C.

**(A)** AlphaFold2's predictions for *Hs*ZC3HC1, *Dd*ZC3HC1 and *Sc*Pml39p, with each homolog's predicted structure aligned to those of the other via each homolog's first BLD and there, in particular, to the zinc coordination sphere and the neighboring β-sheets. In addition, the structures are shown in two different perspectives, allowing for also visualizing additional α-helices, seemingly also evolutionarily conserved, beyond the BLDs' boundaries, as newly defined in this study further below. The functions of some of these non-BLD α-helices, a research topic beyond the scope of the current study, will be presented elsewhere.

**(B)** Structure and sequence characteristics of the BLD1:BLD2 interface of *Hs*ZC3HC1, *Dd*ZC3HC1, and *Sc*Pml39p.

**(B1)** AlphaFold2's predictions for *Hs*ZC3HC1, *Dd*ZC3HC1, and *Sc*Pml39p as in S9A, with the BLDs' positions relative to each other remaining the same, but here having blinded out all those parts not regarded as belonging to the BLDs central parts, the latter essentially as defined in Figure 6C. Also in contrast to S9A, the BLDs are here shown colored, corresponding to their coloring in Figure 6C. The dashed squares mark those parts shown at higher magnification in S9B2.

**(B2)** BLD1 and BLD2 residues of the BLD1:BLD2 interface are shown at higher magnification, with side chains and structures colored as in S9B1. Chimera's structural analysis tool was used to compute and illustrate potential contacts of designated atoms of selected aa side chains with neighboring aa residues. Several possible contacts, on the basis of atom-to-atom distances of ≤ 4 Å, are here depicted as dashed red lines. Note that among the residues most likely involved in such inter-BLD contacts were aromatic ones positioned between the histidine and cysteine of the H-$X_{(3)}$-C pentapeptide of BLD2, with these then in contact with a distinct P-L dipeptide as part of the three ZC3HC1 homologs' BLD1. In the case of, e.g., *Hs*ZC3HC1, the corresponding aromatic residue of BLD2 is W428, while the L-P dipeptide of BLD1 is composed of P99 and L100. Corresponding residues are also present in *Dd*ZC3HC1 and *Sc*Pml39p, where AlphaFold2 also predicts them to contribute to a BLD1:BLD2 interface, with the corresponding residues and their potential contacts here also shown cataloged on the right side. As an aside, note that these aromatic residues are not among those for which single aa substitutions were presented in the current study.

Additional residues seemingly contributing to the BLD1:BLD2 contacts are part of a BLD1 α-helix that is here shown orange-colored, representing the same BLD1 α-helix also shown orange-colored, e.g., in Figure 6C. In the BLD1 of *Hs*ZC3HC1, those helix residues whose side chains point towards BLD2 include F79, V82, and E83, with these then potentially in contact with M276, K278, N433, and, in particular, again with W428. Furthermore, contacts that can be regarded as corresponding, are also evident at the BLD1:BLD2 interfaces of *Dd*ZC3HC1 and *Sc*Pml39p.

While an α-helix equivalent to this particular one in the BLD1 also exists in the BLD2 (see, e.g., Figure 6C) and even in the BIR domains (e.g., Supplemental Figure S10E), where they are then also shown orange-colored, neither the BLD2 nor the BIR domains' orange-colored α-helices contribute to the BLD1:BLD2 nor a BIR:BIR domain interface, respectively. Instead, the common feature of all of the BLD and BIR domains' orange-colored α-helices, which defines their equivalence, are those helix residues, also including an evolutionarily conserved arginine, that are oriented towards the domains' central parts where they allow for distinct intra-BLD contacts (e.g., Supplemental Figure S10F and S10G).

*In toto*, the predicted proximity of each homolog's two BLDs to each other and the numerous potential inter-BLD contacts, with the resulting overarching arrangement then reminiscent of two BLDs tethered to each other, confers the impression of a NuBaID that represents a compact entity of two adjoining modules. As such, it differs notably from the BIR domains within, e.g., those human BIR proteins that possess more than one BIR domain, namely *Hs*BIRC1 to *Hs*BIRC4. In the latter, the BIR domains are predicted to either exhibit no inter-BIR domain contacts at all or only such that appear relatively loose, as specified further below.

**(B3)** Alignment of the three homologs' BLD1 and BLD2 plus some additional BLD-flanking sequence segments. The high-lighting of the G-[WY] dipeptides, the C-$X_{(2)}$-C tetrapeptide, and the H-$X_{(3)}$-C pentapeptide is like in Figure 4B, while those residues high-lighted in grey are the ones predicted potentially capable of inter-BLD contacts. Those contacts between the P-L dipeptide of each of the three homologs' BLD1 on the one hand and the aromatic residues positioned between the histidine and cysteine of the H-$X_{(3)}$-C pentapeptide of BLD2 on the other are accentuated by a connector line.

We further regard it as worth mentioning that the BLD1:BLD2 interface appears notably distinct from the few predicted BIR:BIR domain contacts within the four human BIRC proteins in which more than one BIR domain exists: In the case of *Hs*BIRC1 (https://alphafold.ebi.ac.uk/entry/Q13075), we found only a few contacts predicted to occur between the *Hs*BIRC1 protein's BIR1 and BIR2 domain, with these contacts notably dissimilar from those of the BLD1:BLD2 interface. In *Hs*BIRC2 and *Hs*BIRC3 (https://alphafold.ebi.ac.uk/entry/Q13490; https://alphafold.ebi.ac.uk/entry/Q13489), the BIR domains were predicted so far apart from each other that contact between them appeared unlikely. Merely for *Hs*BIRC4 (https://

alphafold.ebi.ac.uk/entry/P98170) some likely contacts were predicted to occur between its BIR2 and BIR3 domain that involved an aromatic residue located between the histidine and cysteine of the H-$X_{(6)}$-C octapeptide of BIR3 and residues of an adjacent α-helix of BIR2. The latter α-helix, though, is not equivalent to the orange-colored one of BLD1, with this latter instead equivalent to yet another α-helix of the BIR domains, as described further below, where this other BIR domain α-helix is then shown as orange-colored too (Supplemental Figure S10E). Apart from that, also note that the BLD1:BLD2 interface does not overlap with the IBM-reminiscent groove of the human BLD1 (see later Supplemental Figure S10D), with this particular groove located on the BLD1 surface side opposite to the one contributing to the BLD1:BLD2 interface.

**(C)** The positioning of the evolutionarily conserved aromatic residues flanking the zinc ion coordination spheres of BLD1 and BLD2.

**(C1)** Structures predicted for central regions of the BLD1 and BLD2 modules of *Hs*ZC3HC1 and *Sc*Pml39p. The outer boundaries of the here shown part of BLD1 are S74 and G168 for *Hs*ZC3HC1 and D82 and Y189 for *Sc*Pml39p. The outer boundaries of the part of BLD2 shown here correspond to P175 and S472 of *Hs*ZC3HC1 and S192 and D312 of *Sc*Pml39p. The major loop-like insertion (E288 to S417) and a minor loop (I434 to E455) of the human homolog's BLD2 have been blanked out, as in Figure 6C. Side chains are shown for those histidines (highlighted in blue) and cysteines (green) that are engaged in the likely zinc ion coordination, next to the flanking tryptophans' or tyrosines' side chains (highlighted in magenta). These aromatic residues are W107, W158, W256, and W431 in *Hs*ZC3HC1 and W119, W178, Y257, and Y294 in *Sc*Pml39p.

**(C2)** Residues flanking the zinc coordination spheres of both homologs' two BLDs, shown at higher magnification with side chains colored as in S9C1. Note that the *Hs*ZC3HC1 residues W107 and W256 and the *Sc*Pml39p residues W119 and Y257, which we had found crucial for NE-association and TPR/Mlp-binding (Figures 2 and 4; Supplemental Figures S2 and S5), appear to be involved in protecting and shielding the histidines of the zinc ion coordination spheres, with the histidines' and cysteines' side chains again shown as well. In addition, we found the now-emerging role of two *Hs*ZC3HC1 NuBaID signature residues, W158 and W431, and the corresponding ones from *Sc*Pml39p, namely W178 and Y294, of particular interest. In our single aa substitution experiments, the loss of the tryptophan's indole ring at such a position had been found to still allow for the mutant proteins' NE-association in ZC3HC1 KO cells and impaired binding to TPR (Figure 2; Supplemental Figure S2). Now, AlphaFold2's structure predictions indicate that these residues, while also located within the BLDs' central regions, do not appear to be engaging in such type of BLD core-stabilizing intramolecular interactions with other side chains that one would instantaneously regard as being essential. In other words, these particular residues appear less required than others for maintaining the structural integrity of the BLDs' core structures. While the indole ring of both W158 and W431, and correspondingly the W178 indole and Y294 phenol ring of *Sc*Pml39p, do appear to be in contact with an evolutionarily conserved arginine of a particular α-helix occurring in both BLD1 and BLD2 (see later Supplemental Figure S10F and S10G), such contacts rather provide the impression of primarily keeping the aromatic residues in place. There, they appear to occupy, like a lid, a hollow space at the "entrance" to the zinc coordination sphere in a manner that would shield this entrance, suggesting that the function of these residues might be the seemingly less crucial protection of the coordinated zinc ion from water.

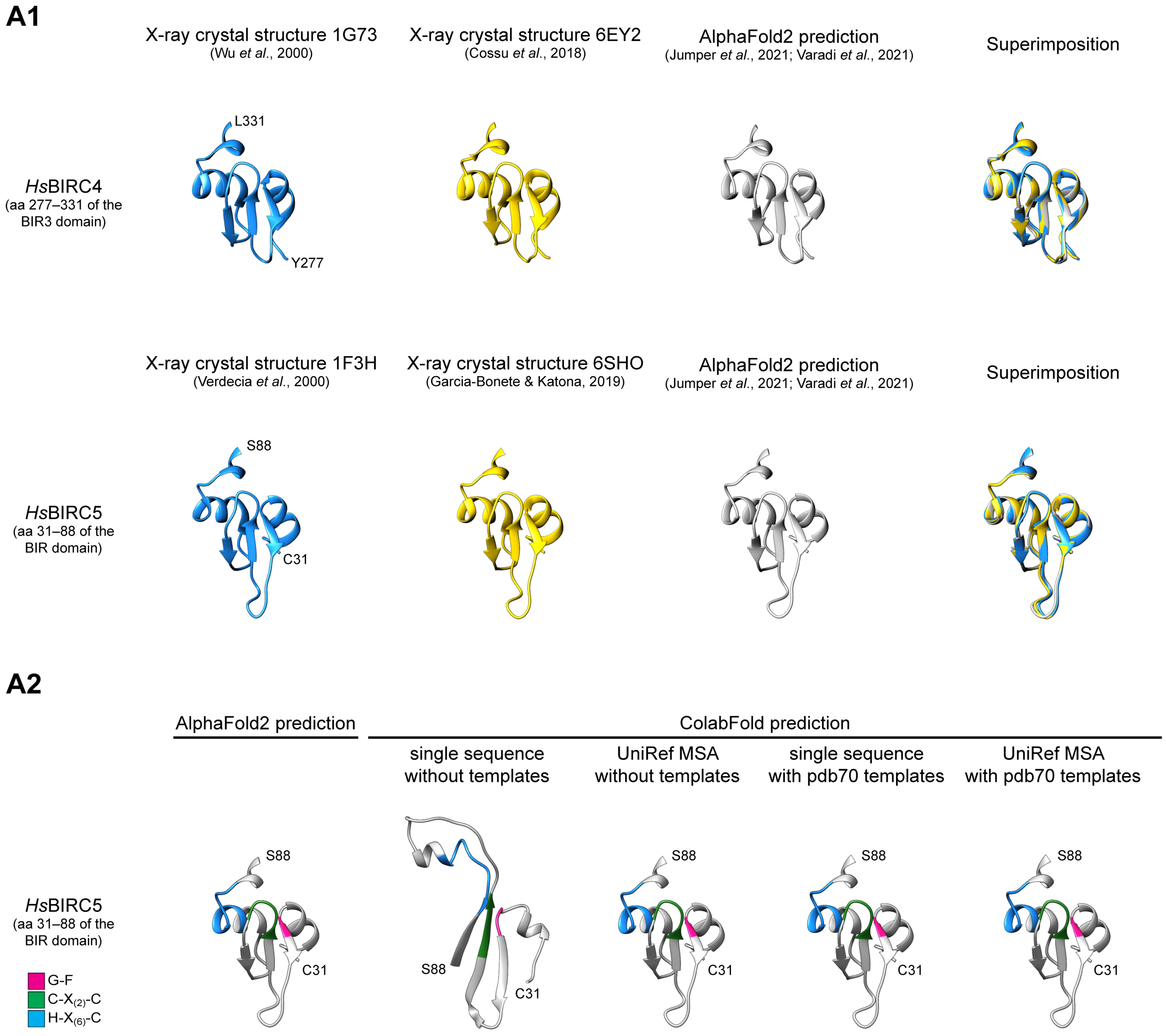
A1
X-ray crystal structure 1G73
(Wu *et al.*, 2000)
X-ray crystal structure 6EY2
(Cossu *et al.*, 2018)
AlphaFold2 prediction
(Jumper *et al.*, 2021; Varadi *et al.*, 2021)
Superimposition
*Hs*BIRC4
(aa 277–331 of the BIR3 domain)
L331
Y277
X-ray crystal structure 1F3H
(Verdecia *et al.*, 2000)
X-ray crystal structure 6SHO
(Garcia-Bonete & Katona, 2019)
AlphaFold2 prediction
(Jumper *et al.*, 2021; Varadi *et al.*, 2021)
Superimposition
*Hs*BIRC5
(aa 31–88 of the BIR domain)
S88
C31
A2
AlphaFold2 prediction
ColabFold prediction
single sequence without templates
UniRef MSA without templates
single sequence with pdb70 templates
UniRef MSA with pdb70 templates
*Hs*BIRC5
(aa 31–88 of the BIR domain)
G-F
C-X$_{(2)}$-C
H-X$_{(6)}$-C
S88
C31

S10 (1/7)

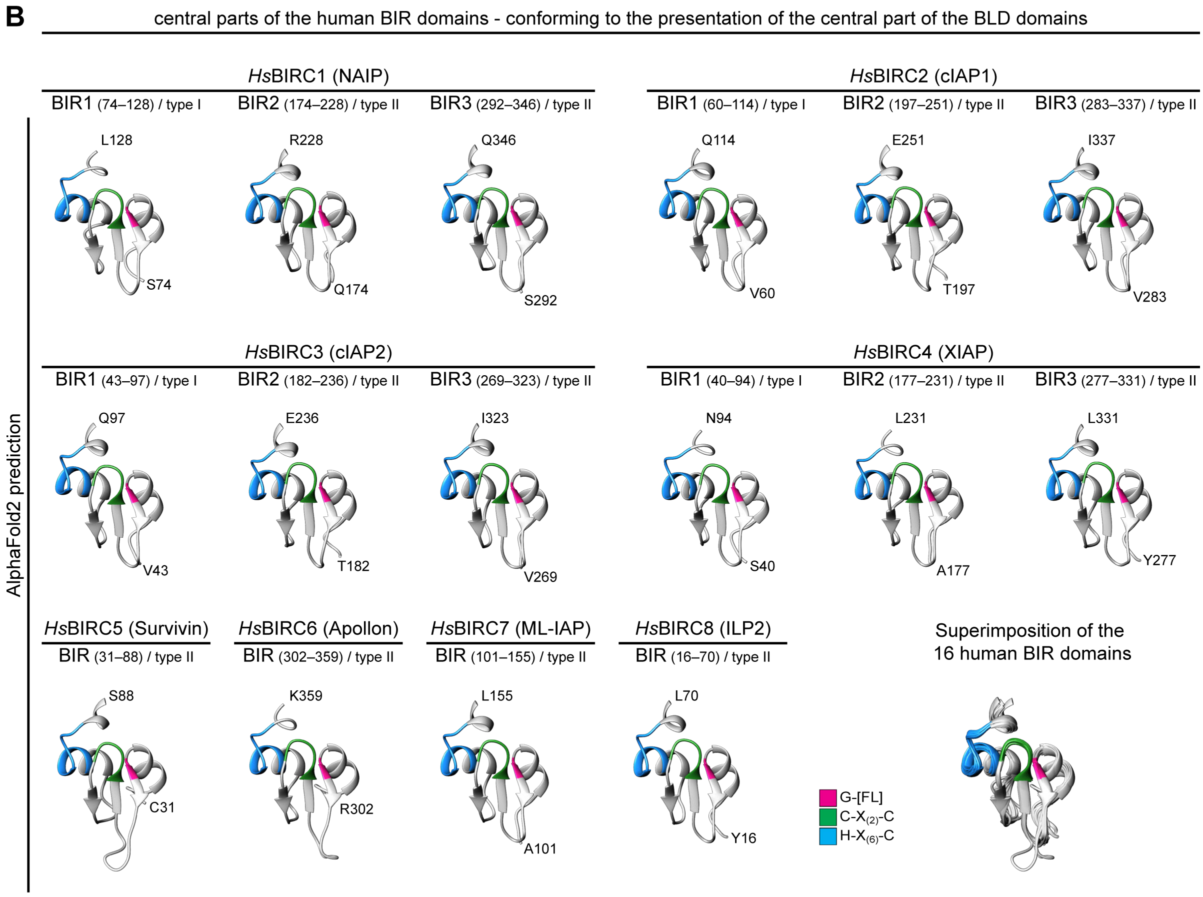
B
central parts of the human BIR domains - conforming to the presentation of the central part of the BLD domains
HsBIRC1 (NAIP)
BIR1 (74–128) / type I
BIR2 (174–228) / type II
BIR3 (292–346) / type II
HsBIRC2 (cIAP1)
BIR1 (60–114) / type I
BIR2 (197–251) / type II
BIR3 (283–337) / type II
L128
S74
R228
Q174
Q346
S292
Q114
V60
E251
T197
I337
V283
HsBIRC3 (cIAP2)
BIR1 (43–97) / type I
BIR2 (182–236) / type II
BIR3 (269–323) / type II
HsBIRC4 (XIAP)
BIR1 (40–94) / type I
BIR2 (177–231) / type II
BIR3 (277–331) / type II
Q97
V43
E236
T182
I323
V269
N94
S40
L231
A177
L331
Y277
AlphaFold2 prediction
HsBIRC5 (Survivin)
BIR (31–88) / type II
HsBIRC6 (Apollon)
BIR (302–359) / type II
HsBIRC7 (ML-IAP)
BIR (101–155) / type II
HsBIRC8 (ILP2)
BIR (16–70) / type II
Superimposition of the
16 human BIR domains
S88
C31
K359
R302
L155
A101
L70
Y16
G-[FL]
C-X(2)-C
H-X(6)-C

**S10 (2/7)**

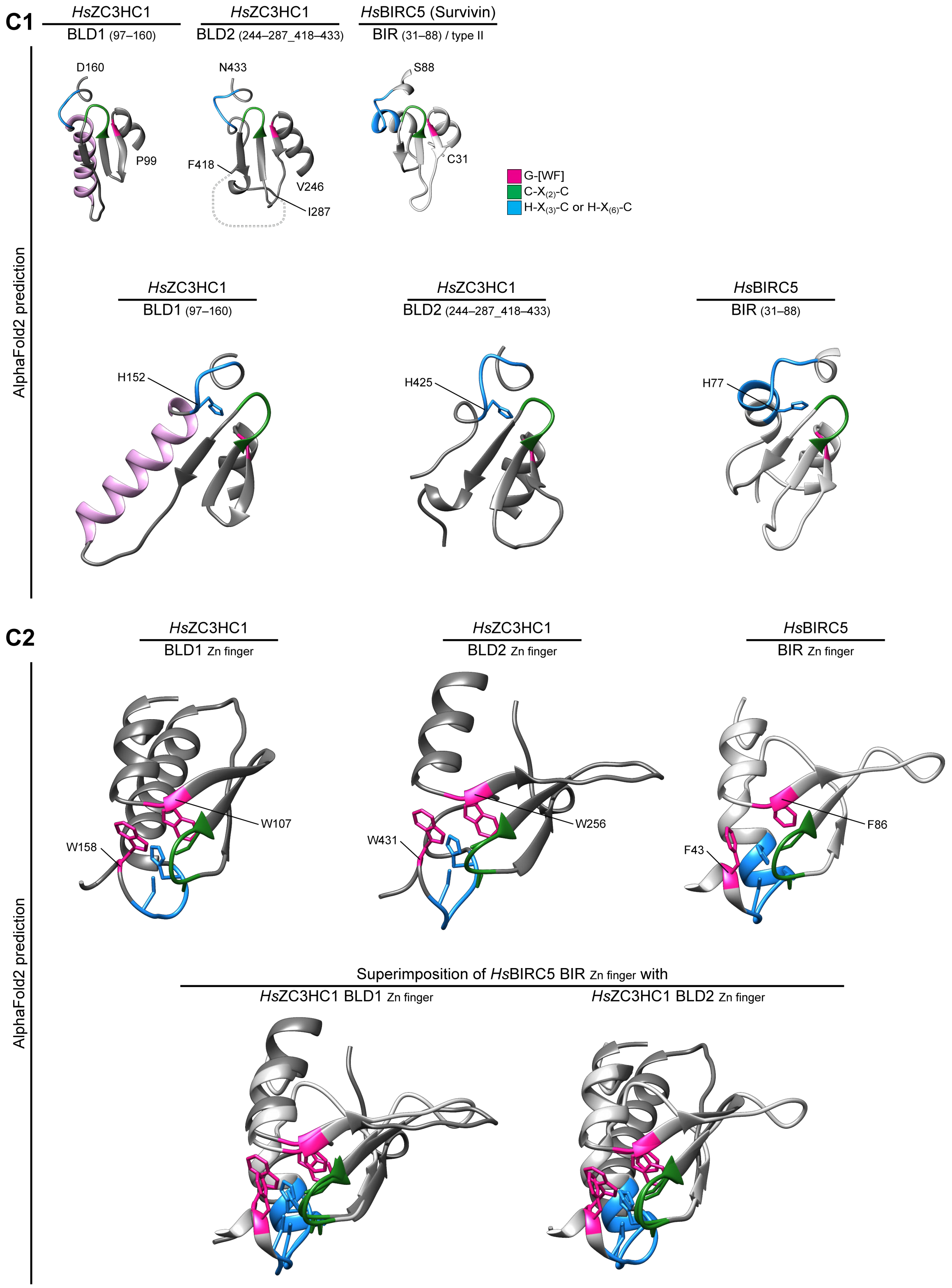
C1
HsZC3HC1
BLD1 (97–160)
HsZC3HC1
BLD2 (244–287_418–433)
HsBIRC5 (Survivin)
BIR (31–88) / type II
D160
P99
N433
F418
V246
I287
S88
C31
G-[WF]
C-X(2)-C
H-X(3)-C or H-X(6)-C
AlphaFold2 prediction
HsZC3HC1
BLD1 (97–160)
HsZC3HC1
BLD2 (244–287_418–433)
HsBIRC5
BIR (31–88)
H152
H425
H77
C2
HsZC3HC1
BLD1 Zn finger
HsZC3HC1
BLD2 Zn finger
HsBIRC5
BIR Zn finger
W107
W158
W256
W431
F86
F43
AlphaFold2 prediction
Superimposition of HsBIRC5 BIR Zn finger with
HsZC3HC1 BLD1 Zn finger
HsZC3HC1 BLD2 Zn finger

**S10 (3/7)**

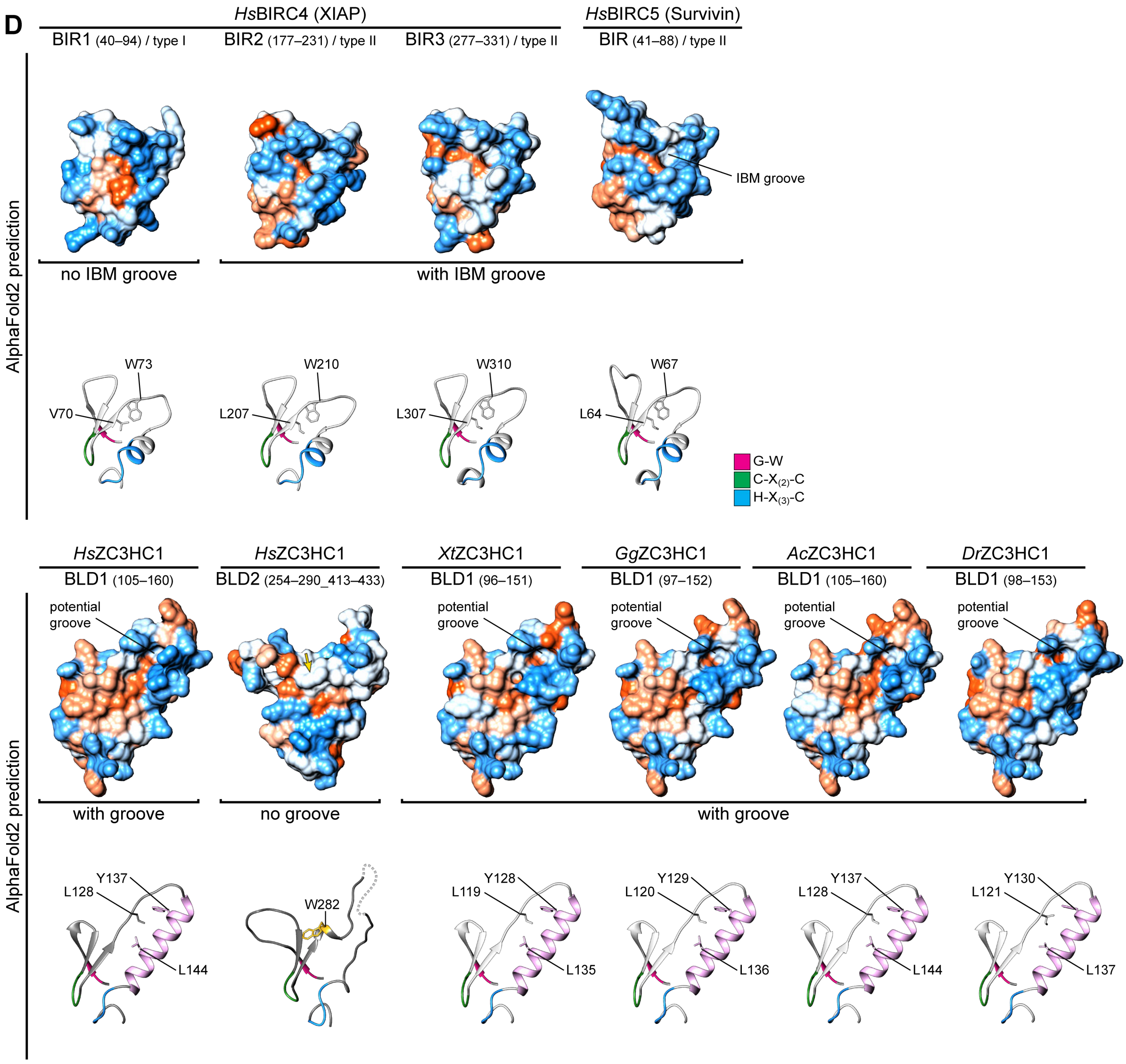
D
HsBIRC4 (XIAP)
HsBIRC5 (Survivin)
BIR1 (40–94) / type I
BIR2 (177–231) / type II
BIR3 (277–331) / type II
BIR (41–88) / type II
AlphaFold2 prediction
IBM groove
no IBM groove
with IBM groove
W73
V70
W210
L207
W310
L307
W67
L64
G-W
C-X(2)-C
H-X(3)-C
HsZC3HC1
BLD1 (105–160)
HsZC3HC1
BLD2 (254–290_413–433)
XtZC3HC1
BLD1 (96–151)
GgZC3HC1
BLD1 (97–152)
AcZC3HC1
BLD1 (105–160)
DrZC3HC1
BLD1 (98–153)
potential groove
with groove
no groove
with groove
Y137
L128
L144
W282
Y128
L119
L135
Y129
L120
L136
Y137
L128
L144
Y130
L121
L137

**S10 (4/7)**

**E1**

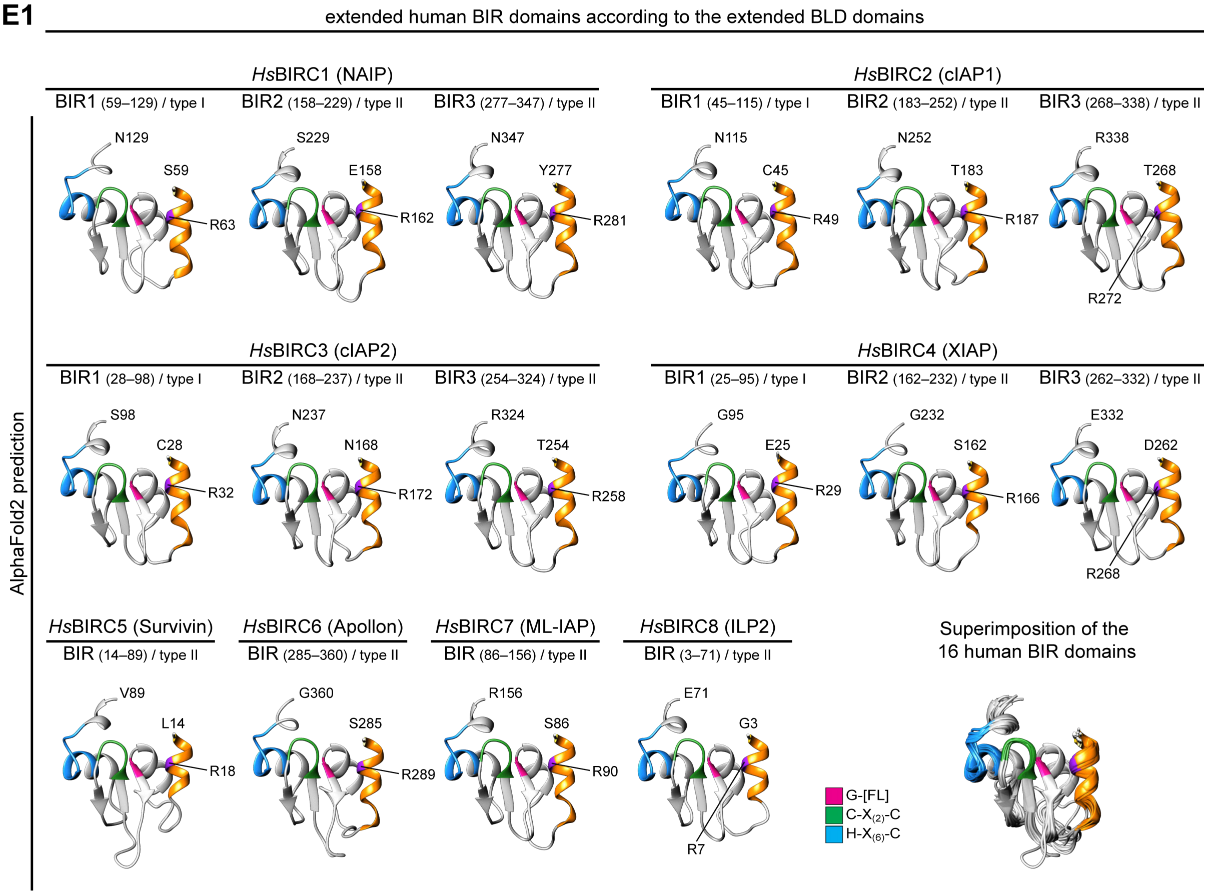
extended human BIR domains according to the extended BLD domains
HsBIRC1 (NAIP)
BIR1 (59–129) / type I
BIR2 (158–229) / type II
BIR3 (277–347) / type II
HsBIRC2 (cIAP1)
BIR1 (45–115) / type I
BIR2 (183–252) / type II
BIR3 (268–338) / type II
N129
S59
R63
S229
E158
R162
N347
Y277
R281
N115
C45
R49
N252
T183
R187
R338
T268
R272
HsBIRC3 (cIAP2)
BIR1 (28–98) / type I
BIR2 (168–237) / type II
BIR3 (254–324) / type II
HsBIRC4 (XIAP)
BIR1 (25–95) / type I
BIR2 (162–232) / type II
BIR3 (262–332) / type II
S98
C28
R32
N237
N168
R172
R324
T254
R258
G95
E25
R29
G232
S162
R166
E332
D262
R268
AlphaFold2 prediction
HsBIRC5 (Survivin)
BIR (14–89) / type II
HsBIRC6 (Apollon)
BIR (285–360) / type II
HsBIRC7 (ML-IAP)
BIR (86–156) / type II
HsBIRC8 (ILP2)
BIR (3–71) / type II
Superimposition of the
16 human BIR domains
V89
L14
R18
G360
S285
R289
R156
S86
R90
E71
G3
R7
G-[FL]
C-X(2)-C
H-X(6)-C

**E2**

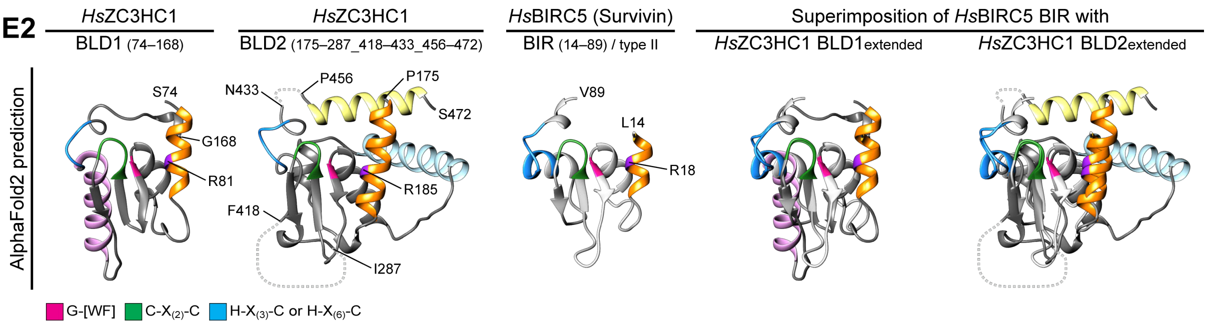
HsZC3HC1
BLD1 (74–168)
HsZC3HC1
BLD2 (175–287_418–433_456–472)
HsBIRC5 (Survivin)
BIR (14–89) / type II
Superimposition of HsBIRC5 BIR with
HsZC3HC1 BLD1extended
HsZC3HC1 BLD2extended
AlphaFold2 prediction
S74
G168
R81
N433
P456
P175
S472
R185
F418
I287
V89
L14
R18
G-[WF]
C-X(2)-C
H-X(3)-C or H-X(6)-C

**S10 (5/7)**

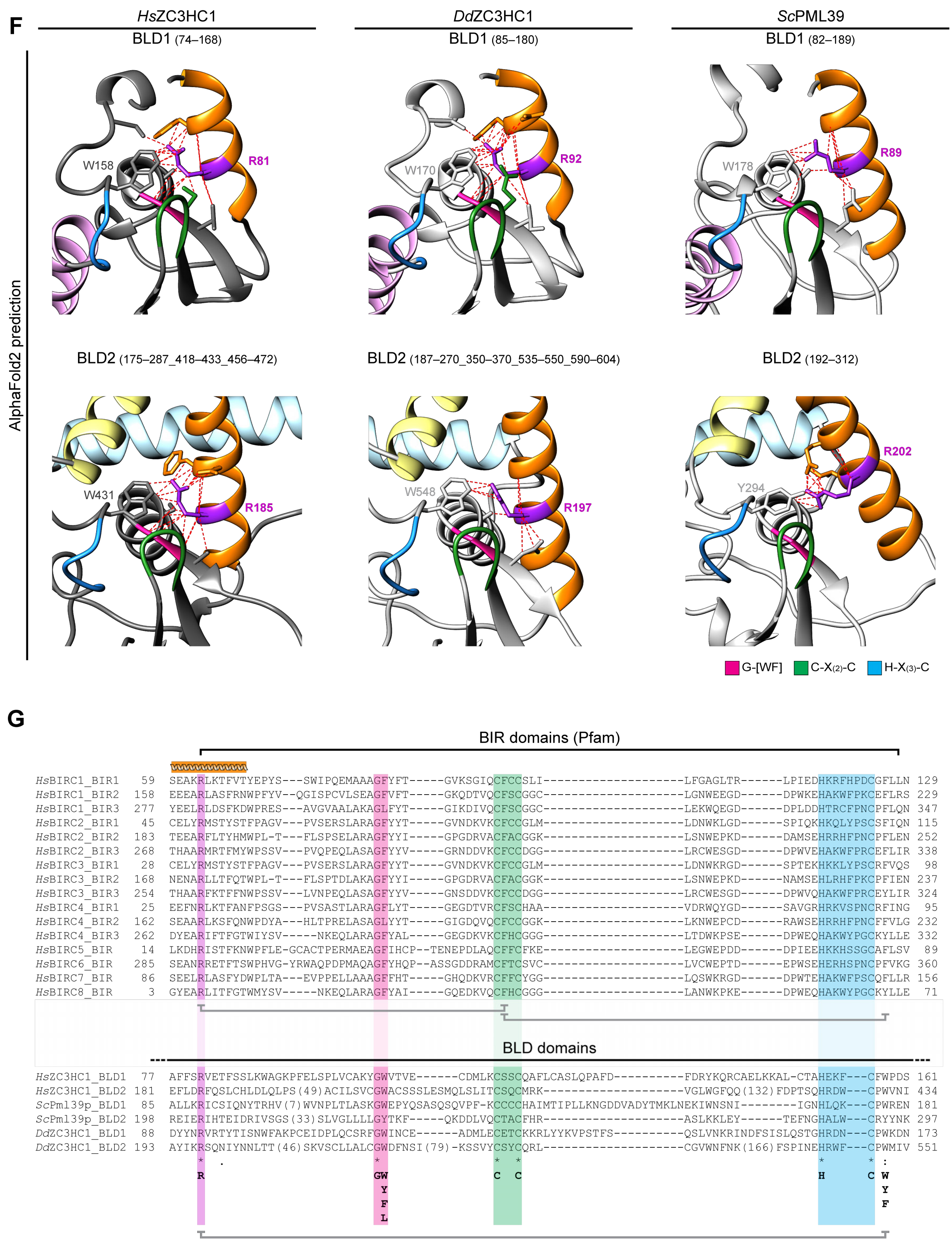
F
HsZC3HC1
BLD1 (74–168)
DdZC3HC1
BLD1 (85–180)
ScPML39
BLD1 (82–189)
AlphaFold2 prediction
W158
R81
W170
R92
W178
R89
BLD2 (175–287_418–433_456–472)
BLD2 (187–270_350–370_535–550_590–604)
BLD2 (192–312)
W431
R185
W548
R197
Y294
R202
G-[WF]
C-X(2)-C
H-X(3)-C
G
BIR domains (Pfam)
HsBIRC1_BIR1 59 SEAKRLKTFVTYEPYS---SWIPQEMAAAGFYFT-----GVKSGIQCFCCSLI--------------------LFGAGLTR------LPIEDHKRFHPDCGFLLN 129
HsBIRC1_BIR2 158 EEEARLASFRNWPFYV--QGISPCVLSEAGFVFT-----GKQDTVQCFSCGGC--------------------LGNWEEGD------DPWKEHAKWFPKCEFLRS 229
HsBIRC1_BIR3 277 YEELRLDSFKDWPRES---AVGVAALAKAGLFYT-----GIKDIVQCFSCGGC--------------------LEKWQEGD------DPLDDHTRCFPNCPFLQN 347
HsBIRC2_BIR1 45 CELYRMSTYSTFPAGV---PVSERSLARAGFYYT-----GVNDKVKCFCCGLM--------------------LDNWKLGD------SPIQKHKQLYPSCSFIQN 115
HsBIRC2_BIR2 183 TEEARFLTYHMWPL-T---FLSPSELARAGFYYI-----GPGDRVACFACGGK--------------------LSNWEPKD------DAMSEHRRHFPNCPFLEN 252
HsBIRC2_BIR3 268 THAARMRTFMYWPSSV---PVQPEQLASAGFYYV-----GRNDDVKCFCCDGG--------------------LRCWESGD------DPWVEHAKWFPRCEFLIR 338
HsBIRC3_BIR1 28 CELYRMSTYSTFPAGV---PVSERSLARAGFYYT-----GVNDKVKCFCCGLM--------------------LDNWKRGD------SPTEKHKKLYPSCRFVQS 98
HsBIRC3_BIR2 168 NENARLLTFQTWPL-T---FLSPTDLAKAGFYYI-----GPGDRVACFACGGK--------------------LSNWEPKD------NAMSEHLRHFPKCPFIEN 237
HsBIRC3_BIR3 254 THAARFKTFFNWPSSV---LVNPEQLASAGFYYV-----GNSDDVKCFCCDGG--------------------LRCWESGD------DPWVQHAKWFPRCEYLIR 324
HsBIRC4_BIR1 25 EEFNRLKTFANFPSGS---PVSASTLARAGFLYT-----GEGDTVRCFSCHAA--------------------VDRWQYGD------SAVGRHRKVSPNCRFING 95
HsBIRC4_BIR2 162 SEAARLKSFQNWPDYA---HLTPRELASAGLYYT-----GIGDQVQCFCCGGK--------------------LKNWEPCD------RAWSEHRRHFPNCFFVLG 232
HsBIRC4_BIR3 262 DYEARIFTFGTWIYSV-----NKEQLARAGFYAL-----GEGDKVKCFHCGGG--------------------LTDWKPSE------DPWEQHAKWYPGCKYLLE 332
HsBIRC5_BIR 14 LKDHRISTFKNWPFLE-GCACTPERMAEAGFIHCP--TENEPDLAQCFFCFKE--------------------LEGWEPDD------DPIEEHKKHSSGCAFLSV 89
HsBIRC6_BIR 285 SEANRRETFTSWPHVG-YRWAQPDPMAQAGFYHQP--ASSGDDRAMCFTCSVC--------------------LVCWEPTD------EPWSEHERHSPNCPFVKG 360
HsBIRC7_BIR 86 SEELRLASFYDWPLTA---EVPPELLAAAGFFHT-----GHQDKVRCFFCYGG--------------------LQSWKRGD------DPWTEHAKWFPSCQFLLR 156
HsBIRC8_BIR 3 GYEARLITFGTWMYSV-----NKEQLARAGFYAI-----GQEDKVQCFHCGGG--------------------LANWKPKE------DPWEQHAKWYPGCKYLLE 71
BLD domains
HsZC3HC1_BLD1 77 AFFSRVETFSSLKWAGKPFELSPLVCAKYGWVTVE------CDMLKCSSCQAFLCASLQPAFD----------FDRYKQRCAELKKAL-CTAHEKF---CFWPDS 161
HsZC3HC1_BLD2 181 EFLDRFQSLCHLDLQLPS(49)ACILSVCGWACSSSLESMQLSLITCSQCMRK--------------------VGLWGFQQ(132)FDPTSQHRDW---CPWVNI 434
ScPml39p_BLD1 85 ALLKRICSIQNYTRHV(7)WVNPLTLASKGWEPYQSASQSQVPF-KCCCCHAIMTIPLLKNGDDVADYTMKLNEKIWNSNI--------IGNHLQK---CPWREN 181
ScPml39p_BLD2 198 REIERIHTEIDRIVSGS(33)SLVGLLLLGYTKF-----QKDDLVQCTACFHR--------------------ASLKKLEY------TEFNGHALW---CRYYNK 297
DdZC3HC1_BLD1 88 DYYNRVRTYTISNWFAKPCEIDPLQCSRFGWINCE------ADMLECETCKKRLYYKVPSTFS----------QSLVNKRINDFSISLQSTGHRDN---CPWKDN 173
DdZC3HC1_BLD2 193 AYIKRSQNIYNNLTT(46)SKVSCLLALCGWDFNSI(79)-KSSVYCSYCQRL--------------------CGVWNFNK(166)FSPINEHRWF---CPWMIV 551
R
GW Y F L
C C
H C W Y F

S10 (6/7)

**H**

zf-C3HC / PF07967 (Pfam 35 release)

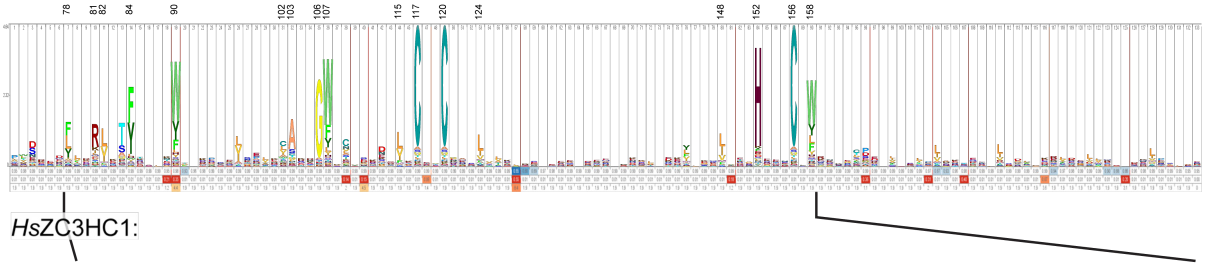

residues in range for potential contacts

| F78 | R81 | V82 | T84 | F85 | W90 | C102 | A103 | G106 | W107 | L115 | C117 | C120 | L124 | L148 | H152 | C156 | W158 |
|---|---|---|---|---|---|---|---|---|---|---|---|---|---|---|---|---|---|
| R81 | F78 | F78 | F85 | T84 | F85 | F85 | F78 | R81 | Y105 | F85 | W107 | C117 | W107 | L97 | Y105 | C117 | E71 |
| V82 | A103 | L100 | V108 | W90 | K93 | W90 | R81 | W158 | L115 | W90 | S119 | A122 | L115 | V101 | W107 | C120 | S72 |
| L100 | G106 | A103 | | P99 | P99 | L148 | V82 | | C117 | W107 | C120 | H152 | A147 | C102 | C117 | H152 | R81 |
| A103 | V108 | | | C102 | C102 | | | | L124 | T109 | H152 | E153 | L148 | Y105 | C120 | W158 | G106 |
| S163 | S118 | W428 | | T109 | L115 | | | | L148 | L148 | C156 | C156 | | W107 | C156 | | C117 |
| F167 | W158 | | | L115 | A126 | | | | H152 | | W158 | | | L115 | W158 | | S118 |
| | S163 | | | | | | | | | | | | | L124 | P159 | | S119 |
| | | | | | | | | | | | | | | | | | H152 |
| | | | | | | | | | | | | | | | | | C156 |
| | | | | | | | | | | | | | | | | | F157 |
| | | | | | | | | | | | | | | | | | S163 |

orange helix (F78–T84); G-W (G106–W107); C-X-X-C (C117–C120); H-X-X-X-C (H152–C156)

■ surface exposed ■ partially surface exposed ■ BLD1:BLD2 connection ■ not surface exposed

Rsm1 / PF08600 (Pfam 35 release)

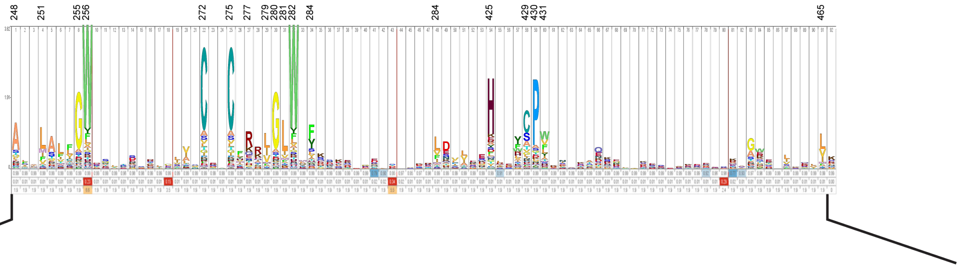

residues in range for potential contacts

| R185 | A248 | L251 | G255 | W256 | C272 | C275 | R277 | V279 | G280 | L281 | W282 | F284 | H425 | C429 | P430 | W431 | L465 |
|---|---|---|---|---|---|---|---|---|---|---|---|---|---|---|---|---|---|
| E181 | F186 | T247 | R185 | C254 | Q274 | C272 | C275 | W256 | L269 | L192 | G280 | P197 | W256 | C272 | Q274 | R185 | V179 |
| F182 | L189 | L189 | W431 | I270 | C275 | R277 | V279 | R277 | W282 | P197 | G283 | V279 | C272 | C275 | C429 | G255 | F182 |
| S252 | C190 | L196 | | Q274 | H425 | R426 | Q424 | F284 | | S268 | | F419 | C429 | H425 | | L270 | L217 |
| G255 | L196 | F419 | | V279 | C429 | W428 | R426 | F419 | | I270 | | | W431 | W431 | | C272 | L220 |
| A257 | | P421 | | H425 | W431 | C429 | | | | | | | V432 | | | Q274 | V461 |
| W431 | | | | | | | | | | | | | | | | H425 | |
| | | | | | | | | | | | | | | | | C429 | |
| | | | | | | | | | | | | | | | | V461 | |

orange helix (not part of the Pfam Rsm1 motif) (R185); G-W (G255–W256); C-X-X-C (C272–C275); H-X-X-X-C (H425–C429)

**Supplemental Figure S10. Comparing the predicted structural characteristics and sequence features of the BLDs of ZC3HC1 with the BIR domains' structures and sequences.**

Altogether, the structural features of the BLDs' central parts, as predicted by AlphaFold2, turned out similar to those characterizing the BIR domains of the IAPs, the latter here exemplified by the eight BIR domain-containing proteins in humans. We, humans, possess a total of 16 different BIR domains, of which four belong to the so-called type I and twelve to the type II of BIR domains (e.g., Oberoi-Khanuja *et al*, 2013). Such type I and II BIR domains exhibit some differences, like a small binding groove of physiological relevance that exists in the type II BIR domains while absent in type I (e.g., Cossu *et al*, 2019; and see further below). However, apart from such few relatively minor structural differences, the overall construction of all these 16 BIR domains, particularly the construction of their zinc coordination spheres, was already known to be highly similar to each other (e.g., Cossu *et al*, 2019). In the following, such similarity is now illustrated by these BIR domains' AlphaFold2-predicted structures, presented in S10B and S10E. Furthermore, the close similarity between some exemplary BIR domain structures, as determined by X-ray crystallography and predicted by AlphaFold2, is shown in S10A1. In addition, S10A2 shows a representative comparison of a BIR domain, as present in the AlphaFold2 database, with the same domain's structures predicted via the ColabFold platform, having used either only this domain's sequence alone or an UniRef MSA, with and without additional consideration of PDB70 templates, as outlined in Supplemental Figure S7. The rationale for this approach, namely assessing the contributions of the sequence database search-derived MSAs, on the one hand, and the PDB-deposited structures as templates, on the other, to the computationally predicted BIR domains' structures, has been described further above (Supplemental Information 3 and Supplemental Figure S7).

In line with the former proposal that the BIR domain of human survivin/BIRC5 and the BLD1 of *Hs*ZC3HC1 are most likely to be structurally very similar (Higashi *et al*, 2005), the outcome of the here presented comparisons of the AlphaFold2/ColabFold-predicted structures illustrates that the BIR domains' structural elements, and their arrangements relative to each other, resemble the BLDs' corresponding parts. In the following, in S10C, this is exemplified by the comparison of the two AlphaFold2-predicted structures of the *Hs*ZC3HC1 BLDs' central parts with those of the three BIR domains of *Hs*XIAP/BIRC4 and the single one of *Hs*BIRC5.

Furthermore, the predictions of the BLDs' structures by Alphafold2 allowed for unveiling yet other features shared by the one or other BLD with either the BIR type I or type II domain beyond the similarities of their central, zinc ion coordination spheres. Among such features are, e.g., a conspicuous groove in the vertebrate homologs' BLD1, located on the side opposite the abovementioned BLD1:BLD2 interface with its hydrophobic residues. This groove appears akin to the so-called IAP binding motif (IBM) peptide-binding groove that characterizes the type II BIR domain, where it functions as a binding site for IAP antagonists (e.g., Cossu *et al*, 2019). On the other hand, such a groove is absent from the type I BIR domains, and we found it to appear similarly absent, altered, or masked in the ZC3HC1 homologs' BLD2. In the latter, a few surface-exposed protruding residues, evolutionarily relatively conserved, are predicted to be positioned at the IBM and BLD1 groove-corresponding positions instead. Of note, though, the BIR domains of type II nonetheless share notable sequence similarities with the BLD2 at precisely those positions that correspond to the IBM groove, with a tryptophan forming the bottom of the BIR domains' IBM groove being equivalent to a tryptophan instead protruding from the BLD2 surface, here W282 of *Hs*ZC3HC1. Supplemental Figure S10D illustrates the grooves' absence and presence in the different BLD and BIR domains, thereby pointing to also some of the residues involved.

Then, comparing the other structural elements of the BLDs and BIR domains, beyond the central parts focused on in S10C, we noted, apart from some apparent differences, also yet another evident similarity. Having identified the *Hs*ZC3HC1 α-helix aa 175–190 as a structural element of the BLD2 within its newly defined boundaries (e.g., Figure 6D), equivalent to a corresponding α-helix in the human BLD1, and the two latter as also correspondingly present in the amoebic and budding yeast homolog's BLDs (e.g., orange-colored α-helices in Figure 6C), we noted these to be potentially equivalent to an α-helix that represents an extension of the BIR domain at its N-terminus. Since part of this α-helix extends beyond the BIR domains' currently defined boundaries, we now propose redefining the BIR domain's N-terminal boundary accordingly. This additional structural similarity is illustrated in S10E, where these particular α-helices are again shown as orange-colored.

Of particular note, these orange-colored α-helices share an evolutionarily conserved arginine of apparently similar function. In the ZC3HC1 homologs, this arginine appears to be involved in stabilizing the position of its α-helix relative to the BLD's more central parts. Moreover, this arginine is also predicted to contribute to additionally stabilizing the position of one of those aromatic residues we had already studied in our single aa substitution experiments. These potential intra-BLD contacts that this particular arginine might be capable of are illustrated in S10F.

Furthermore, based on such structural similarities and our reinspection of the BLD and BIR domains' sequences, we present the corresponding sequence alignments in S10G. For such alignments, we had considered the newly defined BLD boundaries and the orange-colored α-helices now rated as equivalent, which in turn allowed for revealing the additional sequence similarity we regard as noteworthy.

Finally, regarding those residues of the zf-C3HC and Rsm1 motifs that appear evolutionarily most conserved, according to current Pfam release 35.0 information, we present the corresponding HMM logos and these residues' positions relative to the BLDs' central parts and surfaces in S10H, thereby allowing to assess which of these residues are more likely to permit intra-BLD and inter-BLD contacts or such between a BLD and other proteins.

**(A)** Comparison of representative BIR domains' crystal structures with structures predictable by AlphaFold2 and ColabFold using default and altered parameter settings.

**(A1)** Comparison of the central parts of the structures of BIR3 of *Hs*BIRC4 and the single BIR domain of *Hs*BIRC5, as predicted by AlphaFold2, with the same domains' structural elements formerly determined as part of the *Hs*BIRC4 and *Hs*BIRC5 proteins by X-ray crystallography (Wu *et al*; Verdecia *et al*, 2000; Corti *et al*, 2018; Garcia-Bonete & Katona, 2019). For clarity, only the BIR domain segments aa 277–331 of *Hs*BIRC4 and aa 31–88 of *Hs*BIRC5 are shown for comparison, with the protein's other parts, including some side chains presented as parts of the PDB-deposited crystal structures, rendered invisible. As an aside, note that for these BIR domain examples, the Pfam database' BIR motif (https://pfam.xfam.org/family/BIR) comprises aa 268–331 for the BIR3 of *Hs*BIRC4 and aa 18–88 for *Hs*BIRHC5, with the here presented structures thus lacking several BIR signature residues at their N-terminus. The crystallographic data represent parts of the structures found with the identifiers 1G73, 6EY2, 1F3H, and 6SHO in the PDB database (https://www.rcsb.org). Superimposition of the structures again was achieved with Chimera's MatchMaker tool. Note that the close similarity between the structures determined by crystallography and the one predicted by AlphaFold2 appears evident.

**(A2)** Assessment of the contributions of the MSA on the one hand and the PDB database-deposited structures as templates on the other, in BIR structure predictions by AlphaFold2 via ColabFold. The AlphaFold2-predicted structure of the *Hs*BIRC5 BIR domain is shown next to the structures predicted via the ColabFold platform, having used either only the BIR domain's sequence without further alignment with other sequences or having started the prediction from the UniRef MSA provided by ColabFold, with and without additional consideration of the PDB70 templates, including the BIR domains' crystal structures. Note that using neither the MSA nor the template information resulted in a prediction that strongly deviates from the domain's actual structure. By contrast, predictions based either merely on the information provided by the MSA or by the templates alone resulted in structures that appear highly similar, if not hardly distinguishable, from those initially determined by X-ray crystallography, as shown in S10A1.

**(B)** The AlphaFold2-predicted structures of the central parts of the sixteen BIR domains existing in humans, comprising the three BIR domains each of *Hs*BIRC1 (NAIP), *Hs*BIRC2 (cIAP1), *Hs*BIRC3 (cIAP2), and *Hs*BIRC4 (XIAP), and the only one single BIR domain-containing proteins *Hs*BIRC5 (Survivin), *Hs*BIRC6 (Apollon), *Hs*BIRC7 (ML-IAP), and

*Hs*BIRC8 (ILP2). Note that all but one of the BIR domain structures were retrieved from the AlphaFold2 database (https://alphafold.ebi.ac.uk), except for *Hs*BIRC6, which was predicted *via ColabFold.* Further note that those parts of these structures actually shown correspond to the following parts of the Pfam database's BIR signature. For *Hs*BIRC1, aa 74–128 of 63–128 for BIR1, 174–228 of 162–228 for BIR2, and 292–346 of 281–346 for BIR3. For *Hs*BIRC2, 60–114 of 49–114 for BIR1, 197–251 of 187–251 for BIR2, and 283–337 of 272–337 for BIR3. For *Hs*BIRC3, 43–97 of 32–97 for BIR1, 182–236 of 172–236 for BIR2, and 269–323 of 258–323 for BIR3. For *Hs*BIRC4, 40–94 of 29–94 for BIR1, 177–231 of 166–231 for BIR2, and 277–331 of 268–331 for BIR3. For *Hs*BIRC5, 31–88 of 18–88; for *Hs*BIRC6, 302–359 of 289–359; for *Hs*BIRC7, 101–155 of 90–155; for *Hs*BIRC8, 16–70 of 7–70. These central parts of each BIR domain are presented both individually and again superimposed onto each other, once again illustrating the domains' evident structural similarity.

**(C)** Comparison of the BIR domain of BIRC5 with the two BLDs of human ZC3HC1.

**(C1)** The AlphaFold2-predicted structures of the central parts of the two BLDs of *Hs*ZC3HC1, as identically presented in Figure 6A, and here compared with the corresponding central part of the representative BIR domain of *Hs*BIRC5. The structures are shown in two different magnifications and perspectives (upper and lower row), with the lower row also presenting the side chain of the histidine of the BLDs' H-$X_{(3)}$-C pentapeptide and the BIR domain's H-$X_{(6)}$-C octapeptide. Note the structural similarity of these domains' most central parts, including the anti-parallel arrangement of several β-sheets flanked by an α-helix, the latter here colored in grey, next to differences regarding the α-helices that flank the domains' central zinc coordination spheres. In particular, the prominent α-helix of BLD1, colored in light pink as in Figure 6C, cannot be regarded as likely equivalent to the BIR domain's short α-helix here partially colored in blue, with this blue-colored part reflecting the H-$X_{(6)}$-C octapeptide, with its histidine positioned in the middle of this short α-helix, in contrast to the BLDs' zinc ion-coordinating histidines that are positioned beyond the BLDs' α-helix. Further note that additional BLD-specific α-helices distinguishing the one and other BLD from the BIR domains are presented in S10E.

**(C2)** Aromatic residues flanking the zinc coordination spheres of the two BLDs of *Hs*ZC3HC1, shown at higher magnification, compared to corresponding residues in the BIR domains, the latter here again exemplified by the one from *Hs*BIRC5. Note that the aromatic side chains W107, W158, W256, and W431 of *Hs*ZC3HC1 and F43 and F86 of *Hs*BIRC5, all highlighted in magenta, appear relatively similarly arranged to each other and the zinc ion-coordinating residues.

**(D)** Space-filling surface presentations of parts of the type I and type II BIR domains, here represented by those of *Hs*BIRC4 and *Hs*BIRC5, compared to surfaces of parts of the BLD1 and BLD2 of *Hs*ZC3HC1, next to the BLD1 domains of other vertebrates. The BIR domains' structures shown correspond to the following parts of the BIR signature. For *Hs*BIRC4, 40–94 of 29–94 for BIR1, 177–231 of 166–231 for BIR2, and 277–331 of 268–331 for BIR3. For *Hs*BIRC5, 41–88 of 18–88. Surface coloring is according to hydrophobicity, with hydrophobic residues in orange and hydrophilic ones in blue. For *Hs*BIRC5, the so-called IAP binding motif (IBM) peptide-binding groove is marked, the latter characterizing the type II BIR domain while absent from the type I BIR domains. Also marked is a conspicuous groove in the BLD1 of *Hs*ZC3HC1. Such a groove is also detectable in the BLD1 of the other vertebrate homologs. However, it is neither conspicuous in the BLD1 of *Dd*ZC3HC1 and *Sc*Pml39p nor evident in the BLD2 of *Hs*ZC3HC1 and the other vertebrates. In the latter, an evolutionarily relatively well-conserved group of residues (see also S10H), including a surface-exposed, protruding tryptophan, W282 in *Hs*ZC3HC1, here marked by a yellow arrow, which corresponds to W365 in *Dd*ZC3HC1 (not shown here), is predicted to be located at the BLD1 groove-corresponding position instead.

Tempting to regard it as playing a distinct role in the interaction with TPR homologs, such a tryptophan appears evolutionarily conserved in the majority of ZC3HC1 homologs and, as such, it is also part of the Rsm1 motif signature (see also Supplemental Figures S2D1 and S10H). Some homologs, though, exhibit another protruding residue at this position instead, like in *Sc*Pml39p, where this position appears occupied by K278 (not shown here).

The corresponding parts of the BLDs and BIR domains are also presented as ribbons to facilitate correlating the surfaces with the domains' secondary structures. For further orientation, the α-helix of the ZC3HC1 homologs' BLD1 is colored in light pink, corresponding to the coloring of this α-helix in, e.g., S10C1 and Figure 6C. Moreover, those side chains shown additionally as part of the ribbons are, on the one hand, the W282 of the *Hs*ZC3HC1 BLD2, and on the other those residues located at the central bottom of each groove, which in both the BLD1 and BIR domains' grooves represents a residue with an aromatic side chain. In the *Hs*ZC3HC1 BLD1, where residues between L128 and L144 form this groove, a tyrosine, Y137, is centrally positioned at the groove's bottom. Such a tyrosine at the corresponding position is evolutionarily conserved in all vertebrates. In the type II BIR domains, the IBM groove harbors a tryptophan at the groove's dip, for example, W67, in the case of BIRC5. Of note, in the BIR domains, this tryptophan is positioned at the same site relative to the C-$X_{(2)}$-C tetrapeptide as W282 is in the human ZC3HC1 BLD2 (e.g., S10G).

Furthermore, since the IBM grooves of the BIR type II domains are known to interact with IAP antagonists like Smac/DIABLO (see Supplemental Discussion 1 for further information), we consider it of note that minor amounts of Smac/DIABLO, beyond background levels, had been among those materials we found co-sedimented with immunoprecipitated FP-tagged ZC3HC1 polypeptides (our unpublished data). However, whether this might also point to a naturally occurring interaction of physiological relevance still needs to be determined.

**(E)** The AlphaFold2-predicted structures of the two BLDs of *Hs*ZC3HC1 in their entirety, as identically presented in Figure 6C, and here compared with the human BIR domains, as defined by the Pfam database's BIR motif, with now additionally considered BIR motif-flanking aa residues, causing us to refer to these structures as the BIR domains' extended versions.

**(E1)** The AlphaFold2-predicted structures of the central parts of the sixteen BIR domains in humans, here now shown in the extended version. Note that the presented structures again include additional residues flanking the BIR signature, the latter so far having started at its N-terminal side with an evolutionarily conserved arginine, here colored in purple. For *Hs*BIRC1, the predicted structure corresponds to aa 59–129 instead of 63–128 for only the BIR signature of BIR1, 158–229 instead of 162–228 for BIR2, and 277–347 instead of 281–346 for BIR3. For *Hs*BIRC2, it is 45–115 instead of 49–114 for BIR1, 183–252 instead of 187–251 for BIR2, and 268–338 instead of 272–337 for BIR3. For *Hs*BIRC3, it is 28–98 instead of 32–97 for BIR1, 168–237 instead of 172–236 for BIR2, and 254-324 instead of 258–323 for BIR3. For *Hs*BIRC4, it is 25–95 instead of 29–94 for BIR1, 162-232 instead of 166–231 for BIR2, and 262–332 of 268–331 for BIR3. For *Hs*BIRC5, it is 14–89 instead of 18–88; for *Hs*BIRC6, 285–360 instead of 289–359; for *Hs*BIRC7, 86–156 instead of 90–155; for *Hs*BIRC8, 3–71 instead of 7–70. These extended versions of each BIR domain are here presented both individually and again superimposed onto each other, once again illustrating the domains' evident structural similarity. Note that the inclusion of additional aa residues at the N-terminal side of the BIR signature allowed for a prominent α-helix to become apparent in all of the BIR domains. Here colored in orange, we regard this α-helix as corresponding to the orange-colored α-helices of the BLDs shown in S10E2 and Figure 6C.

**(E2)** Comparison of the AlphaFold2-predicted structures of the *Hs*ZC3HC1 BLDs, as presented in Figure 6C and defined in Figure 6D, with the extended versions of the human BIR domains. Note that the orange-colored α-helix as part of both BLDs appears equivalent to the BIR domains' orange-colored α-helix, despite the latter being shorter than in the BLDs. Further note that this particular α-helix, positioned in the proximity of the BLDs' and BIR domains' G-[WF] dipeptide, possesses an arginine, again colored in purple, whose presence appears obligatory for both BLDs and as such equivalent to the BIR domains' arginine already high-lighted in purple in S10E1. Here in S10E2, these arginines are represented by R81 in BLD1, R185 in BLD2, and R18 in BIRC5. Each of these BLD and BIR domains' arginine residues appears involved in stabilizing the position of the neighboring α-helix that leads over to the BLDs' and BIR domains' first β-sheet.

Beyond such similarity between the BLDs and BIR domains, further note, though, that both BLD1 and BLD2 also possess BLD-specific α-helices, present in the one BLD while absent from the other BLD and from the BIR domains. These α-helices include the light pink-colored one of BLD1 and the yellow- and light blue-colored ones of BLD2. The functions of these single BLD-specific features will need to be dissected in future work. Such studies will also have to clarify which parts of the two BLDs engage in direct interactions with TPR as part of the ZC3HC1 protein's TPR-binding interface. Here, we already consider it conceivable that one or the other of each BLD's α-helices will have a penchant for parts of TPR's homodimeric coiled coils, as it will also be discussed from the TPR protein's viewpoint elsewhere (Gunkel *et al*, manuscript in preparation).

**(F)** The evolutionarily conserved arginines that are part of the orange-colored α-helices common to both BLDs are here shown enlarged and colored in purple. Other parts of the BLDs in these image sections are colored as in the preceding Figures. Chimera's structural analysis tool was used to compute and illustrate potential contacts between the conserved arginines' side chain atoms and neighboring residues. Several possible contacts, based on atom-to-atom distances being ≤ 4 Å, are depicted as dashed red lines. Note that among the residues predicted most likely involved in such intra-BLD contacts are the aromatic ones positioned two residues after each BLD's H-$X_{(3)}$-C pentapeptide (see also S10G). In the case of, e.g., *Hs*ZC3HC1, such intra-BLD residue contacts would be R81:W158 and R185:W431, with these aromatic residues also described in Supplemental Figure S9C, and corresponding ones here now shown also present in *Dd*ZC3Hc1 and *Sc*Pml39p. The contacts between one of these aromatic residues and the arginine's side chain provide the impression of a "lid", represented by either the tryptophan's indole ring or the tyrosine's phenol ring, which the arginine appears to contribute to keeping in place (in this context, see also Supplemental Figure S9C2). Apart from that, some of these arginines also appear to engage in additional intra-BLD interactions that would keep the orange-colored α-helix in a distinct position relative to the BLD core, just like some other residues of this α-helix too (not shown here). Again other residues of this α-helix, though, are predicted to contribute to the inter-BLD contacts of the BLD1:BLD2 interface described in Supplemental Figure S9B2. As an aside, also note that W158 and W431 of *Hs*ZC3HC1 were among the residues for which single aa substitutions were presented in the current study (see Supplemental Figure S2). Further note that for the BIR domains, the predictions do not position their orange-colored-α-helix-positioned arginines in direct contact with those aromatic residues located two aa after each BIR domain's H-$X_{(6)}$-C octapeptide that would be the ones equivalent to those in the BLDs. Nonetheless, in this case, the predictions indicate an indirect, i.e., two-step contact via a phenylalanine residue positioned between the two cysteines of the BIR domain's C-$X_{(2)}$-C tetrapeptide (not shown here, but see S10G for further details).

**(G)** Multiple aa sequence alignment of the human BIR domain sequences with BLD sequences of *Hs*ZC3HC1, *Dd*ZC3HC1, and *Sc*Pml39p, having first used for such purpose Clustal Omega (https://www.ebi.ac.uk/Tools/msa/clustalo/), followed by some manual readjustments. While the BIR domain sequences comprise those defined by the Pfam database's BIR motif in their entirety, plus some additionally flanking aa residues, the BLD sequences lack a few residues close to those BLDs' boundaries defined in Figure 6D. The BIR domains' N-terminal α-helix, which has been shown orange-colored in S10E, is here schematically depicted next to the corresponding BIR sequences, with this meant to facilitate correlating parts of the aligned sequences with the structural elements presented. Note that the identification of the *Hs*ZC3HC1 α-helix located between aa 176–190 and its assignment to BLD2, where it would thus be equivalent to the α-helix of BLD1 comprising at least aa 75–83 (Figure 6C and 6D), was followed by relating these α-helices with the BIR domains' orange-colored one. The latter, in turn, allowed for unveiling the here high-lighted arginine residue, as part of these α-helices, as another commonality of the BLD and BIR domains. More precisely, in the *Hs*ZC3HC1 BLD1, this arginine, R81, was already part of the Pfam database's zf-C3HC motif, just like R92 of *Dd*ZC3HC1. In addition, we now found such arginine evolutionarily conserved in *Sc*Pml39p too. Furthermore, we also found such an arginine at the corresponding position of the BLD2 domains, like, for example, the R185 as part of the BLD2 α-helix of *Hs*ZC3HC1. Such correspondence of R185 to R81 had escaped detection during all the alignments between segments of primary ZC3HC1 sequences conducted till then. Subsequent comparison of the BLD and BIR domain structures allowed us to correlate these BLD1 and BLD2 arginines with the arginine at the corresponding position of the BIR domain, like, for example, at R18 of *Hs*BIRC5, where it was already part of the Pfam database's BIR motif. Therefore, we regarded these findings as further supporting our conclusion that the orange-colored α-helices of BLD1 and BLD2 could now be considered equivalent to a likewise positioned shorter α-helix of the BIR domains. Moreover, this finding provided further justification for our new definition of the N-terminal boundary of BLD2, as presented in Figure 6D. As an aside, though, we also need to mention that not all ZC3HC1 homologs harbor such an arginine equivalent to R81 of *Hs*ZC3HC1, with it being absent in a few organisms, like, for example, in C49H3.9, the ZC3HC1 homolog noted for *C. elegans* (Higashi *et al*, 2005; Gunkel *et al*, 2021) and some other roundworms, while again present in other nematodes (our unpublished data). Furthermore, note that one of the connector lines, here colored in grey, marks the predicted direct contact between a BLD's orange-colored-α-helix-positioned arginine and an aromatic residue located two aa after the same BLD's H-$X_{(6)}$-C octapeptide. In the case of the BIR domains, two of such connector lines indicate an indirect contact between the BIR domains' corresponding residues via a phenylalanine residue that is part of the C-$X_{(2)}$-C tetrapeptide.

**(H)** HMM logos of the zf-C3HC and Rsm1 motifs, based on current seed sequences retrievable from the Pfam website. A selection of residues formerly (see Supplemental Figure S2D1) and currently regarded as evolutionarily relatively conserved are numbered corresponding to their position within the *Hs*ZC3HC1 protein sequence. Those conserved residues (bold lettering) that are predicted to be exposed and accessible on the surface of the BLDs are written in magenta, while those appearing only partially exposed outwardly are written in light blue. By contrast, those conserved residues embedded within the BLDs are written in black. In addition, *Hs*ZC3HC1 Y82 and W428, the latter evolutionarily somewhat less well conserved while contributing to the BLD1:BLD2 interface, are written in dark blue. Those conserved residues that are part of the BLDs' G-[WYF] dipeptides, C-$X_{(2)}$-C tetrapeptides, and H-$X_{(3)}$-C pentapeptides, are marked by brackets, as are those residues that are part of both BLDs' orange-colored α-helix. Even though this particular α-helix is not yet part of a current Pfam Rsm1 motif, we here assign the evolutionarily conserved R185 of *Hs*ZC3HC1 to this motif's N-terminal side (hatched rectangle). Moreover, other *Hs*ZC3HC1 residues that could be in contact with the here listed conserved residues' side chains are specified, having again used Chimera's structural analysis tool for computing potential contacts between side chains only, based on atom-to-atom distances being $\leq$ 4 Å. Note that only relatively few of the evolutionarily conserved residues are predicted to be exposed on the surface of either the one and other BLD, except for a cluster of conserved residues that are part of Pfam's Rsm1 motif, with these BLD2 residues located between the C-$X_{(2)}$-C tetrapeptide and the site where the additional, non-conserved sequence loops commonly are found inserted into this BLD. W282, as one of these conserved residues, has already been described in S10D as exposed on the surface of the *Hs*ZC2HC1 BLD2. While these residues and their exposed side chains will need to be a topic of a study to be presented elsewhere, note that the side chains of the majority of the other most conserved residues are predicted not exposed on the BLDs surface but located within the BLDs. With these zf-C3HC and Rsm1 motif residues also being part of the NUBaID signature's minimal versions, these findings allow us to conclude that the current NuBaID signature residues are not likely to engage in intermolecular interactions and thus are unlikely to represent residues of the Mlp- and TPR-binding interfaces.

**A**

*Hs*ZC3HC1 G-W C-X(2)-C H-X(3)-C deleted parts

```
MAAPCEGQAFAVGVEKNWGAVVRSPEGTPQKIRQLIDEGIAPEEGGVDAK   50
DTSATSQSVNGSPQAEQPSLESTSKEAFFSRVETFSSLKWAGKPFELSPL  100
VCAKYGWVTVECDMLKCSSCQAFLCASLQPAFDFDRYKQRCAELKKALCT  150
AHEKFCFWPDSPSPDRFGMLPLDEPAILVSEFLDRFQSLCHLDLQLPSLR  200
PEDLKTMCLTEDKISLLLHLLEDELDHRTDERKTTIKLGSDIQVHVTACI  250
LSVCGWACSSSLESMQLSLITCSQCMRKVGLWGFQQIESSMTDLDASFGL  300
TSSPIPGLEGRPERLPLVPESPRRMMTRSQDATFSPGSEQAEKSPGPIVS  350
RTRSWDSSSPVDRPEPEAASPTTRTRPVTRSMGTGDTPGLEVPSSPLRKA  400
KRARLCSSSSSDTSSRSFFDPTSQHRDWCPWVNITLGKESRENGGTEPDA  450
SAPAEPGWKAVLTILLAHKQSSQPAETDSMSLSEKSRKVFRIFRQWESLC  500
SC                                                  502
```

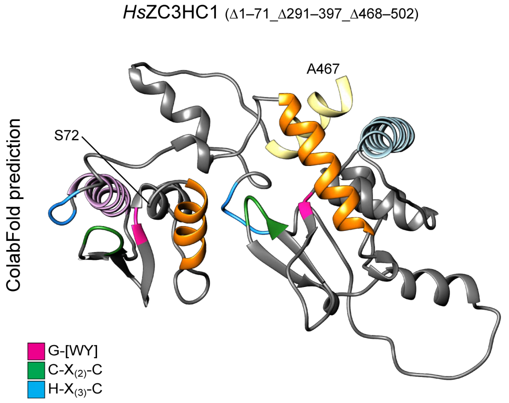

**B**

*Hs*ZC3HC1 G-[WY] C-X(2)-C H-X(3)-C deleted parts

```
MAAPCEGQAFAVGVEKNWGAVVRSPEGTPQKIRQLIDEGIAPEEGGVDAK   50
DTSATSQSVNGSPQAEQPSLESTSKEAFFSRVETFSSLKWAGKPFELSPL  100
VCAKYGWVTVECDMLKCSSCQAFLCASLQPAFDFDRYKQRCAELKKALCT  150
AHEKFCFWPDSPSPDRFGMLPLDEPAILVSEFLDRFQSLCHLDLQLPSLR  200
PEDLKTMCLTEDKISLLLHLLEDELDHRTDERKTTIKLGSDIQVHVTACI  250
LSVCGWACSSSLESMQLSLITCSQCMRKVGLWGFQQIESSMTDLDASFGL  300
TSSPIPGLEGRPERLPLVPESPRRMMTRSQDATFSPGSEQAEKSPGPIVS  350
RTRSWDSSSPVDRPEPEAASPTTRTRPVTRSMGTGDTPGLEVPSSPLRKA  400
KRARLCSSSSSDTSSRSFFDPTSQHRDWCPWVNITLGKESRENGGTEPDA  450
SAPAEPGWKAVLTILLAHKQSSQPAETDSMSLSEKSRKVFRIFRQWESLC  500
SC                                                  502
```

*Dd*ZC3HC1

```
MDERIKKALSDLDNATVLNQLPILSNDLTTTCGSSSGSSSNDNNNNNNKN   50
NNQYSTLNLIDESNNSTSNSTTSPSLLITSYRPWSNTDYYNRVRTYTISN  100
WFAKPCEIDPLQCSRFGWINCEADMLECETCKKRLYYKVPSTFSQSLVNK  150
RINDFSISLQSTGHRDNCPWKDNGCPSFFSRLLDIPFQTQLEAYIKRSQN  200
IYNNLTTLPMLSSDFYQQWVNKQNLMEPPITSRTNNILNIIVKIAKLPTD  250
EVKSKVSCLLALCGWDFNSISNSNNNNNNNNNNNNNNNNNNNNNNNNNNN  300
NNNNNNNNNNNNNNNNNNNNNNNNNNNNNDDDDKNEKDKNKNKIKENEKEKDKK  350
SSVYCSYCQRLCGVWNFNKIKPNSTSPFNEENINTTNNKGFNNSNIGSKR  400
KREEDIEEEKRNIQFEKVLNQSFARTNNNNNNNNNNNNNNNNNNNNNNNN  450
NNNNNNNNNNNNNNNNKKLNSNSNSNNNSNNNSNNNSLFSIVGNGFSKVTS  500
NSQSTGWDWGNSFSNRISDFKAALEIANATEKKKEFSPINEHRWFCPWMI  550
VVDSNRLIIDNNDILGENSQDNNNNSGSSNNSSNSNSSISGWENLLKLLL  600
NQSTFDSKDFIDLKNDKKFHSIVNSLTTSIHHYRK                 635
```

*Sc*Pml39p

```
MEKDALEVRLKSIRHSLDKNTKLLPGKYRNTLGERLITKWRYKKKSHNGS   50
SMLPEKCKSHVQLYDDLVQESSKHFVGFRLHDLRALLKRICSIQNYTRHV  100
LIEWDVRWVNPLTLASKGWEPYQSASQSQVPFKCCCCHAIMTIPLLKNGD  150
DVADYTMKLNEKIWNSNIIGNHLQKCPWRENQVDLNKEYYLSSQNLIREI  200
ERIHTEIDRIVSGSNEFSLKRNSSRIFHYLSEKEIQKLAFFFDCKDYSLV  250
GLLLLGYTKFQKDDLVQCTACFHRASLKKLEYTEFNGHALWCRYYNKELL  300
PTMLLELIGKEDKLITKLGVGERLNKLEAVLQTL                  334
```

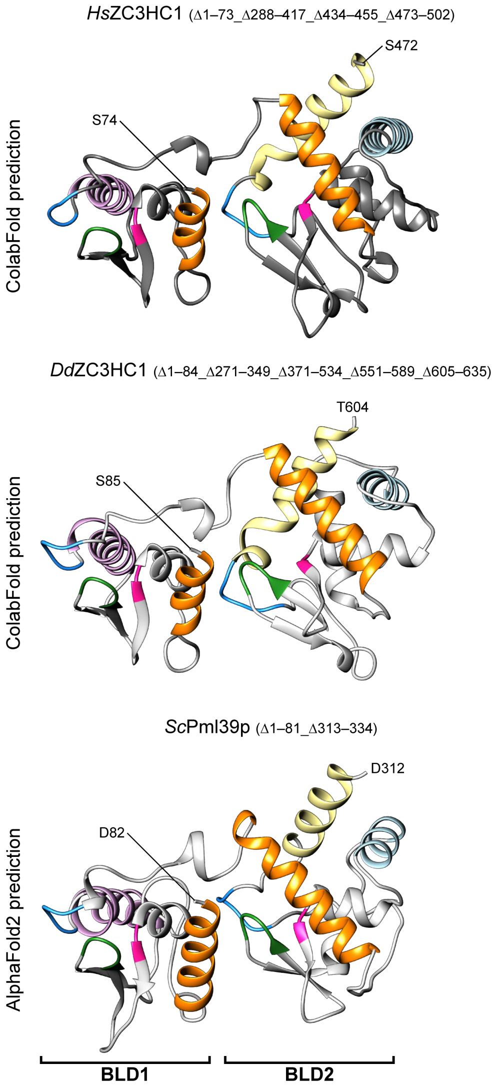

**Supplemental Figure S11. *In vivo* and *in silico* deletion mutants of ZC3HC1 homologs and their tertiary structures predicted by AlphaFold2 via ColabFold.**

**(A)** The sequence of the still NB-binding-competent *Hs*ZC3HC1 deletion mutant 72–290_398–467 and the structure for this residual sequence as predicted by AlphaFold2 via ColabFold.

Those regions highlighted in grey represent the residues removed by cloning and thus excluded also from this structure prediction. Residues highlighted in magenta, green, and blue again represent the positions of the two BLDs' G-W, C-$X_{(2)}$-C, and H-$X_{(3)}$-C sequence elements. The coloring of the α-helices corresponds to the same helices' colors in Figure 6C and Supplemental Figures S7 and S10. Note that the structure predicted for this mutant, comprising only 289 aa and mainly encompassing the two BLDs of *Hs*ZC3HC1, resembles a compact version of a ZC3HC1 homolog, reminiscent, e.g., of the central parts of *Sc*Pml39p with its two BLDs, with its second one lacking large sequence insertions. Such prediction further underscored the conclusion that the integrity of the two BLDs is essential and sufficient for the initial binding of *Hs*ZC3HC1 to the NB.

**(B)** The sequences of *in silico*-created deletion mutants of *Hs*ZC3HC1, *Dd*ZC3HC1, and *Sc*Pml39p and the corresponding structures predicted by either AlphaFold2 or via ColabFold. The largely "loop-free" versions of each homolog here presented, primarily comprising the BLD-corresponding sequences according to the newly defined BLD boundaries, include a mutant, only 247 aa long, of *Hs*ZC3HC1 (Δ1–73_Δ288–417_Δ434–455_Δ473–502) and a 238 aa-long *Dd*ZC3HC1 mutant (Δ1–84_Δ271–349_Δ371–534_Δ551–589_Δ605–635). In addition, the naturally "loop-less" *Sc*Pml39p, which here though lacks its N- and C-terminal parts, thus representing an *in silico* mutant of *Sc*Pml39p only comprising 231 aa (Δ1–81_Δ313–334), is shown for comparison. Note that all three deletion mutants represent polypeptides of about similar length. Further note, in particular, that despite the still relatively low end-to-end sequence identities between the three mutants' sequences (see below), their predicted structures appear, not unexpected, once again very similar, with each homolog's two BLDs as separate yet closely abutting entities now particularly evident.

Having removed, as outlined above, the sequences from the BLD2 domains that represented the larger ones of the loop-like insertions, next to all BLD-flanking N- and C-terminal sequences, some standard sequence alignment tools, even then, did not find any significant similarity between the residual 247 aa-long sequence of *Hs*ZC3HC1 Δ1–73_Δ288–417_Δ434–455_Δ473–502 and the residual 231 aa of Pml39p Δ1–81_Δ313–334. Again other tools that aligned these sequences almost correctly, like the EMBOSS local alignment program Water (https://www.ebi.ac.uk/Tools/psa/emboss_water/) and the end-to-end global alignment tools Needle (https://www.ebi.ac.uk/Tools/psa/emboss_needle/) and GGSEARCH2SEQ (https://www.ebi.ac.uk/Tools/psa/ggsearch2seq/), only yielded end-to-end sequence identities of about 16.5–18.4%. Moreover, even some of these mentioned tools did not allow for a proper pairwise alignment when attempting to match the residual 238 aa-long, largely "loop-free" and primarily

BLD-corresponding sequence of *Dd*ZC3HC1 Δ1–84_Δ271–349_Δ371–534_Δ551–589_Δ605–635 with that of Pml39p Δ1–81_Δ313–334. Again others, like the GGSEARCH2SEQ program, provided an almost correct alignment of the two homologs' NuBaID signatures but with an end-to-end sequence identity again of only 17.3%.

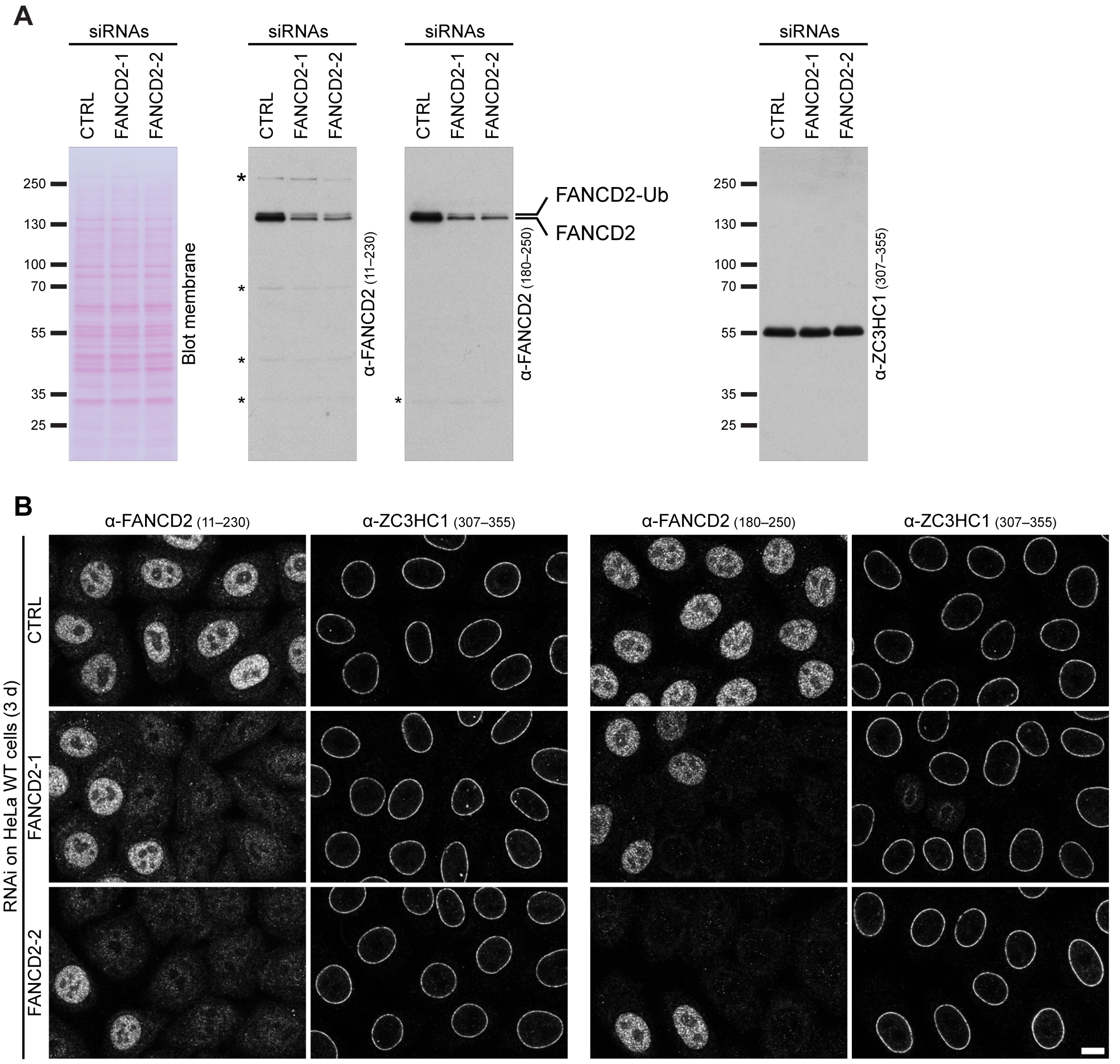
A
siRNAs
CTRL
FANCD2-1
FANCD2-2
250
130
100
70
55
35
25
Blot membrane
α-FANCD2 (11–230)
α-FANCD2 (180–250)
FANCD2-Ub
FANCD2
α-ZC3HC1 (307–355)
B
α-FANCD2 (11–230)
α-ZC3HC1 (307–355)
α-FANCD2 (180–250)
α-ZC3HC1 (307–355)
RNAi on HeLa WT cells (3 d)
CTRL
FANCD2-1
FANCD2-2

**S12 (1/3)**

C

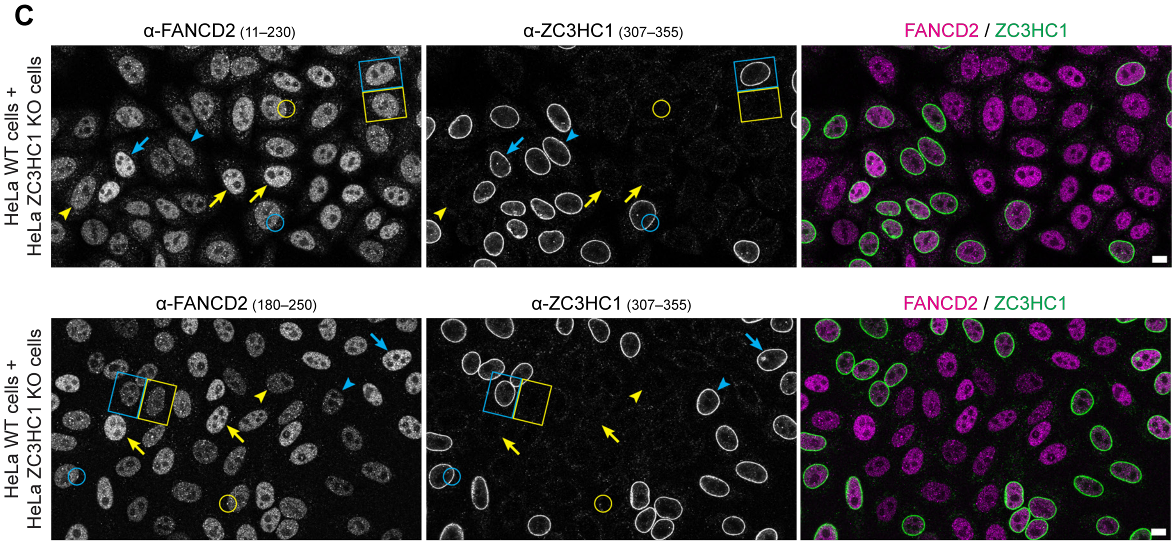
α-FANCD2 (11–230)
α-ZC3HC1 (307–355)
FANCD2 / ZC3HC1
α-FANCD2 (180–250)
α-ZC3HC1 (307–355)
FANCD2 / ZC3HC1
HeLa WT cells +
HeLa ZC3HC1 KO cells
HeLa WT cells +
HeLa ZC3HC1 KO cells

D

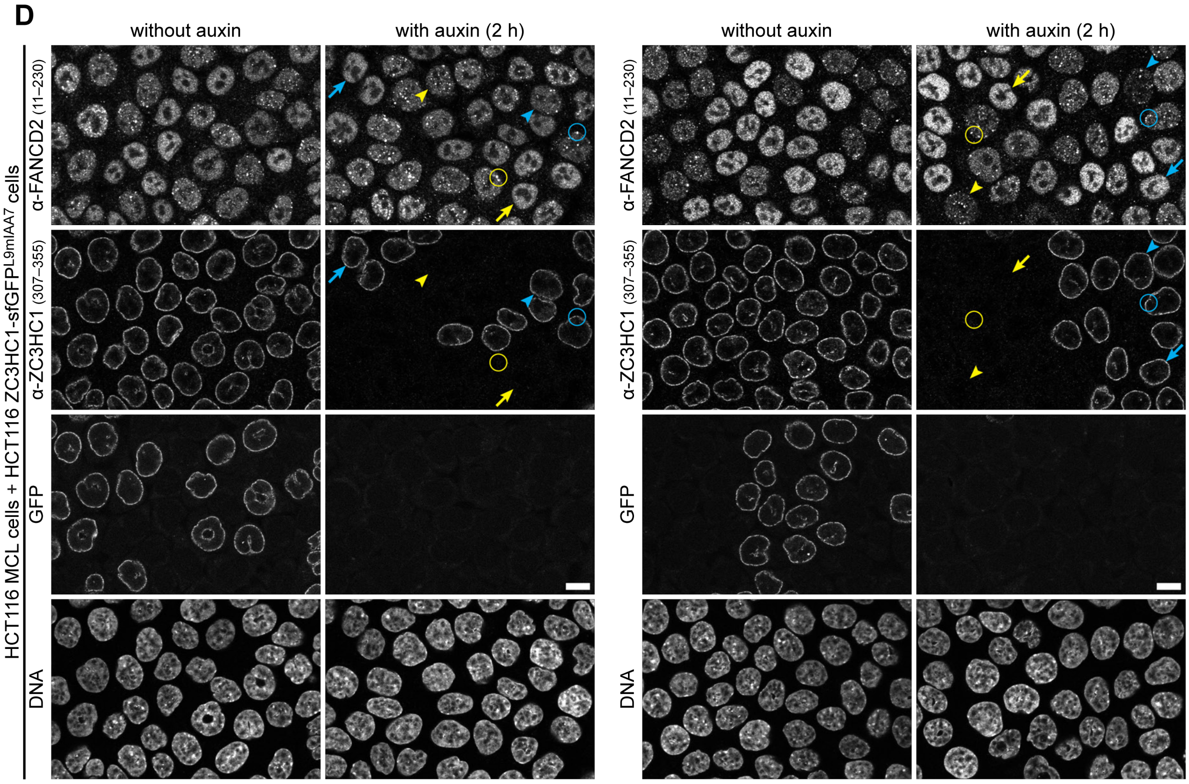
without auxin
with auxin (2 h)
without auxin
with auxin (2 h)
HCT116 MCL cells + HCT116 ZC3HC1-sfGFPL9mIAA7 cells
α-FANCD2 (11–230)
α-ZC3HC1 (307–355)
GFP
DNA
α-FANCD2 (11–230)
α-ZC3HC1 (307–355)
GFP
DNA

S12 (2/3)

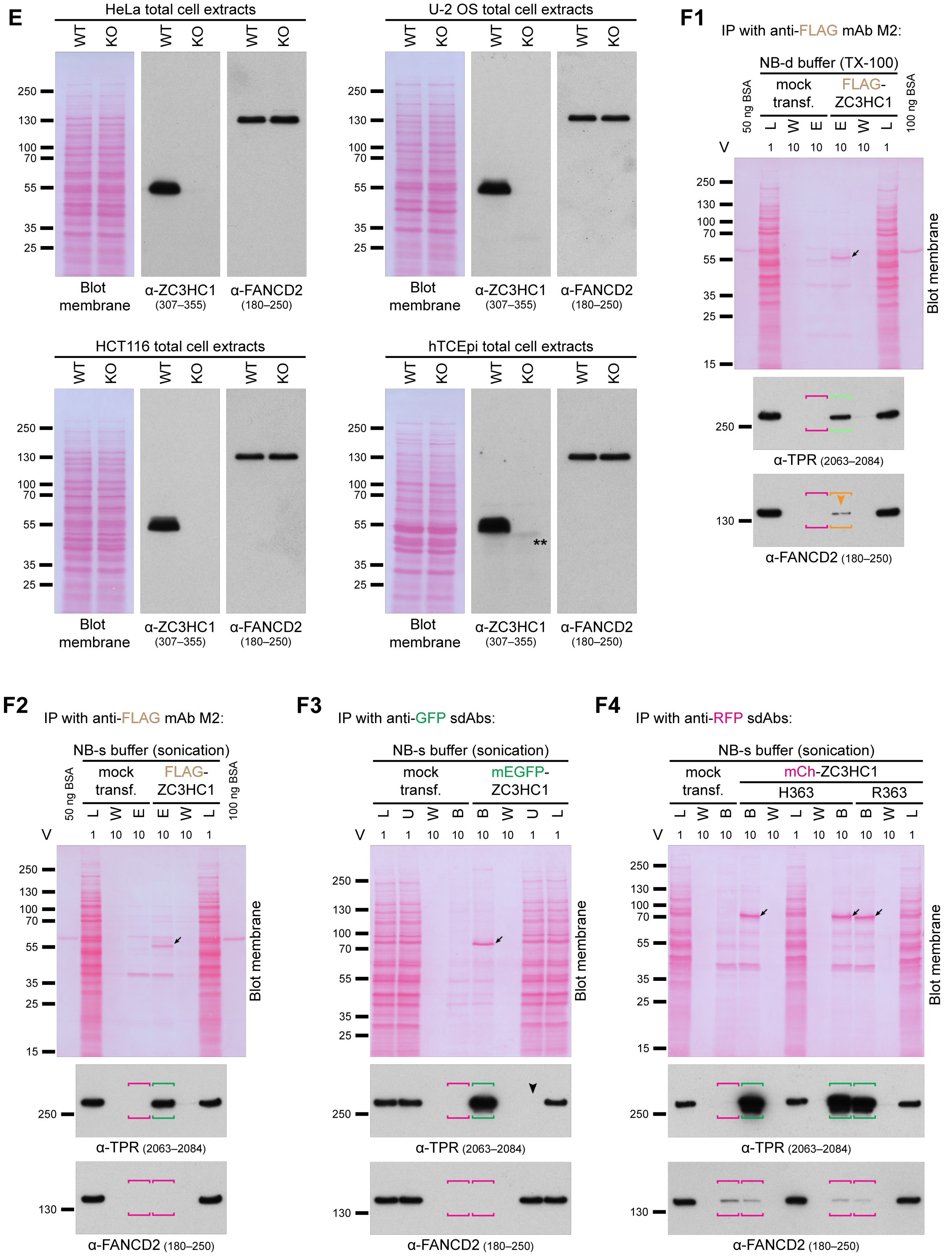

**Supplemental Figure S12. ZC3HC1 deficiency in human cell lines not affecting the cellular amounts and subcellular distribution of FANCD2, in line with no evident robust interaction between ZC3HC1 and FANCD2 at the NE or in cell extracts.**

In a recent study, FANCD2, a protein of 164 kD in humans, has been described as a binding partner of NIPA (Kreutmair *et al*, 2020), the latter protein also known as ZC3HC1. Among this study's data, ZC3HC1 knockdown in HeLa cells has been presented to cause a significant reduction of FANCD2 protein levels in total cell extracts, as demonstrated by IB, and in the HeLa cells' nuclei, as shown by IFM. Moreover, immunoprecipitation (IP) of ectopically expressed, FLAG-tagged ZC3HC1 from HEK293T Phoenix cells has been reported resulting in co-IP of endogenous FANCD2, then regarded as representing a robust ZC3HC1:FANCD2 interaction (Kreutmair *et al*, 2020). The authors considered ZC3HC1 necessary for FANCD2 protein stability and half-life, with ZC3HC1 acting as a scaffold protein for FANCD2 and stabilizing its nuclear abundance.

On the other hand, we had till then not encountered evidence indicating an interaction between ZC3HC1 and FANCD2 in our studies. We had neither found FANCD2 a component of manually isolated and mass spectrometrically analyzed *Xenopus laevis* and *Xenopus tropicalis* oocyte NEs, even though database-deposited *Xenopus* FANCD2 sequences were already available at the time. Therefore, we had also not detected this protein among those co-detached together with *Xenopus* ZC3HC1 and other proteins after having disassembled the oocytes' NEs by physicochemical means (Gunkel *et al*, 2021, and our unpublished data). Similarly, we had neither found FANCD2 among those proteins that one could identify, via comparative proteomics, to have been detached from the NEs of CRISPR/Cas9-edited human cell lines expressing degron-tagged versions of TPR after having, and not having, auxin-induced the degradation of TPR (Gunkel & Cordes, 2022, and our unpublished data). Furthermore, following the IP of FP-tagged versions of *Hs*ZC3HC1, protein FANCD2 had not been detectable by mass spectrometry among those materials co-sedimented with ZC3HC1 (our unpublished data).

However, to assess whether we might have overlooked a ZC3HC1:FANCD2 interaction, we addressed this question in a more detailed manner. To this end, we used the two commercial FANCD2 antibodies (S12A and S12B) that had also been employed in the recent study on ZC3HC1 and FANCD2 (Kreutmair *et al*, 2020). With these antibodies, we investigated the fate of FANCD2 in human cell lines, including HeLa P2, HCT116, U-2 OS, and hTCEpi (for cell line details, see Gunkel *et al*, 2021) in the absence of ZC3HC1 (e.g., S12C to S12E). Furthermore, we investigated whether the IP of ectopically expressed FLAG-tagged and FP-tagged versions of ZC3HC1 from HEK293T cells would allow for detecting co-immunoprecipitated FANCD2 (S12F). In parallel, we also addressed whether the absence of

FANCD2 itself might affect the cellular amounts and the positioning of ZC3HC1 at the NB (S12A and S12B).

In the course of these experiments, we applied a range of different IFM and IP protocols, including those used earlier (Kreutmair *et al*, 2020; Gunkel *et al*, 2021). For some of the FANCD2 immunoblots (S12E and S12F), we eventually decided to use the same cellular materials and nitrocellulose membranes (Gunkel *et al*, 2021) that we had already used for investigating the interaction between ZC3HC1 and TPR and between ZC3HC1 and other proteins formerly reported binding partners of ZC3HC1. These materials thus allowed for directly comparing the here presented data with those we had obtained earlier. Furthermore, since the underlying rationale for our former experiments and their procedural details, followed by an in-depth discussion of the conclusions to be drawn from such experiments' results, had already been outlined in all detail (Gunkel *et al*, 2021), this further allowed for here confining oneself to a brief description of the experimental specifics and the here obtained data related to FANCD2. A representative selection of these data is presented in the following.

**(A)** IB of whole-protein extracts from HeLa P2 cells that had been transfected with control siRNAs (CTRL; Ambion Silencer Select negative control #2, Thermo Fisher Scientific, Waltham, MA, USA; cat. no. 4390846), and two pairs (FANCD2-1 and FANCD2-2) of FANCD2 siRNAs (Ambion Silencer Select siRNAs s4988 [GCACCGUAUUCAAGUACAA] and s4889 [CAGCCUACCUGAGAUCCUA], Thermo Fisher Scientific). Rabbit polyclonal antibodies against *Hs*FANCD2, raised against aa 11–230, (NB100-182, Novus Biologicals, Abingdon, UK) and rabbit monoclonal antibody EPR2302 against an *Hs*FANCD2 epitope located between aa 180–250 (ab108928, Abcam, Cambridge, UK), had already been verified to target FANCD2 by using FANCD2 KO cells, according to the suppliers' information. The RNAi experiments presented here in S12A were conducted to confirm FANCD2 antibody performance and to assess whether a reduction in the cellular amounts of FANCD2 might also affect those of ZC3HC1. The guinea pig antibodies against *Hs*ZC3HC1 have been described earlier (Gunkel *et al*, 2021). Immunolabeling for the three immunoblots shown on the right was performed on the Ponceau S-stained uncut membrane shown here on the left and on an identical duplicate, comprising the entire length of gel-electrophoretic sample separation, in order to also illustrate each of the FANCD2 antibodies' degree of target-specificity, apart from their target-verification by RNAi. The membranes were first incubated with the FANCD2 antibodies, then recovered by quantitatively detaching the bound antibodies through incubation at low pH, and then re-incubated with the ZC3HC1 antibodies. Asterisks mark minor cross-reactions with yet unknown polypeptides. The band marked as FANCD2-Ub represents a minor subpopulation of mono-ubiquitinated FANCD2 polypeptides (e.g., Vandenberg *et al*, 2003). Note that the reduction in the total cellular amounts of FANCD2 did not appear to have affected the cellular amounts of ZC3HC1. As an aside, IB of total cell extracts from HeLa cells treated with ZC3HC1 siRNAs, resulting in a substantial KD in total ZC3HC1 amounts, did not have a notable effect on the cellular amounts of FANCD2 either (our unpublished data), with these findings equivalent to those with cell extracts of ZC3HC1 KO cells, presented further below in S12E.

**(B)** Double-labeling IFM of ZC3HC1 and FANCD2 in HeLa P2 cells. The cells had been treated with either control siRNAs or the two different pairs of siRNAs targeting FANCD2 and then harvested on day 3 post-transfection. FANCD2 was detected with the rabbit antibodies presented in S12A and ZC3HC1 with formerly described *Hs*ZC3HC1-specific guinea pig antibodies (Gunkel *et al*, 2021), which here have been used for S12A, too. Some cells not transfected with the FANCD2 siRNAs are shown as a reference. Note the characteristic dotted staining for FANCD2 throughout the nuclear interior, except for the nucleoli, in the control cells. By contrast, in the cell populations treated with the FANCD2 siRNAs, such staining is only visible in the subpopulation of cells that apparently had remained untransfected. Also, note that knockdown of FANCD2 had not affected the presence and immunolabeling intensity of ZC3HC1 at the NE. Bar, 10 µm.

**(C)** IFM of asynchronous HeLa P2 WT cells grown together with cells of a stable HeLa cell line in which all *ZC3HC1* alleles have been disrupted by CRISPR/Cas9n technology. Allowing for a side-by-side comparison of the WT and ZC3HC1 KO cells on the same coverslip, the mixed cell populations were double-immunolabeled with the FANCD2 and ZC3HC1 antibodies also used for S12A. Some HeLa WT cells are marked by blue arrows and arrowheads, while arrows and arrowheads in yellow mark some ZC3HC1 KO cells. The arrows mark those cells in which the signal intensities for nuclear FANCD2 appear especially pronounced, whilst the arrowheads mark some representative cells with less intense nuclear FANCD2 immunolabeling. Furthermore, blue and yellow circles mark some similarly conspicuous FANCD2 foci at the NEs of both the WT and ZC3HC1 KO cells. In addition, pairs of rectangles, again in blue and yellow, mark some cell pairs, consisting of one WT and one neighboring ZC3HC1 KO cell, in which nuclear FANCD2 levels appear very similar. These arrows, arrowheads, circles, and rectangles are also included same-positioned on those micrographs that show the ZC3HC1 immunolabeling, allowing for directly comparing FANCD2 with ZC3HC1. Furthermore, the same FANCD2 and ZC3HC1 micrographs are also shown as overlays on the right side.

Note that, when regarding these mixed populations of WT and ZC3HC1 KO cells as a whole, neither the cellular amounts nor the subcellular location of FANCD2 appeared altered by the absence of ZC3HC1, with immunolabeling of FANCD2 varying between individual WT cells to the same extent as between individual ZC3HC1 KO cells, accompanied by FANCD2-containing NE-associated foci being similarly evident in the presence and absence of ZC3HC1. As an aside, when using a formerly described protocol for cell fixation and permeabilization (Kreutmair *et al*, 2020), we could not detect any evident difference between the WT and ZC3HC1 KO cells of a mixed population of cells immunolabeled for FANCD2 in such a way either (our unpublished data). Furthermore, we could also not detect any evident difference following ZC3HC1 RNAi in HeLa WT cells, when comparing the siRNA-transfected cells with (i) those cells that had remained non-transfected within the same population and with (ii) a separate population of cells that we had transfected with control siRNAs (our unpublished data). Bars, 10 µm.

**(D)** IFM of cells from an HCT116 progenitor cell line expressing the naturally tag-free ZC3HC1 and cells of a homozygous HCT116 progeny line, in which all ZC3HC1 polypeptides were C-terminally tagged with a GFP-degron tag, called sfGFP$^{L9mIAA7}$ (for details regarding these cell lines and the sfGFP$^{L9mIAA7}$-tag, see Gunkel & Cordes, 2022). These cells had been co-cultured as mixed populations together on the same coverslip, followed by an additional incubation of 2h in the absence or presence of auxin, the latter inducing the rapid degradation of proteasomal degradation of ZC3HC1 (for details, see Gunkel & Cordes, 2022). Specimens were then double-immunolabeled for ZC3HC1 and FANCD2 and analyzed in parallel, using identical microscope settings. Assigned to the same features as in S12C, the blue arrows, arrowheads and circles mark the nuclei of some progenitor cells, i.e. those expressing the tag-free version of ZC3HC1, while the yellow arrows, arrowheads and circles mark some of the progeny cells' nuclei, i.e. those with the sfGFP-tagged ZC3HC1. Note that the auxin-treatment resulted in the elimination of NE-associated GFP and immunostaining for ZC3HC1 in those cells that had been expressing the tagged version of ZC3HC1, while the progenitor cells' untagged ZC3HC1 remained unaffected. Further note that the elimination of the tagged ZC3HC1 neither affected the nuclear amounts nor the subcellular distribution of FANCD2 notably. In particular, note that also upon the rapid loss of ZC3HC1, the more finely punctate nuclear staining for FANCD2 remained largely unchanged, with no evident more diffuse distribution throughout the nuclear interior, in contrast to what a model might have expected in which ZC3HC1 would act as a FANCD2-protecting scaffold. Bars, 10 µm

**(E)** IB of total cell extracts from HeLa P2, HCT116, U-2 OS, and hTCEpi WT and ZC3HC1 KO cells. The Ponceau S-labeled membranes and the IBs for ZC3HC1 are identical to those already presented earlier (see Figures 6B and S15H in Gunkel *et al*, 2021). The double asterisk

marks a cross-reaction of the polyclonal guinea-pig ZC3HC1 antibodies, just beneath the band for ZC3HC1, which was only seen in hTCEpi cell extracts and which also arose when only using secondary antibodies (for further details, see S15I in Gunkel *et al*, 2021). Here, these membranes were recovered by quantitatively detaching the bound ZC3HC1 antibodies through incubation at low pH, followed by re-incubating such membranes with rabbit antibodies for FANCD2. Note that while ZC3HC1 had not been detectable in any of the KO cells' extracts, with each cell type's pair of KO and WT cell extracts representing about the same number of cells, the cellular amounts of FANCD2 appeared highly similar in the WT and KO cells of each cell line. Altogether, these data demonstrated that the integrity of FANCD2 in three aneuploid tumor cell lines of different tissue origins (HeLa, HCT116, and U-2 OS) and a non-tumor cell line of normal diploid karyotype (hTCEpi) does not depend on ZC3HC1 being present.
**(F)** A representative selection of IP experiments that had been performed with HEK293T cell extracts containing differently tagged versions of ectopically expressed intact *Hs*ZC3HC1. The Ponceau S-labeled membranes and the IBs for TPR presented in S12F1–4 are identical to those already presented earlier (see Figure S11D2 and S11D3 in Gunkel *et al*, 2021). Here, these membranes were recovered by quantitatively detaching the bound TPR antibodies through incubation at low pH, followed by re-incubating such membranes with rabbit antibodies for FANCD2. Buffers used for the actual IP experiments included such that we had found to result in the destabilization of the nuclear basket (NB-d buffers), with the composition of these buffers, and the mode of their application, being notably different in several aspects from the physicochemical conditions within the living mammalian cell. The composition of the NB-d buffer for the here presented IP experiment in S12F1 lacked, for example, Mg cations while containing high concentrations, i.e. 1%, of Triton X-100. As such, this buffer and the corresponding IP protocol (for further details, see Gunkel *et al*, 2021) were representatives of very similar or essentially identical IP buffers and protocols used for earlier studies on ZC3HC1 and its alleged interactions with other proteins (e.g., Bassermann *et al*, 2005a, 2007; Illert *et al*, 2012; Kreutmair *et al*, 2020). Other buffers and protocols used for our IP experiments allowed for maintaining the NB's integrity, with the corresponding buffers termed NB-stabilizing (NB-s; Gunkel *et al*, 2021). Of note, while the use of the NB-d buffers only resulted in minor amounts of TPR being co-immunoprecipitated upon IP of tagged versions of ZC3HC1, as exemplified in S12F1, the NB-s buffers, more closely resembling the physiological conditions within the cell, often allowed for a quantitative co-IP of TPR. Figure S12F3 exemplifies such removal of all soluble TPR from the cell extract due to co-IP with the immunoprecipitated ZC3HC1 (for further details, again see Gunkel *et al*, 2021). By contrast, in none of these and other buffers, in

any combination with different IP protocols, did we find FANCD2 co-immunoprecipitated in amounts that we would regard as significant, as outlined further below.

**(F1)** IB of materials obtained from an IP experiment with anti-FLAG IgG-coated immuno-magnetic beads, following ectopic expression of FLAG-tagged ZC3HC1, and subsequent cell extract preparation and incubation under NB-d conditions (for details, see the Supplemental Material and Methods section of Gunkel *et al*, 2021). Lanes had been loaded for SDS-PAGE with an aliquot of the total soluble cell proteins not yet treated with the magnetic immunoaffinity beads (L, for load), with an aliquot of those materials released during the third of three successive washing steps (W), and with the proteins obtained after final elution (E). The arrow on the image of the Ponceau S-stained membrane shown here, like on those presented in S12F2–4, marks the immunoprecipitated tagged ZC3HC1 polypeptides. Loadings in L represented one volume fraction of the respective samples' total amount (1 V), while the loadings in lanes W and E represented ten-fold higher relative amounts (10 V). TPR regarded as inefficiently co-immunoprecipitated with the FLAG-tagged ZC3HC1, as the IP's actual target protein, is framed with brackets in light green, while those cases in which no co-IP had occurred, like upon incubating FLAG-ZC3HC1-deficient cell extracts with the anti-FLAG IgG-coated immuno-magnetic beads, are accentuated by brackets in magenta. The orange-colored brackets frame some trace amounts of FANCD2, also marked by an arrowhead. Note that trace amounts like those seen here in S12F1 represent what one had been able to detect at most in such kinds of experiments relative to the total FANCD2 amounts in the corresponding cell extracts. Even though one might consider such trace amounts of co-sedimented FANCD2 as having been co-immunoprecipitated with the FLAG-tagged ZC3HC1 specifically, one needs to look at the actual numbers of these FANCD2 polypeptides in the context with the numbers of the ectopically expressed ZC3HC1 polypeptides; the latter determined to be present in millions, on average, within a transfected cell at the time of harvest (Gunkel *et al*, 2021). In fact, the availability of quantitative mass spectrometric data for HEK293 cells (Bekker-Jensen *et al*, 2017), paired with the knowledge of the absolute copy numbers within HEK293 cells for some representative NPC proteins, allowed us to deduce the HEK293T cell's approximate total number of FANCD2 polypeptides and thus also the approximate number of the few FANCD2 polypeptides co-sedimented together with FLAG-ZC3HC1. This information, in turn, allowed us to conclude that, at most, only a few hundred FANCD2 polypeptides per transfected cell had been co-sedimented during the IP of the same cell's millions of ectopically expressed ZC3HC1 polypeptides. In other words, for every about 10,000 FLAG-ZC3HC1 polypeptides immunoprecipitated, only one endogenous FANCD2 polypeptide had been co-sedimented.

**(F2)** IB of materials obtained from an IP experiment with anti-FLAG IgG-coated immuno-magnetic beads, following ectopic expression of FLAG-tagged ZC3HC1, subsequent cell extract preparation, and incubation for IP under NB-s conditions more closely resembling the physicochemical conditions within the cells' cytoplasm and nucleoplasm (for further details, also regarding cell extract preparation, see the Supplemental Material and Methods section of Gunkel *et al*, 2021). While not yet reaching the quantitative co-IP of TPR achievable via "nano-trapping", i.e. when using single-domain antibodies (sdAbs) in combination with NB-s buffers (see S12F3 and S12F4 below), these NB-s conditions already allowed for recurrently co-immunoprecipitating notably higher TPR amounts, here now framed by dark green brackets, when immunoprecipitating FLAG-ZC3HC1, as compared to IPs conducted under NB-d conditions, like in S12F1. Further note, in particular, that no co-IP of FANCD2 was detected.

**(F3)** IB of materials obtained from IP experiments with anti-GFP sdAb-coated agarose beads, after ectopically having expressed a monomeric EGFP-tagged version of ZC3HC1, followed by cell extract preparation and incubation for IP under NB-s conditions. Like in S12F1 and S12F2, lanes had been loaded for SDS-PAGE with an aliquot of the total soluble cell proteins not yet treated with the sdAb-coated immunoaffinity beads (L), with an aliquot of those materials released during the third of three successive washing steps (W), and with the proteins obtained after final elution (E). In addition, we had here also loaded the proteins that had remained unbound after incubation with such beads (U). Loadings in L and U represented one volume fraction of the respective samples' total amount (1 V), while the loadings in lanes W and E represented ten-fold higher relative amounts (10 V). Note that these IP conditions had allowed for quantitative co-IP of all soluble TPR polypeptides initially present in interphase cell extracts (L), which were then absent from such extracts (position marked by arrowhead) after the incubation (U) with the anti-GFP beads. By striking contrast, no co-IP of FANCD2 was detected.

**(F4)** IB of materials obtained from IP experiments with anti-RFP sdAb-coated agarose beads, after ectopically having expressed two versions of ZC3HC1, namely the H363 and the R363 variants, here both tagged with mCherry. Note that both ZC3HC1 variants allowed for quantitative co-IP of TPR, while trace amounts of FANCD2 were in this experiment even co-sedimented with the sdAb-coated agarose beads after their incubation with cell extracts lacking ectopically expressed ZC3HC1.

Finally, not having intentionally stressed the cells for our FANCD2-related experiments, we need to remark that our current results do not yet permit excluding scenarios in which FANCD2 might engage in stress-induced interactions with some structures at the NE. For example, one might conceive such interactions as a result of DNA replication stress or DNA damage-induced stresses and localization of DNA lesions to the NPC (e.g., Freudenreich & Su, 2016; Lamm *et al*, 2021; Whalen & Freudenreich, 2020). However, based on our current FANCD2 results, including those presented here in S12, we regard it justified to conclude that FANCD2 is not a regular, customary binding partner of ZC3HC1 in different human cell lines, neither at the NE nor elsewhere within such cells in interphase, and that ZC3HC1 does not function as a regular scaffold protein for FANCD2.

A

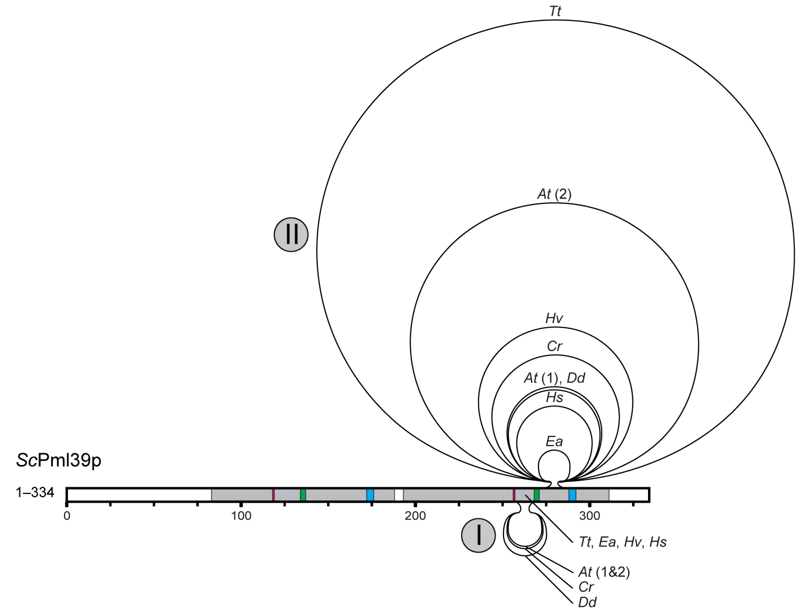

| Species | | total length (aa) | accession number |
|---|---|---|---|
| *Saccharomyces cerevisiae* | M-[117]-GW-[14]-CCCC-[34]-HLQKCPW-[_77]-GY-[10]-CTAC-[_16]-HALWCRY-[_40] | 334 | NP_013600.2 |
| *Elsinoe ampelina* | M-[138]-GW-[_9]-CKGC-[52]-HSESCPW-[101]-GW-[12]-CDAC-[_69]-HREHCPY-[103] | 511 | KAF2227358.1 |
| *Homo sapiens* | M-[104]-GW-[_9]-CSSC-[31]-HEKFCFW-[_96]-GW-[15]-CSQC-[149]-HRDWCPW-[_71] | 502 | NP_057562.3 |
| *Arabidopsis thaliana* (1) | M-[113]-GW-[_9]-CESC-[31]-HKLLCPW-[100]-GW-[72]-CKLC-[178]-HRHFCPW-[_64] | 594 | NP_175325.2 |
| *Dictyostelium discoideum* | M-[115]-GW-[_9]-CETC-[32]-HRDNCPW-[_93]-GW-[89]-CSYC-[183]-HRWFCPW-[_87] | 635 | ON368701 |
| *Chlamydomonas reinhardtii* | M-[_90]-GW-[_9]-CEYC-[29]-HTATCPW-[172]-GW-[76]-CPIC-[243]-HRSWCPW-[_62] | 708 | PNW78929.1 |
| *Hydra vulgaris* | M-[_65]-GW-[_9]-CVTC-[31]-HEKLCPW-[_87]-GW-[13]-CTIC-[289]-HRFWCPW-[_69] | 590 | XP_002154618.2 |
| *Arabidopsis thaliana* (2) | M-[115]-GW-[_9]-CEYC-[32]-HESSCPW-[_98]-GW-[72]-CSLC-[524]-HNCYCPW-[_81] | 958 | NP_173164.1 |
| *Timema tahoe* | M-[_66]-GW-[10]-CTSC-[23]-HYKFCRW-[_75]-GW-[10]-CDFC-[836]-HRYWCIW-[171] | 1217 | CAD7459619.1 |

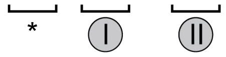

B

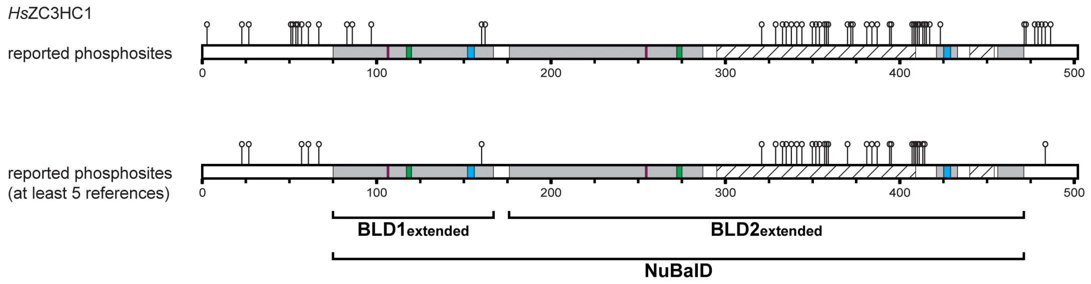

**Supplemental Figure S13. Size variations of loop-like sequence insertions within exemplary ZC3HC1 homologs, and the BLD2 loop of *Hs*ZC3HC1 as a prime target for phosphorylation.**

**(A)** Schematic depiction of the two BLDs of *S. cerevisiae* Pml39p as a representative of one of the shortest ZC3HC1 homologs. The areas highlighted correspond to those already described in Figure 6D, with the boxes in green and blue, respectively, representing the positions of the C-$X_{(2)}$-C and H-$X_{(3)}$-C sequence elements, and the boxes in magenta depicting the position of the evolutionarily conserved dipeptide that in *Sc*Pml39p either reads G-W or G-Y. In addition, loops are shown inserted at positions I and II, where they can be found in other exemplary ZC3HC1 homologs, with the small selection chosen here comprising the ones from *Elsinoe ampelina* (*Ea*), Homo sapiens (*Hs*), *Arabidopsis thaliana* (*At*) with its two homologs, *Dictyostelium discoideum* (*Dd*), *Chlamydomonas reinhardtii* (*Cr*), *Hydra vulgaris* (*Hv*), and *Timema tahoe* (*Tt*; see also Supplemental List of Sequences for ZC3HC1 Homologs). The lengths of these species' loops are drawn approximately to scale relative to the length of the *Sc*Pml39p scheme. The alignment of the homologs' corresponding minimal NuBaID sequence signature, including the G-W, C-$X_{(2)}$-C, and H-$X_{(3)}$-C peptides, and the number of residues located between them, next to each homolog's accession number, is provided for comparison. In contrast to loops I and II that are predicted, in numerous cases, to be essentially unstructured in their entirety, other sequence insertions, like those collectively marked with an asterisk, can encompass both structured parts and additional α-helices present in some species while absent in others. Further note that loops inserted at position II appear to be especially large in some species of the insect order Orthoptera, like possibly in *Gryllus bimaculatus* (not shown here) and, in particular, in the order Phasmatodea. Protein sequences so far deduced and assembled from whole genome shotgun contigs (WGS) for a range of species of the phasmid genus *Timena*, and for species of other stick insect genera, like *Clitarchus hookeri*, *Medauroidea extradentata* and *Dryococelus australis* (not listed in the Supplemental List of Sequences, except for *M. extradentata*), are currently indicating a possible range of 496 to 994 aa located between the phasmids' second BLD's C-$X_{(2)}$-C, and H-$X_{(3)}$-C peptides.

As evident, a necessity for flexible linkers cannot explain the large loops' persistence in a wide range of species since short flexible stretches like those located between the different structural elements of the BLDs of *Sc*Pml39p would suffice as the unstructured elements that all ZC3HC1 homologs would require for allowing their NuBaIDs to fold into their final conformation, with inter-BLD interactions at the BLD1:BLD2 interface. Therefore, with evolution having tolerated such ZC3HC1 loops evolving in numerous species, we are currently wondering whether and which tasks there might be at the NB for such loops to fulfil, with demand and possibly accompanying antagonistic properties then varying between different species.

The existence of huge loops in only a few insect orders, while ZC3HC1 homologs appear to have been lost altogether in most others, is cause for some thought. First, the seeming absence of ZC3HC1 in most insects probably reflects the outcome of separate evolutionary events, perhaps even with different underlying causes. As can be deduced from Figure 3B2, at least one *ZC3HC1* gene loss event appears to have occurred along the evolutionary path leading to Paleoptera like the Odonata, while another one would have happened at some point early in the evolution of the Neoptera, prior to the arising of the Eumetabola, the latter including for example Diptera like *Drosophila melanogaster*. Concerning the gene's loss during the Dipterans' evolution, one could now speculate how this might have come about and whether loop-like insertions within the BLD2 of early Dipterans might have played a role, as outlined in the following.

Such a scenario would be based on several assumptions. One would be that ZC3HC1 would always have been a non-essential protein whose existence reflected the outcome of an evolutionary balancing act. Another one would be that a loop-free ZC3HC1 would represent the initial version of this protein, already functioning as an interconnector of TPR polypeptides at the NB, while the insertion of a loop-forming sequence into a BLD2 would represent an event that happened later. Then, at some point, possibly when beyond a certain length, such a loop would have turned into playing an additional role at the NB, meaning that the protein would have acquired a second functional property. Next, further expansion of the loop's length would have been advantageous concerning the protein's ability to execute the second function. However, beyond a certain length, with the loops in the Orthoptera perhaps defining the length possibly just so still acceptable, such a loop would also gradually come along with increasing problems in correctly assembling the BLD's zinc ion coordination sphere, with the second H-$X_{(3)}$-C pentapeptide widely separated from its corresponding C-$X_{(2)}$-C tetrapeptide. In other words, the loop's evolutionary expansion, to exploit the advantages such a loop could provide, would thus have been expedited at the cost of the protein's functionality regarding its initial task as a structural element of the NB. In again other words, gradually increasing the proportion of those ZC3HC1 polypeptides that no longer function correctly as structural NB components would eventually have neutralized the advantageous effects of an increasingly long loop. Provided then such a neutral point had been reached or overstepped if too many NB binding-incompetent ZC3HC1 polypeptides would represent a handicap, it is imaginable, both with and without adaptive forces at work, that the protein and its large loop would turn into a target for mutations. These could then, e.g., include such that would cause losing the second H-$X_{(3)}$-C pentapeptide, thereby eliminating a functional BLD2 and thus the protein's central function depending on its bimodular construction. Depending on whether the resulting truncated polypeptides would then be more readily tolerable as "molecular garbage" or still represent a problem for the cell, evolution could then dispose of these remnants more gradually or rapidly, blurring and eventually obliterating the traces of the insects' ZC3HC1 gene over time.

Now, we wonder whether this or yet other speculative scenarios might recapitulate the fate that ZC3HC1 experienced very early in Neoptera evolution, near the lineage splitting point, about 260–250 million years ago (e.g., Thomas *et al*, 2013), beyond which the Orthoptera still continued possessing a ZC3HC1 homolog while the other Neoptera had lost their ZC3HC1.

Furthermore, as part of another mind game, we are now also wondering whether such thoughts regarding the loop's advantages and disadvantages can also be adapted for the Viridiplantae that have undergone genome duplication events and then possess two ZC3HC1 homologs. We consider it particularly remarkable that the loop of their one paralog is then of rather "normal length", similar to that in many other organisms, whilst the other paralog's loop is always far longer (see Supplemental List of Sequences), though generally not reaching the extreme lengths in some Phasmatodea. We now wonder whether such plants, having two such ZC3HC1 paralogs at their disposal, take advantage of both opportunities, with the normal-sized loop paralog being the one readily fulfilling its task as a structural element at the NB and the second one additionally or solely exploiting the loops' advantages, with less balancing required between the two functions.

**(B)** Endogenous ZC3HC1 polypeptides have been recurrently identified as being specifically and varyingly phosphorylated at specific sites not only during normal cell cycle progression (Bassermann *et al*, 2005a; Dephoure *et al*, 2008; Blethrow *et al*, 2008; Chi *et al*, 2008; Illert *et al*, 2012; Zhou *et al*, 2013; Sharma *et al*, 2014) but also upon a range of different stimuli and the activation of kinases that operate in different cell signaling pathways (Christensen *et al*, 2010; Moritz *et al*, 2010; Weintz *et al*, 2010; Yu *et al*, 2011; Robitaille *et al*, 2013; Sos *et al*, 2014; Sharma *et al*, 2014; Hu *et al*, 2015). While ZC3HC1 phosphorylation at the onset of mitosis likely represents one of the steps eventually leading to complete ZC3HC1 solubilization along with NB disassembly, the multisite-phosphorylation of ZC3HC1 during interphase in response to extracellular stimuli suggests that ZC3HC1 in general, and the loop, in particular, could play some role in cellular stress response and along specific signal transduction pathways. Accordingly, and in addition to TPR, which is a target for stress-activatable phosphatases (Yadav *et al*, 2017; Wigington *et al*, 2020), we deem it possible that ZC3HC1, which also possesses prototypic binding sites for stress-activated phosphatases, like, e.g., for calcineurin at aa 390–393, and thus also within the loop region, might eventually turn out being a target too for such phosphatases, and other enzymes, in interphase.

The upper scheme, corresponding to Figure 6D, includes all serine and threonine residues of *Hs*ZC3HC1 so far reported (July 2022) to have been found phosphorylated (https://www.phosphosite.org/proteinAction.action?id=3471&showAllSites=true). The lower scheme depicts those phosphosites for which at least five corresponding datasets have been deposited at https://www.phosphosite.org.

| Identified structures: | | | Query structures: | | | | | | | | | | | | | |
|---|---|---|---|---|---|---|---|---|---|---|---|---|---|---|
| | | | *Hs*ZC3HC1 | | *Hs*ZC3HC1_minimal | | *Dd*ZC3HC1 | | *Dd*ZC3HC1_minimal | | *Sc*Pml39p | | *Sc*Pml39p_minimal | | *Dm*Diap2 | |
| Target species | Identifier (afdb-proteome) | Protein name | Score | TM-score | Score | TM-score | Score | TM-score | Score | TM-score | Score | TM-score | Score | TM-score | Score | TM-score |
| ***Homo sapiens*** | **AF-Q86WB0-F1-model_v2** | **ZC3HC1** | **3458** | **5,56E-78** | **1635** | **4,25E-35** | **757** | **8,90E-16** | **649** | **6,89E-14** | **383** | **4,03E-07** | **348** | **1,88E-06** | 197 | 9,67E-03 |
| *Homo sapiens* | AF-Q13075-F1-model_v2 | BIRC1 | - | - | - | - | - | - | - | - | 188 | 1,57E-02 | 184 | 1,62E-02 | 844 | 5,66E-18 |
| *Homo sapiens* | AF-Q13490-F1-model_v2 | BIRC2 | - | - | - | - | - | - | - | - | 172 | 3,75E-02 | 156 | 7,59E-02 | 1398 | 5,12E-31 |
| *Homo sapiens* | AF-Q13489-F1-model_v2 | BIRC3 | - | - | - | - | - | - | 151 | 8,92E-02 | 173 | 3,55E-02 | 158 | 6,80E-02 | 1454 | 2,46E-32 |
| *Homo sapiens* | AF-P98170-F1-model_v2 | BIRC4 | - | - | 200 | 1,26E-02 | - | - | 204 | 4,58E-03 | 248 | 6,09E-04 | 200 | 6,68E-03 | **1550** | **1,35E-34** |
| *Homo sapiens* | AF-O15392-F1-model_v2 | BIRC5 | - | - | - | - | - | - | - | - | - | - | - | - | 260 | 3,18E-04 |
| *Homo sapiens* | AF-Q9NR09-F2-model_v2 | BIRC6 | - | - | - | - | - | - | - | - | - | - | - | - | 221 | 2,63E-03 |
| *Homo sapiens* | AF-Q9NR09-F1-model_v2 | BIRC6 | - | - | - | - | - | - | - | - | - | - | - | - | 207 | 5,62E-03 |
| *Homo sapiens* | AF-Q96CA5-F1-model_v2 | BIRC7 | - | - | - | - | - | - | - | - | - | - | - | - | 696 | 1,73E-14 |
| *Homo sapiens* | AF-Q96P09-F1-model_v2 | BIRC8 | - | - | - | - | - | - | - | - | - | - | - | - | 797 | 7,23E-17 |
| *Homo sapiens* | AF-Q99675-F1-model_v2 | CGRF1 | - | - | - | - | - | - | - | - | - | - | - | - | 189 | 1,49E-02 |
| *Homo sapiens* | AF-Q6UWE0-F1-model_v2 | LRSAM1 | - | - | - | - | - | - | - | - | - | - | - | - | 215 | 3,64E-03 |
| *Homo sapiens* | AF-Q6ZN04-F1-model_v2 | MEX3B | - | - | - | - | - | - | - | - | - | - | - | - | 165 | 5,48E-02 |
| *Homo sapiens* | AF-Q86YT6-F1-model_v2 | MIB1 | - | - | - | - | - | - | - | - | - | - | - | - | 221 | 2,63E-03 |
| *Homo sapiens* | AF-Q969V5-F1-model_v2 | MUL1 | - | - | - | - | - | - | - | - | - | - | - | - | 210 | 4,78E-03 |
| *Homo sapiens* | AF-Q8WZ73-F1-model_v2 | RFFL | - | - | - | - | - | - | - | - | - | - | - | - | 224 | 2,24E-03 |
| *Homo sapiens* | AF-Q5VTB9-F1-model_v2 | RNF220 | - | - | - | - | - | - | - | - | - | - | - | - | 161 | 6,81E-02 |
| *Homo sapiens* | AF-Q9BY78-F1-model_v2 | RNF26 | - | - | - | - | - | - | - | - | - | - | - | - | 269 | 1,95E-04 |
| ***Dictyostelium discoideum*** | **AF-Q54PS8-F1-model_v2** | **ZC3HC1** | 835 | 2,17E-17 | 607 | 7,75E-12 | 4037 | 2,35E-92 | 1385 | 8,68E-32 | 284 | 8,64E-05 | 250 | 4,22E-04 | - | - |
| *Dictyostelium discoideum* | AF-Q55EJ5-F1-model_v2 | MYLIP | - | - | - | - | - | - | - | - | - | - | - | - | 207 | 5,62E-03 |
| ***Saccharomyces cerevisiae*** | **AF-Q03760-F1-model_v2** | **Pml39p** | - | - | **405** | **2,88E-07** | - | - | **388** | **1,54E-07** | **2786** | **1,07E-63** | **1927** | **2,48E-44** | **220** | **2,78E-03** |
| *Saccharomyces cerevisiae* | AF-P47134-F1-model_v2 | Bir1p | - | - | - | - | - | - | 155 | 7,13E-02 | 191 | 1,34E-02 | 178 | 2,25E-02 | 269 | 1,95E-04 |
| *Saccharomyces cerevisiae* | AF-P54074-F1-model_v2 | Asi1p | - | - | - | - | - | - | - | - | - | - | - | - | 200 | 8,22E-03 |
| ***Drosophila melanogaster*** | **AF-Q24307-F1-model_v2** | **Diap2** | - | - | **210** | **7,45E-03** | - | - | **211** | **3,10E-03** | **225** | **2,12E-03** | **223** | **1,88E-03** | **4007** | **1,92E-92** |
| *Drosophila melanogaster* | AF-Q24306-F1-model_v2 | Diap1 | - | - | - | - | - | - | - | - | - | - | - | - | 1030 | 2,36E-22 |
| *Drosophila melanogaster* | AF-Q9VEM2-F1-model_v2 | Deterin | - | - | - | - | - | - | - | - | - | - | - | - | 265 | 2,42E-04 |
| *Drosophila melanogaster* | AF-Q9VCV3-F1-model_v2 | RNF220 | - | - | - | - | - | - | - | - | - | - | - | - | 208 | 5,32E-03 |
| *Drosophila melanogaster* | AF-Q9VUX2-F1-model_v2 | Mib1 | - | - | - | - | - | - | - | - | - | - | - | - | 187 | 1,66E-02 |
| *Drosophila melanogaster* | AF-Q9VZJ9-F1-model_v2 | Mul1 | - | - | - | - | - | - | - | - | - | - | - | - | 208 | 5,32E-03 |
| *Drosophila melanogaster* | AF-P29503-F1-model_v2 | Neur | - | - | - | - | - | - | - | - | - | - | - | - | 179 | 2,57E-02 |
| *Drosophila melanogaster* | AF-P20193-F1-model_v2 | Su(var)3-7 | - | - | - | - | - | - | - | - | - | - | - | - | 199 | 8,67E-03 |
| *Drosophila melanogaster* | AF-Q9VIK5-F1-model_v2 | CG2617 | - | - | - | - | - | - | - | - | - | - | - | - | 186 | 1,76E-02 |

**Supplemental Figure S14. Searching AlphaFold2's protein structure datasets via Foldseek, using ZC3HC1 and BIR protein structures as queries.**

In order to search for proteins with ZC3HC1-reminiscent structures in those insect species, like *Drosophila melanogaster*, that appear to lack a homolog of ZC3HC1 recognizable as such at the protein sequence level, we used the Foldseek tool (https://search.foldseek.com/search; van Kempen *et al*, 2022) for conducting some first searches among the PDB files of the AlphaFold/ Proteome v2 database (here abbreviated as afdb-proteome). As query structures, we used those of the human, amoebic and budding yeast ZC3HC1 homologs (Uniprot identifiers Q86WB0, Q54PS8, and Q03760, respectively), which we also used in their truncated versions (here referred to as the minimal structures), corresponding to those presented in Supplemental Figure S11B, lacking the loops and most of the other parts not regarded as belonging to the NuBaID. In addition, we used the *Drosophila melanogaster* BIR protein Diap2 (Q24307) as a query structure for comparison. Searches via Foldseek were conducted in the 3Di/AA mode, using the taxonomic filter for the respective other species, with searches among the *D. melanogaster* structures having been complemented by searching the human, amoebic and budding yeast structure datasets for comparison. All identified structures and their scores, as obtained with the default settings of Foldseek and retrieved from Foldseek's web server, are presented. Note that these searches, for now, did not reveal a *Drosophila* protein structure that we would regard as a likely ZC3HC1 equivalent, with the only non-ZC3HC1 structures identified with the ZC3HC1 query structures being known BIR proteins. By contrast, when searching the database-deposited human, amoebic, and budding yeast structures with either the *Hs*ZC3HC1, *Dd*ZC3HC1, or

*Sc*Pml39p structure as the only query, the two other species' ZC3HC1 structures were in each case identifiable as the best matches when using the homologs' loop-free NuBaID structures.

### Supplemental Discussion

**Supplemental Discussion 1. A comparison of the NuBaID- and BIR-type of zinc fingers.** The BIR domains have been described as comprising approximately 70 amino acids, including three invariant cysteine residues plus one histidine for tetrahedrally coordinating a zinc ion (Birnbaum *et al*, 1994). They are a characteristic feature of the IAPs, which also exist in insects (e.g., Orme & Meier, 2009; Berthelet & Dubrez, 2013), while absent in plants *(e.g., Higashi et al*, 2005; Cao *et al*, 2008; https://pfam.xfam.org/family/BIR). Each of the IAPs possesses at least one and often two or three of these zinc finger modules, and in some rare cases, perhaps more (e.g., Mace *et al*, 2010a; Silke & Vucic, 2014; https://pfam.xfam.org/family/BIR#tabview=tab1).

It appears evident that the BIR domains share a common ancestor with the BLDs of the NuBaID, with such kinship already proposed in the past (Higashi *et al*, 2005; Kokoszynska *et al*, 2008) and with such notion now substantiated by further findings. While the BIR domain's most common H-$X_{(6)}$-C spacing of its zinc-coordinating histidine and third cysteine distinguishes it from the H-$X_{(3)}$-C arrangement of the NuBaID's two zinc finger modules, it had been noted early on that the BIR domain shares some additional, seemingly conserved residues with either the one or the other or both of the two potential zinc finger modules of several ZC3HC1 homologs. Based on such similarity, and since BIR domain-containing IAPs are absent in plants, the two ILP proteins in *Arabidopsis* had even been considered to take on tasks equivalent to those of the IAPs in other species (Higashi *et al*, 2005). Furthermore, such local similarities between the BIR domains' and the ZC3HC1 homologs' sequences thus had led to naming each of the latter's two zinc finger modules a BLD (Higashi *et al*, 2005; Kokoszynska *et al*, 2008).

As we now know, some of the NuBaID residues common to both the BLDs and BIR domains are essential for *Hs*ZC3HC1 and *Sc*Pml39p to adopt a conformation enabling them to bind to the NB. However, as has been reasoned in this study's main text, these BLD residues, among which are also the invariant ones of the NuBaID's minimal sequence signature, are likely not directly interacting with TPR. Instead, in line with a former homology model of the BLD1 of *Hs*ZC3HC1 (Higashi *et al*, 2005) and now also according to AlphaFold2's predictions, these residues appear to participate in intramolecular interactions and to play, like the BIR domains' corresponding ones (Supplemental Figures S9C and S10C2), both direct and indirect roles in the establishment of shielded zinc ion coordination spheres. Such a similarity between the central parts of the BLDs and the BIR domain's core construction appeared particularly evident after having compared the BLD and BIR domains' structure predictions provided by AlphaFold2 (Supplemental Figure S10C and S10E), and these with the BIR domains' structures determined by X-ray crystallography (e.g., Cossu *et al*, 2019; https://www.rcsb.org; Supplemental Figure S10A1). Therefore, even though the current version of the NuBaID signature allows for distinguishing the ZC3HC1 homologs of numerous, if not most, species from their BIR domain-possessing proteins, one must keep in mind that such a distinction via this signature only relates to a few residue preferences and their spacing within the core structures of these two closely related domains.

With the current versions of the here presented NuBaID signature thus not describing a TPR-binding interface, we expect that reports to come will unveil those sequence elements that define the BLDs' specificity for TPR and distinguish the BLDs' binding interfaces from those of the BIR domains. Some of these target specificity-defining sequence features are possibly already conjecturable from the hidden Markoff models (HMM) of the Pfam database's current versions of its zf-C3HC and Rsm1 motifs, and we can imagine them becoming more evident once the sequences of the loop-free BLDs in their newly defined boundaries are taken into account for updating such motifs.

The NPC-associated TPR protein being a specific binding partner of the BLDs and rather not a target of any BIR domain, was actually in line with us not having found any of the eight vertebrate IAP/BIR proteins (e.g., Deveraux & Reed, 1999; Dubrez-Daloz *et al*, 2008) as naturally interacting with TPR at the NB, neither in *Xenopus* oocytes nor in human tumor cell lines in standard growth conditions (our unpublished data). Furthermore, we also did not find ZC3HC1 stably binding to any other protein than TPR in normally growing human cells in interphase (our unpublished data; but see also Gunkel *et al*, 2021), and neither did we find ZC3HC1 to act as an inhibitor of apoptosis, at least not in its typical physiological concentrations within the cell (Gunkel *et al*, 2021).

However, we currently cannot exclude the possibility that ZC3HC1 might transiently interact with one or another of those proteins interacting with the IAPs. Instead, we even consider it tempting to speculate that the IAPs and ZC3HC1 might use similar mechanisms to regulate the interplay with their respective binding partners. In this context, we also have those proteins in mind that act as IAP antagonists and that, upon stress stimulation, bind to the IBM groove of the type II BIR domains, like in the case of the natural BIRC4 antagonist Smac/DIABLO and thereby displace an IAP's actual binding partner (e.g., Verhagen *et al*, 2001; Gyrd-Hansen & Meier, 2010; Damgaard & Gyrd-Hansen, 2011; Cossu *et al*, 2019). Here then, in the current study, we noted that the BLD1 possesses a similarly positioned conspicuous groove

(Supplemental Figure S10D). Moreover, even though such a groove at the BLD1-corresponding position is not to be seen in the BLD2, the residues at the BLD2 position directly corresponding to the BIR domains' groove turned out conspicuously similar to those defining the IBM groove. We now wonder whether the BLD1 groove or even certain residues of the corresponding part of BLD2 might have a function equivalent to that of the IBM groove, allowing for regulating the interaction between ZC3HC1 and TPR at the NB as part of a cellular stress response. This notion was also inspired by a few Smac/DIABLO polypeptides that we had found among those proteins co-precipitated with soluble ZC3HC1 when the latter had been immunoprecipitated from cell extracts (Gunkel *et al*, 2021, and our unpublished data). However, whether such findings reflect an interaction of physiological relevance still needs to be determined. We already propose, though, to test the IBM antagonist molecules used for cancer research (e.g., Cossu *et al*, 2019) to investigate whether these might interact with ZC3HC1 and perhaps result in destabilization or even displacement of ZC3HC1 and TPR polypeptides from the NB.

However, apart from the evident similarities and those possibly still to emerge, the BLDs and the BIRs represent proteins that nonetheless conspicuously differ in several respects. Such differences not only relate to the BLDs being the characterizing feature of a generally unique, one-of-a-kind protein per species with a non-duplicated genome, in contrast to the BIRs, which can be part of several different proteins. Beyond that, the BIR domains are known to interact with different primary binding partners. In mammals, for example, such binding partners include, among others, tumor necrosis factor (TNF) receptor-associated factors (TRAFs), which interact with type I BIR domains, and effector caspases, which bind to the BIR domains of type II (e.g., Rothe *et al*, 1995; Roy *et al*, 1997; Takahashi *et al*, 1998; Chai *et al*, 2001; Huang *et al*, 2001; Riedl *et al*, 2001; Samuel *et al*, 2006; Gyrd-Hansen & Meier, 2010; Mace *et al*, 2010a, 2010b; Silke & Vucic, 2014; Lalaoui & Vaux, 2018). By contrast, current evidence does not suggest that the BLDs of the vertebrate ZC3HC1 homologs stably bind directly to a regular binding partner other than TPR, also since other proteins formerly proposed as regular ZC3HC1 binding partners (e.g., Bassermann *et al*, 2005a, 2007; Kreutmair *et al*, 2020) were refuted (Gunkel *et al*, 2021) or assessed as unlikely (Supplemental Figure S12). In other words, while the BIR domains act as binding modules for several different proteins within a given species, the NuBaID with its two BLDs currently appears monogamous for only one stably to be bound target protein, namely TPR.

Furthermore, some of the IAP's BIR domains have been shown capable of forming homodimers, with examples, among others, including the BIR1 and the BIR3 domain of protein BIRC4 (Lu *et al*, 2007; Lin *et al*, 2007; Mastrangelo *et al*, 2008). However, whether such capability would further distinguish the BIRs from the BLDs remains uncertain. So far, neither our experimental data nor predictions by AlphaFold2 have allowed us to answer for sure whether ZC3HC1 can dimerize or not.

Clearly, however, another property again distinguishes the IAPs and their BIR domains from ZC3HC1 with its two BLDs strikingly: An individual BIR domain, either as a naturally occurring one as part of a single-BIR domain IAP or when part of an IAP with several of them, represents an autonomous binding unit. Even when separated from their neighboring BIR domains, the individual ones of an IAP, like BIRC4, can still interact with their respective target proteins (e.g., Mace *et al*, 2010a). By contrast, even though each of the two BLDs of ZC3HC1 on their own could well be capable of zinc ion coordination, neither of them, when separated from each other, is capable of a sufficiently robust standalone interaction with TPR *in vivo*, which in the current study even held for the Y2H interactions in yeast cells. Our findings thus confirmed the conclusions of the study in which the BLDs had been described first (Higashi *et al*, 2005) and in which the BLD repeat, i.e., the existence of two BLDs, had been predicted to be essential for the ILPs', i.e., the ZC3HC1 homologs' function.

In other words, while the affinity between one BIR domain and its regular target protein suffices for a lasting interaction, the bipartite NuBaID only allows for a lasting interaction with its corresponding TPR homolog, at least in humans and yeast, when both of its two BLDs are intact and connected. Again, in other words, while an IAP with several BIR domains does not necessarily require cooperativity between its different BIR domains for target protein binding, the two BLDs need to act in concert, either by both contributing to one complex TPR binding interface or by each binding separately but cooperatively to the NPC-anchored homodimers of TPR. Only the avidity of such a bivalent interaction might provide the required strength of interaction that allows for a lasting engagement of ZC3HC1 with TPR *in vivo*.

Furthermore, yet another criterion distinguishes the ZC3HC1 homologs from numerous members of the eukaryotic realm's IAPs. Many of the latter possess not only one or several BIR domains but also one or several other types of domains, among which are some that additionally enable homodimerization, allow for interaction with yet other proteins, or play a role in ubiquitination (e.g., Oberoi-Khanuja *et al*, 2013; Silke & Vaux, 2015; Cossu *et al*, 2019). The presence of such other domains applies, for example, to seven of the eight IAPs in humans, with only the small, single BIR domain-possessing BIRC5 lacking such an additional one. Five human IAPs even possess yet another type of zinc ion coordination system, namely the RING finger (e.g., Oberoi-Khanuja *et al*, 2013). By striking contrast, hardly any of the eukaryotic realm's ZC3HC1 homologs appear to possess, next to their BLDs, an additional protein domain of those currently known. In fact, upon careful inspection of the relatively few sequences corresponding to those illustrations that show, in the Pfam database, a zf-C3HC or an Rsm1 motif as part of a protein possessing different types of domains (https://pfam.xfam.org/family/zf-C3HC, https://pfam.xfam.org/family/Rsm1), we found most of these assemblages explainable differently. The underlying sequences, the majority of which were genomic, harbored computational errors in sequence interpretation, including misassembled sequences, wrongly predicted or assigned exons, or other types of errors (our unpublished data). Therefore, we now dare to conclude that, at least in higher eukaryotes, a genuine ZC3HC1 homolog can also be described by its lack of other known protein domains, thereby further distinguishing it from many of the BIR-possessing IAPs.

Finally, the other prominent feature that distinguishes the IAPs from many ZC3HC1 homologs is the latters' capability to harbor large sequence inserts at different sites within their bounds, including loop-like insertions within their second BLD that can be extraordinarily long, with only a few examples presented in the current study (Supplemental Figure S13A). Even though such loops are also missing in some ZC3HC1 homologs, like in *Sc*Pml39p, they appear absent in the BIR domains far more commonly, if not categorically (see also, https://pfam.xfam.org/family/BIR#tabview=tab1).

**Supplemental Discussion 2. Further thoughts regarding the presence of ZC3HC1 in many organisms and its absence in others.**

While in some organisms, like in most insect orders, all signs of a former ZC3HC1 and its NuBaID signature appear to have disappeared, suggesting the protein's ultimate loss, other organisms possess ZC3HC1 homologs with aa substitutions or deletions that would render the human homolog incapable of binding to the NB and TPR. Notably, these potential mutant versions of ZC3HC1 appear mostly impaired with regard to their BLD2, with parts of the latter sometimes still recognizable in some species and absent in others *in toto*, while the appertaining BLD1 appears unaffected. Moreover, having scrutinized those database-deposited sequences that suggested the existence of truncated ZC3HC1 versions comprising only BLD2 while lacking an intact BLD1, which also included 48 cases listed so far in the Pfam database for the Rsm1 motif (https://pfam.xfam.org/family/PF08600#tabview=tab1), we found these BLD2-only versions of ZC3HC1 to be incomplete merely for procedural reasons, as we could identify an associated BLD1 for each of them (our unpublished data). Based on these findings, we initially concluded that it was mostly the BLD2 that had been a target for mutations in those species in which they appeared to have occurred, like in several unicellular organisms and in some marine invertebrates.

For example, in all tunicates for which sequence information was available by the end of this study, like for the genus *Ciona* of the class Ascidiacea, some of the second BLD's zinc ion-coordinating residues were found exchanged for such not capable of zinc ion coordination. In other tunicates again, like in the genus *Oikopleura* of the class Appendicularia, the second BLD was found lost in its entirety. We further noted such putative signs of ZC3HC1 homologs being in a state of disintegration, by either a complete loss of their BLD2 or by accumulating mutations that one would regard as abolishing their former zinc ion coordination ability, to exist also in very different organisms. Among them are, as another example, the xerophilic fungi of the genus *Wallemia*, in which we additionally confirmed the second BLD's absence by cDNA cloning and sequencing of the "residual" ZC3HC1 homolog of *Wallemia mellicola*, thereby supporting an already database-deposited sequence (see Supplemental List of Sequences).

We also need to note, though, that we do not exclude the existence of naturally occurring single aa substitution mutations also within the BLD1 of some organisms, as exemplified by the ZC3HC1 homolog from *Zygosaccharomyces mrakii* (XP_037145256.1; see Supplemental List of Sequences), which despite its sequence mutation, has a Pfam zf-C3HC motif assigned to it nonetheless. Assuming this sequence can be confirmed to be correct, one could further imagine this homolog no longer capable of binding to an NB, with this assumption, in turn, based on presuming that this species' aa substitution, when introduced into *Hs*ZC3HC1, would abolish the NB-binding capability of the latter, with this particular substitution though still having to be tested. Moreover, apart from that, it certainly will require further efforts to systematically screen and verify the sequences of ZC3HC1 homologs for those that harbor aa substitutions that might abolish a homolog's ability to bind to the NB. Currently, we can only tell that the frequency of potential NB-binding-competence-abolishing mutations as yet noted within the different species' BLD domains appears conspicuously higher in the BLD2 when compared to the then relatively few ones so far detected in the BLD1.

Such findings and conclusions, though, then raise further questions. For example, if the functions of *Hs*ZC3HC1 and *Sc*Pml39p, and the sequence prerequisites for a NuBaID with two BLDs both required for binding TPR and Mlp1p, would also hold for other species' ZC3HC1 homologs, and if one of the latter would then be destined for decay by evolution for whatever reasons, one might not necessarily expect the detectable mutations in some clades primarily occurring within or even confined to the BLD2 alone. One could, therefore, ask whether the remaining, seemingly intact BLD1 might still be good for something in these organisms. If this were the case, one could imagine the single-BLD version having compensated for the second BLD's loss and evolved the means to bind to TPR on its own, thus still allowing for fulfilling distinct tasks at the NB. Of course, one could also imagine such a single-BLD ZC3HC1 variant to be located somewhere else within the cell, where it might fulfill an NB-unrelated function instead of spending merely a non-functional existence as molecular garbage prior to its complete elimination over time. Studying organisms in which such seemingly truncated ZC3HC1 versions exist, and determining their subcellular locations, should now rather straightforwardly allow distinguishing between some of these possibilities.

However, if the seemingly mutated ZC3HC1 versions reflect a state of ongoing evolutionary decay, or even if they represent polypeptides with some residual functional activity or novel tasks, this brings us back to the question of why no selective pressure has preserved the protein's original version and function. One could further ask whether the ZC3HC1 homolog's original might merely have been lost incidentally or whether evolutionary forces might actually have been selecting against it in certain species. Such questions, of course, also apply to those organisms in which ZC3HC1 homologs are no longer detectable at all.

In one scenario, the protein would simply no longer have been of use for a particular species during its replicative lifespan or reproductive phase, with no remaining selective pressure maintaining its existence as a functionally intact protein. Even if minor deficits might have come along with its absence, one could imagine these to have been compensated for by co-evolved counter-steering adaptations of other proteins**.** In other words, during the adaptation to its current environment, such a species' ZC3HC1 homolog, formerly favorable in another environment, would have turned into a gene whose contribution to the species' fitness would eventually have been neutral, with this then also describable as conditional neutrality (e.g., Bargiello & Grossfield, 1979; Anderson *et al*, 2013). Related to this notion, we wonder, with such thoughts also specified elsewhere (Supplemental Figure S13), whether the existence of the extremely long BLD2 insertions in the ZC3HC1 homologs of the Orthoptera might hint at how such a fate of becoming dispensable over time might have come to pass and eventually have led to the ZC3HC1 homologs' absence in the other insect orders of the Neoptera.

In a different scenario, the presence of a species' ZC3HC1 homolog, while again advantageous in some situations, would be disadvantageous in others, resulting in fitness trade-offs. Evolutionary forces would then have expedited the gene's elimination once the disadvantages started overwhelming, with this reflecting the concept of antagonistic pleiotropy (e.g., Roff & Fairbairn, 2007; Anderson *et al*, 2011; see also Carter & Nguyen, 2011). However, we do not regard either one or the other of these different scenarios as solely possible, and we can conceive of ZC3HC1 homologs in different organisms that have been lost for different reasons. In fact, we can imagine that a ZC3HC1 homolog in the one species had indeed become dispensable, its possession thus no longer providing a fitness advantage, resulting in the accumulation of incidental mutations over time. In other species, though, we can imagine a trade-off between the ZC3HC1 homologs' pros and cons that has tilted to the disadvantageous side, causing evolutionary forces to exert selective pressure for the protein's elimination. This notion is outlined in further detail in this study's main discussion.

### Supplemental Tables

#### Supplemental Table S1: Expression vectors

| Description (promoter > expressed protein) | Backbone (resistance) | Source |
|---|---|---|
| ADH1>GAL4-BD (empty vector) | pGBT9 (Amp) | Clontech, Mountain View CA, USA |
| ADH1>GAL4-BD-*Dd*ZC3HC1(1–271_346–426_486–635) | pGBT9 (Amp) | This study |
| ADH1>GAL4-BD-*Hs*ZC3HC1(1–502) | pGBT9 (Amp) | This study |
| ADH1>GAL4-BD-*Hs*ZC3HC1(1–502 \| C102S) | pGBT9 (Amp) | This study |
| ADH1>GAL4-BD-*Hs*ZC3HC1(1–502 \| W107A) | pGBT9 (Amp) | This study |
| ADH1>GAL4-BD-*Hs*ZC3HC1(1–502 \| C112S) | pGBT9 (Amp) | This study |
| ADH1>GAL4-BD-*Hs*ZC3HC1(1–502 \| C117S) | pGBT9 (Amp) | This study |
| ADH1>GAL4-BD-*Hs*ZC3HC1(1–502 \| C120S) | pGBT9 (Amp) | This study |
| ADH1>GAL4-BD-*Hs*ZC3HC1(1–502 \| C125S) | pGBT9 (Amp) | This study |
| ADH1>GAL4-BD-*Hs*ZC3HC1(1–502 \| H152A) | pGBT9 (Amp) | This study |
| ADH1>GAL4-BD-*Hs*ZC3HC1(1–502 \| C156S) | pGBT9 (Amp) | This study |
| ADH1>GAL4-BD-*Hs*ZC3HC1(1–502 \| W158A) | pGBT9 (Amp) | This study |
| ADH1>GAL4-BD-*Hs*ZC3HC1(1–502 \| C249S) | pGBT9 (Amp) | This study |
| ADH1>GAL4-BD-*Hs*ZC3HC1(1–502 \| W256A) | pGBT9 (Amp) | This study |
| ADH1>GAL4-BD-*Hs*ZC3HC1(1–502 \| C272S) | pGBT9 (Amp) | This study |
| ADH1>GAL4-BD-*Hs*ZC3HC1(1–502 \| C275S) | pGBT9 (Amp) | This study |
| ADH1>GAL4-BD-*Hs*ZC3HC1(1–502 \| H363R) | pGBT9 (Amp) | This study |
| ADH1>GAL4-BD-*Hs*ZC3HC1(1–502 \| H425A) | pGBT9 (Amp) | This study |
| ADH1>GAL4-BD-*Hs*ZC3HC1(1–502 \| C429S) | pGBT9 (Amp) | This study |
| ADH1>GAL4-BD-*Hs*ZC3HC1(1–502 \| W431A) | pGBT9 (Amp) | This study |
| ADH1>GAL4-AD (empty vector) | pGAD424 (Amp) | Clontech, Mountain View CA, USA |
| ADH1>GAL4-AD-*Sc*Mlp1p(1–143) | pGAD424 (Amp) | This study |
| ADH1>GAL4-AD-*Sc*Mlp1p(1–190) | pGAD424 (Amp) | This study |
| ADH1>GAL4-AD-*Sc*Mlp1p(1–297) | pGAD424 (Amp) | This study |
| ADH1>GAL4-AD-*Sc*Mlp1p(287–499) | pGAD424 (Amp) | This study |
| ADH1>GAL4-AD-*Sc*Mlp1p(287–584) | pGAD424 (Amp) | This study |
| ADH1>GAL4-AD-*Sc*Mlp2p(1–120) | pGAD424 (Amp) | This study |
| ADH1>GAL4-AD-*Sc*Mlp2p(1–210) | pGAD424 (Amp) | This study |
| ADH1>GAL4-AD-*Sc*Mlp2p(199–626) | pGAD424 (Amp) | This study |
| ADH1>GAL4-BD-*Sc*Pml39p(1–334) | pGBT9 (Amp) | This study |
| ADH1>GAL4-BD-*Sc*Pml39p(1–334 \| W119A) | pGBT9 (Amp) | This study |
| ADH1>GAL4-BD-*Sc*Pml39p(1–334 \| C134S) | pGBT9 (Amp) | This study |
| ADH1>GAL4-BD-*Sc*Pml39p(1–334 \| C134–137SGGS) | pGBT9 (Amp) | This study |
| ADH1>GAL4-BD-*Sc*Pml39p(1–334 \| C176S) | pGBT9 (Amp) | This study |
| ADH1>GAL4-BD-*Sc*Pml39p(1–334 \| Y257A) | pGBT9 (Amp) | This study |
| ADH1>GAL4-BD-*Sc*Pml39p(1–334 \| Y257W) | pGBT9 (Amp) | This study |
| ADH1>GAL4-BD-*Sc*Pml39p(1–334 \| C271S) | pGBT9 (Amp) | This study |
| ADH1>GAL4-BD-*Sc*Pml39p(1–334 \| C292S) | pGBT9 (Amp) | This study |
| ADH1>GAL4-AD-*Hs*TPR(1–60) | pGAD424 (Amp) | This study |
| ADH1>GAL4-AD-*Hs*TPR(1–74) | pGAD424 (Amp) | This study |
| ADH1>GAL4-AD-*Hs*TPR(1–88) | pGAD424 (Amp) | This study |
| ADH1>GAL4-AD-*Hs*TPR(1–102) | pGAD424 (Amp) | This study |
| ADH1>GAL4-AD-*Hs*TPR(1–111) | pGAD424 (Amp) | This study |
| ADH1>GAL4-AD-*Hs*TPR(1–175) | pGAD424 (Amp) | This study |
| ADH1>GAL4-AD-*Hs*TPR(11–109) | pGAD424 (Amp) | This study |
| ADH1>GAL4-AD-*Hs*TPR(20–111) | pGAD424 (Amp) | This study |
| ADH1>GAL4-AD-*Hs*TPR(29–175) | pGAD424 (Amp) | This study |
| ADH1>GAL4-AD-*Hs*TPR(43–175) | pGAD424 (Amp) | This study |
| ADH1>GAL4-AD-*Hs*TPR(54–175) | pGAD424 (Amp) | This study |
| ADH1>GAL4-AD-*Hs*TPR(110–342) | pGAD424 (Amp) | This study |
| ADH1>GAL4-AD-*Hs*TPR(110–377) | pGAD424 (Amp) | This study |
| ADH1>GAL4-AD-*Hs*TPR(172–651) | pGAD424 (Amp) | This study |
| ADH1>GAL4-AD-*Hs*TPR(233–342) | pGAD424 (Amp) | This study |
| ADH1>GAL4-AD-*Hs*TPR(233–499) | pGAD424 (Amp) | This study |
| ADH1>GAL4-AD-*Hs*TPR(275–450) | pGAD424 (Amp) | This study |
| ADH1>GAL4-AD-*Hs*TPR(275–481) | pGAD424 (Amp) | This study |
| ADH1>GAL4-AD-*Hs*TPR(275–539) | pGAD424 (Amp) | This study |
| ADH1>GAL4-AD-*Hs*TPR(347–499) | pGAD424 (Amp) | This study |
| ADH1>GAL4-AD-*Hs*TPR(347–543) | pGAD424 (Amp) | This study |
| ADH1>GAL4-AD-*Hs*TPR(361–539) | pGAD424 (Amp) | This study |
| ADH1>GAL4-AD-*Hs*TPR(386–481) | pGAD424 (Amp) | This study |
| ADH1>GAL4-AD-*Hs*TPR(386–539) | pGAD424 (Amp) | This study |
| ADH1>GAL4-AD-*Hs*TPR(411–539) | pGAD424 (Amp) | This study |

<table>
<tr><td>ADH1>GAL4-AD-*Hs*TPR(432–539)</td><td>pGAD424 (Amp)</td><td>This study</td></tr>
<tr><td>ADH1>GAL4-AD-*Hs*TPR(450–543)</td><td>pGAD424 (Amp)</td><td>This study</td></tr>
<tr><td>ADH1>GAL4-AD-*Hs*TPR(608–940)</td><td>pGAD424 (Amp)</td><td>This study</td></tr>
<tr><td>ADH1>GAL4-AD-*Hs*TPR(926–1178)</td><td>pGAD424 (Amp)</td><td>This study</td></tr>
<tr><td>ADH1>GAL4-AD-*Hs*TPR(1129–1632)</td><td>pGAD424 (Amp)</td><td>This study</td></tr>
<tr><td>ADH1>GAL4-AD-*Hs*TPR(1177–1632)</td><td>pGAD424 (Amp)</td><td>This study</td></tr>
<tr><td>ADH1>GAL4-AD-*Hs*TPR(1618–1917)</td><td>pGAD424 (Amp)</td><td>This study</td></tr>
<tr><td>ADH1>GAL4-AD-*Hs*TPR(1894–2138)</td><td>pGAD424 (Amp)</td><td>This study</td></tr>
<tr><td>ADH1>GAL4-AD-*Hs*TPR(2110–2363)</td><td>pGAD424 (Amp)</td><td>This study</td></tr>
<tr><td>CMV>EYFP-*Hs*KPNB1(1–876)</td><td>pEYFP-C1 (Kan)</td><td>This study</td></tr>
<tr><td>CMV>EYFP-*Hs*SKP1(1–163)</td><td>pEYFP-C1 (Kan)</td><td>This study</td></tr>
<tr><td>CMV>EYFP-*Hs*ZC3HC1(1–502)</td><td>pEYFP-C1 (Kan)</td><td>This study</td></tr>
<tr><td>CMV>EYFP-*Hs*ZC3HC1(1–490)</td><td>pEYFP-C1 (Kan)</td><td>This study</td></tr>
<tr><td>CMV>EYFP-*Hs*ZC3HC1(1–477)</td><td>pEYFP-C1 (Kan)</td><td>This study</td></tr>
<tr><td>CMV>EYFP-*Hs*ZC3HC1(1–467)</td><td>pEYFP-C1 (Kan)</td><td>This study</td></tr>
<tr><td>CMV>EYFP-*Hs*ZC3HC1(1–462)</td><td>pEYFP-C1 (Kan)</td><td>This study</td></tr>
<tr><td>CMV>*Hs*SKP1(1–163)-EGFP</td><td>pEYFP-C1 (Kan)</td><td>This study</td></tr>
<tr><td>eEF1α>*Hs*ZC3HC1(1–502)-EGFP</td><td>pEGFP-N1 (Kan)</td><td>This study</td></tr>
<tr><td>eEF1α>*Hs*ZC3HC1(1–502 | C102S)-EGFP</td><td>pEGFP-N1 (Kan)</td><td>This study</td></tr>
<tr><td>eEF1α>*Hs*ZC3HC1(1–502 | W107A)-EGFP</td><td>pEGFP-N1 (Kan)</td><td>This study</td></tr>
<tr><td>eEF1α>*Hs*ZC3HC1(1–502 | W107F)-EGFP</td><td>pEGFP-N1 (Kan)</td><td>This study</td></tr>
<tr><td>eEF1α>*Hs*ZC3HC1(1–502 | W107Y)-EGFP</td><td>pEGFP-N1 (Kan)</td><td>This study</td></tr>
<tr><td>eEF1α>*Hs*ZC3HC1(1–502 | C112S)-EGFP</td><td>pEGFP-N1 (Kan)</td><td>This study</td></tr>
<tr><td>eEF1α>*Hs*ZC3HC1(1–502 | C117S)-EGFP</td><td>pEGFP-N1 (Kan)</td><td>This study</td></tr>
<tr><td>eEF1α>*Hs*ZC3HC1(1–502 | S118A,S119GG)-EGFP</td><td>pEGFP-N1 (Kan)</td><td>This study</td></tr>
<tr><td>eEF1α>*Hs*ZC3HC1(1–502 | C120S)-EGFP</td><td>pEGFP-N1 (Kan)</td><td>This study</td></tr>
<tr><td>eEF1α>*Hs*ZC3HC1(1–502 | C125S)-EGFP</td><td>pEGFP-N1 (Kan)</td><td>This study</td></tr>
<tr><td>eEF1α>*Hs*ZC3HC1(1–502 | H152A)-EGFP</td><td>pEGFP-N1 (Kan)</td><td>This study</td></tr>
<tr><td>eEF1α>*Hs*ZC3HC1(1–502 | C156S)-EGFP</td><td>pEGFP-N1 (Kan)</td><td>This study</td></tr>
<tr><td>eEF1α>*Hs*ZC3HC1(1–502 | W158A)-EGFP</td><td>pEGFP-N1 (Kan)</td><td>This study</td></tr>
<tr><td>eEF1α>*Hs*ZC3HC1(1–502 | C249S)-EGFP</td><td>pEGFP-N1 (Kan)</td><td>This study</td></tr>
<tr><td>eEF1α>*Hs*ZC3HC1(1–502 | W256A)-EGFP</td><td>pEGFP-N1 (Kan)</td><td>This study</td></tr>
<tr><td>eEF1α>*Hs*ZC3HC1(1–502 | W256F)-EGFP</td><td>pEGFP-N1 (Kan)</td><td>This study</td></tr>
<tr><td>eEF1α>*Hs*ZC3HC1(1–502 | W256Y)-EGFP</td><td>pEGFP-N1 (Kan)</td><td>This study</td></tr>
<tr><td>eEF1α>*Hs*ZC3HC1(1–502 | C272S)-EGFP</td><td>pEGFP-N1 (Kan)</td><td>This study</td></tr>
<tr><td>eEF1α>*Hs*ZC3HC1(1–502 | C275S)-EGFP</td><td>pEGFP-N1 (Kan)</td><td>This study</td></tr>
<tr><td>eEF1α>*Hs*ZC3HC1(1–502 | H363R)-EGFP</td><td>pEGFP-N1 (Kan)</td><td>This study</td></tr>
<tr><td>eEF1α>*Hs*ZC3HC1(1–502 | H425A)-EGFP</td><td>pEGFP-N1 (Kan)</td><td>This study</td></tr>
<tr><td>eEF1α>*Hs*ZC3HC1(1–502 | C429S)-EGFP</td><td>pEGFP-N1 (Kan)</td><td>This study</td></tr>
<tr><td>eEF1α>*Hs*ZC3HC1(1–502 | W431A)-EGFP</td><td>pEGFP-N1 (Kan)</td><td>This study</td></tr>
<tr><td>eEF1α>*Hs*ZC3HC1(49–502)-EGFP</td><td>pEGFP-N1 (Kan)</td><td>This study</td></tr>
<tr><td>eEF1α>*Hs*ZC3HC1(61–502)-EGFP</td><td>pEGFP-N1 (Kan)</td><td>This study</td></tr>
<tr><td>eEF1α>*Hs*ZC3HC1(72–502)-EGFP</td><td>pEGFP-N1 (Kan)</td><td>This study</td></tr>
<tr><td>eEF1α>*Hs*ZC3HC1(82–502)-EGFP</td><td>pEGFP-N1 (Kan)</td><td>This study</td></tr>
<tr><td>eEF1α>*Hs*ZC3HC1(1–101_159–502)-EGFP</td><td>pEGFP-N1 (Kan)</td><td>This study</td></tr>
<tr><td>eEF1α>*Hs*ZC3HC1(1–169_211–502)-EGFP</td><td>pEGFP-N1 (Kan)</td><td>This study</td></tr>
<tr><td>eEF1α>*Hs*ZC3HC1(1–169_189–502)-EGFP</td><td>pEGFP-N1 (Kan)</td><td>This study</td></tr>
<tr><td>eEF1α>*Hs*ZC3HC1(1–169_179–502)-EGFP</td><td>pEGFP-N1 (Kan)</td><td>This study</td></tr>
<tr><td>eEF1α>*Hs*ZC3HC1(1–202_237–502)-EGFP</td><td>pEGFP-N1 (Kan)</td><td>This study</td></tr>
<tr><td>eEF1α>*Hs*ZC3HC1(1–221_237–502)-EGFP</td><td>pEGFP-N1 (Kan)</td><td>This study</td></tr>
<tr><td>eEF1α>*Hs*ZC3HC1(1–235_252–502)-EGFP</td><td>pEGFP-N1 (Kan)</td><td>This study</td></tr>
<tr><td>eEF1α>*Hs*ZC3HC1(1–248_276–502)-EGFP</td><td>pEGFP-N1 (Kan)</td><td>This study</td></tr>
<tr><td>eEF1α>*Hs*ZC3HC1(1–290_398–502)-EGFP</td><td>pEGFP-N1 (Kan)</td><td>This study</td></tr>
<tr><td>eEF1α>*Hs*ZC3HC1(1–285_398–502)-EGFP</td><td>pEGFP-N1 (Kan)</td><td>This study</td></tr>
<tr><td>eEF1α>*Hs*ZC3HC1(1–340_412–502)-EGFP</td><td>pEGFP-N1 (Kan)</td><td>This study</td></tr>
<tr><td>eEF1α>*Hs*ZC3HC1(1–279_412–502)-EGFP</td><td>pEGFP-N1 (Kan)</td><td>This study</td></tr>
<tr><td>eEF1α>*Hs*ZC3HC1(1–419_450–502)-EGFP</td><td>pEGFP-N1 (Kan)</td><td>This study</td></tr>
<tr><td>eEF1α>*Hs*ZC3HC1(72–467)-EGFP</td><td>pEGFP-N1 (Kan)</td><td>This study</td></tr>
<tr><td>eEF1α>*Hs*ZC3HC1(72–290_398–467)-EGFP</td><td>pEGFP-N1 (Kan)</td><td>This study</td></tr>
<tr><td>GAL1>yECitrine / TEF>hphNT1</td><td>2µ/pMB1 (Amp)</td><td>This study</td></tr>
<tr><td>GAL1>yECitrine-*Sc*Pml39p(1–334) / TEF>hphNT1</td><td>2µ/pMB1 (Amp)</td><td>This study</td></tr>
<tr><td>GAL1>yECitrine-*Sc*Pml39p(1–334 | W119A) / TEF>hphNT1</td><td>2µ/pMB1 (Amp)</td><td>This study</td></tr>
<tr><td>GAL1>yECitrine-*Sc*Pml39p(1–334 | C134–137SGGS) / TEF>hphNT1</td><td>2µ/pMB1 (Amp)</td><td>This study</td></tr>
<tr><td>GAL1>yECitrine-*Sc*Pml39p(1–334 | C176S) / TEF>hphNT1</td><td>2µ/pMB1 (Amp)</td><td>This study</td></tr>
<tr><td>GAL1>yECitrine-*Sc*Pml39p(1–334 | Y257A) / TEF>hphNT1</td><td>2µ/pMB1 (Amp)</td><td>This study</td></tr>
<tr><td>GAL1>yECitrine-*Sc*Pml39p(1–334 | C271S) / TEF>hphNT1</td><td>2µ/pMB1 (Amp)</td><td>This study</td></tr>
<tr><td>GAL1>yECitrine-*Sc*Pml39p(1–334 | C292S) / TEF>hphNT1</td><td>2µ/pMB1 (Amp)</td><td>This study</td></tr>
<tr><td>yEGFP / TEF>hphNT1</td><td>pYM25 (Amp)</td><td>(Janke *et al*, 2004)</td></tr>
<tr><td>yEGFP / TEF>kanMX4</td><td>pYM27 (Amp)</td><td>(Janke *et al*, 2004)</td></tr>
<tr><td>mCh / TEF>natNT2</td><td>based on pYM43 (Amp)</td><td>This study (based on Janke *et al*, 2004)</td></tr>
</table>

#### Supplemental Table S2: Yeast strains

| Description | Genotype | Source |
|---|---|---|
| **Y187** | *MATα, ura3-52, his3-200, ade2-101, trp1-901, leu2-3,112, gal4Δ, met–, gal80Δ, URA3::GAL1*$_{UAS}$*-GAL1*$_{TATA}$*-lacZ* | Clontech |
| **CG-1945** | *MATa, ura3-52, his3-200, ade2-101, lys2-801, trp1-901, leu2-3,112, gal4-542, gal80-538, cyhr2, LYS2::GAL1*$_{UAS}$*-GAL1*$_{TATA}$*-HIS3, URA3::GAL4*$_{17\text{-}mers(x3)}$*-CYC1*$_{TATA}$*-lacZ* | Clontech |
| **AH109** | *MATa, trp1-901, leu2-3, 112, ura3-52, his3-200, gal4Δ, gal80Δ, LYS2::GAL1UAS-GAL1*$_{TATA}$*-HIS3, GAL2*$_{UAS}$*-GAL2*$_{TATA}$*-ADE2, URA3::MEL1*$_{UAS}$*-MEL1*$_{TATA}$*-lacZ* | Clontech |
| **wild-type (BY4742)** | *MATα, his3Δ1, leu2Δ0, lys2Δ0, ura3Δ0* | Dharmacon (Lafayette, CO, USA) |
| ***pml39Δ*** **(BY4742)** | *MATα, his3Δ1, leu2Δ0, lys2Δ0, ura3Δ0, pml39::kanMX4* | Clone ID 16507 (Dharmacon; Winzeler *et al*, 1999) |
| ***nup60Δ*** **(BY4739)** | *MATα, leu2Δ0, lys2Δ0, ura3Δ0, nup60::kanMX4* | Clone ID 10407 (Dharmacon; Winzeler *et al*, 1999) |
| ***mlp1Δ*** **(BY4742)** | *MATα, his3Δ1, leu2Δ0, lys2Δ0, ura3Δ0, mlp1::kanMX4* | Clone ID 17104 (Dharmacon; Winzeler *et al*, 1999) |
| ***mlp2Δ*** **(BY4742)** | *MATα, his3Δ1, leu2Δ0, lys2Δ0, ura3Δ0, mlp2::kanMX4* | Clone ID 12308 (Dharmacon; Winzeler *et al*, 1999) |
| **wild-type Mlp1p-mCh** | *MATα, his3Δ1, leu2Δ0, lys2Δ0, ura3Δ0, MLP1-mCh:natNT2* | This study |
| ***pml39Δ*** **Mlp1p-mCh** | *MATα, his3Δ1, leu2Δ0, lys2Δ0, ura3Δ0, pml39::kanMX4, MLP1-mCh:natNT2* | This study |
| ***nup60Δ*** **Mlp1p-mCh** | *MATα, leu2Δ0, lys2Δ0, ura3Δ0, nup60::kanMX4, MLP1-mCh:natNT2* | This study |
| ***nup60Δ mlp1Δ*** **yEGFP-Pml39p** | *MATα, leu2Δ0, lys2Δ0, ura3Δ0, nup60::kanMX4, mlp1::URA3, yEGFP-PML39:hphNT1* | This study |
| **wild-type Mlp1p-yEGFP** | *MATα, his3Δ1, leu2Δ0, lys2Δ0, ura3Δ0, MLP1-yEGFP:hphNT1* | This study |
| ***pml39Δ*** **Mlp1p-yEGFP** | *MATα, his3Δ1, leu2Δ0, lys2Δ0, ura3Δ0, pml39::kanMX4, MLP1-yEGFP:hphNT1* | This study |
| **wild-type Mlp1p-yEGFP, Mlp2p-mCh** | *MATα, his3Δ1, leu2Δ0, lys2Δ0, ura3Δ0, MLP1-yEGFP:hphNT1, MLP2-mCh:natNT2* | This study |
| ***pml39Δ*** **Mlp1p-yEGFP, Mlp2p-mCh** | *MATα, his3Δ1, leu2Δ0, lys2Δ0, ura3Δ0, pml39::kanMX4, MLP1-yEGFP:hphNT1, MLP2-mCh:natNT2* | This study |
| ***nup60Δ*** **Mlp2p-yEGFP, Mlp1p-mCh** | *MATα, his3Δ1, leu2Δ0, lys2Δ0, ura3Δ0, nup60::URA3, MLP2-yEGFP:hphNT1, MLP1-mCh:natNT2* | This study |
| ***nup60Δ*** **Mlp2p-yEGFP, Mlp1p-mCh** | *MATα leu2Δ0 lys2Δ0 ura3Δ0, nup60::kanMX4, MLP2-yEGFP:hphNT1, MLP1-mCh:natNT2* | This study |
| ***pml39Δ nup60Δ*** **Mlp2p-yEGFP, Mlp1p-mCh** | *MATα, his3Δ1, leu2Δ0, lys2Δ0, ura3Δ0, pml39::kanMX4, nup60::URA3, MLP2-yEGFP:hphNT1, MLP1-mCh:natNT2* | This study |
| **wild-type Mad1p-yEGFP, Mlp1p-mCh** | *MATα, his3Δ1, leu2Δ0, lys2Δ0, ura3Δ0, MAD1-yEGFP:hphNT1, MLP1-mCh:natNT2* | This study |
| **wild-type Ulp1p-yEGFP, Mlp1p-mCh** | *MATα, his3Δ1, leu2Δ0, lys2Δ0, ura3Δ0, ULP1-yEGFP:hphNT1, MLP1-mCh:natNT2* | This study |
| **wild-type Sac3p-yEGFP, Mlp1p-mCh** | *MATα, his3Δ1, leu2Δ0, lys2Δ0, ura3Δ0, SAC3-yEGFP:hphNT1, MLP1-mCh:natNT2* | This study |
| ***pml39Δ*** **Mad1p-yEGFP, Mlp1p-mCh** | *MATα, his3Δ1, leu2Δ0, lys2Δ0, ura3Δ0, pml39::kanMX4, MAD1-yEGFP:hphNT1, MLP1-mCh:natNT2* | This study |
| ***pml39Δ*** **Ulp1p-yEGFP, Mlp1p-mCh** | *MATα, his3Δ1, leu2Δ0, lys2Δ0, ura3Δ0, pml39::kanMX4, ULP1-yEGFP:hphNT1, MLP1-mCh:natNT2* | This study |
| ***pml39Δ*** **Sac3p-yEGFP, Mlp1p-mCh** | *MATα, his3Δ1, leu2Δ0, lys2Δ0, ura3Δ0, pml39::kanMX4, SAC3-yEGFP:hphNT1, MLP1-mCh:natNT2* | This study |
| ***mlp2Δ*** **Mlp1p-yEGFP** | *MATα, his3Δ1, leu2Δ0, lys2Δ0, ura3Δ0, mlp2::URA3, MLP1-yEGFP:hphNT1* | This study |
| **wild-type Mlp2p-yEGFP** | *MATα, his3Δ1, leu2Δ0, lys2Δ0, ura3Δ0, MLP2-yEGFP:hphNT1* | This study |
| ***pml39Δ*** **Mlp2p-yEGFP** | *MATα, his3Δ1, leu2Δ0, lys2Δ0, ura3Δ0, pml39::kanMX4, MLP2-yEGFP:hphNT1* | This study |

| | | |
|---|---|---|
| ***mlp1*Δ** **Mlp2p-yEGFP** | *MATα*, *his3*Δ1, *leu2*Δ0, *lys2*Δ0, *ura3*Δ0, *mlp1*::*URA3*, *MLP2-yEGFP*:*hphNT1* | This study |
| **wild-type Mlp2p-yEGFP, Mlp1p-mCh** | *MATα*, *his3*Δ1, *leu2*Δ0, *lys2*Δ0, *ura3*Δ0, *MLP2-yEGFP*:*hphNT1*, *MLP1-mCh*:*natNT2* | This study |
| ***pml39*Δ** **Mlp2p-yEGFP, Mlp1p-mCh** | *MATα*, *his3*Δ1, *leu2*Δ0, *lys2*Δ0, *ura3*Δ0, *pml39*::*kanMX4*, *MLP2-yEGFP*:*hphNT1*, *MLP1-mCh*:*natNT2* | This study |
| **wild-type yEGFP-Pml39p, Mlp1p-mCh** | *MATα*, *his3*Δ1, *leu2*Δ0, *lys2*Δ0, *ura3*Δ0, *yEGFP-PML39*:*hphNT1*, *MLP1-mCh*:*natNT2* | This study |
| ***mlp2*Δ** **yEGFP-Pml39p, Mlp1p-mCh** | *MATα*, *his3*Δ1, *leu2*Δ0, *lys2*Δ0, *ura3*Δ0, *mlp2*::*URA3*, *yEGFP-PML39*:*hphNT1*, *MLP1-mCh*:*natNT2* | This study |
| ***mlp1*Δ** **yEGFP-Pml39p** | *MATα*, *his3*Δ1, *leu2*Δ0, *lys2*Δ0, *ura3*Δ0, *mlp1*::*URA3*, *yEGFP-PML39*:*hphNT1* | This study |
| ***nup60*Δ** **yEGFP-Pml39p, Mlp1p-mCh** | *MATα leu2*Δ0 *lys2*Δ0 *ura3*Δ0 *nup60*::*kanMX4*, *yEGFP-PML39*:*hphNT1*, *MLP1-mCh*:*natNT2* | This study |
| ***nup60*Δ *mlp2*Δ** **yEGFP-Pml39p, Mlp1p-mCh** | *MATα*, *leu2*Δ0, *lys2*Δ0, *ura3*Δ0, *nup60*::*kanMX4*, *mlp2*::*URA3*, *yEGFP-PML39*:*hphNT1*, *MLP1-mCh*:*natNT2* | This study |
| ***nup60*Δ *mlp1*Δ** **yEGFP-Pml39p, Mlp2p-mCh** | *MATα*, *leu2*Δ0, *lys2*Δ0, *ura3*Δ0, *nup60*::*kanMX4*, *mlp1*::*URA3*, *yEGFP-PML39*:*hphNT1*, *MLP2-mCh*:*natNT2* | This study |
| ***nup60*Δ** **yEGFP-Pml39p** | *MATα*, *his3*Δ1, *leu2*Δ0, *lys2*Δ0, *ura3*Δ0, *nup60*::*URA3*, *yEGFP-PML39*:*hphNT1* | This study |
| ***nup60*Δ *mlp1*Δ** **Mlp2p-yEGFP** | *MATα*, *leu2*Δ0, *lys2*Δ0, *ura3*Δ0, *nup60*::*kanMX4, mlp1*::*URA3*, *MLP2-yEGFP*:*hphNT1* | This study |
| ***nup60*Δ *mlp2*Δ** **Mlp1p-mCh** | *MATα*, *leu2*Δ0, *lys2*Δ0, *ura3*Δ0, *nup60*::*kanMX4, mlp2*::*URA3*, *MLP1-mCh*:*natNT2* | This study |
| ***pml39*Δ *nup60*Δ *mlp1*Δ** **Mlp2p-yEGFP** | *MATα*, *his3*Δ1, *leu2*Δ0, *lys2*Δ0, *ura3*Δ0, *pml39*::*kanMX4*, *nup60*::*URA3*, *mlp1*::*HIS3*, *MLP2-yEGFP*:*hphNT1* | This study |
| ***pml39*Δ *nup60*Δ *mlp2*Δ** **Mlp1p-mCh** | *MATα*, *his3*Δ1, *leu2*Δ0, *lys2*Δ0, *ura3*Δ0, *pml39*::*kanMX4*, *nup60*::*URA3*, *mlp2*::*HIS3*, *MLP1-mCh*:*natNT2* | This study |
| ***nup60*Δ** **Mad1p-yEGFP, Mlp1p-mCh** | *MATα*, *leu2*Δ0, *lys2*Δ0, *ura3*Δ0, *nup60*::*kanMX4*, *MAD1-yEGFP*:*hphNT1*, *MLP1-mCh*:*natNT2* | This study |
| ***nup60*Δ *mad1*Δ** **Mlp1p-mCh** | *MATα*, *leu2*Δ0, *lys2*Δ0, *ura3*Δ0, *nup60*::*kanMX4*, *mad1*::*LEU2*, *MLP1-mCh*:*natNT2* | This study |
| ***nup60*Δ *mlp1*Δ** **Mad1p-yEGFP** | *MATα*, *leu2*Δ0, *lys2*Δ0, *ura3*Δ0, *nup60*::*kanMX4*, *mlp1*::*URA3*, *MAD1-yEGFP*:*hphNT1* | This study |
| ***pml39*Δ *nup60*Δ** **Mad1p-yEGFP, Mlp1p-mCh** | *MATα*, *his3*Δ1, *leu2*Δ0, *lys2*Δ0, *ura3*Δ0, *pml39*::*kanMX4*, *nup60*::*URA3*, *MAD1-yEGFP*:*hphNT1*, *MLP1-mCh*:*natNT2* | This study |
| **wild-type Nup1p-yEGFP, Mlp1p-mCh** | *MATα*, *his3*Δ1, *leu2*Δ0, *lys2*Δ0, *ura3*Δ0, *NUP1-yEGFP*:*hphNT1*, *MLP1-mCh*:*natNT2* | This study |
| ***nup60*Δ** **Nup1p-yEGFP, Mlp1p-mCh** | *MATα*, *his3*Δ1, *leu2*Δ0, *lys2*Δ0, *ura3*Δ0, *nup60*::*URA3*, *NUP1-yEGFP*:*hphNT1*, *MLP1-mCh*:*natNT2* | This study |

### Supplemental Table S3: Primer sequences

| **Target** | **Sequence** |
|---|---|
| *Dd*ZC3HC1_for1 | ATGGATGAGAGAATTAAAAAAGCACTAAGCGATTTAG |
| *Dd*ZC3HC1_for346 | GAAAAGGATAAAAAATCAAGTGTATATTGTTCATATTG |
| *Dd*ZC3HC1_for486 | AGTTTATTTTCAATAGTTGGTAATGGATTCTCTAAAG |
| *Dd*ZC3HC1_rev271 | TGAAATTGAATTAAAATCCCAACCACATAATGC |
| *Dd*ZC3HC1_rev426 | TGTTCGGGCAAATGATTGATTTAAAACTTTTTC |
| *Dd*ZC3HC1_rev635 | TTTCCTATAATGATGGATTGAAGTTGTTAATGAGTTTAC |
| *Dd*TPR_for1 | ATGACATCTGTTAGTGATTCAAATAATC |
| *Dd*TPR_for282 | GCATCACTTTACCAAGAGAGATCAGAGGAA |
| *Dd*TPR_for660 | GGTATGATGACATTATCAGATTTATC |
| *Dd*TPR_for921 | GAAACCTCAATAGCAATGACTCATCAAATC |
| *Dd*TPR_for1219 | CTCTCAAGAATTGGAACAAGCCAAAC |
| *Dd*TPR_for1351 | ATGCGTACTCTTACCGTTAG |
| *Dd*TPR_for1630 | ACCACTCCAACTGTTGTTTCAACTCCAACT |
| *Dd*TPR_rev659 | ACCACTACTGCTATTATTGTTG |
| *Dd*TPR_rev1350 | ATTCTCTTGTTCTTCTTGAAGTTTC |
| *Dd*TPR_rev2052 | TTATTCTTGCGATGGTTGATTATC |

### Supplemental List of Sequences for ZC3HC1 Homologs

#### Representative sequences for Figure 3A:

**>NP_057562.3[*Homo_sapiens*]**
MAAPCEGQAFAVGVEKNWGAVVRSPEGTPQKIRQLIDEGIAPEEGGVDAKDTSATSQSVNGSPQAEQPSLESTSKEAFFSRVETFSSLKWAGKP
FELSPLVCAKYGWVTVECDMLKCSSCQAFLCASLQPAFDFDRYKQRCAELKKALCTAHEKFCFWPDSPSPDRFGMLPLDEPAILVSEFLDRFQS
LCHLDLQLPSLRPEDLKTMCLTEDKISLLLHLLEDELDHRTDERKTTIKLGSDIQVHVTACILSVCGWACSSSLESMQLSLITCSQCMRKVGLW
GFQQIESSMTDLDASFGLTSSPIPGLEGRPERLPLVPESPRRMMTRSQDATFSPGSEQAEKSPGPIVSRTRSWDSSSPVDRPEPEAASPTTRTR
PVTRSMGTGDTPGLEVPSSPLRKAKRARLCSSSSSDTSSRSFFDPTSQHRDWCPWVNITLGKESRENGGTEPDASAPAEPGWKAVLTILLAHKQ
SSQPAETDSMSLSEKSRKVFRIFRQWESLCSC
**>XP_041459241.1[*Lytechinus_variegatus*]**
MAASNFETSKPRRIKALLSSFLKGISREEKKESETKQVIFESLDTEGFADGFVVVDEVSSEQTSTVQPLNQELFFNRVETFSISSWFAKPDEVC
PLRCAQYGWENIDVDSLKCVSCKEVLYGGLPPKWETDLYENACKKLVDSLKTGHSKICPWQSNPSPASFLEVNLVSSQNAVNDFLHRVASIRCF
GTSVPAVDLSCLQQVDENEDALSRIVSGVLGEEPTCDYKERIESVCFMAACGWSRSSPEGSQSPTMSCQYCRRNVGLWNFTPYEQNAKTVTEDD
SEPTAKRLKVDKGLFNPIEEHRSWCPWIKPTSSTQKVKSLPQDKNQDDERPVRPAWHELLVLLHQRSSPDKQGLLTNKQVTPPSQAWKAVRRIT
NFWQSRNAVNKT
**>Own_assembly[*Saccoglossus_kowalevskii*]**
MVSDKMAEKSSVLVTPKRVHDLLSSFIHKEETGVDERSDEPNQENIAQSSRQFLPRNREAFFARLETFSAFTWFAKPIELSPLKCAQYGWENTD
NDIVKCVSCKEIVCASLPKTWDPDLYAKRCEELRAALVKSHSNICPWRDSPSPDIFLSIPLLNQSEVQGDVLSRCTSLEKLGRKLPVIETCDIE
SQITAEALSGVVTRQINLLEDIRNDDDSSAINTVCVLSLCGWSTSSCETSQYPTVSCQYCRRQAGLWNFTPVVEPEPDKMTSEEQPTVDSSVES
SDADSAQEPSMKKRKISESKKSAFNPITEHRAWCPWVISYVMQQSDDHKSSPVPTYNNIPGWQAVFTLLAPKTSPLKENIQRLTDKDQTPPNQV
WKAVRKILSFW
**>XP_005104988.1[*Aplysia_californica*]**
MAATSTTNDTQTHTPEKVKNLLSSFLASSEQTRTEAITRGNASVQGEKTFPSDGTAYDLRPRLACSTSVLRSYDSYLQRLDTFSSLTWFNKPAE
LNPLICARYGWENIDTDMLQCVGCKAFLCGQLPVKTNPEVYEESLSKLKKNLLAAHDKFCALAVNPCPESFCRVPLHDPTNLTAEYTERASKLS
QIQERLPVIDYTRLHELEYDEGQGAAYCKKHLMPDSDVSPQAVTLAFTGWTTSSQNVLVCCMCRRQVGLWNYAQESSRGLAPDSTTKESSDEDE
ESKESGEPEAKKRKMKVSIKQKFDPIEEHWHWCPWVTETEVPCSSPPSGQQAAPAQRQKSLAFITAIKVTAPGLMDNNTGLAHAMKTSPMVEGL
RCFRRVMKSWSSPKLLSAGNSPT
**>XP_013395661.2[*Lingula_anatina*]**
MQAGFAQRVRTILSSFLHNETDDKKKELKNGHAVEEGDQSPAPSSSPCEAGVTARSGEVQLPTHARDRTAYFHRVETYSAVSWFAKPPSLCPLQ
CARYGWQNVDTDMLRCVSCKAVLCASLPKTYDHDVYDESCKKIAESLVSSHRKICPWPSNPSPVHFMSILYSGRDEALDDLLSRHQNLKQLGSQ
LPKLDIQADHELSGPSVQNLFKIVKQQGSSGGFNLDSALTDQVSCVLALCGWDISSLKKTDILICHFCRRQIGLWNFLSHSSLEQETDQSSQSV
IENGSHSKKEGSASPSPKKRKFLNDSIAKTSFNPIGEHRSWCPWITSTQQDKSNSQKSDQREVVSSYVGDEDSRPGWKRLLDWLVPNSASTDKL
TPKFVRNVKTTPPSEGLKNIRKVLHDWSSPTIGKL
**> Own_assembly[*Capitella_teleta*]**
MDERIKSALKTLSDTAALTKSGDGPNNSLDEADAPSSIHEDRAAFFDRLQSFSFANWFAKPLWLSPIACARYGWKNVDKDLLECVGCKKPMAGK
LPQSADPIISAKCHERLYQALTSAHHNSCAWKFHPTPVRVFAVPHHNHANATDQFVSRGSSFLALASRLPQIDPSVMESFGISLPVLETLYHVT
ATDGAYLLHDGNWQLLTEKASCDEDLATIPNLLQSACLLSITGWMLHSSKPGNEQIRCSLCQRLIRLAEVQNFTEKHPEMEAESSADHFLESII
ALDAVMKMQRTERSETDQQMAPQTPTVKSTEVVESIMNELLTAVCARNEEQREEPMEIDFRDAISKAKRSAAKKAKLDSKPTLNLLTEHRNWCP
WVSDGSCAHLDKDFKESSSDDFVPGWKTLLHILLPKESANQLNQSVEGLKKIRRSLHQGI
**>PAA86276.1[*Macrostomum_lignano*]**
MSQSEVESAIALFKTTVDDFDDSDNELHLLVSTQPLTSTEGIIKKRTLAGLLARLKTFSSLSWSVKPIELSPLVCAQYGWIGKKPNLLVCVTCK
NNLCIKLPSDSKLYNHCLKEAIEKLSSNHDNLCPWKLFPAIDAEPVLKFTNPAQDFEILNSIADSLSCLANEIDDSLQFSIGVDADTAEDFLNC
IAGAAATSSEVGSDEQLQIRRRQRCFQLAITGWSRPSPDSDELLKHFKSSQCPSILSCQFCLRRLGVWNLVRLANRQERQQHGEESTESEVNLS
TGNLFNNDSNSHIGDQLEGGVEQSRDDADNSDDAVVSQSSEQPASKRARLSEAPATAAAKLELLSLHWLWCPWSSGQVTKAWCDSVLLASPKRR
RRKPAIPTTTDQQQISTVAADGCAKVMQLLRSIV
**>NP_501317.1[*Caenorhabditis_elegans*]**
MEVDTASSRHSTVLKRKATDSINEILNYGQSSTSPQKRCKKAASLHKYRDMETYHKIIKTYKAPTWYGCAVSPRDLADYGWACVKKDCVKCIEC
EQYLSTVLPNICKVSFNVYNSSLQDIHEKMTTAHRTTCKLRCGAPPFRIVEPTAKEVMDGIQRRLSDSKKIIDEDLKADIPSDVNLPKIEGVPE
QLIYVAALGWHVSMPKRGSLLFGCDNCARELAIRCGNKFDPIHNHERWCPRIEMDDHGEPSWQSDLNTVLNTKNHVTNRYTGSSIFKEAYAARR
LLDSSLSTIITPNYI
**>XP_046451818.1[*Daphnia_pulex*]**
MATLNRKRKLDQTLIQKLYHDVPSKDPTRESFLKRLKTYDVFNWSGKPVDPPLCALHGWEIAEKDVLKCVMCHQFMSVTLPSPTKDAPYKHACS
KLKSRLASAHSKFCLYSTNQVPDSVLEIEHVSNMELQDTIQQQLLAFKNIEALCKVSELQDEIKEIFDWFLETTAIPDVHLSSFTFVLTGWKFL
QDDMLLKCDYCNRKWSIEPYLSQKNLTEKNDSVRTTVDPVAQHQRWCAWRAPTRGWKSRLLQLQQLKESRCREKRSRLSSDSCDSLTDRMRTVR
KLLNGTL
**>XP_002154618.3[*Hydra_vulgaris*]**
MSSANTKNKITDILSSLVSTTPEKSSSHVTCNPQSKDLFLQRVKTFTSSNWVAKPVGLSPLHCAQYGWCTEYLDQLRCVTCNATLDAGLPDEWD
EAAYNEICNKVQNKLQIGHEKLCPWPDNPCPPSFLSLPSYTSEQWCAEMKLSFESLMTLRGNLPELNEDEIASLGVLDNSSIETMLNQVFKWSS
ENDDDALQAKVASILAICGWSVCQPVVEDPSIICTICGMEAGLWNYKSLSSRINHQKRFKTSLEYTNSQQSLTELQENMSTDVSVKSEKSETEK
SDNSGQGSMLNELSRLATSSDSLAQSDLLRAAESQESLLNLRVSDSKVMATDGLEHHIETTHEDFNQLKNQFSEPRLPQATHPELMKELLLSRA
PGSRTRSRYSDSIFSEVGSECFEKRLEETCEKDIEHERAITDNPPPNHPNSSQQQWMKELLFITADACHSEPTESVISEIGSVAFDRRIEDNSS
LNSPRLLTPVHHGMHDYDEDSEPSRIKKMRFQVPDISMFSVIGEHRFWCPWVCSTERFTDHETSLSDSTAFISCKKIPGWKYVLYQLLPYNRPS
PCSTQRHEAWRYVRSTLTECISNKIT
**>Own_assembly[*Trichoplax_adhaerens*]**
MAEISLVPRKFLSLLDSFIFDKPACPENECKQDKVAEDYFERVETYSAYTWLAKPPALSPLQCARYGWKNVDIDRLKCVTCSATLTVSLPLPST
RSQPYEEAVRTFTDALISRHNPLCPWKDAPVREELIRLPNTYAKCVPILRQRLKSYSSAPHLLPRLELNSTLNTLLSCKEAVETLKDILDCQVI
LTDVDEEAQMVACILAFCGWVLKPSDKLKSSLLICSICRRRAGIWNYIMAISDSSTDSLELSSESIEEQSSSSPSKRRKIEKKKFNPIEEHRYW
CPYFIVDDNNAESREDVERSLGNVHQDNNNDTAGLKSMLSYLFPKRHSRTTDPNLNSCSAKSLRISTIPHQDLCSRSYIIMSLEEEV
**>NP_013600.2[*Saccharomyces_cerevisiae_S288C*]**
MEKDALEVRLKSIRHSLDKNTKLLPGKYRNTLGERLITKWRYKKKSHNGSSMLPEKCKSHVQLYDDLVQESSKHFVGFRLHDLRALLKRICSIQ
NYTRHVLIEWDVRWVNPLTLASKGWEPYQSASQSQVPFKCCCCHAIMTIPLLKNGDDVADYTMKLNEKIWNSNIIGNHLQKCPWRENQVDLNKE
YYLSSQNLIREIERIHTEIDRIVSGSNEFSLKRNSSRIFHYLSEKEIQKLAFFFDCKDYSLVGLLLLGYTKFQKDDLVQCTACFHRASLKKLEY
TEFNGHALWCRYYNKELLPTMLLELIGKEDKLITKLGVGERLNKLEAVLQTL

**>XP_024513553.1[*Cryptococcus_neoformans_var._neoformans_JEC21*]**
MELSSNTDDDLRDVFKLLYADDDWALTSDSELDDSEQLGNADGSEIDVADDEEQHTIRIYSGRITKKRLFSALDSLLSPGYETDTKRQRIYNPP
APSIPSLILSTQPMPALPLSKVYAPFSALSLLSRLMTFQPYTYSPQHPLTLSPVRAAMKGWVNEGREGLKCDVCGARWGLGGLEKVRDEAMKSN
LGERLAKGFEERHEKNCAWRICASPGNLYEQLRHLVHPPITSSLAPLASHLLLECLALPSLRLLSPLNPLQVERLVSLFKPSSTFSIPSPATDV
ASQLALFGWFPYHPNYPTIQISLNTPSSRTEIVCCRICHRRIGLWNFSNEKDGVKRFDVLNEHLVWCPVRIQDGEKEWWSESGLLDGQSTQAKR
IGEGGIKGLVKVSEKMEKRSWRRS
**>ON368701[*Dictyostelium_discoideum_AX4*]**
MDERIKKALSDLDNATVLNQLPILSNDLTTTCGSSSGSSSNDNNNNNNKNNNQYSTLNLIDESNNSTSNSTTSPSLLITSYRPWSNTDYYNRVR
TYTISNWFAKPCEIDPLQCSRFGWINCEADMLECETCKKRLYYKVPSTFSQSLVNKRINDFSISLQSTGHRDNCPWKDNGCPSFFSRLLDIPFQ
TQLEAYIKRSQNIYNNLTTLPMLSSDFYQQWVNKQNLMEPPITSRTNNILNIIVKIAKLPTDEVKSKVSCLLALCGWDFNSISNSNNNNNNNNN
NNNNNNNNNNNNNNNNNNNNNNNNNNNNNNNNNNNNNNNNNNNDDDDKNEKDKNKNKIKENEKEKDKKSSVYCSYCQRLCGVWNFNKIKPNSTS
PFNEENINTTNNKGFNNSNIGSKRKREEDIEEEKRNIQFEKVLNQSFARTNNNNNNNNNNNNNNNNNNNNNNNNNNNNNNNNNNNNNNNKKLNS
NSNSNNNSNNNSNNNSLFSIVGNGFSKVTSNSQSTGWDWGNSFSNRISDFKAALEIANATEKKKEFSPINEHRWFCPWMIVVDSNRLIIDNNDI
LGENSQDNNNNSGSSNNSSNSNSSISGWENLLKLLLNQSTFDSKDFIDLKNDKKFHSIVNSLTTSIHHYRK
**>NP_175325.2[*Arabidopsis_thaliana*]**
MAQDSEKRFHQIMDKLFTPSKSQLPSSSTSSSVEQQSRGKKRQNPSSALALVEPKIVLATIDRSSALKVPAGTSPSGLCRPWDRGDLMRRLATF
KSMTWFAKPQVISAVNCARRGWVNDDADSIACESCGAHLYFSAPSSWSKQQVEKAASVFSLKLESGHKLLCPWIENSCEETLSEFPLMAPQDLV
DRHEERSEALLQLLALPVISPSAIEYMRSSDLEEFLKRPIAPACSDTAAESSQTESLTNHVGASPAQLFYQAQKLISLCGWEPRALPYIVDCKD
KLSETARGTETIDLLPETATRELLSISESTPIPNGISGNNENPTLPDTLNSDPSSVVLDCKLCGACVGLWVFSTVPRPLELCRVTGDTEINIEK
HPKGGTLQHQPSSLKFTIAGGPPATKQNFKATISLPIIGRNLRSRFASYSRDHDHGDVSSIQDQQSRTAENNGDVTQNSNQVMNDIGEKADGGR
NSTDVESDIALQNKDKQMMVVRSNLPENNKPRDSTAEKSATSNKQMEFDPIKQHRHFCPWIWSTGRRGPGWRQTLSALQRHKGSCQTPPSSSSL
FKVDDPLTSVRNLFKSPSPKKRKLNGGSSS
**>NP_173164.1[*Arabidopsis_thaliana*]**
MKEEDVSSQNVNPRSNRNSVASASASASATPVDRFRRRARSPSPPQTAAASSAGASSPAVLVNAGSVDWTGHGLALSVRSCRTWDRGDLLRRLA
TFKPSNWLGKPKTASSLACAQKGWVSVDLDKLQCEYCGSILQYSPPQDSLNPPEADTTGEKFSKQLDDAHESSCPWVGKSCSESLVQFPPTPPS
ALIGGYKDRCDGLLQFYSLPIVSPSAIDQMRASRRPQIDRLLAHANDDLSFRMDNISAAETYKEEAFSNYSRAQKLISLCGWEPRWLPNIQDCE
EHSAQSARNGCPSGPARNQSRLQDPGPSRKQFSASSRKASGNYEVLGPEYKSESRLPLLDCSLCGVTVRICDFMTTSRPVPFAAINANLPETSK
KMGVTRGTSATSGINGWFANEGMGQQQNEDVDEAETSVKRRLVSNVGLSFYQNAAGASSSAQLNMSVTRDNYQFSDRGKEVLWRQPSGSEVGDR
AASYESRGPSTRKRSLDDGGSTVDRPYLRIQRADSVEGTVVDRDGDEVNDDSAGPSKRTRGSDAHEAYPFLYGRDLSVGGPSHSLDAENEREVN
RSDPFSEGNEQVMAFPGARDSTRASSVIAMDTICHSANDDSMESVENHPGDFDDINYPSVATAQSADFNDPSELNFSNQAQQSACFQPAPVRFN
AEQGISSINDGEEVLNTETVTAQGRDGPSLGVSGGSVGMGASHEAEIHGADVSVHRGDSVVGDMEPVAEVIENLGQSGEFAPDQGLTDDFVPAE
MDREGRLGDSQDRVSQSVVRADSGSKIVDSLKAESVESGEKMSNINVLINDDSVHPSLSCNAIVCSGYEASKEEVTQTWESPLNAGFALPGSSY
TANDQGPQNGDSNDDIVEFDPIKYHNCYCPWVNENVAAAGCSSNSSGSSGFAEAVCGWQLTLDALDSFQSLENPQNQTMESESAASLCKDDHRT
PSQKLLKRHSFISSHGKK
**>XP_042921239.1[*Chlamydomonas_reinhardtii*]**
MSSVYERITSALSSLGKRRERDSSAHEGDGAGASAAGGGPTSPGGRSAAARTPKRFRPWEQADLHKRLETYKPLTWFGKPASVGPVPCALKGWV
NDGSDCLTCEYCGSKLVYPPHVAYDQRQAAADMFSPSLTTKHTATCPWRQTACQPKLLAYVPSTTPEQLCSLFYSLADKLMRVDVLPDMDTLAI
QTLRSTAMPYGSYDDFITAAAPGGGAVVGGGGAAGYSHDLAPRRRQMPSATIRELDQNGDEVMTPSGSAAGAAAAMAPAAAAGGGGDAAAVLQA
LVAAGDAGEGQAVLVQTSKLAPAQKARLLALLGWDVDVLQPDSASGMAVAPFAAGGSYSLSHLGVKPKAAAAAAAGAGAGGAAVPGTPGGAGGK
GGKGSSKVPSSQVVLKCPICNSRMGLWNYSGVRPVPVGRLTAPPPPAAGGAAALMLSPRPAAASSGGGAAAAAAPAVPATIGSDPLSCTIAGGQ
YGQFGFGGAASAAKPFGSAAAAAPFRFGSAASTAPVFGLAAMDVDAQRAASAASGPFGSAAAAAAATPSMPAPSGSATPAPAGRKRKAEAPEPM
ALDAQHTPSAGMATPVAAPDGKRQRMAATPLWGGAGFGAVGGPAASPSGLGLGGASALAAASAAGQPRELDPVAQHRSWCPWVYTGSGDEKHMS
GWQHMLSALSQHQQHQQQQANVAATPGAAAASPADARQLRDNALEAIRKL

#### Representative sequences for Figure 3B1:

**>NP_057562.3[*Homo_sapiens*]**
MAAPCEGQAFAVGVEKNWGAVVRSPEGTPQKIRQLIDEGIAPEEGGVDAKDTSATSQSVNGSPQAEQPSLESTSKEAFFSRVETFSSLKWAGKP
FELSPLVCAKYGWVTVECDMLKCSSCQAFLCASLQPAFDFDRYKQRCAELKKALCTAHEKFCFWPDSPSPDRFGMLPLDEPAILVSEFLDRFQS
LCHLDLQLPSLRPEDLKTMCLTEDKISLLLHLLEDELDHRTDERKTTIKLGSDIQVHVTACILSVCGWACSSSLESMQLSLITCSQCMRKVGLW
GFQQIESSMTDLDASFGLTSSPIPGLEGRPERLPLVPESPRRMMTRSQDATFSPGSEQAEKSPGPIVSRTRSWDSSSPVDRPEPEAASPTTRTR
PVTRSMGTGDTPGLEVPSSPLRKAKRARLCSSSSSDTSSRSFFDPTSQHRDWCPWVNITLGKESRENGGTEPDASAPAEPGWKAVLTILLAHKQ
SSQPAETDSMSLSEKSRKVFRIFRQWESLCSC
**>NP_001186366.2[*Gallus_gallus*]**
MAAPSAEEAGGGRSRPPAVTPQQIRDLIDGGIASEGSGPEGKGTSDWSESANGSLQIDALSSESTSKEAYFSRVETFTPLKWAGKPHELSPLVC
AKYGWTNVECDMLKCSSCQAFLCVSLQLTFDFNKYKERCVELKKSLCTAHEKFCFWPDSPCPDRFALLLVDEPRALLQDFLERFQNLCQLELQL
PSLRAEDMKNMSLTEEKISLLLQLIKEELEHRTEGEKPPMKFASESLQVHIPACVLALCGWTCSAVSGSVLSVITCSRCMRKVGLWGFHQLESA
GLELDSWSPSTASTASGERGPPVPTSPRRMLTRSQDTNSPGSEQEKSPSPSISRLKGSDPPSSPVERGGELEATSPTQRNRPITRSMGQGDNVE
VPSSPLRRAKRPRLCSSSSSDTSPRSFFDPSSQHRDWCPWVNAVEGGETPEDPEKEPAKGEPGWQVVLSTLLASRKCDRVPETEPVSLSVKSCK
VFRIFRQWESINPS
**>XP_003228779.1[*Anolis_carolinensis*]**
MAAPSPAAVSSALPSEAEESKGKPASVTPQKIRELIDGGIAPEETSLEGKDLSALYEVANGSPKTEELPFEATSKEAYFNRVETFTSLKWAGKP
HELSPLICAKYGWTNTECDMLKCASCQAYLCASLQLAFDFSKYKERCLELKKALSTAHEKFCFWPDNPCPDRFSVLLVDEPLALLSDFLERFHS
LCRLELQLPSLKPEDLKSMSLTEEKISQLLQLIEEEADCKAEGEKTPSRKPLDLLQIHITACVLALCGWTSSPSSGSIQLPLISCCRCLRKAGL
WGFHQIESAPPETEVSPGLATPDGRPSSSSDKAGPTVPTSPRRMMTRSRDTTLPPGSEQQEKSPSPVISRMRSWDSTSSGERGEPESASPVPSP
RSRPVTRSMGQGDISGLGAEVPSSPLRKAKRARLCSSSSTDSCARSVFDPASQHRDWCPWVNAVKERPALEAEAGNQVEEGKAALGWQAVLKAL
LATKQSEGPADAESENLSAKSRKVFRIFRQWEAACSS
**>NP_001011259.1[*Xenopus_tropicalis*]**
MATSCEDVSPVKSPAVTPLKIRELINEGIVTGERSSIGRKETAVVPEENNGFEDPLSNSSYESTSKDAFFGRVESFSSLKWAGKPSELCPLICA
KYGWSNIECDMLKCSSCNAYLCASLQPVLDFSKYKQRCVELQEALRKAHEKFCFWPDSPCPDYFWALMVTEPSSVLSDFVGRFDNLCHLEIQLP
SIKHEDLKNMDITEETVSHLLRLIEDELKSKDGREDNSRLASDSLQVHISACILALCGWSTSYTSGSLCIINCPRCMRKVGLWAFQQLEAVELD
NSLSAPNTPVSPAEGHERSPFGIMSPNRRVTRSRDAEQSPALAYGRTRSSDLLSPADSEAVRSRPVTRSMGQGESSGLSNELHSSPLRRSKRPR
LCSSSSSDTSPRGCFDPLSQHRSWCPWVNVCQASETSTLGSEIQEEASRKEYGWKEVLNVLLAEENSRTLSDPDTSSVPEKSHKVFRIFRQWQM
AASASENP
**>NP_001070846.1[*Danio_rerio*]**

MAALGSRANRPENGEKQTKSPLVSPLKVRELLNEGVASEDSVLNCSQQDPNTASPNGLGKAPCEAANKEAFFNRVESYSCLKWAGKPSVLSPLR
CARYGWINVDCDMLKCCSCQAFLCASIQATLDFQKYKGRISEVRQQLQTQHEKFCSWPDFPCPDRFWMVPINEPTVLLAAFLERYKSACLLEQQ
LPAMKPEQLKAMTLTEDIISVLLQLIEDEQAKQGSSPSKVSSDPLSVQVAACILALCGWAASPSLHALNLPILACSYCMRKVGMWNFYQMDTAL
EAENPPQSPASSTSVTSTQGMSGDKGTPTSPSQSPTPCRMKLRSQDSTRSEQAESTPLRTRSRDSPTPHDEHPSPLSRGKRPMTRSRGQGEGQG
TDVPSSPQRKTKRPRLSSASGPEGPLHRNMFDPVAQHRDWCPWVSVEKEEDNQDASDFVGCEAELPQPGWKAVLALFLSMKQSLNPVGASPSQG
PHDKSKRVFSIFRQWQVSSPSQ

**>XP_042191218.1[*Callorhinchus_milii*]**
MATEGGEENAETKSGKDAARTPQKVRELLSDSVAPTDQTNSISESPNASLEESIPPCDSANKEAFFLRVETFTSLKWAGMSFEFSPLYCAKYGW
VNVDCDMLKCSSCQALLCLSLQPTQDSTKYKERVTELHKALKTAHEKFCYWPDSPCPGRFWALPFKEPSVLLSGLTERFRGLCQLEFQLPTLKH
DDLKDMTLTEDTISFLLQLIEDEVKGSAANESSGLKNTTDILSTHVAACILALAGWAAGPSDSLQLPIIMCSYCMRKVGLWSFQQIETLGGGSE
IDLPISLCNTPISPQENKAGRSTPTSLTISPHRMVTRSQDAAQSLPGEQQELSSSPITPRTRSRDTHSPTPVDRSESDSVSPGLRGKRPATRSK
GQGEVPSSPQRKPKRLRLSSSTSSDSPKSYFDPVFQHRDWCPWITRDGEPDRSEDPEQGEVGAVPDTLERRELGWRTVLQVLLSLQPSRSPDEE
SDSFSLSEKSRKVFRIFRQWQVTCSS

**>XP_032822323.1[*Petromyzon_marinus*]**
MWKSVSGWVPIPPPFTTMDASQTPSSPERATPEKIRQLLSTFISPEKSPDVGQKSPETRVILSRNRERFLKRVETFTPSAWAAKPSCLSPLCCA
ALGWECVARDVLCCCSCRTAQCIVLPPIWETARYDEKLKEVTESLKTSHAKYCTWPDDPCPDWLFLLPLHDPAQLLSSFCSRFQQLHALPSLSP
AALHDVGISEDTMKLLLQALAPAKRAEHGGDAETDGKNVCGREGSNGKNDSLEGATKGEVEIAGATSAMDNGEGKVSGGRGTNGTGGHDASASG
KDDGKEVLSETEGGEGETNGRSGKGEGTPSNVDADSGVQGKDVEGEAAVVQKEAAPGASSEVGEKVDDGVKVDDVSNVDGGATVDDGAKVDTGA
KVDEGAKVDDGAKVDTGAKVDEGEKVDDGAKVDDGAKVDDGAKVDDGAKVDDGAKVDDGAKVDDGAKVDDGAKVDDGAKVDDGAKVDDGAKVDD
GAKVDDGAKVDDGAKVDEVTVDMEPGAVAACVLALCGWDASPNVKSIVSCTICCRRIGLWNFKTQDADSERSCDTMTVGNVTPCGRPSPASAAV
ISPASQQNLVSPPKQQVSPGPRVAVGTKRPIVEDASGGHGGRRDDCASPAKRGRTDVDNITFHPISEHRHWCPWVCPVTVPGPSEADEGASQDK
VPGWRAALNTMLPMIVGEGTVSTLEPTDVWKKVHRSIMSWQVSGSG

**>XP_002128832.1[*Ciona_intestinalis*]**
MELTSENSSENIPPAIRPCKTVVDQVKNTLSFLLTSAVNSIKKADSEKNEIPVEKSSSVDNPQILARKNKNLGDSPKVVSRSEKLRSKEAFQKR
VKSFTLQNWCGKPLLLNPMLYAQYGWRCSGEDMVCCSSCGAVQCVQLPWPNATNYEEQCLKTRSKIVSGHMKVCSWSSSYCNDSFIIPFHPRYS
NTQQQTCMLEEFQARAKKLLHLKNDLPLITDDVKEEMELSEKILVQLCAAADFNDNDDIENLMVVSSIILALSGWDTKSDGDSSNPGIFCNESS
RLVGLWNFYSVGYLPNETDAPVAKKQKKDEDFTAIEKSYFHPLKQHHVWSPWVVTIKPMSADELSDERTSATYTDDDVLPGWKMLKNLICSDGL
TTSKPTMKTPPRAAVKQARRILSEWSSPM

**>CAG5088580.1[*Oikopleura_dioica*]**
MSAADIIKETETALKNFLNKLPDPKNEEVRIFDPSAIILLRTFPRYQQRLGTFDVTKWSGKPACFSPTFCAIYGWSCKEKDVLKCPDCGEVLLA
ELPPRNNDYFPKKVDHLLTMLQSGHADYCEFANSHEPFEFLKKNDSPYLFKKRLQTFPGTLDFLPKVTSGISEEDLDFLSRYFSYCVKNKKLQR
EILNLAVNGWSFHHEDETDYLLCEDDVRQIPER

**>XP_041459241.1[*Lytechinus_variegatus*]**
MAASNFETSKPRRIKALLSSFLKGISREEKKESETKQVIFESLDTEGFADGFVVVDEVSSEQTSTVQPLNQELFFNRVETFSISSWFAKPDEVC
PLRCAQYGWENIDVDSLKCVSCKEVLYGGLPPKWETDLYENACKKLVDSLKTGHSKICPWQSNPSPASFLEVNLVSSQNAVNDFLHRVASIRCF
GTSVPAVDLSCLQQVDENEDALSRIVSGVLGEEPTCDYKERIESVCFMAACGWSRSSPEGSQSPTMSCQYCRRNVGLWNFTPYEQNAKTVTEDD
SEPTAKRLKVDKGLFNPIEEHRSWCPWIKPTSSTQKVKSLPQDKNQDDERPVRPAWHELLVLLHQRSSPDKQGLLTNKQVTPPSQAWKAVRRIT
NFWQSRNAVNKT

**>Own_assembly[*Saccoglossus_kowalevskii*]**
MVSDKMAEKSSVLVTPKRVHDLLSSFIHKEETGVDERSDEPNQENIAQSSRQFLPRNREAFFARLETFSAFTWFAKPIELSPLKCAQYGWENTD
NDIVKCVSCKEIVCASLPKTWDPDLYAKRCEELRAALVKSHSNICPWRDSPSPDIFLSIPLLNQSEVQGDVLSRCTSLEKLGRKLPVIETCDIE
SQITAEALSGVVTRQINLLEDIRNDDDSSAINTVCVLSLCGWSTSSCETSQYPTVSCQYCRRQAGLWNFTPVVEPEPDKMTSEEQPTVDSSVES
SDADSAQEPSMKKRKISESKKSAFNPITEHRAWCPWVISYVMQQSDDHKSSPVPTYNNIPGWQAVFTLLAPKTSPLKENIQRLTDKDQTPPNQV
WKAVRKILSFW

#### Representative sequences for Figure 3B2:

**>XP_046992160.1[*Schistocerca_americana*]**
MVLGKVSMMADDGYEKVRKIKSLLAECMVPPRESQFLNLDRYIEENSEVSFIPDPPSWCNISFRGFQERVLTYRPQHWGADPNAVLWFSKYGWR
CTEKFVIKCDICSSTVECSSLIQKPDLWGDIAHKAHDNLCYWSFCPCPDRFIQIITHPKILCKNICHSWKKIIMYESELPKIQQVVLDKMGLSR
DILEHLFKIIETNCSEEGISALVLVICGWLKSSEDNTLECAYCERHLALRHFFSIADSSKINDSVPVQDDVRPGDEATKMDISETNAGKSLSSE
TNRDNSVADSGCKIESVHTLHPMKRIRRGQPAVHTGIARHRARALSCVTKQKFPSFCNMLHLRYGVKLPKHRGSRKNKKYLGRKYSGSDIIVED
EITTNGNESASEKPTQELYYIDKTDGEDSGPPNGKNGIKRKFPDNEDTENIEICQIDKQQKVAADVSKIEDNNSNMHKVDIDTVSNVKCSTEDS
VKPESDSEMITRNLISENVEISRNMTESNLPVSEQLVSPHELEGKEASSSSNCVQNTLRNKSDDVPPAEEEYRGDSNFEKKQNEIAKIAAISAS
EEIGPNVGKRQLICDDDQQTEGPNKRLNISSENSPLRKKNLNPVVEHRFWCIWRIKTVDRLLGSPKEGWRQILDLLKFGVSPQNSEAEEEAGDM
FEEVKKIHYMMSGW

**>CAD7459619.1[*Timema_tahoe*]**
MEFEDRIKRYKKLPFETLNGLESSDVSDDIKTAVNQDFDKFVERVATFGVLKWGAYPQLVLQLAQFGWQSEMKDYMIYCTSCSTRLSFSAYLKE
PDDWTDKIKSSHYKFCRWISLGHAYPEEFVKVPSDLKLIRDRVITRTKALLELGTSLPKMIALEHGALQLIVKNILKVPLTEESLSALILSLCG
WGPTTSHNVLSCDFCKRQVGLWNFITIQDEPTLADDYLCILNSPGSCTDEPLSILESSPIQDVDMQLTDPQTTEGSELHEDATEKEESSFIDDQ
YDYGHYDENQDSCDDENEGSVEQYTIENNDEENLDSDQRESAMKGYQDPNLMIDNINLVPDNFSEQDCEDFPGIEPEDGTTSVDTKVSTKPNLL
EGENGGVEILELTSGSEESDGSLEDEEDEEDYPSDDEEAEDEEMYMEEGNIRFMTDEKGLIQKYRKDEESFEDEEEDIEEHCDEKDDPEGEEEE
EATEEEDDPEGEEEEEATEEEDDKEEGTEEEDDDDDDDDDMSDGERVEQVGGNGSPSNGVVDEQSNPQTFEMYDSERDGQNSSVIRAKGENETT
TNVVGSTMLVYSENKVKGENVIEQILVETQNVQAVVNKPIKLMIEEMEIAVTPNTEVAMNHTSHVDKAENKREVKEINNKSDKACILDETMVSS
KDIESESEMLVKSTKETFHYLTNDLLNNDKKTKVFDGPKTLTQQIRREDVEAKPGTAFEKEDSSETTKNLINVKHTNVRPGTSVESQEDLDNQT
RSGLELESGKVEQEPGPPIEIETKTCENVPAEQTISNESDGPECMEAESRTEMISKNSTSELEPCLVSERANSEPMEIDKVETACTGKRLINVE
EEMEVKTENQNVGDDSRQDVKGDINLKDSQSQAWEDEINLKEGNGQEMKEDINLKDSQSQDVGGSKDIQEQDVSTLPSDGSDCVTKEKEHSVSE
ESVEKLLVVSGEVDCQHVLDETEDKTETTYSTCNGPVVDMDITGLGTSVDIGANQDAEDRVHAAGQSRKRRRVSTPLPEIDAKRWRMESVKLDF
DPVQEHRYWCIWGLPVDRGQEDSLIGWQRVLADLVRYTLQKSTIVSEEETTAEEKNIFEELVKYRDVLSTLVLKRGCSLQWEGGVNKHCLPWLI
TPWWSQASQAEGLFIESERVEKGEIAGTLTIVLELPLLYKEPFESEVFHLEVLMSLRTATSTQPYSRSALEFPCDQDRVLVTNLLVEHS

**>Own_assembly[*Catajapyx_aquilonaris*]**
MNSHSASTTILCLFRSELVRPRCIFKLSKRCAPTAKPWSHHDFVGRLKTFSPINWPYKRVSPIECAMRGWVIYGMNRITCITCKGRIYGGIMDT
KYTDLYEQWVQKLRSNLQERHELCCPWRVNPTRLEFEYIFDPGKVPQLGFIRRLKKFSQVNMLPELSEEFLESISEDTETINSISSRLESVPEE
RVALILALCGWEVKSADVLCCGICLRDVFIRTFGQTVSRASTPSQSGADDSYQNNLTDSDLATDISTSVLEPNTSVNEESVGDVGERNGDCDIG
HLVAQLELNGKVAGDTEKISEEIVVNSDTAEMEVQSESNEDNLGAKTVDTIEMKDSLACSEKDSPSAGVVREDLKIDPKSEERDEAADVMDTDM
TEENTCKQADSDSNATDLEQKLRPEVGEGTKIKSDAMDAVSNEEVNSPSNKKGHQGEDKDISVEPDKNGISLDKNDENVTAEGDSTSDIPSEEV

KKNEIPAEGDEKNAIPAEDDDKKGISIDEGLPAEEKSLPNSNCKNEAVQLEVDANVNTTTSELGEEKCSETGISEAEPSEEAPKNEKNDDTIFV
APKIECHESDVHSEADTVENTFLADDMEAETSSVVAPSTEPNSPNTSFLIPPDPNAASKKRRRSTSIIQMKKFRPSSFHPLNSHRFWCPWAAPN
DIVKHMSNTKLPGWKLYMSNINNNLIRSPDTIRLCHSSTYVSMEHINVRVVANINFFLVDSGRRTAKDTPRPWKVPLAVYRNNSPTKAYFYVYK
IRNFF

**>CAG7825680.1[*Allacma_fusca*]**

MLLKRKRSLDPGSERGIKLVRESLEAGVSDGKDLNRRRSLESASKADIETFINGIRKPSVDSGLPVSFDEFEASKFVAKENLVRDSQQTALSIE
DFKARLATFDMRRWSRKPDCIASFHCARFGWKLCEKGDALICVTCKQLIPVSGDGNAFAASSSGLKCGTQLRQAIEGNCHANSCPWKKYFSPVT
ILCLPQGFDMFYFLREGVASFQRTFTLPTLHSHYGHIMDNLDLTWMECIAGLKTLGVNIKSEDSENEKILHQQTRLICFLVLSGWSMKATNVED
GIGCYLCLREIPLWLCKSTADANSVCSLSTPVGSLKLGGRKYCSELDPINNHWWWCPWRRFLEFSDAATIVSSNVRKDLKDADFFEQRFEDSLC
LKPHADKEDYLRLKVLIEQECSQAASISEVKVISPQVVSTVGSIMDAFLSPSRCSTLTSLADESLLLPCDFSMKSEGTEAKPVQFKSNSNSMQE

**>CAG7712993.1[*Allacma_fusca*]**

MISESEMDDDEQEGPSSSKIRRVEKEDHLIIERRVDVYRKSNLGKSLAGALQELLNLKKIEAGLAEVVLQQFDRSVGRNLPHTRNHLVFSAERV
NRYRLIADNWKVTLQNVTFSTPYSELVIYYWRTQDLKPDMLPTSVVGNNRTFNSKSSFILQIKYYLKCAGFNLLIMAEQEVDILEKVRIALDSG
IVNLAKPAPDAGGDPDQNGASCGTTTKQPPGDYIDIDEYCKMLQDFRDDGIGPMDSRNLIGKQQTKLDSQQTSHSVADLHDRVSSFDILKWSQK
PDCISSFQCARFGWKLTEWGFLACVTCKNILPVPSQIPKINGDWANRLRQGIESSGHLPTCPWVRMPSAEDILAVPKGSDLFKFLCESVGSFKM
MKTVPTLENHYKMVSVDLSLKWLCELTSATLPDVAEEKEKGVTPEMVRNMALFLVLNGWSMSEVEDIMTCQLCLREVPLWLCQPTPPQHEILDL
LSPPASPVLNKSFSYKEFNPVSNHWSWCPWRKRLLTNVDESLMSMTPTKDSQQEFIRIKNQPSVDDYEKLLEDFKKECASITLNYDKETSNVRL
IDSVTSLFDNFIGSSVQPLSDGIEDSNDTNGGVDGNGLSKNHDNSEKECNSEKATEEDPADVVTPPKKRIRRELFEAGADI

**>XP_046451818.1[*Daphnia_pulex*]**

MATLNRKRKLDQTLIQKLYHDVPSKDPTRESFLKRLKTYDVFNWSGKPVDPPLCALHGWEIAEKDVLKCVMCHQFMSVTLPSPTKDAPYKHACS
KLKSRLASAHSKFCLYSTNQVPDSVLEIEHVSNMELQDTIQQQLLAFKNIEALCKVSELQDEIKEIFDWFLETTAIPDVHLSSFTFVLTGWKFL
QDDMLLKCDYCNRKWSIEPYLSQKNLTEKNDSVRTTVDPVAQHQRWCAWRAPTRGWKSRLLQLQQLKESRCREKRSRLSSDSCDSLTDRMRTVR
KLLNGTL

**>tr|T1J7V6|T1J7V6_STRMM[*Strigamia_maritima*]**

MASVSHSEISVHQLLQAFLVDSSDKASKEDFLKRLATFSISLRCARYGWYNKDADFLKCVSCDSVLCGKLPKRFQFELHKKSLNKLRNQLADAH
HKHCPWRNNPSPEPYYNISRWSKEEVTNEFLSNLNRLKSLEGVLPVLQDSCVNAIEDSLETLQKISDDPKLTLPVAQLAIRGWTCDLNSDANCS
TLYCAYCVRRVGLWNYKSNEEDSCDNPNKKQKFNGDTVEKKAFDPILEHQIWCPWVSTSQENDKLGWKVFLHVLTNKSSKLDHDSSELSSKQHL
GQIRKWLRGLMADSNCGKKFAGNISPTLRRPGCRNPEFLVTFTLRPRSPPKALVVELPGPHSLWGRPTLIERPPIVPSSLTKEDELEKLLEEKS
PVPVIGPTIYTLPKGRTNPKGWWSLYEPQILNLDFVTSNSEYSRYPPNPYTMPISYSVKDRSFTNVSI

**>XP_029831140.3[*Ixodes_scapularis*]**

MTAFVDAECSQLMDSLTTFSLPGISAKDYADFKSRIETFFDDYGTLSRWPCKPPELSPPQCARFGWTCANESLLVCAACKEYLDCEVSSSLGRK
LHKECLSRLVSLLEGAHKPCCPWKTAPCPKSYTVMQPVLRKDALSQLRERLETLRPILSTLPVLNTDKILSLLSPEDILRIGKLVDKDRGTETQ
RADLLLAITGWQAGAGSGKMKLVTCEYCSRKVATFFYKSAPISESRDEVTTSQDKGACSPGHGTKRKREEDELDPVHEHRPWCIWVLNDDSGKP
GWLVFSECLLRNVDSSHDDSRLSTSSVDAFKHDVEKIISSWREVVKHPTIQKTTTS

#### Representative sequences for Figure 3C:

**>XP_004235730.1[*Solanum_lycopersicum*]**

MAEESQKRFQDAMDKIFRTPPKSKLNSSASGVQLSRDKERLDMSSIGKAVSKYNLLATKGSGEAPPCRPWDRDDLFTRMSTFKSMTWFAKPQAI
SAVNCARRGWINVDMDTIACEACGSRMLFTTPPSWAQQQVDKAALVFSLKLDSGHKLLCPWIDNVCDEKLADFPPTATVMLVDQYKIRHSVLSQ
LAALPVISPKAIDFLRNPQLEQFLRESLTVEHDESMHTPQEETRNAPTSVSSLTYYQVQKLISLCGWELRRLPYMVDPKDQLNQSSKDANLSEK
SILSRKSEIITVYGSCTDKTSESKTDDDNRASEEAIINPNSVVLDCKLCGACIGLWDFSMVSRPLEFLRVSGYTQVNNDHINHTHGDKNHFSGN
SGRDKSRECTGQVTTSANTMLDRRPPNFNLTIAGGPPPVTHDYRAKISLPIIGRNLRAWFIAESELKDDLVTKSSSGVSKNPEFLAGENTEEGS
SLSTSEVSTEAQLENNQAAAQVSGNTTEMADNTESMNKVDPAVTDPCKDKVGNDFGSSSRGKELPILSLDKALEFDPFKLHRYFCPWIASNGVS
PSGWEQTLSALERHEESSSPLSNHAPSSLIKVDDPVASVQKLFTSPQAKRRKLVRPS

**>XP_010323636.1[*Solanum_lycopersicum*]**

MKEEAISSSHDPQLPPKSSSPPPIPTPAASSVGASSPAVPTNAGGTDWFAQAQGSKAASLSRIGSQPMWTSVSNSAGGSALGSSQPSCRPWERG
DLLRRLSTFQPTNWFGKPKASSSLACARRGWVNVDADTIECEACGANLRFVSSATWTSGEADIAGEEFAKKLDEGHKATCPWRGNSCAESLVQF
PPTPPSALIGGYKDRCDGLLQFPSLPIVAASAIEHIKVSRSPEIDRLLAQSQAFGGMEPIFRLEIMSGTETNTEDVFLVYSRANKLISLCGWEP
RWLPNVQDCEEHSAQSARSGYSIGPTKYHTSLQDFGHGENVLPSSKKKVHSKNEAVGPRSKGESRSPLLDCSLCGATVRIWDFLTVVRPACFAP
NSNDIPETSKKMALTRGASAASGISGWVAADGVEKEQTEDLDEAATNDVGRSLSNIGVDLNLTMAGGLSSSQVNMDAKPEQFEDGHKRRYPVTG
QPSSSEVGGQAASYESRGPSSRKRNLEEGGSTVDRPQLPLQPADSVEGTVIDRDGDEVNDGSQYSAGPSKRPCQSDAFGTHHTSYGKDSSGAGP
SLSLGFEIGTSAPRDDTFGRRHEQLTGVPSTRDSTHVSSVIAMDTVHGTDDSMESVENLPGDFDDVHFPSTSMLRSADPVETSELNYSNQAQQS
TCPAVVRSAGEMGVSSTNDEEVVNADTATANVRDGPSFGISGGSIGMGASHEAEIHGTDASVHRADSVAGEVAVAEITENQGQTGEFAPDPGL
MGDYVPEEVDRGDPNGDSQDLTSRSVGRADSGSKVVGSAKAESIESGEKNCHVQPMLPNSPHPSLSCNAVVCSAHEASKEEVTQNNAPATDDCG
FVESDYMLANGTGPPIGESNYEEAVEFDPIKHHNFFCPWVNGNVAAAGCSNSGSSSSNSGAIALCGWQLTLDALDSFQSLGHIPVQTVESESAA
SLYKDDHRAPGRKLLARHSFSKHHGHN

**>XP_008374705.2[*Malus_domestica*]**

MSKDSEKKFHLIMDKLFFAPKSAPSSASSSSGVQTSRGKKRANPSSALALVEPKSRGDRMEVSRHFSAPAVAAHAPLCRPWDRGDLMRRVATFK
SMTWFAKPKVVSALNCARRGWINVDADILACESCGARLFFSTPSSWNQQQVEKAALVFSLKLDNGHKILCPWIDNACVETLAEFPPTPPPVLVD
KFRERCYALLELSVLPVISSSAIEYMKSPQLEQFLGQSSMFYGNGSGDISRTEHSDNEGNADSAKLYYQAQKLISLCGWEPRLLPYVVDSGNRL
NHSATNRQNPSISVHSASNDEHKNASACTNIQAEHDSVVLDCKLCGASVGLWAFSTVPRPVECFRLVGYAEVNSESHSGTHDSNAESHCDSRID
VLNAGVDGATLSKDRFANLKLTIAGGPPPTNQNFKAIISIPVIGRNLRARISYDSELRDCLSVGQEGMQSDTQMEKEENHYQENAENGGLENSE
VCGPGTPDANATHLNGEMDKSDPLVMVSSKGDLLHSGTIVEHSEEHESTSVPSSFEANADLNSSRTDPEPTSNQEASEDTVQIPANGELVACSS
GKDLKHVVPGSMMEFDPIRQHRYFCPWIASTGNGAPGWKQTLSALQRQEGGSPSSASIIKVDDPITSIRNLFTSPSPKRTKPTVLTTRTSEQ

**>XP_028947922.1[*Malus_domestica*]**

MSKDSEKKFHSIMDKLFFAPKSAPSSDSSSSGVQTSRGKKRANPSSALALVEPKSRGDRMGVSRHFSAPAVAAHAPLCRPWDRGDLMRRVATFK
SMTWFAKPKVVSALNCARRGWINVDADILVCESCGARLLFSTPSSWNQQQVEKAALVFSLRLDNGHKILCPWIDNACDETLAEFPPMPPPVLVD
KFRERCYALLELSVLPVISSSAIEYMKSPQLEQFLGQSSMFYGNGSGDISRTEHSVNEDSADSAKLYYQAQKLISLCGWEPRLLPYVVDSENRQ
NYSATNRQNLGINVHSASNDELMKSHNTSIQSEHNSVVLDCKLCGASVGLWAFSTVPRPVECFRLVGFAEVNSESRSGTHDSNTESHCDSRIDI
LNAGVDGATLSKDRFSNLNLTIAGGPPPTNQNFKAKILLPVIGRNLRARISYDTELRDCLSVGQEGMQSDSQMEKEEDHYRENAGHGGLENSEV
SGPGTPDADITHLNGEIDKSDSLVMVSSKGDLLHSGTIVEHSEEHESPSVPSSFEANADLSSSRNDPQPTSNQEASEGIVQIPANNELVACSSG
KDLKHVVPDGRMEFDPIRQHRYFCPWIASTVNGAPGWKQTLSALQRQEGGSPSSTSIIKVDDPITSIRNLFMSPSPKRMKPTVLTTRSSDQ

**>XP_008371629.2[*Malus_domestica*]**

MREEVISSGGNIDPAPAASSAGASSPTVPANVGSVDGSIHGQGSKGASISCVGSQPPVTSLSTSAGGGGGGDSSVFGSSRLSCRPWERGDLLRR
LATFKPSNWFSKPKVISSLACARRGWVNVDVDKIACESCGASLGFALLPSWTPDEVQNAGDAFVKQLDSGHKAACPWRGNSCPESLVQFPPTPQ

SALIGGYKDRCDGLLQFHSLPNVAASAIEQMLVSRGPQVDRFLMAGEVDFKPESIPEQESSRDGSICLYSRAQKLISLCGWEPRWLLNAQDCEE
HSAQSARNGYSLGPTYAQVHLSQEPGPSKKAVSASARKDAGKSKMLVKESRRDPRSPLLDCSLCGATVRILDFLTIPRPVRFTPNNIDIPDTSK
KLGLIRGASAASGISGWVATDDAEKEQTEDRDEVATTTEGSLFPKTDVDLNLTMGGGFTFNRFGRPEMSENIQDADMGRDLMIGQPAGSEVGDR
AASYESRGPSSRKRSLEKGGSSVDRPHLRTQHADSFEGTVIDRDGDEVTDGGQYSAGPSKRARDSDMFDTYCSSGAGPSHSMGLDIYADANRVA
SFPQGSDQFGIHSNMDSARASSVIAMDTIGHGTDDDSMESVENYPGDVDDVHFPTSSTYGNLDMNDTSELNYSNQAQQSVGFQPVADVVIGEIG
VSSTNDGEEIFNTETVTAQARDGISFGISVGSVGMCASHEAEIHGADVSVHRADSVVGDVEPRTEDAENQGQTGESAPDPGLMDEIVPDEINRE
DPHGDSQEMISRSIGRADSGSKIDGSTKAESVESGEKISQGFKFENSARPSLSCNANVFSNYRTTKEVKSAGKSSFTNNCVYQESEYAVANGLG
PPKGESNYEEPMEFDPIGHHNQFCPWVNGNVAAAGSSSCGHGSSVAAVALCGWQLTLDALDALRSLGQDAIQTLQSESAASLYKDDHQTPSQKL
LQNHSISRSQGQY
**>XP_003609078.2[*Medicago_truncatula*]**
MSQDSEKRFRSIMDKLFHSSKSSSNNPDKSSSGVQLSSSRGKKRGFQSIVDRRGDEQYLSATAVSESQGHLCRPWDRADFMRRLATFKSISWFA
KPKKVSAVNCARRGWINVDVDTIACEECGARLLFSTPASWNHHQVEKAALVFSLKLDNGHKLLCPWIDNACSETLARFPPTSPPVLVDNFRERC
SALLELSTLPRIASSALDHMQSPYMDDFLGQSLMQECGNGSAENFGIEDVSSQEELKLYYQAQRLISLCGWELRYLPYAVDCRDVSDQSHKNST
IVYSPRVVSDARNNNLTVYSADNNESSKMDENSKHSIGEQMDPNSAVLDCSLCGATVGLWAFCTVPRPVESIRLVGYAEVNVDNDLESRQGVNN
ALSDIANSSKDTSLGLNMTIAGGPPPTKQNFKAIISLPIIGQNLRARLSYDYDIRDHFFVDRGGSQSDSQEIKIQEKTDNTVDASIGQLVPVSS
EIREISNCETGSQQASICDSVLDNDLEGTSSAGQPSGFKEKMPVQAETGGLKNSSAEDPSSSQTDMAEDEALSHKTKEGSHVETSGVKERAENP
INREDVHNSIGKFKNPSLPGKAMEFDPIRQHRHFCPWIASEDGVEPGWKQTLSALYRPKEHLRHSSNTSPSSMSIMKVDDPVGSIRKLFMSPPT
SRRKLTHISSQNAEHR
**>XP_024629233.1[*Medicago_truncatula*]**
MREEVISSGGTVDPTTAASSSAGASSPTVPMNVGSIDGSSHGQGSKAASLSCVGSQPPWTSISTSVGGSAFGSSRSSCRPWERGDLLKRLATFA
PLNWSGKPQVIDSLACAQKGWMNIGEDKIACESCGACLSFTSLLSWTVAEAQDASESFARQLDSGHKANCPWKGNSCPESLVQFPPTSQSALIG
GYKDRCDGLLQFHYLPVVAISAIELMRVSRGPQIERFLSQSQNFMSGTDFKPENISELESSQDEAYCSFTRAQKLISLCGWEPRWLLNVQDCEE
HSAQSERNGYSFGPSKTQLRLAQDPGPKAVSASTKMDPRKGKEPFKESSLEYRSPMLDCSLCGATVRILDFLTVPRPSRFAPNNIDNPDTSKKI
GLTRGGSAASGINGWIAADDAEKDQTEDRDEVATTNEGKSLANTDLDLNLTMAGGFRCTPFGRTATSENMHDVDMGRDLMIGQPSGSEIGGRAA
SYESRGPSSRKRNLEKGGSSDDRLVLRSQQQADSVEGTVIDRDGDEVTDGGQYSAGPSKRVRDSDIFDTYCSPLQRDSSGAGPSNSLGFEGYVT
GNRVSSFHQGSDGLIGIQSARDSARASSVIAMDTICHSVNDDSMESVENYPGDLEEVHLPSSSTYGNVDMNETSELNNSNAAQQSTCLQTAPEV
VRGEVGVSSTNYGEENFNAETVTAQARDGFSLGISGGSVGMCASHEAEIHGADVSVHRTNSVVGDMEHRVEDAENQGQTGESVPDPGLMDEIIP
DDINREYPVGDSQEMMSHSAGRADSGSKIGCSTKAESVESGEKISQNCKLPPANNSHPSQSCNANIYSDCGTTKEEIMKDGKSSFTNNCALVGS
DFATANRIGPPKGDNNYEEAVEFDPIVYHNQYCPWVNGNVAAAGCPSSFPGTGSDAIALCGWQLTLDALQSLGNAIPTVQSESAASLYKQNDPQ
APRKKLLHNHSMSRSHGQL
**>XP_024632064.2[*Medicago_truncatula*]**
MKEDDVVTSSKKKNHKPHSAASSAGASSPPYDTTGEASRRDKSSADSYMLIASALHGASNPSCRPWERCDLLRRLSTFKIAGKLPKVGGPLACA
KRGWVNVDVSKIECELCGVQLDYALPSASSAEEADASSEELSKQLDRGHKINCPWRGNSCPESLVQFPPTSHSALIGGFKDRCDGLLQFYSLPI
VSSSAVEQMRVTHGPQIDRFIAQLQIQTAGELGYRAETSLTGEQAPHSYSHAQKLISLCGWEPRWLPNVLDCGEQSAESAKNGYNSDPAKGSAP
GPAPSKEFSNSSRKDTGDNDVLGSEFNCESRSPLLDCSLCGATVRIWDFLTAPRPVHLTPCGTDTPQTSKKIASMRGISAASGINEWAAADGVE
KERTGDRDEATTSGKRKLVSNKGLDLNLKMASGPRRSLINVTSTLDHVQYAGEGSNLRNRGPSGSDVGGPAASYESQGPNVRKRRLDDGATRAD
RPPLSMQQADSADRTVVNHDNNEISGGQQYSAGPSKRARDANHLETLQFSLRNTSGAVPSYSANIQSEAEENTVNQLNAEKDHVTSMPFTREST
HASSVIAMNGRYHSSDDESMESVENSPADFNEVNFPSVDLNETSELNSSYQAQQSACNQPPLERTGGEAGLSSSNVCGEVLNTEILTAQARDGP
SFGISGGSVGMGASHEAEIHGTDVSVHRVDSLGDAEQIAEVIENHGHVSEFTPYHGHNGDFVPEEMSREDPQGDSQAVVSQSTARVDSGSKTIA
STKVESVESGEKTSCSMETPGLENSAHPSLSCNAVVCSAYEVSKEEVAQTGKPSYIDDGAHPSLSCNAVVCSAYEVSKEEVAQTGKPSYIDDGA
HPSLSCNAVVCSAYEVSKEEVTQTGKPSYIDDGAHPSLSCNAVVCSAYEVSKEEVTQTGKESYIDVSTYHESGNLDADVVGTPYRDNSSGRVEF
DPIKLHNDYCPWVNGVVAAAGSDSPCSTSDVGPAARCGWQLTLEALDSFQLLGHLPVQTLESESAASMCKGDRFTSSQKLLARNSFVRHQGKN
**>XP_021625743.1[*Manihot_esculenta*]**
MADDPEKRFHSIMDKLFHAPKSLSNPSSSSGVELSRGKKRPNPESALALVEPRTRGDVVGSSQRSLAPADAPLCRPWDRGDLMRRMATFKSMTW
FAKPKVVSAVNCARRGWINLDMDIIGCEACGARLLFSTPSSWTQQQVEKAAMVFSLKLDNGHKLLCPWIDNACDERLAEFPPTPPPVLVDKFRE
RSSALLQLLGLPMISSSALEYMKSSQLEEFLRQAPTLDCGNGSIKISQVEYPGNESEAYSANLYYQAQKLISLCGWEPRLLPYVVDCKAKPKKR
IKDANTLNSSHIFTNGQNTSIGFYSATTNENAEATEDFNAPGGLQADPHSIVLDCKLCGASVGLWTFSTVPRPVELFRLVGYTEVNSRKNYGQD
SENESQVNDRQVINSSSNGVLSSIDRPSTLNFTIAGGPPPTKQNFKATISLPVIGRNLRARFSHDSGFRDHTFNDLEPQSRPDKYLCMEESSIT
ENFGEQVSLPESVGMLKSKTTDQGQCSSASGDQSSCLNIENGKKGSDLRKDSDSNRECTTESTADAAQGFDQSNRLPENALNVGSLDSPAGSLG
SSQIIVSSMSGPGATVTAGNGNSTRDSLALVTSEGGNQQQVPGADVLCGKDVNLKIDLTKGKELKEISDEGMEFDPIRQHRHFCPWIVSTESWA
AGWKQTLSALFRLKDLSSPSTKSPSSTSTVKVDDPITSVRKLFMSPSAKKMKPTRGSS
**>XP_021598693.1[*Manihot_esculenta*]**
MREEVISSGGTMDPTPAARYQITRLPSFLPRSAGASSPAVPANTHASKAASLSGVGSQLPWTSLSTSAGGSVLGSSRPSCRPWERGDLLRRLAT
FKPSNWFGKPKIASSLACAQRGWMNIEIDKIVCESCGACLSFVLLPSWTPTEVESAGEVFARQLDDGHKTSCPWKGNSCPESLVQFPPTPQSAL
IGGYKDRCDGLMQFQFLPIVAASAVEQMRVSWGPVVDRFLSYSQNFTFGEGDFKPEGIQELENSRDGASYLYSRAQKLISLCGWEPRWLLNVQD
CEEHSAQSARNGCSFGPAQAQVHLSHDPGPSKRAHSASATKNTGKNRLVAESRCDSRSPLLDCSLCGATVRILDFLTVPRPACFAPNNIDIPDA
SKKMALTRGVSAASGISGWVAVDDTEKEPTEDRDEVATTDKGKLLQNTEVDLNLTMAGSLPFYLPDKAAIPESVRHLEMGRDLIIGQPSGSEVG
DRAASYESRGPTRKRSLEIGGSSDNRPHLMMQPVDSVEGTVIDRDGDEVTDGGQFSAGPSKRARDSDFDTHCSPCQRDSCGAGPSHSVGMEIYA
DGNMVNLFRQGSDQVVGIPSARDSTRASSVIAMDTVCHSTDDSMESVENYPGDIDDVHFPSSSTHGNLDMNETSELNYSNQAQQSISVKYAAEV
AHGEMGVSSTNDGEEIFNAETVTVQARDGPSFGISGGSVGMCDSHEAEIRGVDVSVHRTDSVVGDVEPRVEDVENQGQTGESAPGPGLMDEVVP
DEINREDPHGGDSQEMFSRSVERADSGSKIDGSAKAESVESGEKASQSCKLALGNNDGPSLSCNANMYSGYQTTKKGVGKAGKSSSTNNGIGPP
KGESNYEEAIEFDPIIHHNQFCPWVNGNVAAAGCSSRSSGNNADADALCGWQLTLEALDALQSLGHIPIQTVQSESAASLYKDDHQTPGQLLRR
HSMNRSHGQH
**>XP_043807289.1[*Manihot_esculenta*]**
MREEVISSGGTMDPTPAASSAGASSPAVPGNICGMERSSHAHTSKAASVSGVGSQLPRASLSTSAGGSVLGSSRPSCRPWERGDLLRRLATFKP
SNWFGKPKMANSLACAQRGWMNVDVDKIVCESCGACLSFVLLASWTPAEVESSGEAFAKQLDDGHKASCPWRGNSCPESLVQFPPTPQSALIGG
YKDRCDGLLQFLFLPVVAASAVEQMRVSRGPVVDRFLSQSHNFTSGEGDFKSEGMPEFETSRDGASCLYSRAQKLISLCGWEPRWLLNVQDCEE
HSAQSARNGCSFGPAQAQVHLSHDPGPGKKAHSASAKKDTEKNKLLAESRCDSRSPLLDCSLCGATVRILDFLTVPRPARFAPNNIDIPDASKK
MVLTRGVSAASGISGWVAADDTDKEHTEDRDEVATTDKGKLLQNTEVDLNLTMAGALPFTQADRLAITDNVHDVEMGRDLMIGQPSGSEVGDRA
ASYESRGPSSRKRSLEIGGSSDDRPNLTQPADSVEGTVIDRDGDEVTDSRQFSAGPSKRTRDSDFFDTHCSPYKRDSCGAGPSHSVGMEMYAEG
NRVNLFHQGSDQVVGITSVRDSTRASSVIAMDTVCHSADDDSMESVENYPGDIDVHFPSSSTYGNLDMNETSELNYSNQAQQSICFRHTAEVAP
GEMGVSSTNDGEEIFNAETATAHARDGPSFGISGGSVGMCASHEAEIHGADVSVHRTESVVGDVEPRIEDVENQGQTGESAPDPGLMDEVVPDE
INREDPHGDSQEMLSRSMERADSGSKVDGSTKAESVESGEKEASQSFKLALDSNAHPSLSCNANMYSGYQTNKKGVSKAGKSSSTNNFPCLESD
YTIANGIGPPKGESNYEEAIEFDPIIHHNQFCPWVNGNVAAAGCSSHDSGNNADADALCGWQLTLDALDALRSLGNVPIQTVQSESAASLYKDD
HQTPGQKLLRRHSMSRSHGQY
**>NP_175325.2[*Arabidopsis_thaliana*]**

MAQDSEKRFHQIMDKLFTPSKSQLPSSSTSSSVEQQSRGKKRQNPSSALALVEPKIVLATIDRSSALKVPAGTSPSGLCRPWDRGDLMRRLATF
KSMTWFAKPQVISAVNCARRGWVNDDADSIACESCGAHLYFSAPSSWSKQQVEKAASVFSLKLESGHKLLCPWIENSCEETLSEFPLMAPQDLV
DRHEERSEALLQLLALPVISPSAIEYMRSSDLEEFLKRPIAPACSDTAAESSQTESLTNHVGASPAQLFYQAQKLISLCGWEPRALPYIVDCKD
KLSETARGTETIDLLPETATRELLSISESTPIPNGISGNNENPTLPDTLNSDPSSVVLDCKLCGACVGLWVFSTVPRPLELCRVTGDTEINIEK
HPKGGTLQHQPSSLKFTIAGGPPATKQNFKATISLPIIGRNLRSRFASYSRDHDHGDVSSIQDQQSRTAENNGDVTQNSNQVMNDIGEKADGGR
NSTDVESDIALQNKDKQMMVVRSNLPENNKPRDSTAEKSATSNKQMEFDPIKQHRHFCPWIWSTGRRGPGWRQTLSALQRHKGSCQTPPSSSSL
FKVDDPLTSVRNLFKSPSPKKRKLNGGSSS
**>NP_173164.1[*Arabidopsis_thaliana*]**
MKEEDVSSQNVNPRSNRNSVASASASASATPVDRFRRRARSPSPPQTAAASSAGASSPAVLVNAGSVDWTGHGLALSVRSCRTWDRGDLLRRLA
TFKPSNWLGKPKTASSLACAQKGWVSVDLDKLQCEYCGSILQYSPPQDSLNPPEADTTGEKFSKQLDDAHESSCPWVGKSCSESLVQFPPTPPS
ALIGGYKDRCDGLLQFYSLPIVSPSAIDQMRASRPQIDRLLAHANDDLSFRMDNISAAETYKEEAFSNYSRAQKLISLCGWEPRWLPNIQDCE
EHSAQSARNGCPSGPARNQSRLQDPGPSRKQFSASSRKASGNYEVLGPEYKSESRSLPLLDCSLCGVTVRICDFMTTSRPVPFAAINANLPETSK
KMGVTRGTSATSGINGWFANEGMGQQQNEDVDEAETSVKRRLVSNVGLSFYQNAAGASSSAQLNMSVTRDNYQFSDRGKEVLWRQPSGSEVGDR
AASYESRGPSTRKRSLDDGGSTVDRPYLRIQRADSVEGTVVDRDGDEVNDDSAGPSKRTRGSDAHEAYPFLYGRDLSVGGPSHSLDAENEREVN
RSDPFSEGNEQVMAFPGARDSTRASSVIAMDTICHSANDDSMESVENHPGDFDDINYPSVATAQSADFNDPSELNFSNQAQQSACFQPAPVRFN
AEQGISSINDGEEVLNTETVTAQGRDGPSLGVSGGSVGMGASHEAEIHGADVSVHRGDSVVGDMEPVAEVIENLGQSGEFAPDQGLTDDFVPAE
MDREGRLGDSQDRVSQSVVRADSGSKIVDSLKAESVESGEKMSNINVLINDDSVHPSLSCNAIVCSGYEASKEEVTQTWESPLNAGFALPGSSY
TANDQGPQNGDSNDDIVEFDPIKYHNCYCPWVNENVAAAGCSSNSSGSSGFAEAVCGWQLTLDALDSFQSLENPQNQTMESESAASLCKDDHRT
PSQKLLKRHSFISSHGKK
**>XP_009107327.1[*Brassica_rapa*]**
MAQDSEKRFHQIMDKLFTPSKSPLPSSPSSTSSPVEQQSRGKKRPNPSSALALVEPKTALATTIDRSLKVPATGTSQSGLCRPWDRGDLMRRLA
SFKSMTWFAKPQVISALNCARRGWVNDDTDTISCESCGAHLYFSAPASWSKQQVEKAASVFSLKLDNGHKLLCPWIENSCEETLSEFPSMTPQD
LVDRHEERSEALLQLLALPVISPSAIDYMKSSDLEEYLRRHIASGDTTAECSQTESLINHVGASPAQLFYQAQKLISLCGWEPRALPYIVDCKD
KSGEAAKGTDTIDLLPETATRELLSSSSSTSNPNGVSENSENPVVPDTLNSDPSSVVLDCKLCGACVGLWVFSTVPRPLELCRVTGDTEVNTEK
NSRDDTLQRQTSSLQFTIAGGPPATKQNFKATISLPIVGRNLRSRFASYSRDRDHGTDNSIQDQQCRTPERNGGGMENSDQDMIDVGEKADGGR
NASDLVSNTTPQTKDKQLMVVTSSLPENYKPKDSTGDTGISNKQMEFDPINQHRHFCPWIWSTGRRGPGWRQTLSALQRQKGSCQTPPAPSSIF
KVDDPLTSVRNLFKSPSPKKARLNRGSSS
**>XP_033132690.1[*Brassica_rapa*]**
MKEEGESSQNVKPRSKRNSVASASASASATPVNRFRHRSARSPSPPLTAAASSMFNSSVGASSYAVPVNAGSVDWTGQGMGSSGRPCRPWDRGD
LLRRLATFKPSNWLAKPKTASSLVCAQKGWVGVDLDKIQCEFCGSSLHYSPPQNSLKRPEADSNGEEFSKQLDVAHESSCPWVGNCCPESLVQF
PPTPPSALIGGFKDRCDGLLQFYSLPIVSVSAIDQMRASRGPQIDRLLALPQVYSNDDPSFRMGNISATETSKEEALSNYARAQKLISLCGWEP
RWLPNIQDCEEHSAQSTRNGCPSGTARNQSRLQDPGPSMKQFSASSRKASGNYEVLGPEYKSESRSPLLDCSLCGVTIRIWDFLTTSRPVPLAP
INANLPETSKKMGVTRGTSETSGINGWFANGGMAQQQNEEVDEAETSGKRKLVSNTGTSFYQTAAGASSSAQLNMSVTRDNYQFSDRGKEVMRR
QPSGSETGDRAASYESRGPSTRKRNLEDGGSTADRPPYLRIQHADSVEGSVVDRDGDEVNDDSAGPSKRTRGSEVQDTCLPFYGRDLSVGGPSH
SVDAENEREVNRSEGNEQALAFRGARDSARASSVIAMDTICHSANDDSMESVENRPGDFDDVNYPSVATAQSADLNDPSEFNLSNQAQQSACFQ
PAPVRSNAEQGISSINDGDEVLNTETVTAQGRDGPSLGVSGGSVGMGASHEAEIHGADVSVHRGDSVVGSMEPVAEVIENLGEFAPDQGVTDDF
VPEEMDREDRLGDSQDRVSQSVAKADSGSKIVDSSKAESVESGEKMSNMNVYDSVHPSLSCNAIVCSGFEASKDEVTQTWNESPLNAGFALPGS
SYTANGQGPPNGDSNDEIVEFDPIKYHNCYCPWVNENVTAAGCSSNSSSSSSVAEALCGWQLTLDALDSFQSLENAQIQPMESESAASLCKDDH
RAPSQKLLKRSFISSHGKK
**>XP_009149168.2[*Brassica_rapa*]**
MKEDDVSSRNVNPRSNRNSVASASASASAAPVDSLRRRARSPSPPQTAAASSVGASSPAVPVNAGSVDWTGHGLGSSGRSCRPWDRGDLLRRLA
TFKPCNWLGKPKTASSLACAQKGWVSIDLDKIQCEYCGSSLHYSPPQHQLNHPQADSSREEFSKKLDDAHEGSCPWIGNCCPESLVQFPPTPPS
ALIGGYKDRCDGLLQFYSLPIVSVSAIDQMRASRRLQIDRLLAQPQVYAHDDPSLRMDNTLAAETSKQEALSNYSRAQKLISLCGWEARWLPNI
QDCEEHSAQSARNGCPSGPSRNQSRFQDPGPSRKQLSASSRKASGNYEVLGPEYKSESRSPLLDCSLCGVTIRIWDFLTTSRPVPLAPINANLP
ETSKKTALTRGNSATSGINGWFANEGMEQQQNEDVDEAETSVKRRLASNAGISFYQTAAGASSSAQLNMSVTRDNYQFSDRGKEILLRQPSGSE
VGDRAASYESRGPSTHKRNLEDGGSTADRPYLRVQHTDSVEGTVVDRDGDEVNDDSAGPSKRSRGSEVHETYLPSYGRELSVGGPSHSVDAENE
REVNRSDPFSEGNEQAMAFPGARDSARASSVIAMDTICHSANDDSMESVENQPGDFDDVNYPPAATGQSADPSELNFSNQAQQSACFQPAPVRS
NAEAGISSINDGEEVMNTETVTAQGRDGPSIGVSGGSVGMGASHEAEIHGADLSVHRGDSVVGDMEPVAEVIENLGQSGEFAPDQGVTDDFVPE
EMDREGRVGDIQDRVSQSVARADSGSKIVDSLKAESVESGEKMSNVNALMNEDSVHPSLSCNAIVCSGFEASKEEVTQTWNESPLNAGFALPGS
SYTANDQGPPNGDSNDEIMEFDPIKYHNCYCPWVNKNVAAAGCSSNSSSSSSIAEALCGWQLTIDALDSFQSLENAQIQPMESESAASLCKDDH
RTPSQKLLKRHSFISRHGKK
**>XP_012466404.1[*Gossypium_raimondii*]**
MADDPEKRFYSIMDKLFHSSKSTTPFSSSPPAPGTGGQRQLLRAKKRPVPSYTTAVEKPQHCLAASEAPLCRPWDRGDLLRRLSTFKSMTWFAK
PKVVNAVNCARRGWVNVDMDIIACESCGARLLFSTPSSWKRQQVEKAALVFSLKLDSEHKLLCPWIDNTCDERLAEFPPSVPADLVDKFRERSD
SLFQLIALPVISSLAIEFMRSPQLEQFLRQPLMLDCLKGNAEFSHLERIEDGSAVDSAILYYQAQKLLSLCGWEPRSLPYVVDCKDGQNQFVKD
ADILSSSQGVGYGLNLHLSFRPTDENENLEANKGFENSFGLQYDPKSVVLDCRLCGASVGLWAFSTVQRPVELFRLFGCEEVNPGVHDSGHESD
VCEVPFNSGSSSMEQSSNSKLTIAGGPPPTRQNFKARIYVPVIGESLRARLLYHPEIRDQIYSNPKNTLVESNCNRILGEIDCFNNSVNQLGVP
LADLRTLNGKKDGQVNCNSKSSDQSPCSNYDVCSGDDTFRNVTPLEGTDFTAKENSPYTGIDDSNIGGQIESSQNLVLDSCQSNNFPEKVDNDR
TCNLAVKNSDAMLVGESSVMTQGANVSPRNEGAEANDSSVMVTSEKYYPEQNAEPDKVCDKKNCFSNRDSTCVASCLEADVNVDGTNKMNSRED
KTCSNSEEGVIAEVQAVQNNKVLSCPKGKDLKRLHMDKISEFDPIRQHRHFCPWIAPMSGGAPGWQQTLSALLYGKDFPHSSPVCSTSTVSMIK
VDDPIASVRKLFMSPTAKRTKITRE
**>XP_012449448.1[*Gossypium_raimondii*]**
MREEVISSGGTMDPTPAASSAGASSPAVPTNVGSVDWSGHGQNSKAASQSCVGSQAQWISLYSTAGGSALGSSRTSCRPWERGDLLRRLATFKP
VNWFGKPKVASSLSCARRGWINIDVDKIACETCGACLNFASSPSWATSEAEDAGQAFSKQLDVGHKVACPWRGNSCPESLVQFPPTPQSALIAG
YKDRCDGLMQFQSLPVVAASAVEHMRVSRGPQLDRLLYQLQNHMAEFESRSESILEADSARDGAFCLYSRSQKLISLCGWEPRWLLNVQDCEEH
SAQSARNGCSFGPNTTKVHRSQDPGPSKNALIASGKDIGKNKLIVVEARSEYRSPLLDCSLCGATVRILDFLTVPRPARVAPNIDIPDTSKKMG
LTRGVSAASGISGWVAIDDPEKEPTEDRDEVGTTDERNLMQKTDVELNLTMAGSLSFSQLGRAATSRNMNDADMGRDLMIGQPSDSEVGDRAAS
YESRGPSSRKRSLEIGASSEDRPQLRPQQADSVEGTVIDRDGDEVTNARQYSAGPSKRARDSDIFDTYCSPYPRDSSDAGPSHAMGCETSVDGN
KVALFRQGSSHVIGIPSARDSTRASSVIAMDTVCHSADDDSMESVENYRGDVDDIHFPSSSIYGHLDVNDTSELNYSNQAQQSICFQQTAEAVP
GEVGISSTNDGDEIFNAETVTAHARDGLSFGISGGSVGMCASHEADIHGADVSVHRTDSVVGDIEPRIEDAENQGQTGESAPDPGLMDEVVPDE
IDREDPLGDCREMLSRSLGRDDSGSKVDGSAKAESIESGEKISQSCKVIPDNNALPSLSCNANVYSGNETTKEIKNAGKSSSINNCTYPDPDSD
LAVATGIGPPKGESNYEEAIEFDPIIHHNQFCPWVNGNVAAAGCSGYSSSSSCSNADVVALCGWQLTLDALDALRSLGHIPVQTVQSESAASLH
KDDQQTPGKRLLQRHSVNKSHGQH
**>XP_012453776.1[*Gossypium_raimondii*]**
MREEVISSGGTIDPTPAASSAGASSPAVPTNVGSVDWSGHGQNSKAASQSCVGSQAPRISFSTSAGGSALGSSRTSCRPWERGDLLRRLATFKP
MNWFGKPKVASSLACAQRGWVNIDVDKIVCETCGACLHFASSPSWATSEVEDAGEAFSKQLDIGHKVSCPWRGNSCPESLVQFPPTPQSALIAG
YKDRCDGLVQFQSLPIIATSAMEHMRVSRGPQVDRLLSLLQNYVSEFESRSESVPELDVTRDGAFCLYSRSQKLISLCGWEPRWLLNVQDCEEH

SAQSARNGCSFGPNRAQVHLSQDPGPSKNALAPSAKDTGKNKVLVMESRSEFRVPLLDCSLCGATVRILDFLIVPRPARVAPNNIDIPDTSKKM
GLTRGLSAASGISGWVAADDPEKELTEDRDEVGTTDERKLVPKTDVDLNLTMAGGLSFYKLGRATSSRNMNDADMGRDLMIGQPSGSEVGDRAA
SYESRGPSPRKRSLEIGASSDDRPQLCTQQADSVEETVIDRDGDKFNDCRQYSAGPSKRARDSDFFDTYCSPYPRDSSEAGPSHSVGFETHGDG
GNRVALFRQGSNQVIEIPSVRDSMRASSVIAMDTLCHSAGGDSMESVENYRGDVDDIHFPSSSTYGHLDMNETSELNYSNQAQQSICFQPAAEE
VPGEMGISSTNDGEEIFNAEPETVTAQARDGLSFGISGGSVGMCASHEAEIHGADVSVHRTDSVVGDVEPRIEDVENQGQTGESAPDPGLMDEV
VPGEINREDPHGDSQEMLSRSLGRADSGSKVDGSVKAESVESGEKISQSCKLAPDNGAHPSLSCNANMYSGNETPKKEEKDAGKSSSINNCPEP
ESDFAVANGIGPPKGESNYEEAVEFDPVIHHNQFCPWVNGTVAAAGCNGSSADVVALCGWQLTLDALDALRSQGHIPVQTVQSESAASLYRDDH
QTPGKKLHRRRPMNKNHGQ

**>XP_015622239.1[*Oryza_sativa*_Japonica_Group]**

MATGGGGGGDIGADSERRLKKAMDKLYHFPKPKAGTGPGSSKPSSASTSSALSIGRAGKAAGAGGRRFGMVRGSRLPSQLAAMSAISPPPPCRP
WDRADLMRRLATFKAMTWFAKPKVISPVNCARRGWINIEPDVITCEACEARLLFSTPSSWAPQQVEKAAAVFSLKLDNGHKLLCPWIDNICDES
LALFPPTPPPVLVENYHEGFSSLLRLSALPRISCSSLESMKKRSPQLEQFLLKPFSSSVVLKGGFILTEDSTIKDLDHTFQDADTYYQALKIIS
LCGWEPRLLPYAVDCGTKSHSDANSSSTLTQPGLINNSMEDRVVVYAPNEVDGSTVIADARQAYQHYDPLSVVLDCQFCGACVALWPFSLVERP
LQLFKLISDSSRQDEQTEGHAGRVSGAGPSKTANIGFNFTIAGGPPPTRQNFRPRVSLPVVSRHLKADLSSHGHFISSGSDNHMVPVTLHASGL
TKHKRSMDESHMLEGNNTISTDAGTTTNGADHQRENSVNGTSNLVANPEHQQGGSHSDTSRVTSTGEVSNEESETGHAAIKSHTSTDELGQHGS
DPKSLPVEDSSNAHDLAKTCTNNSRPVQAATLTKSSNDGEKGASQPSGSQGLYDKLNEFDPMKQHRTFCPWICPDGGETLPGWRLTLPALLSQD
KRIDEDSQVEPQISLLSEEDDPVTSVRKLFMTPPSKKLRIHRAEKG

**>NP_001389926.1[*Oryza_sativa*_Japonica_Group]**

MREEVRSSSAAPPDPPPRSASPPATPVASSAGASSPPAQTNAASIDWLGGEPISKVESSSQIAPHAPRPSLSTNAAGAAVDFSQPSCRPWERGD
LLRRLATFKSSTWASKPKAASSLACARRGWVNIEMDKIACESCGAHLIFTALTSWSPAEVANAGEAFAEQLDASHLGDCPWRGNSCADSLVQFH
LTPSALVGGFKDRCDGLLQFISLPVIAKSAIESMKLTRSPQIDRVLSQAITILSGELGYKTDSTTGIDINHQDESCSYSQAQKLISLCGWEPRW
LPNVQDWEENSTRSAKHTASADPDQIHSRLPEHKQNSYSASVKKDKGKGKIHVKDSGCSMRSPLLDCSLCGATVRIWDFRSVPRPSHLSINNID
APDMRKGVLTRGISATSGINGWVAEGTERENVEGRGEAGTDEGKSLSNAQVDLNLTMAGGLPSTHSVMPSMHDHFNDGGMGRDLMIGQPTGSEL
GGFAASFESRGPSSRKRNLEEGGSTADKPLNRLHPADSIEGTVIDRDGDEVDDGAQDSDIRSNKRPRGFNLFDVNQPSSSGAGPSRNLSFDLDI
DVNKFDTYKAEGPSALHNPSASMRASSVIAMDTVHSAEENSTESVEYHPCDVDDVHKPSSAVRSGGMSEALDLNYSNQAPQSSFVQPAAESNAR
EIGGSSMNGGEEVLNAETAPAFARDQLSLGVSGGSVGMGASHEAEIHGVDVSEHKTDSVVGDVEPAPELTENMGNTGESAPGPGMMDEFVPEDV
GREEPQGDSQDVASRLVGRADSGSKICGSTKADSVESGEKMSHGIGHESNLQHSLSRNARVYSGIDLSKDEVTQIAKLPANDDYDPGDDLAANG
GNDYEAGLPEFDPISHHNNYCPWVNGHVAAACCINTGSSTSTGLSGWQLTVDALETIQSLAQAQNQIMPSDSAASLYKDDHVAPSRKLLKRASH
SKC

**>XP_015630572.1[*Oryza_sativa*_Japonica_Group]**

MREEVRSSSGAAAEPPPTPVASSAGPSSPAMQANVASIDWSGSRQASRVDSSSHVAPHAHQPSHSFDATGTALDSAPSCRPWERGDLLRRLATY
KPTTWASRPKAASSLACARRGWVNVDMDKIECESCGAHLIFSTLTSWSPAEVSNAGEAFAEQLDASHHNSCPWRGNSCADSLVQLHLTQSALIG
GFKDRCDGLLQFTSLPVIASSAIEHMRLTRSSQIDRLLSQSITFLSGELSYKAESTTGIDIQQDSSCSYSKARKLISLCGWEPRWLPNVQDCEE
NSTHSAKNADSVEPFFPRFAEHQKNSFSGSAKKDKGKGKRPLKDSGCSMRSPLLDCSFCGSTVKIWDFRSVSRPCRFSPNNIDAPETGKKLALT
RGISAASGINEWVTDGMERDPAEGRDEEATNEGKSLSNAGVDLNLTMAGGLPSIQSSIPIASERFNGGLGRDLMIGQPTGSEVGDHATSYESRG
PSSRKRNHEEGGSTVDKPQDRLQHADSIEGSVIDRDGEEVDDAAQDSDIPNKRSRGFDLFGSYLPSSSGAGPSRNFCFDPDADAGKFSHARAAG
LAAVDRDSMRESSVAAMDTVHSADEDSMESVEYYPGDGNDIDMPSSSAHRNIEMDDVLGLNYSNQAQQSACVQPASGSDGREIGGSSTNEGEEV
LDAVTAPAFARDQLSVGISGGSVGMGASHEAEIHGIDVSLQRAESVVGDAEPNTELTETMGHTGESVPGPGLMDEFVPDEVDRQEPHGDSQDMV
SQSVGQADSGSKIYGSTKADSVESGEKIGGHAVGHASRMHPSLSCNAGMQTGLDVSKEEVTQAGKLLIAGDVPMGLDYDPQNGLGATNGENDFE
SGLPEFDPVKHHNSYCPWVNGTVAAACCSNTESSSSSSPLSGWQLTVDALDTFQSLGQAQNHAMRSDSAASLYMDDHVTPNHKLARRASVSRSH
GKC

**>XP_039810120.1[*Panicum_virgatum*]**

MAAGGGGGGDIGADSERRLKKAMDKLYHFPKPKPSGPGGSKPSSSSAPAPSSGRPVGKAAAEAARRFGLVRGSRLPPQVTAMSAISPPPPCRPW
DRADLMRRLGSFKAMTWFAKPKVISPVNCARRGWINIEPDVITCEACGARLLFSTPSSWTTQQVEKAAAVFSLKLDTGHKLLCPWIDNICDESL
ALFPPTPPPVLVENYYECFSSLLRLVALPRISCSSLEIMKKRSPQLEQFLSEPFSSSVVLKGRFVLTEDSTIKDLDDAFQDADTYYQALKIISL
CGWEPRLLPYAIDCGTESHSDASSSPKLAQPQQISKTMEDRIILYSPNDANGARASADANREDQHYDPLSVVLDCQFCGACVALWPFSLVERPL
QLFKLVSDSNGQDDKDNGHANVVCGVGHSKDANIGFNFTIAGGPPPTRQSFRPKVSFPVVSRHLKADLNSRGNSLSSGSDGHMVPVASKALGSM
KRKRSTDQPDLLEGDTDDVDTSPIGAKSHQPGDNSEKSMPNPEVMNEQEQGGSHSDTDKYINMDGASNEKQPESSSPSRKSITSTDAALDQHGS
EPRFPSVQGTNEEPSNGVTLAETHANNSRPTELSTVTKSLVNKEKGAYGPSEKQGLYDRMNEFDPIKQHRTFCPWVSPDISESLPGWRLTLTAL
LAQDKRYDGDSRGEVQIGLLDEEDDPLTSVRKLFMTPPPKRRRIQQSEKS

**>XP_039848146.1[*Panicum_virgatum*]**

MPNGRQQLPAAAACKRITTPSPSLPPPIPHPAPATSMAAGGGGGGDIGADSERRLKKAMDKLYHFPKPKPSGSKPSSSSAPAPSSGRPVGKAAA
EAARRFGVVRGSRLPPQMAAMSAISPPPPCRPWDRADLMRRLGSFKAMTWFAKPKVISPVNCARRGWTNIEPDVITCEACGARLLFSTPSSWTT
QQVEKAAAVFSLKLDTGHKLLCPWIDNICDESLALFPPTPPPVLVENYYECFSSLLRLLALPRISCSSLEMMKKRSPQLEQFLSEPSSSSVVLK
GRFVLTEDSTIKDLDDAFQDADTYYQQALKIISLCGWEPRLLPYAIDCGTESHSDASSTPKLAQPHQISKTMEDRVILYSPNDANGARASADAN
QEDQHYDPLSAVLDCQFCGACVALWPFYLVERPLQLFKLVSDSNGQDDKDNGHANVVSGVGLSKDANIGFNFTIAGGPPPTRQSFRPKVSFPVV
SRHLKADLNYRGNLLSSGSDSHMVPVASETSGSMKRKRSTDQPDLLEGDTDDVDTSPIGAKPHQPGDNSEKSIPNSEVGNGQEQGGSHSDTDKN
INMDGASNEKQPESSSPSRKSITSTDAALDQHGSEPRFPSVQGTNEEPSNGATLAETHANNSRPTELSTVTKSLANKEKGAYGPSEKQGLYDRM
NEFDPIKQHRTFCPWVSPDDSKSLPGWRLTLAALLTQDKRSDGDSRGEVQIGLLDEEDDPLTSVRKLFMTPPPKRRRIQPSEKS

**>XP_039827893.1[*Panicum_virgatum*]**

MREEVRSSSGAAAEPPLAVARSSSPPHTPVASSAGASSPALQTNIGRQASRVDSSSQVAAHAYHPSHSFDAAGTAMDSAPSCRPWERGDLLRRL
ATFKPSTWASKPKAASSLACAQRGWVNIDLDKIECESCGAHLIFNALMSWSPVEVASAGEAFAEQLDAAHQNSCPWRGNSCADSLVQLPLTQSA
LIGGFKDRCDGLLQFTSLPVIASSAIENMRTTRSAQIDRLLSQSITFLSGVLGCKAESAAGVEIQQDSSCSYSQAQKLIGLCGWEPRWLPNVQD
CEENSTHSAKNAPSVGPDEPFYPHFVDHIKNSFSASAKKDKGKGKLPLRDSGCSMRSPLLDCSLCGATVRMWDFRPVLRPSRLSPNNIDVPETG
RKLTLICGISAASGINEWLTDGVERGQEEGRDEAATNEGKSPLIIGVDLNLTMAGGLPSPRSATPAASERFNNGGMGRDLMIGQPTGSEVGDCE
TSYESRGPSSRKRNLEEGGSTADNPQDRLHHADSIEGNFIDHDGEEVDDAAQDSDVPNKKSRGFDLFDAYRPSSGAGPSRNLSFDPDVGAGMFS
SSRSIDLAVERPAARDSLRASSVIAMDTVRTSEEDSMESVEYYPGDGNDIDMPSSSAHRNIEMNDVLDLNYSNQVQQSANAHAAAGSDAREIGG
SSINEGEEVINAETAPAFGRDQLSIGISGGSVGMGASHEAEIHGNAASLHRAESVVGDAEPIAELTETMGQTGESGPGPGLMDEFVPEEVNREE
PHGDSQDMVSRSVGQADSGSKIYGSTKADSVESREKIGHATGIESSMRPSLSCNAGMCAGFDPAKDDVTQSGRIILTTDDTLMGLDYDPGNGLG
ATNGENDYEAGLLEFDPVKHHNSYCPWVNGIVAAACCNNIGSSSSSSALSGWQLTIDALDTFHSLGQAQNQIMQSDSAASLYMDDQNQITNNRR
LGRRPSVSRSYGKC

**>XP_039782290.1[*Panicum_virgatum*]**

MREEVRSSSGAAAEPPLAVARSSSPLHTPVASSAGASSPAMQANIGRQASRVDSSSQVAAHVYHPSHSFDAAGTAMDSAPSCRPWERGDLLRRL
ATFKPSTWASKPKAASSLACAQRGWVNIDLDKIECESCGAHLIFNALMSWSPVEVASAGEAFAEQLDAAHQNSCPWRGNSCADSLVQLPLTQSA
LIGGFKDRCDGLLQFTSLPVIASSAIENMRMARSAQIDRLLSQSITFLSGVLGCKAECTAGVEIHQDASCDYSQAQKLIGLCGWEPRWLPNVQD
CEENSTHSAKNAPSVGPDEPFYPHFVDHIKNSFSASAKRDKGKGKLPLRDSGCSMRSPLLDCSLCGATVRMWDFKPVLRPSRLSPNNIDVPETG
RKLTLTRGISAASGINEWVADGVERGQDEGRDEAATNEGKSPSIIGVDLNLTMAGGLPSPRSATPAASERFNNGGMGRDLMIGQPTGSEVGDCE

ISYESCGPSSRKRNLEEGGSTADNPQDRLQHADSIEGNFIDHDGEEVDDAAQDSDVPNKKSRGLDLFDAYRPSSGAGPSRNLSFDPDVGAGMLS
PSRTIDLAVERPAARDSLRASSVIAMDTVRTSEEDSMESVEYYPGDGNDTDMPSSSAHRNIEMNDVLDLNYSNHAQQSANAHAAAGSDAREIGG
SSINEGEEVINAETAPAFGRDQLSIGISGGSVGMGASHEAEIHGNAASLHRVESVVGDAEPIAELTETMGQTGESGPGPGLMDEFVPEEVNREE
PHGDSQDMVSRSVGQADSGSKIYGSTKADSVESREKIGHATGIESSMRPSLSCNAGMCAGFDPAKDDVTQSGRKILTTDDALMGLDYDPGNGLG
ATNGENDYEAGLLEFDPVKHHNSYCPWVNGIVAAACCNNFGSSSSSSALSGWQLTIDALDTFHSLGQAQNQIMQSDSAASLYMGDQITNNRRLG
RRPSVSRSYGKC
**>XP_039793133.1[*Panicum_virgatum*]**
MSVSVGKTRVGRYELGRTLGEGTFAKVKFARNVETGENVAIKILDKEKVLKHKMIAQIKREISTMKLIRHPNVIRMYEVMASKTKIYIVMELVT
GGELFDKIASRGRLKENDAKKYFQQVINAVDYCHSRGVYHRDLKPENLLLDASGTLKVSDFGLSALSQQVREDGLLHTTCGTPNYVAPEVINNK
GYDGAKADLWSCGVILFVLMAGYLPFEDSNLMSLYKKIFKADFSCPSWFSTSAKKLIKKILDPNPNTRITIAELINNEWFKKGYQPPRFETVDV
NLDDVNSIFDESGDPAQLVVERREERPSVMNAFELISTSQGLNLGTLFEKQTGSVKKETRFASRLPANEILSKIEAAAGPMGFNVQKRNYKLKL
QGENPGRKGQLAIATEVFEVTPSLYMVELRKSNGDTLEFHKFYHNISNGLKDVMWKPDGSIVEGDEARHRRTATSKKKNSSHTPRARPASSAPP
PDPTRRGLLPRRPRERASAAAMREEVRSSSAAPPDPPPARSASPPPTPVASSAGASSPPAQAIVASIDWLGSDQVSKAGSSHVAPPASQPALST
NADGAAADFFQSSCRPWERGDLLRRLAMFKHSTWASKPKAASSLACAQRGWVNIDVDKIECESCGAHLIFTALTSWSPAEVANAGEAFAEQLDA
SHQNDCPWRGNSCADSLVQFHLTPSALVGGFKDRCDGLLQFVSLPVIASSAIESMKLTRSFQIDRILSQSVTILSGELGYRTDITTGIDINQQD
ETCCYSQAQKLISVCGWEPRWLPNVQDWEENSTRSARNAGSAEPDGQFHSQFAEHRQSSYSASVKKEKGKGKMRVKDSGCSMRSPLLDCSLCGA
TVRIWDFKSVPRPSHFSLNNIDMPDTGRKPVLTRGISATSGINGLVAEGAEKENVEGRDEGGTDEQKSVSNAQVDLNLTMAGGLPSNYSALPPM
PGHFNYGGMGRDLIIGQPTGSELGGHAASFESRGPSSRKRNLEEGGSTADKPINRLQPADSIEGTVIDRDGDEVDDAAHDSGARNKRPRGFNLF
DINRPSSSGAGPSRNLSFDLDIDVNRFDTSNAEGPSALHNTFPKDSMRESSVIAMDTVHSAEENSMESVEYHPCDGDDVNKPSSALRSGGMSEA
LDLNYSNQAQQSSFVQPAPETESNAREIGGSSMNGGEEVLNAETTPASARDQFSLGVSGGSVGMGASHEAEIHGTDVSEHKTGSVVGDADPVPE
LIETMGHTGESAPGPTLMDEFAPEEVGREDPHGDSQDMASRLAVRADSGSKVCGSTKADSVESGEKMSHAVGPENSAHPSLSCNARVFSGVDAS
KEEVTGIMLTNDDYDPGNGLGATNGENDYETDLPDFDPIKHHNNYCPWVNGNVAAACCISTGSSTALSGWQLTVDAIETLQSVGQAQNQTRQSD
SAASLYKDDHAPPRRKLLKRANHSRS
**>XP_039834233.1[*Panicum_virgatum*]**
MSVSVGKTRVGRYELGRTLGEGTFAKVKFARNVETGENVAIKILDKEKVLKHKMIAQIKREISTMKLIRHPNVIRMYEVMASKTKIYIVMELVT
GGELFDKIASRGRLKEDDARKYFQQVINAVDYCHSRGVYHRDLKPENLLLDASGTLKVSDFGLSALSQQVREDGLLHTTCGTPNYVAPEVINNK
GYDGAKADLWSCGVILFVLMAGYLPFEDSNLMSLYKKIFKADFSCPSWFSTSAKKLIKKILDPNPNTRITIAELINNEWFKKGYQPPRFETVDV
NLDDVNSIFDESGDPAQLVVERREERPSVMNAFELISTSQGLNLGTLFEKQTGSVKKETRFASRLPANEILSKIEAAAGPMGFNVQKRNYKLKL
QGENPGRKGQLAIATEVFEVTPSLYMVELRKSNGDTLEFHKFYRNISNGLKDVMWKPDGSIVEGNEARHRRTPRRFHQTKSMDNVNEGMCRVSI
GLSKHRRGPAGDLDVLRRSCILSVADPDAVVAQFSCACLTGPSTPALAAATTPTAVGGNGLSSDNDEQEGEDLALVRIIRKNLHRVNKASPASS
LPPGDAPGGGDTSSPRSILFGGSIAETTQPRQEEEPKAQGRSITAQSASAELNGASPRSFAHAALALPADLLCSARPDPTRPPPPTTLPQSERP
EAAMREEVRSSSAAPPDPPPARSASPPPTPVASSAGASSPPAQGSDQVSKAGSSHVVPPASQPALSTNADGAAADFFQSSCRPWERGDLLRRLA
TFKHSTWASKPKAASSLACAQRGWVNIDVDKIECESCGAHLIFTALTSWSPAEVANAGEAFAEQLDASHQNDCPWRGNSCADSLVQFHLTPSAL
VGGFKDRCDGLLQFASLPVIASSAIESMKLTRSVQIDRILSQSVTILSGELGYRTDSTTGIDISQQNESCCYSQAQKLISVCGWEPRWLPNVQD
WEENSTRSDRNAGSAEPDGQFHSRFAENRQSSYSASVKKEKGKGKMRVKDSGCSMRSPLLDCSLCGATVRIWDFKSVPRPSHFSLNNIDMPDTG
RKPVLIRGISATSGINGLVAEGGEKENVEGRDEAGTDECKSVSTAQVDLNLTMAGGLPSSHSALPPMHGHFNYGGMGRDLIIGQPTGSELGGHA
ASFESRGPSSRKRNLEEGGSTADKPINRLQPADSIEGTVIDRDGDEVDDAAHDSGDRSKRPRGFNLFDINRPSSSGAGPSRNLSFDLDIDVNRF
DTSNAEGPSALHNPFPKDSMRESSVIAMDTVHSAEENSMESVEYHPCDGDDVNKPSSALRSGGMSEALDLNYSNQAQQSSFVQPAAETESNARE
IGGSSMNGGEEVLNVETTPASARDQFSLGVSGGSVGMGASHEAEIHGTDISEHKTGSVVGDADPVPELVETMGHTGESTPGPMLMDEFAPEEVT
REDPRGDSQDMASRLAVRADSGSKVCGSTKADSVESGEKMSHAVGPENSAHPSLSCNARVFSGIDASKEEVTGIMLTNDDYDPGNGLGATNGEN
DYETDLPDFDPIKQHNNYCPWVNGNVAAACCINTGSSTALSGWQLTVDAIETLQSAGQAQNQTMQSDSAASLYKDDHAPPRRKLLKRANHSRS
**>XP_002963700.1[*Selaginella_moellendorffii*]**
MTSGDKDDDAEQRIQRAMDRIFSSPIQAASSCSSSPPTKSILERRVVTPLALERSPGEKNAGEKNAGEIASSSTPSCRPWDREDLLRRLGTFKS
VSWFGKPSAAGPVACAQRGWINVDMDLLCCEVCGSRLSFSFPATWSKKEVETAGLEFSRKLHDGHKTSCPWKGNGCGEDLAAFPPTPAPVLVQA
YEARLQSVALLSDLPVISSSTVERMKISRGDQVASLLALPSNDAAVRELEAAQGEAVQKLRQTYEAFLQAQKLISLCGWEVRLLPYAVDSHDSN
VDSHELRVSLATGSDPCSAVLECRLCKASVGLWRFRTLSRSSLSITAILSTIEASAKKNVEVLPAGDVNAHVDDNAAENIDTVNAEATISEDND
AAAVDDSKKNPGLDLTLTIAGGPRPTRLSAPSPASIPIPGLNSQRHDDFQPRKSTLEQEPAEKTTVQDRSSSKRKRDSESAHRKKLKAVDGLPG
SSSVNAVETSYNHNRHENSAESVECSPQGSDEEQDTVLSREKAASQDVAKEAAIFSGSPSEFDPVHHHRYFCPWISSNAADQSGKCGWQMTIDA
IFSCAATNAKSGVVSDRDKAAAVNKMDPLVSVRRMLGGKGKGVAVNGGAISMSRAGT
**>XP_002981891.1[*Selaginella_moellendorffii*]**
MAEDAAERRMERPLFGVSPPTPTASSAGPTLAVQGNYASIDWHGLAQKRPAISGTSAGPPRPAVSTSTAAVSGSSSHRHLCRPWDCGDLLRRLS
TFRTSNWNAQVIGPAVCARKGWVNVDVDMIACEGCDTHLSFALPLQTEVEAASESFRKQLETSHQRSCPWKGNACSESLAQFPSSAMALIGGYN
DRCDALLQLPSLPVVSTFAVDQMRLSRGPQIERLLSLPAMHFPERNGAGPLTDDFVKAQRIISFCGWEARLLPHAGDLEDYSAHSSRKPNKASA
SLKRAKRTMPRSDKEGDSRGARQTASTLLECVYCGASVPILRFQTVARPGGGSTGSEYPSSDNKSLPLVRGASAASSIDIHKRRQRLEGGEAGE
ATEQKPPSLTITAARASPTNLNVTPVARPEQQPEGSEMGDCGVASYESRGPRDDHQGTAAQGDSSTYVPQLRAESAGGTNGDFDNEDEGKNAAE
SSKRKRVPESLAADQHSLGMAVSGNAATCEQEGETEDRKNKKVCIGPATAIERTVSLEVPDNTYNRSARSLPVRQENLRRRVTREYPCSSSVNA
IDTCFQNKMEDSMDSVEFAPQDDAQQHTAASGYTENEWLFDEPNSTVQGQQSNSYAPAAIQEQATGETNAAVISTGTATGCGGSVGMAGIRLGS
QRVQSNEADIQGAELSELQTESVAGEPADAVVDAVVDQGDPGLVCDSTPLTAAQDCAGGGDESGESGSKASLRPRNGFSEERTIDTTNAVVTGC
SIGTIAKEMTEGSVEVAAVGDNVRAVNVNSAPTVKEAEDTLFDPIRCHRSFCPWVNGLVLSAANGTFGPGPVYCGWQLTVDALDAFHQQENASA
GVTESESTASMCKDDGRGRIFAQGSSSQPCNGSSRLEPN
**>XP_024380728.1[*Physcomitrium_patens*]**
MEVTEVSEKRFERAMERLFSGSSTPAASSARVPSPVKEKKEDEENTSGRVFQTAGVATPLSGSHGCRPWDRGDLLRRLATYKSISWFGKPQVAG
PVACARRGWVNVDIDLLACEICGSRLSFPVSSSWSRHQVEQAALVFAEKLDTAHKGLCAWKNNPCAETLAHFPPTPVSVLRGAYTDRCEALLQL
SALPVISDAAKSLMKLSRGPQVDQLLSELNPPSPGFLVGNGASSSSTENEIFANSKAYYEAQRMIAVCGWEPRLLPYTVDCEDRSGAQSIHEQI
GTSHGPGPSVTVHMQGGQQSKVAQGQIQSTGTDVSDPASAVLDCNLCGASVGLWNFATLNRPAPLLNSGLEELFSSKNRSSGSNPVRDDAAGAV
GGPLLASPEAAEMVDAKPPEVVPVNVLERAAPPKGVLDLKLTIAGGPPPTRLIAPASVPPSFGIPGLPHTVMQPRKIEAITYAAASYESRRPAH
QGRGHNDTGTPTYHVEGEVIQNREMKDAEESKDAEYFSKRKREKGNDESEGVFSPNTKRKREVGSQWPGFTVKLPARDLPHASSVNAIDTCYPP
KQENSMESVDNLPLGSDGQGANTAEYHVNGPESRVQAEHSMSRQDGIDAGKSICGGRVEEGVQNNVVVICPGAGISGSHETEIQGVEANLERSE
SVAEFATDVIELMEEHVSGRGLMDEFIPEDTLKAIVTDDHEGSRQAMLGISHTFVKDTSSAGVSERYQRNVAQRDADEDANAGSTLETTNAIIQ
GVDMVANIQVKEVTAGRTVGESPSERNEEVQSHPSFPPGLLLDDVKKLEIQTGEFDPIRQHRHFCPWVNAHVAAATSGTGSSKFCGWQIVLDAL
QPQPPSPHQQHQHQQGSVESEFTGSKYKDDPILSVRKLLGSVSSNYLPSS
**>XP_042921239.1[*Chlamydomonas_reinhardtii*]**
MSSVYERITSALSSLGKRRERDSSAHEGDGAGASAAGGGPTSPGGRSAAARTPKRFRPWEQADLHKRLETYKPLTWFGKPASVGPVPCALKGWV
NDGSDCLTCEYCGSKLVYPPHVAYDQRQAAADMFSPSLTTKHTATCPWRQTACQPKLLAYVPSTTPEQLCSLFYSLADKLMRVDVLPDMDTLAI
QTLRSTAMPYGSYDDFITAAAPGGGAVVGGGGAAGYSHDLAPRRRQMPSATIRELDQNGDEVMTPSGSAAGAAAAMAPAAAAGGGGDAAAVLQA
LVAAGDAGEGQAVLVQTSKLAPAQKARLLALLGWDVDVLQPDSASGMAVAPFAAGGSYSLSHLGVKPKAAAAAAAGAGAGGAAVPGTPGGAGGK
GGKGSSKVPSSQVVLKCPICNSRMGLWNYSGVRPVPVGRLTAPPPPAAGGAAALMLSPRPAAASSGGGAAAAAAPAVPATIGSDPLSCTIAGGQ

YGQFGFGGAASAAKPFGSAAAAAPFRFGSAASTAPVFGLAAMDVDAQRAASAASGPFGSAAAAAAATPSMPAPSGSATPAPAGRKRKAEAPEPM
ALDAQHTPSAGMATPVAAPDGKRQRMAATPLWGGAGFGAVGGPAASPSGLGLGGASALAAASAAGQPRELDPVAQHRSWCPWVYTGSGDEKHMS
GWQHMLSALSQHQQHQQQQANVAATPGAAAASPADARQLRDNALEAIRKL

#### Representative sequences for Figure 3D:

**>OTA34089.1[*Hortaea_werneckii*]**
MAEAIATKKRNFYKSLDAFNNPNPSSTSVATEPATKRPRRNLSAASAASRLTTATANHPATPAKPNQSPKPPPAFSPWSQDTFLARLRTFSRVS
LWHPKPQSISEVKWAKRGWSCVDVNTVACKGCCGKRVVVSLDFAKTESVNRGEDVEGDGDNGEAQENSEQDEDELEAALALKYQALIVDGHSDS
CPWRRTGCPDDIYRLQVIRAASWQLELRRRYQSLHQISDAIREVTLRGSSQDKQSLIPIDQLLADLPADVLGPPGEEQPAPEDSLKALEIAMHG
WKGSEDSGNELLHCDACFQRVGLWMYQPGYKPARSSSDDEDQTAIVDLVELHREHCPWRNPDNQCALGTLKGLNACQVLQTCVSAFVKDERRRD
ERQRKSVHQPETTEDEQEAESSPPSPAPSRDEIEKQDKERESRLKRLKSLFTIKRKSTVVAPPNPKPSIVGKRPATRG
**>OTA24219.1[*Hortaea_werneckii*]**
MAEAIATKKRNFYKSLDAFNNPNASSTSVATEPAAKRPRRKLSTASVASRHTTATANNPATPVKPNQSPKPPPAFSPWSQDTFLARLRTFSRVS
LWHPKPQSISEVKWAKRGWSCVDVNTVACKGCCGKRVVVSLDFAKAEGVNRGEGVKSDDDDDAQGDTEQDEDELETALASKYQALIVDGHSDSC
PWRRTGCPDDIYRLQVIRAASWQPELRRRYQSLHQISDAIRDVTLRGLPQDKQSLVPTDQLLADLPADVLGPPGEEQPAPEDSLKALEIAMHGW
TGSEDSGNELLHCDACFQRIGLWMYQPGYKPARYSSDDDDQTAAVVDLVELHREHCPWRNPDSQCALGTLKGLNACQVLQTCVSAFVKDERRRE
ERQRRGVQQPDTNGDEEGHESAPPSPAPSRDEIEKQDKERESRLKRLKSLFTIKRKSTAVAPPKPKPSLVGRRPATRG
**>KKK13521.1[*Aspergillus_rambellii*]**
MSYALETKKRKFHRVLESLTRPLNGESSPKLTSASSAPSLEHAPSAKRARLSGLGDGDFPSVRKKTFQPARPSSSNSSSSFSPRPSFVPWDRDR
FLERLETFRRVDRWSPKPSAINEVEWAKRGWVCSDVSRVTCAGGCGGSVVVKLPDELDELDGYDSEKIQERKEVRTKLVNEYANLVIQGHGENC
PWRNKGCDATIHRLPLANPDTAISGLQTRYSHLLKMADKLPSLVDLQLPEHWDLQAIISVLPLEGFQGLESQVETTDTQPPAGGDESQQRELQT
STHPEPAVNESAFVLAFFGWDSVADGAVGLAGCGACFRRLGLWMYKPRQEGKSSAHDPLDVVNEHMEYCPWINGKTQSGTGKPSEKMEGLRSGW
ELLAQGLKVKHLRYIRSTEPIGSRAGSEAPSVGDSVAEETSDDTKKAKDREWWAKIRRMRQVLNVKSPKPKT
**>XP_037155881.1[*Letharia_lupina*]**
MPTGTALSTTKRKFHKILDSISNASSTSLATKSNHDNYNASTTTLPTTMDPPAKKPRIVRPASAYVPPSTRILTSQSPNLRAAAATAKQPSPIV
TMNEERKTPNFAPWDRGQFLERLKTYRHVDKWMGKPERINEVQWAKRGWSCVGKERVECVGGCGKEVVITLESSREEKHDNGETQDTEKRPSDE
EEDEDEWREKAQEQLVEKYAEMVVTSHDGGCLWRRRGCDDTIQRLPLAHHKTATDELRQRYTSLVAVASELPPDPSTPEGFDLSSLSQKLAPLL
HPPSPDPPSPTPSSPPNPPPNTTTSLPPINNSALALALFGWRAEEGHVVGLATCTACFRRLGLWLFKPASESPSHSSMDRLDVVGEHRDYCPWI
SPLSQNGATSRRTSLDGLAGWEALLRAVNASAIHKRHGDEDTTPVTARAADGLGSEVASLAHSSATREELGVRDERDKERWAKLKRLKQAFHVK
RRKGKDNGETKGGNGKV
**>XP_957690.1[*Neurospora_crassa*]**
MNATVKRKFNSLLQGIGNRPTNPDSPSSTRDNDSLSSTPASSSSSRFTNMANDSLDYLSKKRRVGGLPSTPSAITLTTPTKGQTTISNVTLRKW
NSHGGPGSSPAPGAGGGSSNAKGDSPVTKLQPPKYCPGDRNQLVRRLATFQELTDWTPKPDRVNEIEWAKRGWVCQGKERVKCTLCNNELAVKL
NRKEVDGKEIPVLIAADIAESVVDQYVELIITSHREDCLWRKKGCDDSLLRLPLPNPKLALETLRQRYDELCQRKDFLPYEFNLRLPKGLDIDI
ILSYLPSNFFAEPPASSTVDSSVASQPASPNPSTSQQPQAVNRTALALALLGWQGLTNPRLGTAVPNSASCHTCLRRLGLWMFKSKQVDPETNT
ILVPAPMDHLDPLREHRFFCPWKNPQAQRNPGAKPLARGETNKAAWEVLVEGLKNESRLREKARDLMHGRSKSSSSGFGFGLGGKKGTPHRATG
STSGFLGVPTTPDGRGVTGQQPNSAPGGLQVGGQEGGGQEEEGFEEEDDDESEEARKKKDQAMMSRLKRVKTLFNTKGSKLKKSGASPSPSAVN
IPDSPRPGSSHSTRTTGTPGTAATTGISTNAPAE
**>PUU81978.1[*Tuber_borchii*]**
MLYSTKRKLHNLINGPPAPSTSISAPSTPAKQPTDLPTTTTAAFDDNDVVLIAPAVADAAKKRRLGGLGNRSRPSVRAVSPTGSIRSTVSTTSS
QQMPTYSPWDRAAFLERLRTYRFVDKWSAKPVDVNEVEWARRGWSCIDKNRVRCGVCKREVVVKVELDEEQDSDITKAVVEKYKEMIVTEHEDR
CLWKKRGCDDTIYRLQLANPSVSRPSFVSRYSSLLRIRPEIPPSLSYPTADFPEHILETIHENTLLLEEQHSGALVLALFGWQNEDPGIPSLVT
CSACFRRLGLWLFRKKVVSIFDSVDEQEEASVCRLDVIGEHRDYCPWINATNQGTEPGWQIMLRILQQPNKGPALGSYTERDAEDQSKLKRLRK
LGAMYLGATKKTKGKGKESKERPKTPKTPKTPTKEAPPPKTPKTPKTPSKDASKTPKTPSRDPPKTPKTPSREAQKTPGRSEE
**>NP_013600.2[*Saccharomyces_cerevisiae*_S288C]**
MEKDALEVRLKSIRHSLDKNTKLLPGKYRNTLGERLITKWRYKKKSHNGSSMLPEKCKSHVQLYDDLVQESSKHFVGFRLHDLRALLKRICSIQ
NYTRHVLIEWDVRWVNPLTLASKGWEPYQSASQSQVPFKCCCCHAIMTIPLLKNGDDVADYTMKLNEKIWNSNIIGNHLQKCPWRENQVDLNKE
YYLSSQNLIREIERIHTEIDRIVSGSNEFSLKRNSSRIFHYLSEKEIQKLAFFFDCKDYSLVGLLLLGYTKFQKDDLVQCTACFHRASLKKLEY
TEFNGHALWCRYYNKELLPTMLLELIGKEDKLITKLGVGERLNKLEAVLQTL
**>NP_588231.1[*Schizosaccharomyces_pombe*]**
MSFPTDMETNEILDQLDKIDERNEDILLKSLKASKCTYKPWSREEFLRRLLTYRSRWAYVNDPQIGEINCCLNGWLCESNNILVCDVCRNKINL
TALQQVDAENDSLNELPEKTKERLEVSLKEEHQDNCLWRLHKFPPDIYHLSVSAELVQVGRRFNSLSTRLVSTHLPEEMTLKRLEKVANKIRVD
WEKEDAAVLLGIALTGWSEQVPGRLYVCNYCHRRLGVWNLQSEGQDFDVLEEHKKSCPWVIPQPFTDLLGWQQIFELLCKESIFQSTTKTMDVS
QYTDYTFSLLQGLR
**>XP_024513553.1[*Cryptococcus_neoformans_var._neoformans*_JEC21]**
MELSSNTDDDLRDVFKLLYADDDWALTSDSELDDSEQLGNADGSEIDVADDEEQHTIRIYSGRITKKRLFSALDSLLSPGYETDTKRQRIYNPP
APSIPSLILSTQPMPALPLSKVYAPFSALSLLSRLMTFQPYTYSPQHPLTLSPVRAAMKGWVNEGREGLKCDVCGARWGLGGLEKVRDEAMKSN
LGERLAKGFEERHEKNCAWRICASPGNLYEQLRHLVHPPITSSLAPLASHLLLECLALPSLRLLSPLNPLQVERLVSLFKPSSTFSIPSPATDV
ASQLALFGWFPYHPNYPTIQISLNTPSSRTEIVCCRICHRRIGLWNFSNEKDGVKRFDVLNEHLVWCPVRIQDGEKEWWSESGLLDGQSTQAKR
IGEGGIKGLVKVSEKMEKRSWRRS

#### Additional sequences:

**>NP_001298015.1[*Mus_musculus*]**
MAATSEGPLFAASIEKTWGSVVRSPEGTPQKVRELIDEGIVPEEGGTEPKDTAATFQSVDGSPQAEQSPLESTSKEAFFHRVETFSSLKWAGKP
PELSPLICAKYGWVTVECDMLKCSSCQAFLCASLQPTFDFGRYKERCAELKKSLCSAHEKFCFWPDSPSPDRFGMLPLGEPAVLISEFLDRFQS
LCHLDLQLPSLRPEDLKTMCLTEDAVSALLHLLEDELDFHADDRKTTSKLGSDVQVQATACVLSLCGWACSSLEPTQLSLITCYQCMRKVGLWG
FQQIESSMTDLEASFGLTSSPIPGVEGRPEHFPLVPESPRRMMTRSQDATVSPGSEQSEKSPGPIVSRTRSWESSSPVDRPELEAASPTTRSRP
VTRSMGTGDSAGVEVPSSPLRRTKRARLCSSSSSDTSPRSFFDPTSQHRDWCPWVNITLVKETKENGETEVDACTPAEPGWKAVLTILLAHKRS
NQPAETDSMSLSEKSRKVFRIFRQWESSSSS
**>CAB55333.1[*Yarrowia_lipolytica*]**
MHNTSTYHHTMPTEEQLHGIMESLREVTRKSGSAPTTPTKHSPRNSTLKTATPNRFTTASSTSGKISKRPSLMERIRQVKEKDYRRTTKEITEL
APTTTATSTPATYTPWSKEDFLDRVSTYTYQKYPIETSLYPKLSPYNVARYGWKCTSSKMLQCVSCGSYLAVVCGEEDDEATIKVVQDKYLGLI

TRNHSTRCLWKNKPCSESLGSIMGNIGRLRKDLGGKIAIPEGVEVTVDGQSVEDFFKKLLLEVDGEVNKEIDENRGLDLKEEVKENGSGSNKDS
NDLTQSAELGSDSKLIPEEVSRITLAAGWRESHNMFKCHVCSRIVKPSETFDVVQEHRDWCPYVVEKEGDKPAWWQVLTKPSKESSKKRLSNIR
EVYFGV

**>XP_002634353.1[*Caenorhabditis_briggsae*]**

MEVDSANNRHAAVLKRKATDSINEVLSFGQSSSSPQKRTKKTPLHKIRDVEAYKKIIKTYKAPTWFGCAVSPRDLADYGWECVKKDCVKCVECE
ELLCTSLPNICKVSFNVYNSRLQEIHDQLSSAHRTTCKLRTGAPPIRFTEPTSKEIMSGIQSRLSDSKSIEDDDFVVHIPSDVNLPKLEGISDR
LMYVAALGWHVTKPRRGTLMFGCDYCARELAIRCGNGFDPIHNHERWCPRIDMDEHGEPSWQSDISIVLNTKNRVSHRYSGSSIFKEAYAARRL
IDSSLSTIISPNLL

**>KAF4260764.1[*Aspergillus_fumigatus*]**

MSYAVETKKRKFHRMLESLTKPSTVEPAAKKIPASPSPARERLPADLSIKKVRLSDKDESNFAAVGRSILKARASSKGSTISSATRPSFVPWDR
ERFLERLETFRRVDRWSPKPAVISEVEWAKRGWICTDVSRVSCVGGCGGSVVVKLPDELDELDGYALDKVQERKEVRDKLVEEYASLLVKGHGE
HCPWRNKGCDATIHRLPLTNTDTAIASLQKRYSNISKIADKLPAEDIIRTPESFNLDDVLKIWPRESFETGDGLRSTESQREQQDQSQDQESSE
EIRGESSQVAPAPSKPIDRAAFALAFFGWDAVSNETAGLVGCSACFRRLGLWMYKPKDNGDATVYDALDVANEHMEYCPWISGKAQSGTGRATE
KQAELRSGWELLVQALKVKHRRQVRSSTSMDTLRAVSETPSGDFPVVDEVSEEQKKASDREWWAKLRRMRQVLNVKSPHAKKSVVP

**>XP_006958443.1[*Wallemia_mellicola*]**

MNKRKIEDALTRLDKAFQSEDSVKAESSKRTKKPKIPAELEERLRWFTMTHNLQSRRSKSTDLLKKKEQSDLFSFHNFILRLQSFSLRTYTSKP
IELSPPAVALRGWQHDEAHRNRILCSRCKVGFIVDLSGSQANSYQKLIDTYVEAIESKHTSHCPWKYQQCASSTYRIHELSLPPNLAANVIAER
ARKIDSNLNFSIRVKHPLSNEQVQSLFNAMPEPKPSEDATILAIYGWEYKSTISSSSYLLTSELDVANVSVKPEKVFDVVREHRFYAPQLMPAN
EGSKNTGVEALLALITRRGRKKEIQESMIVNSVSVGSNTATVSLLI

**>XP_037145256.1[*Zygotorulaspora_mrakii*]**

MDSLSHGRLQSLFGMMLRDSEGDKGRNLRMILGTPETRRIIKKWKYRTKNCERATSRNAFLPEKLQKLQPLYYNVSEIVGKEYKGLQLYSLEQA
LERLNSIVVGTNCVLLSWDDKWINELSLASKGWKFHSLAGETPNTLVLLCSCGRCDTKIYLDLHSTENHNNDNVQNLINERYWEDIVTKSRPSQ
CPWVSNQFDLSDYHLKEANLIHDITRMQGHETVHFGTSGEITHNSTPLFSQEQLFSLSEFFKCADKRKLELMLRGFIPVEGCNDAVICVRCFRK
AFIGSITGSDYLNPHAPWCRYHDETKLPTMITSQLLSAKKTDDIKKRLSDLENYFESV

**>KAF2227358.1[*Elsinoe_ampelina*]**

MPEATATRKRRFYGLLDKLTQSSGPASKPSKTSAPTSRPTTSGGVTSNPRPSTPSSHPSISSPRPSTSSQRPSISAPRPIRPDRPTTLATPGAP
PPIPRIPHFTPWSHEDFLERLKTFAPVTSWFPKPEGVGEVEWAKRGWECVGNETVGCKGCKRRVVVDFTINGIGGDDEQAEDNVDESEEEAEFE
NTFEKALAEKYAEEIIKGHSESCPWRKTGCKDDIYRLPVVRTAWWQDTVRTSFTSVLEIKEDIENLRIHPISASPPADKLLLHIPPIFFNPNAT
PRNNSVLSNGSVDDPSVPSDEIMARALMIALTGWHATCESRTNLLVCDACFQRIGLWMYQSKPSTSPTTIPAGTSPLPTTPSTQSTSLTPAIKS
VSFGPSPPSSPSPTSSHPPLDLLASHREHCPYRSPSSQAATGDYVGFPAWRILWTTAARYADEQRRRSVAAKPGLGMGGLPGLGLGMKEGLSRE
EVARLDKERVNKLRRLKSALGIKLGVPRRGGGGREGEGSWV

**>Mext011234[*Medauroidea_extradentata*]**

MGVAVLLEGKVWTAEGRPVSRGAFVPIPSERKVWSQPYMGVEARKFNKTIVVTGPCLSVEKDDVLKKVEQNTSNPMSFRGFVKRMQTYNIVCWY
VHRRLALQLARRGWENVKDYVVKCCFCTAEVDLRNHISSQEEELLQLLYKRHASFCRWLTITSPEFFACVPHVQSSQGLKNTVATTATQLLELG
QELPKIPEVVLHKMKLEHETLELLISGLLKDVSVSEISISAVALTLCGWSPASTKGILCCCYCDRRLALRHWMSIQEFDEQRRLVLSDSNMKIC
SASYMQSSYADGAEEEDDDSQSPLSDQSIITDCSDEEEEVDEEEEENEEEFEEENEEGNEEMEVEEYELYEKHAEQDNECGNNEISDDGEEQSS
QTRNCQNEDNIENKYEDFQCRDEEMDENYDEINARALEENNFDDDDDEETEDEMPPNENGPSDEEVEEAEVSDESIMSDNEFKNENPKIFRGPN
KHTRYQSYVKRNKDSSNDLQAQYDSCGNEEDVESSEDDLLVENEVEDGDEEEEENEQNEGEEIGEEEDDEDDDDDDEEEEEDDEEEDDDDKDDE
DDEELPDDSKYHENNSDKVIELLSDSDEEIEQQPIEKIQSDSDEEIESQPVEQVQSVSSPTSAVVLETQTVVKENNVSNENKQSHCEKLCVEPK
TETENNDVDVREIIEKEIECDSDIALRSECENQEVKSLSLAEAIIMDDSKVRESQYDGNNSTQQEKVTLKRKRDCDEVETEQSLSIKRQQISLH
ESDKPLRVNENDMVITGGNPDCDKNNIEVEENTCSEGNDKNAEMVNSKALNENDCKTSAACERSVVVELVEHSKSQKECKLNGTLSLSKEKCEL
SVNSCEDSASAAVEGTNNLEQTCIENIEESVSSTAEMQTGVSETPAHSQNVVQKPLPKSEINSEVTVLKIETVCSIDTTQVIQKQKSRGKFVEN
NCTNVPLLLPEMPSENGLVIVQEEVCNVENTSSNIPLLLLENRSSDDDAGQEKGVICDKGGEPEVLATVQAVQDNSGEVEQAQILGEKAENKDA
QLEVPPLPGVQNGGGEVVENNSARGEETTKEDSVLSEVSCSLKGTSLPTEKGSENELSLQAGEDVKDTTVTNEAEDRTVGETTSLNNGNTSEEV
VQEYKSSEGNECSEELNPEECVHNNEESSTCDKSEADKKGEEGVTEMCESEALSVCAEKDGVMLEAMEVCEAGEQSSSRGVKRNRESDLEHSSP
EAKRPHQSTEVRRQPSLLDPLAEHRFWCLWAPSRGSASFGQSGFEIVFKEILDKILLQRREQDEMDIIGKLRSTQEQIMPL
